# Phylogenomic and mitogenomic data can accelerate inventorying of tropical beetles during the current biodiversity crisis

**DOI:** 10.1101/2021.07.15.452170

**Authors:** Michal Motyka, Dominik Kusy, Matej Bocek, Renata Bilkova, Ladislav Bocak

## Abstract

Conservation efforts must be evidence-based, so rapid and economically feasible methods should be used to quantify diversity and distribution patterns. We have attempted to overcome current impediments to the gathering of biodiversity data by using integrative phylogenomic and three mtDNA fragment analyses. As a model, we sequenced the Metriorrhynchini beetle fauna, sampled from ∼700 localities in three continents. The species-rich dataset included ∼6,500 terminals, >2,300 putative species, more than a half of them unknown to science. The phylogenomic backbone enabled the integrative delimitation of robustly defined natural units that will inform future research. Using constrained mtDNA analysis, we identified the spatial structure of α-diversity, very high species-level endemism, a biodiversity hotspot in New Guinea, and high phylogenetic diversity in the Sundaland. We suggest that ∼20 person months of focused field research and subsequent laboratory and bioinformatic workflow steps would substantially accelerate the inventorying of any hyperdiverse tropical group with several thousand species. The outcome would be a scaffold for the incorporation of further data. The database of sequences could set a benchmark for the spatiotemporal evaluation of biodiversity, would support evidence-based conservation planning, and would provide a robust framework for systematic, biogeographic, and evolutionary studies.

## Introduction

The number of known insects surpasses that of all other terrestrial groups (Mora *et al*., 2011), and we need much more detailed information to fully understand their diversity. Currently, the available biodiversity data are far from complete, and the majority of insect species remain undescribed (Novotny *et al*., 2006; Sriwathsan *et al*., 2019). In addition, robust phylogenetic hypotheses are lacking for most lineages, and the genera and tribes are often artificial assemblages which are not relevant to evolutionary and biodiversity research. Therefore, we need to gather new information in order to advance our understanding of evolutionary and genetic relationships, and to build a phylogenetic scaffold for comprehensive taxonomic, biogeographic, and evolutionary studies that would be indispensable for biodiversity management.

Descriptive, morphology-based insect systematics is not keeping pace with the rapid loss and degradation of natural habitats (Theng *et al*., 2020), and with the ongoing decline in insect abundance as a result of human activities and climate change (van Klink *et al*., 2020). To accelerate the cataloguing of biodiversity, it is vital to use innovative methods (Riedel *et al*., 2013; Sriwathsan *et al*., 2019; Yeo *et al*., 2020; Sharkey *et al*., 2021). DNA data are indisputably a valuable source for modern biodiversity research (Tautz *et al*., 2003; Hajibabaei *et al*. 2007). There are two principal sources of data: voucher-based DNA sequences typically produced by systematists (Riedel *et al*., 2013; Yeo *et al*., 2020;, Sharkey *et al*., 2021), and DNA sequences produced by an ecosystem-based sequencing that does not associate individual samples with Linnean names (Andújar *et al*., 2015; Sriwathsan *et al*., 2019). It is the responsibility of systematic biologists to assemble the natural system, i.e., we need to reliably delimit genus-and tribe-level taxa, to make their ecological and distribution attributes informative. Then, a robust and stable natural classification will significantly facilitate detailed research into the spatial and temporal distribution of biodiversity (Morrison *et al*., 2009; Thomson *et al*., 2018). As an ultimate goal we should attempt to construct a complete tree of life, or at least its backbone, which is invaluable in aiding the selection of groups for more detailed analyses (Chesters, 2017; McKenna *et al*., 2019). With a well-defined high-level classification, it is paramount to exploit all accessible data. We assume that voucher-based molecular phylogenies provide much-needed tools to researchers working on site-based biodiversity assessments (Andújar *et al*., 2015; Sriwathsan *et al*., 2019) and that, in turn, the data produced by environmental and ecosystem-focused sequencing contribute to building the tree-of-life (Arribas *et al*., 2016; Bocak *et al*., 2016).

We have used hyperdiverse tropical metriorrhynchine beetles (Coleoptera, Lycidae, Metriorrhynchini) as our model. This net-winged beetle tribe contains >1,500 recognised species, mostly found in the Old-World tropics (Fig. 1A), and their classification is complicated by the complex taxonomic history (Bocak *et al*., 2020). The phenetic plasticity of Metriorrhynchini is relatively high (Fig. 1B–D), but many distant species resemble each other due to convergent selection in Mullerian rings (Bocek *et al*., 2019; Motyka *et al*., 2020, 2021). Therefore, unrelated taxa were often assumed to be closely related due to misleading morphological similarities. Although there are over 40 genera in the tribe, three-quarters of the species have been described in five ambiguously defined genera (*Xylobanus, Cautires, Trichalus, Metriorrhynchus*, and *Cladophorus*). Sometimes a single genus contains species from different subtribes (Bocak *et al*., 2020). In this respect, the Metriorrhynchini is a typical species-rich tropical insect group without well-founded classification and the paucity and inaccuracy of available data (Letsch *et al*., 2020). As a result, unlike vertebrates, these poorly known insect groups have not been considered for use in large-scale, integrative projects and data metanalyses (Myers *et al*., 2000; Holt *et al*., 2013) and have contributed little to our understanding of global biodiversity patterns.

**Figure 1.**
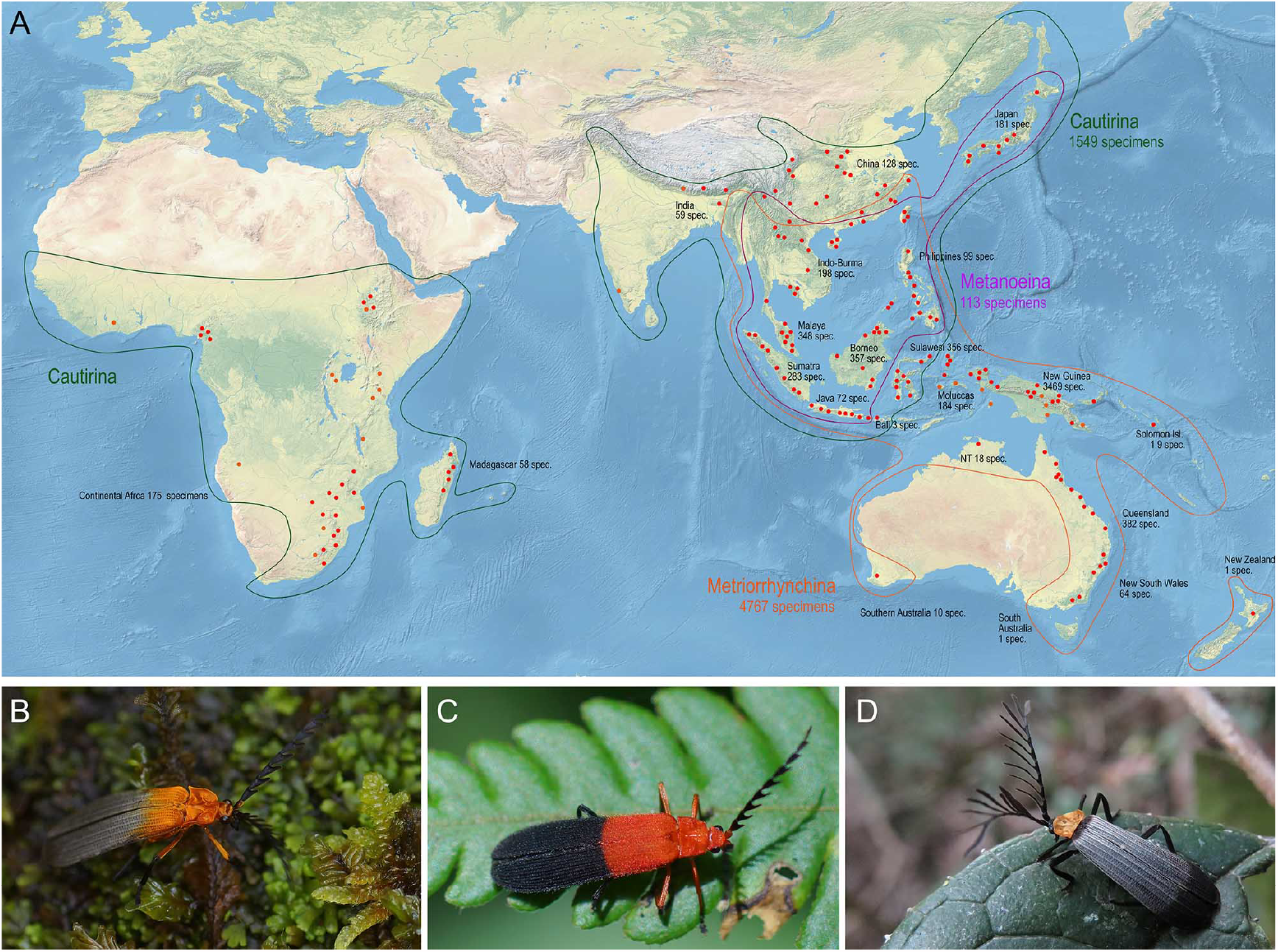
A – Distribution of Metriorrhynchini with major sampled localities designated by red dots. The numbers of analysed specimens from individual regions are shown for regions and subtribes. B–D – General appearance of Metriorrhynchini.

The principal objective of this study is to demonstrate how biodiversity information for a hyperdiverse tropical group can be rapidly expanded via targeted field research and large-scale sequencing. Our investigation comprised four distinct steps, aiming for a DNA-based evaluation of diversity and evolution. First, we assembled material from several hundred localities on three continents (Fig. 1, Tab. 1). Second, as hyperdiverse groups are difficult to tackle and the current classification is unreliable, we attempted to compartmentalise diversity using phylogenomics. We then produced a tree, using all available data, to estimate species limits, intraspecific genetic variability, and species ranges. Finally, the tree was pruned and used to estimate shallow phylogenetic relationships, total and regional α-diversity, and endemicity, and to define generic ranges and continental-scale range shifts. Our information and phylogenetic hypotheses can be a resource for higher-level phylogenetics, population genetics, phylogeographic studies, and biodiversity estimation. At the same time, we want to show how limited our taxonomical knowledge is and how this lack is hindering biodiversity research and management (Thompson *et al*., 2018).

**Table 1.** The numbers of sampled localities per region.

| Area | Localities | Area | Localities |
| --- | --- | --- | --- |
| Australian region | 347 | Palaearctic region | 79 |
| Australia | 118 | China | 51 |
| New Zealand | 1 | Japan | 28 |
| New Guinea | 175 | Russian Far East | 0 |
| New Britain & Ireland | 0 |  |  |
| Solomon Isl. | 4 |  |  |
| Moluccas | 15 | Oriental region | 206 |
| Sulawesi | 34 | Himalayas | 0 |
|  |  | S India & Ceylon | 3 |
| Afrotropical Region | 64 | E. India & Burma | 12 |
| West Africa | 1 | E. Indo-Burma | 44 |
| Guinean Gulf | 9 | Malay Peninsula | 57 |
| Congo Basin | 2 | Sumatra | 23 |
| Sahel | 0 | Borneo | 19 |
| Ethiopia & Somalia | 6 | Java & Bali | 15 |
| East African Highl. | 10 | Philippines | 33 |
| South Africa | 25 |  |  |
| Madagascar | 11 | Total | 696 |

## Results

### Sampling of the Metriorrhynchini range

In total, we monitored almost 800 localities, 696 of them with occurrences of the Metriorrhynchini (Tabs. 1, S1). The distribution of sampling sites was partly biased due to the large extent of the Metriorrhynchini range, limited time and funds, different goals of various expeditions, and logistic problems (inaccessible regions, legal obstacles). The densest sampling is available from the Sundaland and New Guinea, while India and the Afrotropical region are under-sampled.

### Assembly of the phylogenomic tree

The phylogenomic dataset contained 35 Metriorrhynchini terminals, seven outgroups, and ∼4,200 orthologs (1.9–5.7 × 10^6^ aligned positions; Tab. S5). The tree shown in Fig. 2A was produced using maximum likelihood (ML) analyses, whereas the coalescent method produced the topology shown in Fig. 2B; additional trees are shown in Figs. S1–S8. For details on the data sets’ characteristics see Figs. S9–S12. Phylogenomic analyses resolved three subtribes (Metanoeina (Metriorrhynchina, Cautirina)), and five clades were regularly recovered within the Metriorrhynchina, i.e., the procautirines, leptotrichalines, trichalines, porrostomines, and cladophorines. Different settings (see Methods) produced slightly different topologies and shifted the positions of the leptotrichalines and procautirines (Fig. 3D). However, the monophyly of major subclades was not affected. The FcLM analysis favoured a deeper position for the leptotrichaline clade (61.2%; Figs. 2C, S13). The position of the remaining terminals was stable across all analyses.

**Figure 2.**
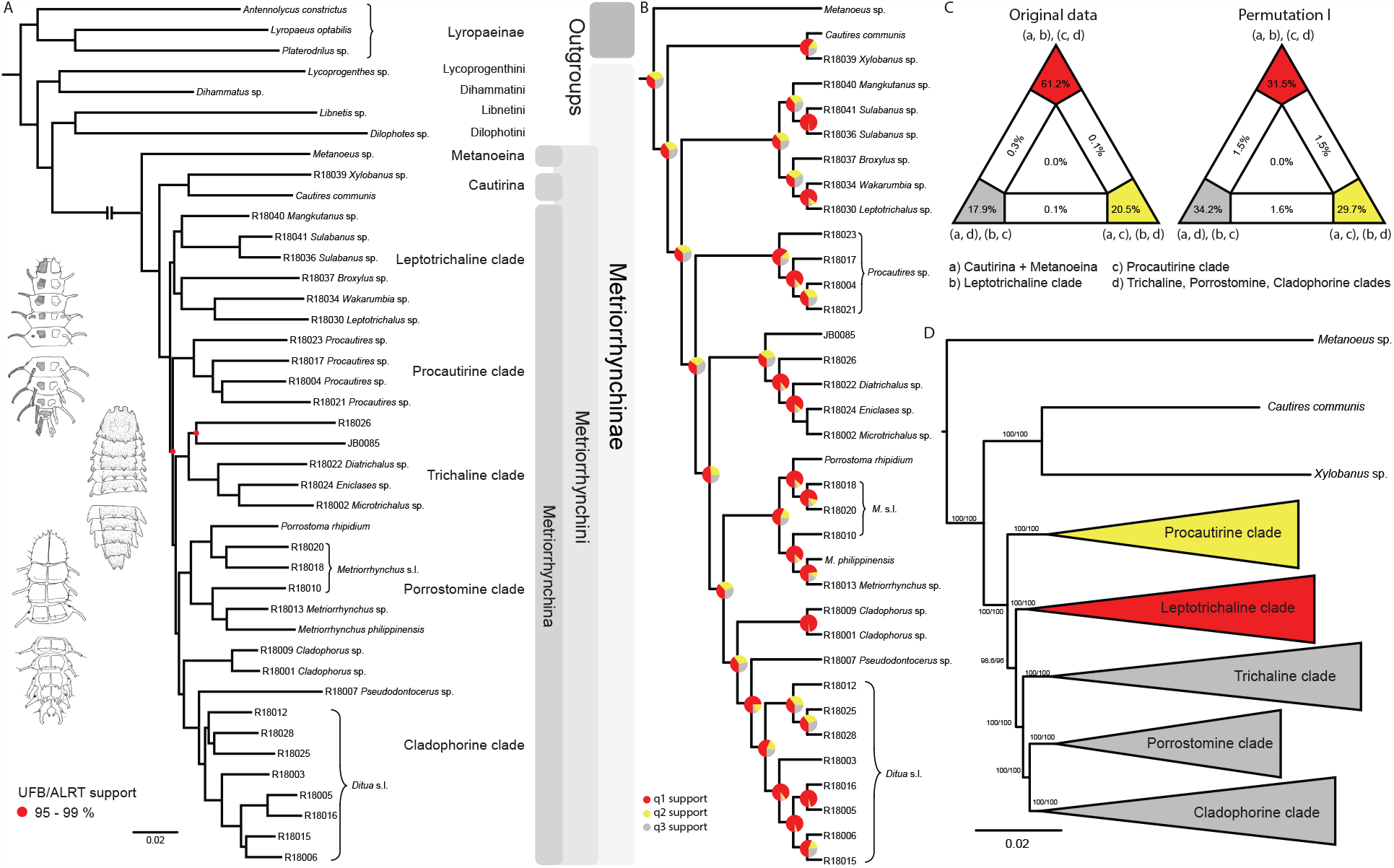
A – Phylogenetic relationships of Metriorrhynchinae based on the ML analyses of the concatenated amino-acid sequence data of supermatrix F-1490-AA-Bacoca-decisive. Unmarked branches are supported by 100/100 UFB/alrt; red circles depict lower phylogenetic branch support. B – Phylogenetic relationships of Metriorrhynchini recovered by the coalescent phylogenetic analysis with ASTRAL when analysing the full set of gene trees (4109 gene trees inferred at the nucleotide level). Pie charts on branches show ASTRAL quartet support (quartet-based frequencies of alternative quadripartition topologies around a given internode). Outgroups taxa are not shown. C – Results of FcLM analyses for selected phylogenetic hypotheses applied at the amino-acid sequence level (supermatrix F). D – Alternative phylogenetic relationships of Metriorrhynchinae based on the ML analyses of the concatenated amino-acid sequence data of supermatrix A-4109-AA. Numbers depict phylogenetic branch support values based on 5000 ultrafast bootstrap replicates.

**Figure 3.**
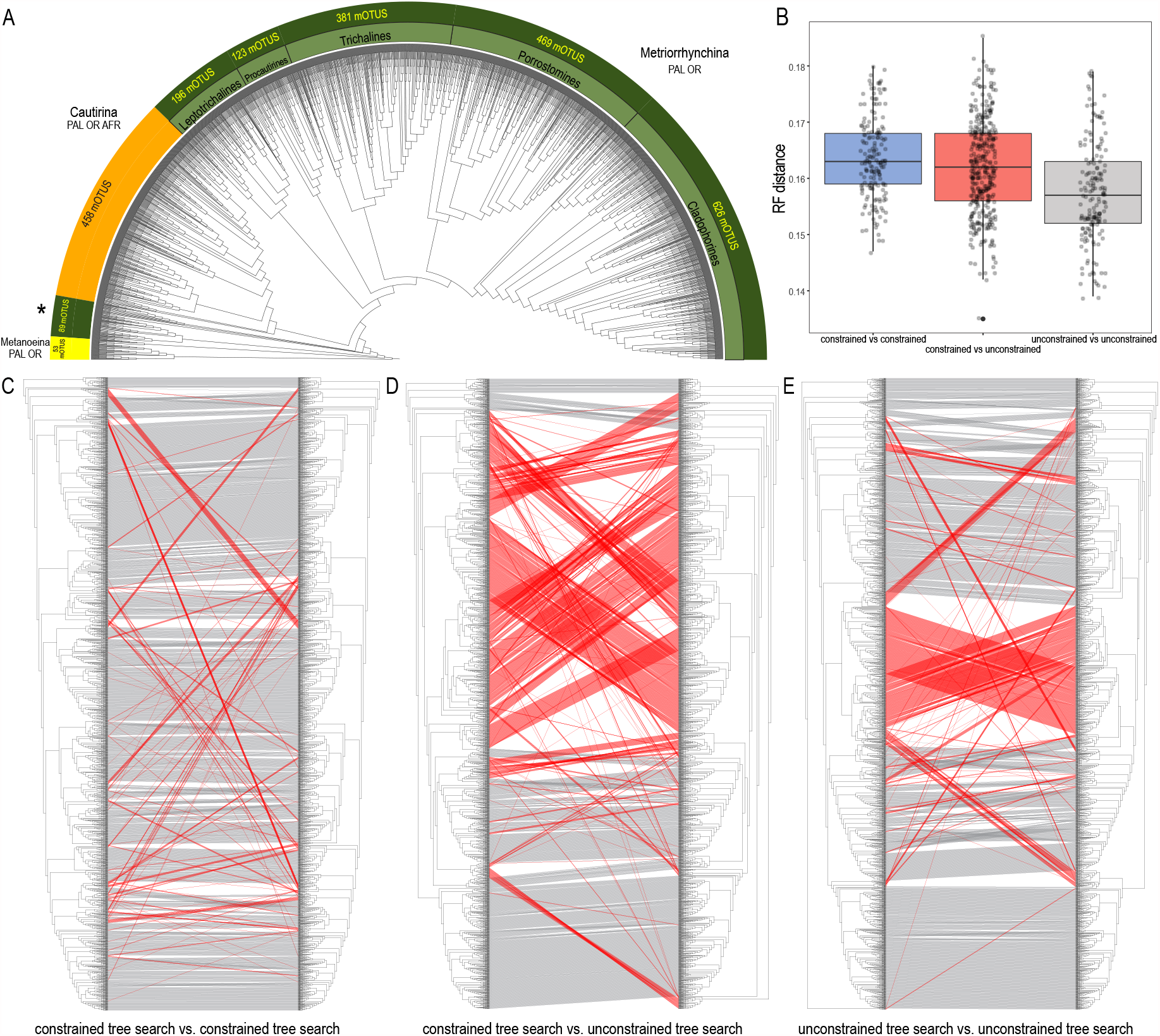
A – Relationships of 2,345 Metriorrhynchini species recovered by the constrained analysis of the pruned dataset (The full resolution tree is shown in Fig. S15 along with a tree recovered from the analysis of a complete dataset of 6,429 terminals in Fig. S14), asterisk designates a grade of Metriorrhynchina-like taxa found in a position in conflict with their morphology; B – A chart of Robinson-Foulds distances among topologies inferred by repeated runs of the constrained and unconstrained analyses; C – A comparison of the results obtained by two runs of the constrained analysis; D – A comparison of trees inferred with/without the phylogenomic backbone; E – A comparison of results obtained by two runs of the unconstrained analysis. The red lines designate terminals with conflicting positions in compared trees.

### Constrained mitogenomics

The mtDNA database contained >11,500 mtDNA fragments (5,935 *cox1*, 2,381 *rrnL*, and 3,205 *nad5*) representing 6,429 terminals (2,930 aligned positions). Using these data, we inferred additional trees using the constrained positions of 35 terminals whose relationships were determined through phylogenomic analyses, and the free positions of the other ∼6,400 terminals (Fig. S14). The units based on uncorrected pairwise distances represent molecular operational taxonomic units (mOTUs), considered to be putative species, or ‘species’ for short. We identified 37 mOTUs in the Metanoeina clade and 456 mOTUs in Cautirina. The major Metriorrhynchina clade (1,763 mOTUs) included procautirines, leptotrichalines, trichalines, porrostomines, and cladophorines. In addition, we identified several deeply rooted lineages, the kassemiines, and another five small clades (89 mOTUs in total; Fig. S15), each of which comprised a limited number of species. As phylogenomic data for these terminals are still lacking, their terminal positions were determined based only on mtDNA data. The number of mOTUs does not include ∼50 mOTUs for which *cox1* was unavailable.

### Pruned mitogenomic tree with and without constraints

The dataset was subsequently pruned to a single terminal per mOTU (see below) and was analysed both with and without topological constraints (Figs. 3A, S15, S16). Repeated runs with different starting seeds identified terminals with unstable positions (Figs. 3A–C). The major clades were generally stable, whereas small, deeply rooted clades were prone to ‘wandering’ around the tree, as were distinct singletons. The trees that resulted from each of the seed-specific 19 ML runs differed slightly; tree similarity was thus evaluated using the Robinson-Foulds index, with values ranging from 0.180 (most similar) to 0.147 (most distant; Tab. S7).

### Tree congruence

The degree of incongruence between selected topologies is shown in Fig. 3C–D. The unconstrained analysis of mitochondrial data yielded a topology with a high number of terminals that were recovered in positions incongruent with their morphology (Fig. 3D, E, S16). The same dataset, when analysed using the constrained position of 35 terminals (based on their relative relationships inferred by prior phylogenomic analyses), produced a topology with a much lower proportion of terminals in dubious positions (Fig. 3C, S15). The composition of the constituent clades is shown in Tab. S8; individual clades are characterised in the Supplementary Text.

### Alpha-diversity

To investigate the total and regional α-diversity of the Metriorrhynchini, we analysed a dataset comprising 5,935 of the 6,429 terminals for which the *cox1* mtDNA fragment was available (Fig. 3A; Tab. S3). We identified 2,345 mOTUs using a 2% distance threshold. We disregarded the presence of ∼50 mOTUs (494 terminals) for which *cox1* was missing. The number of mOTUs based on the *cox1* analysis varied by threshold. For the Metriorrhynchini, we identified 1,848 and 2,356 mOTUs using thresholds of 5% and 2%, respectively (Fig. S17).

Following an extensive literature review, we updated species lists for the Cautirina (641 spp.), Metanoeina (38 spp.), and Metriorrhynchina (895 spp.; Bocak *et al*., 2020). By analysing DNA data, we identified 456 spp. of Cautirina, 37 spp. of Metanoeina, and 1,852 spp. of Metriorrhynchina. The numbers of species per subregion, along with the estimated ratios between formally described and estimated α-diversity, are shown in Tab. 1. Even using a threshold of 5%, the number of putative species surpasses the number of species reported in the literature.

We observed very high species turnover, and no species has been recorded in two landmasses separated by a deep-sea (>200 m). Similarly, the faunas of Sulawesi and of the surrounding large islands do not overlap. Only thirteen species were distributed across two landmasses separated by an inundated shelf (sea depth <100 m). Eleven species were distributed in two or more islands of the Great Sundas and two species were found in both New Guinea and Australia. The centres of α-diversity of the Metriorrhynchini are New Guinea (1,434 spp.) and the seasonally to perennially humid areas of the Sundaland (261 spp.). The results suggest substantial modifications to the generic limits and ranges for numerous taxa that had been previously delimited (Fig. S18).

## Discussion

In the context of the present loss of biodiversity (Sodhi *et al*., 2004; Hallmann *et al*., 2017; Theng *et al*., 2020), large-scale genomic resources are urgently needed for biodiversity assessment and conservation (Hajibabaei *et al*., 2007; Krehenwinkel *et al*., 2019). Molecular data cannot replace morphology-based taxonomy (Fig. 3C–D; Thomson *et al*., 2018), but the analyses of our dataset complement and facilitate traditional biodiversity research in several directions. Our first step is to compartmentalise hyperdiverse Metriorrhynchini into manageable natural units (Fig. 2). The densely sampled phylogeny identifies tribal and generic limits. It provides a useful foundation for detailed taxonomic research through the identification of weak areas in earlier classifications and points out the clades with undescribed diversity (Figs. 3, S18). Furthermore, the analyses of species-rich datasets identify the areas with high α-diversity as one of the critical conservation value parameters (Tab. 2; Baselga, 2010; Srivathsan *et al*., 2019). Traditional taxonomic research costs time and money, and the number of newly described species is relatively low if we consider the enormous diversity of tropical insects (Novotny *et al*., 2006; Sangster & Luksenburg, 2014). Therefore, we use DNA-based units as a provisionary descriptor of α-diversity (Hebert *et al*., 2003; Monaghan *et al*., 2009), and subsequently as a source for integrative taxonomy (Figs. S14–S16; Srivathsan *et al*., 2019). The presented large-scale monitoring project provides information on relationships (Figs. 2, 3), genetic divergence (Figs. S14–S16), turnover (Tab. 2), the extent of generic and species ranges (Fig. S14–S16, S18), and on evolutionary phenomena that are usually studied using a few model organisms (Fig. 4). Using phylogenomics and voucher-based sequencing, we show that taxonomic literature has provided insufficient and sometimes erroneous information, even after the formal consolidation of scattered descriptions (Bocak *et al*., 2020). We show that a taxon-focused continental scale project can effectively assemble comprehensive data for diversity of tropical insects.

**Table 2.** The number of described species and identified mOTUs (molecular operational taxonomic units at 2% difference) per region. * – some species are shared by two regions.

| Region | Metriorrhynchina | Cautirina | Metanoestina | Metriorrhynchini | Seq./Descr. |
| --- | --- | --- | --- | --- | --- |
| **Australian region** | **639/1608** |  |  | **639/1608** | **2.52** |
| Australia | 196/167* |  |  | 196/167 | 0.85 |
| New Guinea | 423/1434* |  |  | 423/1434 | 3.39 |
| Solomon | 21*/9 |  |  | 21/9 | 0.43 |
| **Wallacea** | **162/174** | **14/10** |  | **176/184** | **1.05** |
| **Philippines** | **51/18** | **45/12** | **8/3** | **104/33** | **0.32** |
| **Continental Asia** | **43/52** | **331/330** | **30/34** | **404/416** | **1.03** |
| Sundaland | 36/44 | 201/184 | 24/19 | 261/247 | 0.95 |
| Indo-Burma | 6/7 | 62/52 | 3/4 | 74/63 | 0.85 |
| China, Japan | 1/1 | 53/75 | 1/11 | 55/87 | 1.58 |
| India |  | 35/19 | 2/0 | 37/19 | 0.51 |
| **Afrotropical region** |  | **231/104** |  | **231/104** | **0.46** |
| Africa |  | 178/74 |  | 178/74 | 0.42 |
| Madagascar |  | 53/30 |  | 53/30 | 0.57 |
| **Total** | **895/1852** | **641/456** | **38/37** | **1574/2345** | **1.50** |

**Figure 4.**
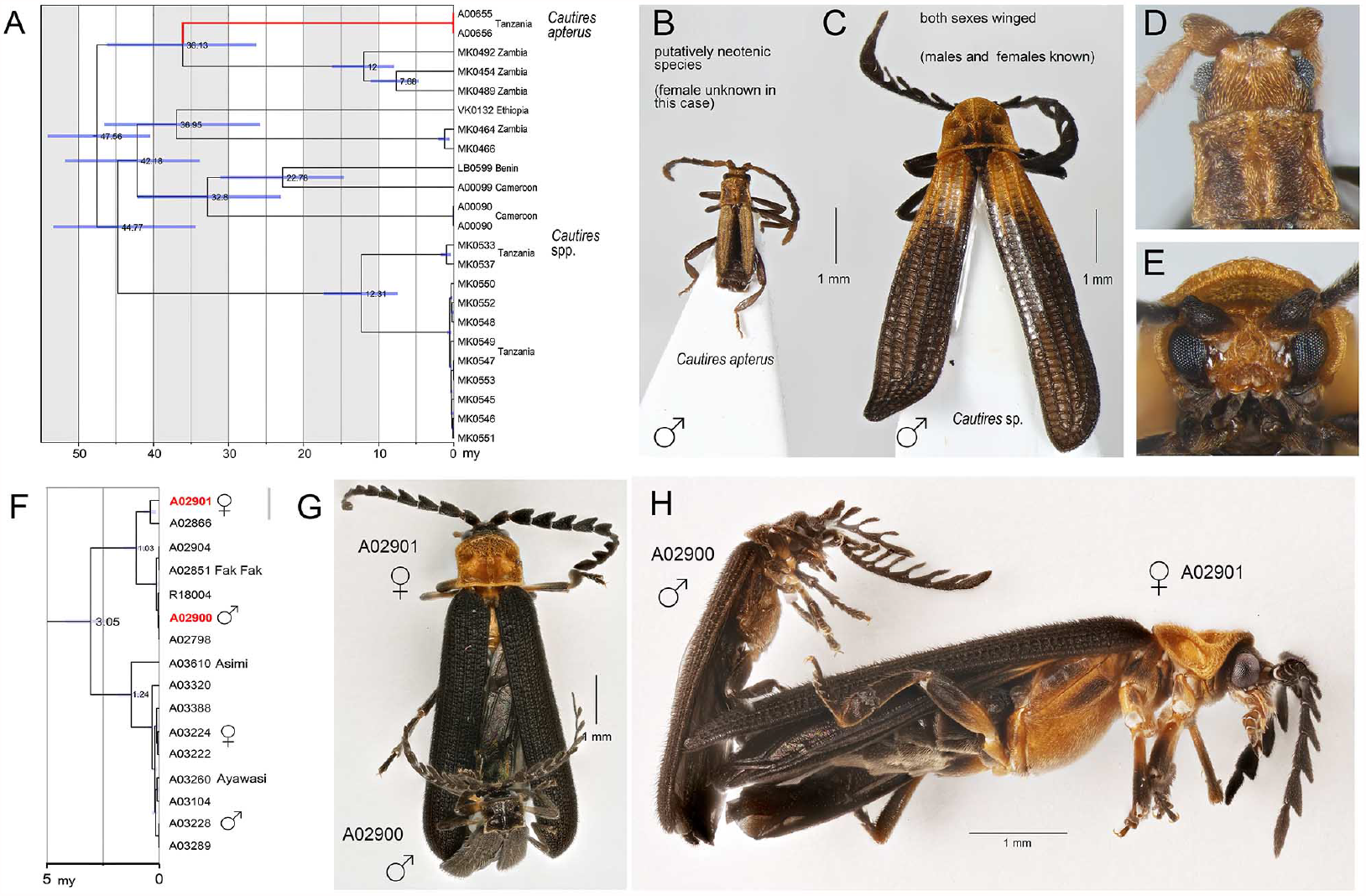
Identification of sexual dimorphism by large scale biodiversity inventory. A – Relationships of lineages with modified ontogeny, the dated tree; B, D – general appearance and head of Cautires apterus, a putative neotenic species; C, E – ditto of the close relative with both sexes winged. Mimetic sexual dimorphism identified during diversity survey. F – the dated tree, red coloured terminal labels designate the individuals shown in G and H; G – dorsal view of individuals in copula; H – ditto, lateral view. Except of collecting individuals in copula, DNA-based assessment of relationships is the only option as the species are sexually dimorphic and no morphological traits indicate their conspecifity.

**Figure 5.**
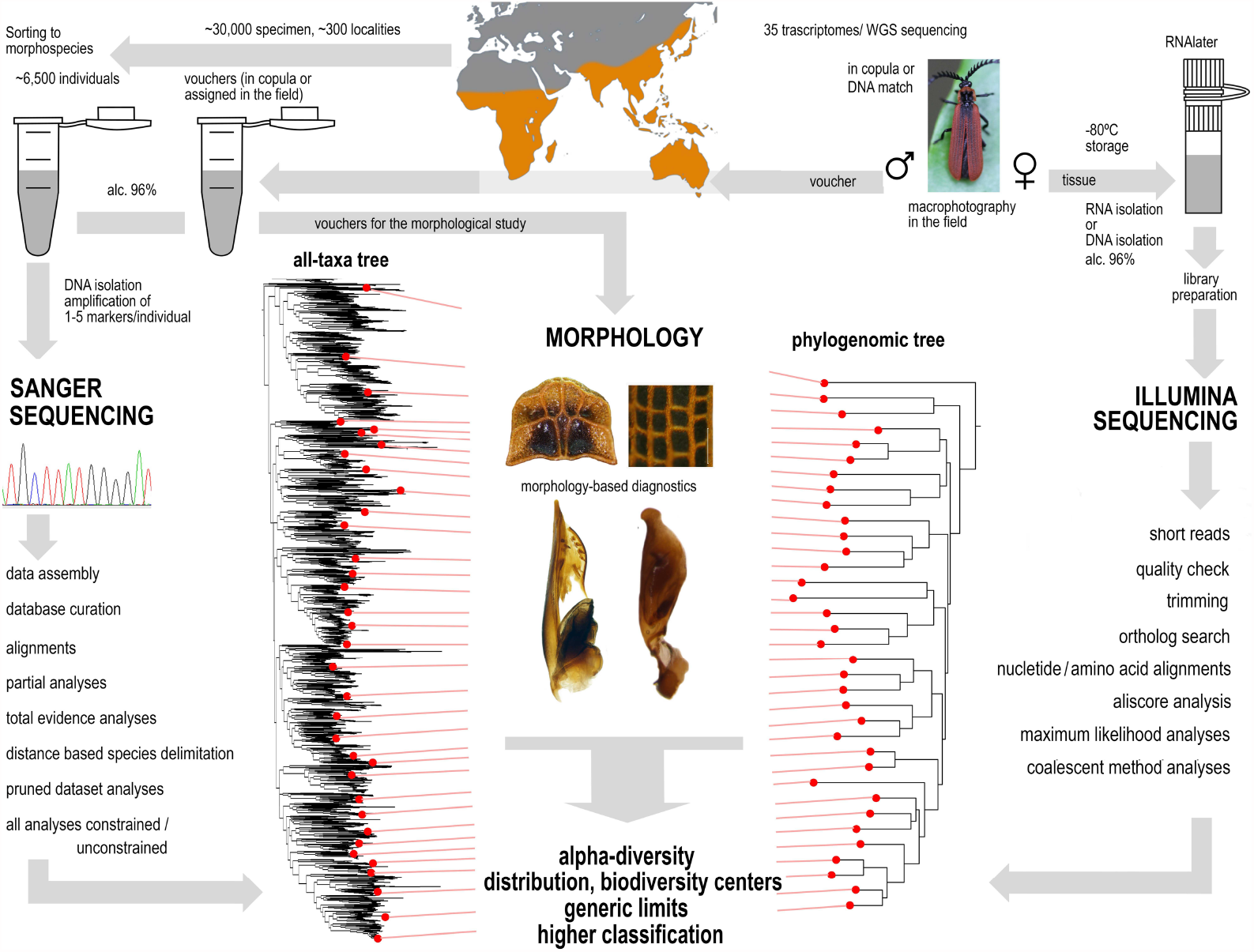
A sequence of applied methods from sampling to hypotheses.

### Continent-wide taxon-specific monitoring of biodiversity: feasibility and impediments

Tissue and DNA archives have become critical in the assessment of biodiversity status (Hajibabaei *et al*., 2007; Blom, 2021). Although museomics is a potentially valuable source of data (Gauthier *et al*., 2020), in our case, museum collections are insufficient for filling data gaps due to the scarcity of material. For example, the Metriorrhynchini collection deposited in the Natural History Museum in London contains <3,000 specimens, whereas there are ∼6,500 terminals in our dataset. At the beginning of our study, we faced critical absence of primary data. Therefore, we conducted intensive field research to obtain samples for a realistic assessment of the extant Metriorrhynchini diversity. We processed samples from our expeditions (most of which were focused on a range of topics over two decades between 2001 and 2019) and samples obtained through extensive collaboration with other researchers, both local and visiting, and with local naturalists whose contribution has increased with the growing number of citizen science projects (Jaskula *et al*., 2021; MacPhail, & Colla, 2020). In such a way we assembled a Metriorrhynchini tissue collection from almost 700 localities in three continents (Tab. 1, Fig. 1). For several reasons our sampling is partly biased. We noted the serious loss of natural habitat in many regions. Previously described species were often collected in vicinity of seaports, but the lowland ecosystems are rapidly disappearing due to human exploitation. Therefore, type localities of many described species could not be sampled during recent expeditions and species known from museum collections are missing in our DNA dataset (Jiruskova *et al*., 2019). The habitat loss in South East Asia also affects other animal groups, and lowland primary forests are seriously endangered in the whole region (Sodhi *et al*., 2004, Theng *et al*., 2020). Further sampling bias is a consequence of the unsafe conditions and logistic problems in large areas of West Africa, Sahel, and the Congo Basin (Fig. 1A), where net-winged beetles have not been systematically studied since the 1930s. Additional data gaps are caused by strict biodiversity research restrictions (Prathapan *et al*., 2018; Laird *et al*., 2020). Regardless of these limitations, we believe that the assembled dataset is a foundation for a robust classification framework and a soundly based assessment of biodiversity. Our results show the importance of field research for biodiversity studies and systematics (Basset & Lamarre, 2019).

### Phylogenetic relationships: a scaffold for targeted research

Unresolved taxonomy is a common reason for the exclusion of specific groups from biodiversity research projects and this omission has an effect on conservation policies (Gutierrez & Helgen, 2013). The current phylogenomic and mitogenomic phylogenetic hypotheses (Figs. 2, 3; S14–S16) supersede the morphology-based topologies (Bocak, 2002). The phylogenomic analysis incorporates a large amount of information, and we favour this method over morphological traits and short DNA sequences, both of which contain uncertainties (McKenna *et al*., 2019). Phylogenomics has resolved subtribe relationships and their internal structures. The analysed 35 transcriptomes and low-coverage genomes were sufficient to identify five major Metriorrhynchina clades (with 100–600 putative species each) and also to identify the limits of genera, which can be tested using traditional taxonomic methods (Figs. 2, 3A, S14–S16).

The sampling strategy is critical for building a phylogenomic backbone, and our goal was to cover as many deep lineages as possible. Therefore, we sequenced RNAlater preserved tissues and conspecific vouchers prior to assigning tissue samples for transcriptomic analyses. In this way, two rounds of sequencing provided us with critical information based on evenly distributed anchor taxa. In the next step, we re-analysed the short-fragment dataset (Tab. S3) using constrained positions for taxa whose relationships had already been recovered through phylogenomics (Figs. 2, 3). A stabilised phylogenomic backbone is critical for inferring a robust topology because the analyses of short mtDNA fragments are sensitive, even to the application of starting seeds, and they often produce topologies incongruent with morphological traits (Figs. 3B, E). Only several small lineages have remained unanchored by genomic data, owing to a lack of properly fixed samples (Figs. S15, S16). For example, four small clades are much more deeply rooted than their morphology suggests (Figs. 3A, S15) and additional data are needed to place them in a phylogenetic context.

Our approach yielded a phylogeny with 6,429 terminals and 2,345 mOTUs, and this provides the basis for the approximation of α-diversity (Figs. 3, S15, S16). Concerning the extent of diversity, phylogenomic and mitochondrial data must be simultaneously analysed to provide a strong foundation for subsequent investigations (Fig. 3C). Phylogenomics cannot deal with thousands of species, and mitogenomic data are insufficient for the construction of robust relationships. The final steps are morphological validation (see Supplementary Text) and, in the future, formal descriptions of biodiversity using the Linnean classification. In such a way, the results of phylogenomic and mitogenomic inventory should be incorporated in the Linnean classification (Godfray & Knapp, 2004).

### Alpha-diversity: literature data and reality

Here, we deal with a tropical beetle tribe with >1,500 described species, and our results indicate that regionally up to 3.4 times that number of species remain undescribed (Tab. 2). When analysing *cox1* mtDNA, we identified 2,345 mOTUs based on an arbitrary distance threshold of 2 % uncorrected pairwise distance (Hebert *et al*., 2003). The application of the threshold is a compromise between estimation accuracy, speed, and sequencing costs, taking into account the feasibility of inventorying a hyperdiverse group within a limited timeframe (Dupuis *et al*., 2012; Eberle *et al*., 2020). We had previously explored a subset of Metriorrhynchini to estimate the congruence between mtDNA and morphology, and between nextRAD-and mtDNA-based species limits. We found similar numbers of species regardless of the approach that was applied (e.g., Bocek *et al*., 2019, Jiruskova et al. 2019). Additionally, the slope representing the relationship between the number of mOTUs and distance thresholds was gradual (Fig. S17) due to a high number of genetically distant, indisputably distinct lineages (Figs. S15, S16). Therefore, we assume that our estimates of α-diversity are realistic, although future taxonomic revisions are needed for validation.

Our approach provides information about the diversity of the internal lineages. Metriorrhynchina is by far the most diverse group, within which the cladophorines comprise the largest clade (626 mOTUs, *Ditua* historically has included 2 spp. now ∼300 spp.). The porrostomine clade is the next diverse group (346 spp.) and contains the speciose *Porrostoma* (156 spp.) and a paraphyletic series of lineages whose species have conventionally been placed in *Metriorrhynchus*. The differences between previously published data and our results are substantial (Figs. S15, S16).

The numbers of mOTUs must be interpreted in the context of the sampling activity in each region. We identified only 104 mOTUs from the Afrotropical region, mainly due to the limited number of collecting trips (5 person months; 64 localities) and the inaccessibility of some areas. Despite intensive field research (4 person months, 33 localities), we collected from the Philippines less than one third of the species described. Our collection activities in the Philippines were hindered by substantial loss of natural habitats, and this is soon expected to be the case in other regions (Sodhi *et al*., 2004). The number of species known from the Sundaland (16 person months, 114 localities) was approximately equal to the number of sequenced mOTUs. Many regions remain unsampled and species ranges are small (Jiruskova *et al*., 2019), so this number will increase in the future. The proportion of new species was regionally ∼70% if DNA data and morphology were considered in detailed taxonomic studies (e.g., Jiruskova *et al*., 2019). While these regions house numerous unknown species, we found New Guinea to be exceptionally diverse, with 3.4 times the number of species reported in the literature (1,434 mOTUs; 7 person months; 175 localities). Despite the relatively large number of sampled localities, many areas of New Guinea remain unexplored (Fig. 1A), and additional species were added to the dataset with each batch of sequenced samples from other localities.

We observed a high turnover between regions, and few species had ranges which included landmasses separated by shallow seas (2 spp. Queensland / New Guinea, 11 spp. Sundaland islands; Fig. S14). Poorly dispersing lycids generally have very small ranges, except for the few genera that visit flowers and fly in open areas (Kusy *et al*., 2021). A similar small-scale turnover has recently been reported along altitudinal gradients (Bocek *et al*., 2019; Motyka *et al*., 2020, 2021). A high turnover indicates a large proportion of hidden diversity, especially in tropical mountains (Merckx *et al*., 2015; Mastretta-Yanes *et al*., 2018). Mountain fauna is especially vulnerable to climate change and its inventorying is urgently needed.

The Metriorrhynchini has recently received considerable attention in taxonomic studies, and 302 species have been described by several authors over the past three decades, making a total of 1,574 formally described species (Tab. 1, Bocak et al. 2020; Supplementary Text and References). Although the recent 24% increase in described diversity appears substantial, the distance-based analysis indicates the presence of 2,345 mOTUs (Fig. S17). An additional ∼50 putative species (494 terminals) were identified, but this identification was only based on divergent morphology because of the absence of *cox1*. We assume that our sampling represents only a subset of all known species (<50%). It means that the dataset contains 1,000–1,500 undescribed species. At the current rate, formal morphological descriptions of an additional 1,000+ species would take 100 years. This is a very long time in the context of the ongoing deforestation and fragmentation of natural habitats, and currently undocumented diversity might be lost long before it can be catalogued (Brooks *et al*. 2002; Sodhi *et al*. 2004; Ceballos *et al*., 2015; Theng *et al*., 2020). The rapid DNA-based inventory is an effective shortcut for obtaining basic information on the true diversity of tropical beetles and for setting a benchmark for future biodiversity re-evaluations.

The results reveal major biodiversity hotspots in New Guinea and the Sundaland. Tropical rainforests currently cover most of New Guinea, a tectonically young island that has not been considered a biodiversity hotspot for vertebrates (Myers *et al*., 2000, Hall, 2011, Toussaint et al. 2014). In the case of net-winged beetles, we show that the New Guinean fauna is phylogenetically diverse, spatially heterogeneous, and extremely rich as regards both the number of species and the endemic genera (Tab. 1). Additionally, the large clades of New Guinean species indicate that the diversification of major lineages preceded the uplift of the islands, and possibly started on the norther margin of the Australian craton and adjacent islands. Southeast Asia is a centre of phylogenetic diversity at the tribal level; its fauna contains all principal lineages and the highest diversity of Cautirina but is smaller than those of New Guinea. The Afrotropical and Palearctic regions represent only recently populated low-diversity outposts.

### Impact of biodiversity inventorying on biogeographical and evolutionary research

Detailed data on Metriorrhynchini diversity indicate low dispersal propensity and this makes Metriorrhynchini a promising model for biogeographic studies. Our densely sampled phylogeny did not find any long-distance dispersal events, in contrast to many studies of flying beetles (Balke *et al*., 2009; Jordal *et al*., 2015). Most recovered overseas dispersal events are limited to distances of less than 100 km and are commonly accompanied by speciation (Fig. S15, S16). The high proportion of erroneous placement of many taxa (Fig. S18; Bocak *et al*., 2020) renders the distribution data cited in previous literature unsuitable for phylogeographic investigations, and revision of the classification is important in order to understand the true distribution of individual taxa. The original and revised ranges of selected genera are compared in Fig. S18 as examples.

Intensive biodiversity research has the potential to fill knowledge gaps concerning evolutionary phenomena that are mainly studied using a small number of model species, and the research can identify the unique attributes of other potential models. We document the contribution of a large-scale biodiversity inventory to evolutionary studies with two examples.

Net-winged beetles include several lineages in which females have lost the ability to completely metamorphose (Bocak *et al*., 2008; McMahon & Hayward, 2016). If a putative neotenic species is discovered, a comprehensive reference database of the group may identify its closest relatives. We used our data to place the East African *Cautires apterus* in a phylogenetic context, and the results indicated that it may be the youngest neotenic taxon of all net-winged beetles (36.1 my, Fig. 4).

Our extensive DNA database of metriorrhynchine diversity may also play an important role in the study of the evolution of mimicry. Our inventory identified an extreme and previously unknown aposematic dimorphism in New Guinean metriorrhynchines (Fig. 4). The placement of sexually dimorphic species in the phylogeny suggests that the shift to dimorphism was very recent (3.0 mya at the earliest) and began when both sexes were small-bodied. Mimetic sexual polymorphism is well understood in butterflies with non-mimetic males and mimetic females (Kunte, 2008), but the advergence of males and females to different aposematic models has only recently been reported in two subfamilies of net-winged beetles (Motyka *et al*., 2018, 2020, 2021). Divergent evolution in Müllerian systems appears to be more common in multi-pattern aposematic rings than was previously believed when morphology was the sole source of information.

## Conclusion

Priority areas for global conservation have usually been identified based on richness, species endemism and vulnerability of vertebrates (Myers *et al*., 2000; Holt *et al*., 2013). We assume that different patterns of biodiversity distribution can be revealed if other animal groups are studied. Reliable information on additional groups can focus our conservation efforts on valuable regions (Morrison *et al*., 2009; Thomson *et al*., 2018). Our model, beetles, is the most speciose group of animals but is much less known than vertebrates. Therefore, new data must be generated, and our research workflow must be innovative. We conducted a worldwide sampling in ∼700 localities, analysed transcriptomes, genomes, and mitochondrial markers (>2,300 species), and validated our results with morphology. We achieved substantial progress with respect to the development of a Metriorrhynchini tree of life (Chesters, 2017; Linard *et al*., 2018). The voucher-based DNA entries established a framework for classifying samples from other studies, such as environmental sequencing (Linard *et al*., 2016; Andujar *et al*., 2015; Arribas *et al*., 2016). Despite limited time and funding, we identified >2,300 mOTUs which indicate that there are at least twice more species than the number previously reported in the literature. This means that, at a conservative estimate, 1,000–1,500 species in the dataset were previously unknown to science. Furthermore, we identified New Guinea as a biodiversity hotspot, which is in clear contrast with studies identifying the biodiversity patterns of vertebrates. Our accelerated inventory shows that the literature records of tropical beetles cannot be used for biodiversity conservation and metanalyses without critical revision. We suggest that if ∼20 person months of focused field research and subsequent workflow steps are applied to any hyperdiverse tropical group, the results can set a benchmark for future evaluation of spatiotemporal changes in biodiversity.

## Material and methods

### Field research

The analysed individuals had been accumulated by numerous expeditions to various regions of the Metriorrhynchini range (Fig. 1A, Tab. S1). The distribution of sampling sites was partly biased, and no samples are available from West Africa, Congo Basin, Sahel, Sri Lanka, and the Lesser Sundas. About 10% of samples were provided by other researchers.

Tissues for transcriptomic analyses were fixed in the field. As field identification is generally unreliable, we preferred to collect pairs *in copula*, then the female was fixed using RNAlater, and the male kept separately in 96% ethanol for Sanger sequencing and the voucher collection. Alternatively, the morphologically similar individual from the same place was fixed in ethanol and the identity of an individual assigned for transcriptomic analysis was confirmed by sequencing *cox1* mtDNA using tissue from the specimen preserved in RNAlater and putatively conspecific voucher (Fig. 2). About 100 tissue samples were fixed and thirty-five of them were used for sequencing (Tab. S2). Earlier published transcriptomes were added (McKenna *et al*., 2019; Kusy *et al*. 2019). Due to the inaccessibility of properly fixed tissue, the two critical samples were shotgun sequenced using isolated DNA.

Almost 7,000 samples from 696 localities (Tab. 1) were included in the sequencing program to obtain short mtDNA fragments. In total, 6,429 yielding at least a single fragment were included in the analysis (Tab. S3). The analysed data set contained some previously published sequences (e.g., Sklenarova *et al*., 2013; Bocek & Bocak, 2019). Voucher specimens are deposited in the collection of the Laboratory of Biodiversity and Molecular Evolution, CATRIN-CRH, Olomouc.

### Genomic and transcriptomic sequencing, data analysis

Libraries for thirty transcriptomes were prepared by Novogene Co., Ltd. (Beijing, China) and sequenced on the HiSeq X-ten platform (Illumina Inc., San Diego, CA). The removal of low-quality reads and TruSeq adaptor sequences were performed using fastp v.0.20.0 (Chen *et al*., 2018) with the following parameters: -q 5 -u 50 -l 50 -n 15. All paired-end transcriptomic reads were assembled using SOAPdenovo-Trans-31mer (Xie *et al*., 2014).

Additionally, the total DNA (∼33 Gb each) of *Metanoeus* sp. and an unidentified sample Metriorrhynchina species (Voucher JB0085) was shotgun-sequenced on the same platform. Reads were filtered with fastp using the same settings as above and quality was visualized with FastQC (http://www.bioinformatics.babraham.ac.uk/projects/fastqc). The draft genomes were assembled using SPAdes v.3.13.1 (Bankevich *et al*., 2012), with k-mer sizes of 21, 33, 55, 77, and 99. Obtained contigs were used to train Augustus (Stanke & Waack, 2003) for species-specific gene models with BUSCO. Predicted species-specific gene models were then used for *ab initio* gene predictions in Augustus and predicted protein-coding sequences were used for subsequent analyses. Outgroup taxa were reported in previous studies (Kusy *et al*., 2019; McKenna *et al*., 2019).

The ortholog set was collated by searching the OrthoDB 9.1 (Zdobnov *et al*., 2016) for single copy orthologs in six beetle genomes (Tab. S4). We used Orthograph v.0.6.3 (Petersen *et al*., 2017) with default settings to search in our assemblies for the presence of specified single copy orthologs. From the recovered 4,193 orthologs, terminal stop codons were removed, and internal stop codons at the translational and nucleotide levels were masked. The amino acid sequences were aligned using MAFFT v.7.471 with the L-INS-i algorithm (Katoh & Standley, 2013). The alignments from each ortholog group were then checked for the presence of outliers. To identify random or ambiguous similarities within amino acid alignments, we used Aliscore v.2.076 with the maximum number of pairwise comparisons –r 10^27^, option -e. and we masked them using Alicut v.2.3 (Kück *et al*., 2010). Alinuc.pl was then used to apply the Aliscore results to match amino acids to the nucleotide data. MARE v.0.1.2-rc was used to calculate the information content of each gene partition (Misof *et al*., 2013). Partitions with zero information content were removed at both levels. Finally, the remaining 4,109 alignments were retained for subsequent multispecies coalescent analyses, and different concatenated datasets were generated for both amino acid and nucleotide levels using FasConCat-G v.1.4 (Kück & Longo, 2014) (Tab. S5 and supplementary methods). The degree of missing data and overall completeness scores (Ca) across all datasets were inspected using AliStat v.1.7 (https://github.com/thomaskf/AliStat).

### Compositional heterogeneity tests

To explore the effect of among species compositional heterogeneity and its possible bias to tree reconstruction, we inspected the data with BaCoCa v.1.105 (Kück & Struck, 2014) to identify the gene partitions that strongly deviate from compositional homogeneity using relative composition frequency variation value (RCFV). Following Vasilikopoulos *et al*. (2019), we considered compositional heterogeneity among species in a given partition to be high when RCFV ≥0.1. The heterogeneous partitions were excluded from the data to generate a more compositionally homogeneous dataset. We used Maximum Symmetry Test (Naser-Khdour *et al*., 2019) to identify the partitions that strongly deviate from compositional homogeneity at the nucleotide level (p-value cut off <0.05), and partitions below the threshold were excluded. The software SymTest v.2.0.49 (https://github.com/ottmi/symtest) was used to calculate the overall deviation from stationarity, reversibility, and homogeneity (SRH) (Ababneh *et al*., 2006)

### Phylogenomic maximum likelihood analyses

For all datasets, phylogenetic reconstruction was performed using the maximum likelihood (ML) criterion with IQ-TREE 2.1.2 (Minh *et al*., 2020). First, we analysed all datasets using the original gene partition boundary. The model selection for each gene was performed with ModelFinder (Chernomor *et al*., 2016; Kalyaanamoorthy *et al*., 2017) implemented in IQ-TREE2 (-MFP option). GTR model was considered for nucleotide supermatrices. For the amino acid supermatrices, the substitution models LG, DCMUT, JTT, JTTDCMUT, DAYHOFF, WAG, and free rate models LG4X and LG4M were tested. All possible combinations of modelling rate heterogeneity among sites were allowed (options: -mrate E,I,G,I+G,R -gmedian -merit BIC). We used the edge-linked partitioned model for tree reconstructions (-spp option) allowing each gene to have its own rate. The optimized partition schemes and best-fitting models were inferred for some datasets using -m MFP+MERGE and the considering same substitution models as above. The fast-relaxed clustering algorithm was used to speed up computation during partition-scheme optimization (Lanfear *et al*., 2017). Ultrafast bootstrap (Hoang *et al*., 2017) and SH-like approximate likelihood ratio test (SH-aLRT) were calculated in IQ-TREE2 (options -bb 5000 and -alrt 5000) to assess nodal supports for focal relationships.

### Coalescent analyses and analyses of the confounding and alternative signal

To account for variation among gene trees owing to incomplete lineage sorting and to account for potential gene tree heterogeneity and discordance (Edwards, 2009), the data were also analysed using the coalescent-based species-tree method. For every single gene partition, we calculated an ML gene tree in IQ-TREE2, with 5000 ultrafast bootstrap replicates (-bb option) and using the same substitution models as predicted by ModelFinder in the above described partitioned analyses. For subsequent coalescent species tree estimation, the Accurate Species Tree Algorithm (ASTRAL-III v.5.7.3; Zhang *et al*., 2018) was used. To account for very poorly resolved branches on gene trees, branches with ultrafast bootstrap ≤ 10 were collapsed using newick utilities v.1.6 (Junier & Zdobnov, 2010) in every ASTRAL analysis. Local posterior probabilities (Erfan & Mirarab, 2016) and quartet frequencies of the internal branches in every species tree were calculated using the parameter ‘-t=2’. Pie charts representing quartet scores for the given topology and two alternatives were plotted to the resulting species trees in R using https://github.com/sidonieB/scripts/blob/master/plot_Astral_trees_v2.R.

Additionally, we studied the effect of potentially confounding signals, like non-random distribution of data coverage and violations of SRH conditions, on our phylogenetic reconstructions with the Four-cluster likelihood mapping (FcLM) approach (Strimmer & von Haeseler, 1997) implemented in IQ-TREE2. Based on the results of our tree reconstructions we tested the hypotheses about the alternative placement of leptotrichaline and procautirine clades.

### Mitochondrial DNA sequencing and data analysis

Total DNA was extracted from the metathorax with a Wizard SV96 kit (Promega Corp., Madison, WI). The yield was measured using a NanoDrop-1000 Spectrophotometer (Thermo Fisher Scientific Inc., Waltham, MA). The PCR settings and cycle sequencing conditions were the same as those used by Bocak *et al*. (2008). Three fragments of mitochondrial genome were sequenced: *cox1* + tRNA-*Leu* + *cox2* (∼1100 bp), *rrnL* + tRNA-*Leu* + *nad1* (∼800 bp), and ∼1210 bp of *nad5* and adjacent tRNA-*Phe*, tRNA-*Glu*, and tRNA-*Ser* mtDNA (the mtDNA fragments are further mentioned as *rrnL, cox1*, and *nad5*). The PCR products were purified using PCRu96™Plates (Merck Millipore Inc., Burlington, MA) and sequenced by an ABI 3130 (Applied Biosystems, Waltham, MA) sequencer using the BigDye® Terminator Cycle Sequencing Kit 1.1 (Applied Biosystems, Waltham, MA). Sequences were edited using Sequencher v.4.9 software (Gene Codes Corp., Ann Arbor, MI). Altogether 6,476 individuals were analysed including some previously published (Sklenarova *et al*., 2013; Bocek & Bocak, 2019).

The *cox1* gene fragment was used to OTUs delimitation (Blaxter *et al*., 2005) using CD-hit-est (Fu *et al*., 2012) and different thresholds (from similarity 0.99 to 0.90 by 0.05 steps). Therefore, we assembled two datasets: A) the dataset containing all sequenced individuals and B) all OTUs delineated by 0.98 similarity of the *cox1* gene. The *rrnL* and *tRNAs* were aligned using MAFFT 7.2 with Q-INS-I algorithm (Katoh & Standley, 2013), protein-coding genes were eye-checked for stop codons and aligned using TransAlign (Bininda-Emonds, 2005). All fragments were concatenated using FasConCat (Kück & Longo, 2014) and analysed under maximum-likelihood criterium in IQ-TREE v.2.1.2 (Minh *et al*., 2020; Tab. S6). To assess the branch supports values, we used SH-aLRT test with 1,000 iterations. ModelFinder tool implemented in IQ-TREE was used to identify the best fit models using the Bayesian Information Criterion (Chernomor *et al*., 2016). The results of the TSA/WGS analyses were used to constrain basal topology among major clades of Metriorrhynchini in both analyses of datasets A and B. Further, we ran unconstrained analyses of the above-mentioned datasets with identical settings except -g option to compare results. We replicated constrained tree search nineteen-times and compared resulting trees using Robinson–Foulds distances in R package phangorn (Schliep, 2011; Tab. S7). Randomly chosen trees were then compared using cophylo script (phytools; Revell, 2012) with argument rotate = TRUE.

## Supplementary Materials

The supplementary files are available online here.

Data Depositories.The mitogenomic dataset and all the analysed supermatrices are deposited in the Mendeley Data repository DOI: 10.17632/ntgg6k4fjx.1.

## Author Contributions

Conceptualization, L.B., DK, MM; formal analyses, M.M. (Sanger data), D.K. (phylogenomic data); data production and curation, L.B., M. B. and R.B.; writing, original draft preparation, L.B., M.M., D.K.; writing—review and editing, L.B., M.M., D.K., M.B., R.B.; visualization, L.B., M.M., D.K.; funding acquisition, L.B., M.M., D.K. All authors have read and agreed to the published version of the manuscript.

## Acknowledgements

Several colleagues provided samples from their respective home countries or their own field research. The field research was enabled by the permits from the Government of Papua New Guinea and the Queensland Ministry of Environment, Malaysian Ministry of Natural Resources, and UP Los Baños. The trips to Papua New Guinea were supported by the Binatang research centre, Nagada and we are obliged to H. Maraia and J. Kua for their field assistance. A substantial part of the research before the start of the project was funded by the senior author and Palacky University is acknowledged for granting the leave. L. Harmackova advised with some analyses in R.

