## Supplementary Materials for "Phylogenomic and mitogenomic data can accelerate inventorying of tropical beetles during the current biodiversity crisis"

**Electronic supplementary information**

**Supplementary text**

Introduction: Overview of Metriorrhynchini systematics and natural history
Methods: Detailed information on handling molecular data
Results: Morphology and composition of Metriorrhynchina subclades
References (selected studies dealing with Metriorrhynchini)
References (Supplementary methods section)

**Supplementary tables and figures**

Table S1. Detailed information on regional sampling
Table S2. List of material for transcriptomic analysis
Table S3. Completeness of the mtDNA dataset used for IQtree analysis
Table S4. Overview of official gene sets of six reference species used for ortholog assessment
Table S5. Phylogenomic datasets overview
Table S6. Characteristics of concatenated super-matrices and used models of the DNA evolution
Table S7. Comparison of the Robinson-Foulds distances among tree searches with constrained topology and tree searches with unconstrained topology
Table S8. Overview of the revised Metriorrhynchini generic classification and distribution

Figure S1-S8. Further trees recovered by phylogenomic analyses

Figure S9. AliStat heat maps of pairwise completeness scores (Cij) in amino acid datasets
Figure S10. AliStat heat maps of pairwise completeness scores (Cij) in nucleotide datasets
Figure S11. Heat maps calculated with SymTest showing p-values for the pairwise Bowker's tests in amino acid datasets
Figure S12. H eat maps calculated with SymTest showing p-values for the pairwise Bowker's tests in nucleotide datasets
Figure S13. Results of four-cluster likelihood mappings (FcLM) and tested phylogenetic hypotheses on the original and the permuted datasets.
Figure S14. Maximum likelihood tree recovered using whole dataset.
Figure S15. Maximum likelihood tree recovered using 98% mOTUs dataset with constrained backbone
Figure S16. Maximum likelihood tree recovered using 98% mOTUs dataset with unconstrained backbone
Figure S17. Subtribal mOTUs delimitation using CD-hit-est

Figure S18. A Distribution of selected Metriorrhynchini genera (originally assumed and revised)

#### Introduction: Overview of Metriorrhynchini systematics and natural history

Lycidae (net-winged beetles) is a diverse, mostly tropical family of Elateroidea and Metriorrhynchini with almost 1,600 described species is the largest tribe placed in the family (Bocak & Bocakova 2008, Masek et al. 2018).

The history of Metriorrhynchini higher classification started with Kleine's proposals of subfamilies and/or tribes Metriorrhynchini/inae, Trichalini/inae, Cladophorini/inae, and Dilolycini/inae (Kleine, 1926, 1928, 1933). Bocak & Bocakova (1990) revised the supergeneric classification of net-winged beetles, merged Conderini with Metriorrhynchini and Trichalini in the widely defined Metriorrhynchinae and synonymized Cladophorini and Dilolycini to Metriorrhynchini. Calder (1998) considered Metriorhynchinae (based on *Metriorhynchus* Guérin-Méneville, 1838) to be a homonym of Metriorhynchinae in Reptilia and used Dilolycinae Kleine, 1926 as a replacement name. Trichalinae were placed as a tribe Trichalini in Metriorrhynchinae (Bocak & Bocakova 1990) or a subtribe Trichalina in Metriorrhynchini (Bocak 2002). Additionally, Hemiconderina Bocak et Bocakova, 1990 were erected for *Hemiconderis* and the Australian genera *Achras* Waterhouse, 1879 and *Synchonnus* Waterhouse, 1879 were placed there along with the type genus. Bocak (2002) revised the generic classification using morphological dataset, but due to a limited amount of material for the study, many taxa were defined based on the type-species and a low number of close relatives. Therefore, the limits of the genus-rank lineages often remained ambiguous. Recently, Sklenarova et al. (2013) studied in detail the phylogeography of Metriorrhynchini and and revised the subtribal classification (Sklenarova et al. 2014).

The lower level metriorrhynchine taxonomy is mostly based on the original, often uninformative descriptions of a high number of species (e.g., Waterhouse, 1878, 1979, MacLeay 1872, 1886; Lea 1909, 1921; Pic, 1923, 1925; Kleine, 1930). Recently, Calder (1998) compiled the catalogue of Australian net-winged beetles and proposed *Metriorhynchus* Guérin-Méneville, 1838 as a junior homonym of *Metriorhynchus* Meyer, 1930 (Reptilia). Subsequently, *Porrostoma* Castelnau, 1838 was used as a replacement name for *Metriorhynchus* Guérin-Méneville, 1838 and new combinations and new replacement names were proposed (Calder 1998). Bocak (1998) reviewed the status of lycid taxa described by Guérin-Méneville (1838) and proposed to replace *Metriorhynchus* Guérin-Méneville, 1838 by *Metriorrhynchus* Gemminger et Harold, 1869. Bocak (2002) additionally considered *Porrostoma* Castelnau, 1838 as a valid name for a separate genus in Metriorrhynchini. The genera *Porrostoma* and *Metriorrhynchus* represent majority of Australian taxa although the placement is often based more on superficial similarity than on relationships.

There are five Metriorrhynchini genera which housed the majority of about 1,600 described species of Metriorrhynchini: *Xylobanus* (231 spp.), *Cautires* (437 spp.), *Trichalus* (120 spp.) *Metriorrhynchus* (194 spp.), and *Cladophorus* (131 spp.; Kleine, 1926, 1933; Calder, 1995; Bocak, 2002; Bocak & Bocakova 2008, Dudkova and Bocak 2010Kazantsev, 2015, 2016). *Trichalus* was recovered as a much smaller clade than *Microtrichalus* Pic, 1921, an originally monotypic genus from Java, into which have been transferred many *Trichalus* species whenever their types become available for study (Bocak 1998a, b, 1999, 2002, Bocek & Bocak 2017). The *Metriorrhynchus* diversity was split between *Porrostoma* Laporte, 1838 and *Metriorrhynchus* (Bocak 1998c). All Australian *Metriorrhynchus* incorrectly transferred to *Porrostoma* Castelnau, 1838 by Calder (1995) were formally returned to *Metriorrhynchus* by Bocak et al. (2020). *Xylobanus* was recovered as a clade that contains all Oriental species with four primary costae, but Afrotropical *Xylobanus* were recovered as a terminal clade within Afrotropical *Cautires* (Sklenarova et al. 2013, 2014) and some species earlier considered related to *Xylobanus* belong to Metanoeina (Kubecek et al. 2015). Numerous New Guinean species designated as *Xylobanus* belong to Metriorrhynchinae and not to Cautirina (Bocak et al. 2020). *Cautires* was recovered as a lineage which colonized Madagascar (Sklenarova et al. 2013) and continental Africa during the drift of Indian northwards. After the accretion of India to the Asian craton, it also colonized the Oriental and eastern Palearctic regions (Sklenarova et al. 2013, 2014). An additional genus, *Procautires* Kleine, 1925, was reported from various disjunctive ranges in Asia, Africa and Australasia (Kleine 1933), but the type species belong to Metriorrhynchina in contrast to the Afrotropical and Asian species which are close to *Cautires* (Cautirina).

*Distribution of Metriorrhynchini*

The tribe Metriorrhynchini dominates the Australian net-winged beetle fauna with ~85% (196 spp.) of the named taxa. They are similarly common in New Guinea (423 spp.) and the Wallacea (176 spp.). Although highly diversified (261 spp. and 231 spp., respectively), they represent only about 30% of described species in the Oriental and Afrotropical regions where similar number of species belong to Platerodini, Calochromini and Lycini, or several small tribes, respectively (Masek et al. 2018).

The tribe Metriorrhynchini has a Gondwanan distribution and the range reaches to the Palearctic East Asia (Russian Far East, Cautirina and Metanoeina). Additionally the tribe is well represented in the Sino-Japanese realm sensu Holt *et al*. (2013) (China, Japan, Cautirina and Metanoeina in the whole area) and in the Oriental realm (most of India, Sri Lanka, the lower elevations of the Himalayas, Indo-Burma, Malaya, the Great Sundas, Lesser Sundas, Philippines). Further, the tribe is widespread in the Oceania realm, in its western part: the Moluccas, New Guinea, the Solomon Islands and a few islands further west) and the Australian realm (Australia, Tasmania, and one introduced species in New Zealand). The Afrotropical realm houses relatively diversified fauna in forest and savannah habitats, but all species belong to Cautirina and represent only two clades which correspond with independent colonization events from drifting India to Madagascar and continental Africa (Sklenarova et al. 2013).

The subtribes differ in their distribution. Metriorrhynchina are mostly Oceanian and Australian, relatively rich Metriorrhynchina fauna is known from the Sulawesi but differs in the generic composition from the New Guinea fauna, much less species occur in the Philippines and only a few species of *Diatrichalus*, *Leptotrichalus*, *Trichalus*, *Microtrichalus*, and *Metriorrhynchus* occur in the eastern part of the Oriental realm, i.e. Indo-Burma, Malaya and the Sundaland (Bocak 1999, 2000a, b, 2007, Bocak et al. 2006, Bocak & Yagi 2010). The Cautirina occur in the Afrotropical, Oriental, Palearctic, and Sino-Japanese realm, but they do not cross Weber's line which was proposed as a border between Oriental and Oceanian realm by Holt et al. (2013). The Metanoeina is Oriental, with a limited number of species in China and Japan. They do not occur in the Philippines.

The Metriorrhynchina is the largest subtribe with 895 species. In the taxonomy of this group, we face in the burden of uninformative descriptions (e.g. the studies by the French entomologist M. Pic in the first half of the 20th century, Pic, 1921, 1922 etc.). The careless classification of newly described species based on poor knowledge of the group (often a single new species described in a large, poorly known genus without a reference to earlier described species, inaccessibility of type-holding collections or their poor organization, and legal regulations limit taxonomic research (Nijar et al. 2017).

**Biology of Metriorrhynchina**

The knowledge on biology of Metriorrhynchina is limited. The adults are volatile, but they are poor dispersers due to their soft-bodiedness and many species remain in shaded situations under the canopy, usually sitting on leaves. Some species of *Porrostoma*, *Metriorrhynchus*, *Leptotrichalus*, and *Trichalus* visit flowers, which they pollinate (Forster 1989). The flower frequenting species are more common in semi-dry areas where nectar represents a source of water for adults instead of water on leaves or in rotten wood. All Metriorrhynchini are unpalatable for predators and most aposematically colored (Moore & Brown 1981, 1989) similarly to other net-winged beetles (Eisner *et al*. 2008). The color patterns are usually geographically restricted and the sympatrically occurring species form Müllerian mimicry rings (Bocak & Yagi, 2010, Motyka et al. 2018, 2020, Bocek et al. 2019). Larvae of *Cautires*, *Leptotrichalus*, *Metriorrhynchus* and *Porrostoma* were described by Bocak & Matsuda (2003) and Levkanicova & Bocak (2009). The larvae live in rotten wood including trunks and logs on soil surface, or in rotten roots in soil in arid regions.

#### Supplementary Methods

Genomic and transcriptomic sequencing, data analysis, individual datasets details
Libraries for thirty transcriptomes were prepared by Novogene Co., Ltd. (Beijing, China) and sequenced on the HiSeq X-ten platform (Illumina). The removal of low-quality reads and TruSeq adaptor sequences were performed using fastp v.0.20.0 (Chen et al., 2018) with the following parameters: -q 5 -u 50 -l 50 -n 15. All paired-end transcriptomic reads were assembled using SOAPdenovo-Trans-31mer (Xie et al., 2014).

Additionally, the total DNA (~33 Gb each) of Metanoeus sp. and an unidentified sample JB0085 was shotgun-sequenced on the same platform by Novogene Co., Ltd. (Beijing, China) for 150 bp paired-end reads. Read were filtered with fastp using same setting as above and quality was visualized with FastQC (http://www.bioinformatics.babraham.ac.uk/projects/ fastqc).
The draft genomes were assembled using SPAdes v.3.13.1 (Bankevich et al., 2012), with all parameters set to default values and k-mer sizes of 21, 33, 55, 77 and 99. Obtained contig sequences were used to train Augustus (Stanke & Waack, 2003) for species specific gene models with BUSCO v.3 (Waterhouse et al., 2018), -long option, Endopterygota set of conserved genes (n = 2442) and -sp tribolium2012 as the closest relative. Predicted species specific gene models were then used for ab initio gene predictions in Augustus and predicted protein coding sequences were used for subsequent analyses. Seven outgroup taxa, Porrostoma and Mangkutanus transcriptomes were reported in previous studies (Kusy et al., 2019, McKenna et al., 2019). The filtering and assembling methods for these samples have been described by Kusy et al. (2019).

The ortholog set was collated by searching the OrthoDB 9.1 database (Zdobnov et al., 2016) for single copy orthologs in six beetle genomes Onthophagus taurus (Poelchau et al., 2014), Tribolium castaneum (Shelton et al., 2015, Richards et al., 2018) Dendroctonus ponderosae (Keeling et al., 2013), Anoplophora glabripennis (McKenna et al., 2017), Leptinotarsa decemlineata (Poelchau et al., 2014) and Agrilus planipennis (Poelchau et al., 2014) (Table S3). OrthoDB 9.1 predicted 4,225 single copy orthologs for beetle species and Coleoptera (Polyphaga) reference node. We used Orthograph v.0.6.3 (Petersen et al., 2017) with default settings to search in our assemblies for presence of specified single copy orthologs. From the recovered 4,193 orthologs, terminal stop codons were removed, and internal stop codons at the translational and nucleotide levels were masked using the Perl script summarize_orthograph_results.pl (Petersen et al., 2017). The amino acid sequences were aligned using MAFFT v. 7.471 with the L-INS-i algorithm (Katoh & Standley, 2013). The alignments from each ortholog group were then checked for the presence of outliers using the script checker_complete.1.3.1.2.pl, according to previously published methods (Misof et al., 2014; Peters et al., 2017). Identified outlier sequences were removed from amino acid and nucleotide alignments. After this step, the sequences of reference taxa and all gap-only sites were pruned. Corresponding multiple sequence alignments of nucleotides were generated using Pal2Nal (Suyama et al., 2006). To identify random or ambiguous similarities within amino acid alignments, we used Aliscore v. 2.076 with the maximum number of pairwise comparisons –r 1027, a special scoring approach for gap-filled amino acid sites option -e, and other parameters set to default values. Any random or ambiguous similarities were masked using Alicut 2.3 (Kück et al., 2010). Alinuc.pl was then used to apply the Aliscore results to match amino acids to the nucleotide data (Peters et al., 2017). Additionally, in each gene alignment, the short randomly aligned fragments were replaced with gaps and sequences with ≥80% missing data (calculated as percentage of '-' and 'X' in the amino acid alignments and '-' and 'N' in the nucleotide alignments) were removed using Python scripts (Zhang et al. 2020). MARE v. 0.1.2-rc was used to calculate the information

content of each gene partition in terms of amino acid coding (Misof et al., 2013). Partitions with zero information content (IC0) were removed at both amino acid and nucleotide level.
Finally, the remaining 4109 alignments were retained for subsequent multispecies coalescent analyses, and different concatenated datasets were generated for both amino acid and nucleotide levels using FasConCat-G v.1.4 (Kück & Longo, 2014). With all 4109 genes, we generated datasets designated as A-4199-AA and B-4199- NT (name-number of partitions-level). To reduce the effect of saturation on phylogenetic reconstruction (Breinholt & Kawahara 2013) we created the dataset C-4199-NT12 and E-4109-NT2, with third-codon or third and first-codon positions excluded.The degree of missing data and overall completeness scores (Ca) across all datasets were inspected using AliStat v.1.7 (https://github.com/thomaskf/AliStat) and heatmaps of pairwise completeness scores for all analysed datasets were generated (see electronic supplementary material, table S4). Further Information content and saturation values (the overall degree of data coverage with respect to gene presence or absence) were calculated with MARE v.0.1.2-rc (MAtrix REduction) (Misof et al., 2013) for each dataset.

Compositional heterogeneity tests

To explore the effect of among species compositional heterogeneity and its possible bias to tree reconstruction, we inspected the data with BaCoCa v.1.105 (Kück & Struck, 2014) to identify the gene partitions that strongly deviate from compositional homogeneity using relative composition frequency variation value (RCFV). Following Fernandez et al. (2016) and Vasilikopoulos et al. (2019), we considered compositional heterogeneity among species in a given partition to be high when the overall RCFV value was ≥0.1. The compositionally heterogeneous partitions were excluded from the dataset A-4109-AA to generate more compositionally homogeneous dataset D-3370-AA. To create more homogenous and decisive dataset, we reduced the dataset D-3370-AA to contain only partitions with all taxa and created dataset F-1490-AA. Additionally, we used MARE to increase information content of dataset D-3370-AA and default output supermatrix is designed as dataset J-2129-AA_MARE. We used tree matched-pairs tests of homogeneity the MPTS (matched-pairs test of symmetry), MPTMS (matched-pairs test of marginal symmetry), and MPTIS (matched-pairs test of internal symmetry) (Naser-Khdour et al., 2019) implemented in IQ-TREE2 to identify the partitions that strongly deviate from compositional homogeneity at nucleotide level (P-value cutoff <0.05) and partitions below the threshold were excluded. And following datasets generated: G-NT-1767_MaxSymTest, H-NT-1645_MaxSymTestmarginal and I-NT-3905_MaxSymTestInternal.

The software SymTest v.2.0.49 (https://github.com/ottmi/symtest) was used to calculate the overall deviation from stationarity, reversibility, and homogeneity (SRH) (Ababneh et al., 2006) between the amino acid and nucleotide sequences. Heatmaps were generated for all datasets to visualize the pairwise deviations from SRH conditions.

Phylogenetic maximum likelihood analyses

For all datasets phylogenetic reconstruction was performed using maximum likelihood (ML) criterion with IQ-TREE 2.1.2 (Minh et al., 2020). First, we analysed all datasets using original gene partition boundary. The model selection for each gene was performed with ModelFinder (Chernomor et al., 2016; Kalyaanamoorthy et al., 2017) implemented in IQ-TREE2 (-MFP option). GTR model were considered for nucleotide supermatrices. For the amino acid supermatrices, the substitution models LG, DCMUT, JTT, JTTDCMUT, DAYHOFF, WAG and free rate models LG4X and LG4M were tested. All possible combinations of modelling rate heterogeneity among sites were allowed (options: -mrate E,I,G,I+G,R -gmedian -merit BIC). We used the edge-linked partitioned model for tree reconstructions (-spp option) allowing each gene to have its own rate.
The optimized partition schemes and best-fitting models were inferred for dataset A-4109-AA using -m MFP+MERGE and considering same substitution models as above. The fast-relaxed clustering algorithm was used to speed up computation during partition-scheme optimization (Lanfear et al., 2017). The top 10% of partitions schemes --rclusterf 10 and maximum 12 327 partitions pairs (3 times the number of partitions) --rcluster-max 12 327 were considered. Dataset A-4109-AA was also analysed without any prior partitioning and under best fitting JTT+F+R9 substitution model. Ultrafast bootstrap (Hoang et al., 2017) and SH-like approximate likelihood ratio test (SH-aLRT)

were calculated in IQ-TREE2 (options -bb 5000 and -alrt 5000) to assess nodal supports for focal relationships (see electronic supplementary material, table S4).

Analyses of the confounding and alternative signal with four-cluster likelihood mapping (FcLM) and sequence data permutations
Additionally, we studied the effect of potentially confounding signal, like non-random distribution of data coverage and violations of SRH conditions, on our phylogenetic reconstructions with the Four-cluster likelihood mapping (FcLM) approach (Strimmer & von Haeseler, 1997) as described by Misof et al. (2014) and implemented in IQ-TREE2. Based on the results of our ML tree reconstructions we tested the hypotheses about alternative placement of Leptotrichaline and Procautirine clades. Permutation of data were executed as described in Misof et al. (2014) Our approach only differs in the use of the LG substitution matrix (Le and Gascuel, 2008) for permuting the supermatrices instead of WAG substitution matrix. For the analysis of the permuted supermatrices we used the option -q for the partition model in IQ-TREE2 and we used the LG model (for amino-acid alignments) and GTR model (for the nucleotide alignments). With original data, we performed the FcLM analyses by applying previously estimated substitution models (edgelinked models, -spp) with IQ-TREE2.

For estimation of *Cautires apterus* and sexually dimorphic *Metriorrhynchus* sp. splits was used subset of the samples which represent the most common lineages. Dating of forementioned diversification was analysed in BEAST 1.8.1 (Drummond et al., 2012) applying HKY +I+ G substitution model, uncorrelated relaxed clock model and birth–death process tree prior and 0.0115 (Brower et al. 1994) rate for cox1 gene. The stationary phase was checked in tracer 1.5 (http://beast.bio.ed.ac.uk/Tracer). The consensus tree was computed with TreeAnnotator 1.8.2 (http://beast.bio.ed.ac.uk/treeannotator) after eliminating a part of trees as a burn-in after evaluating ESS in tracer 1.5. The final trees were visualised in Figtree 1.4.4.

Data deposition: The additional supporting data and all the analysed supermatrices are deposited in the Mendeley Data repository DOI: 10.17632/ntgg6k4fjx.1

#### Supplementary Results

##### Morphology and composition of Metriorrhynchina subclades

*The procautirine clade*
The phylogenomic analysis contained four taxa that clustered as the first or second deepest serial branch in Metriorrhynchina. One of the internal clades contains species that are morphologically highly similar to a typical *Procautires.* They have reduced secondary elytral costae in the middle part of the elytra. Unlike this group of species, the members of other clades have fully developed secondary costae and would not be morphologically placed in *Procautires* relationships based on the structure of elytral costae. The only morphological trait supporting the procautirine clade is a setose patch in the internal sac of the male genitalia.

*The leptotrichaline clade*
The phylogenomic backbone identified close relationships of three endemic genera from the Sulawesi (*Mangkutanus*, *Wakarumbia*, and *Broxylus*), and two genera, *Sulabanus*, *Leptotrichalus*, known from Sulawesi and the Philippines, eventually also from the South East Asia. The mtDNA dataset identified as closely related the genera Austro-New Guinean *Synchonnus* and New Guinean *Falsolucidota*. We have not found any external diagnostic morphological character which would support the relationships of these genera.

Two lineages of the clade occur the Oriental region: *Leptotrichalus* (the Philippines, Sundas, Indo-Burma and China,) and *Sulabanus* (only the Philippines, 2 spp. from Mindanao and Sibuyan identified herein, either undescribed species or currently placed in Cautirina: *Xylobanus*).

*The trichaline clade*

The genomic dataset contained representatives of three genera, *Diatrichalus*, *Microtrichalus*, and *Eniclases*, all of them sharing the characteristic morphological traits of the trichaline clade, i.e. a single median areola in the pronotum and the shortened first primary costa. We identified as close relatives two lineages of small bodied Metriorrhynchina with conspicuous median pronotal areola and either absent or simplified lateral carinae, but with the full-length elytral costa 1. The structure of elytral costae is diverse in the clade: the trichalines *sensu stricto* mostly have well-developed secondary costae, i.e., nine costae at humeri, only a few *Diatrichalus* have only four primary costae in the humeral part. All genera of the here delimited trichaline clade share similar male genitalia. The deepest branches of the trichaline clade contain mostly New Guinean species and several lineages of the core trichalines colonized the Sulawesi, Philippines, Greater Sundas, Malaya, Indo-Burma and southernmost China.

*The porrostomine* clade

Six transcriptomes were analyzed to recover the backbone. *Porrostoma* and *Metriorrhynchus* are distantly related and easy to distinguish with genital morphology. As their close relatives were identified numerous small bodied, morphologically diverse species. Most *Metriorrhynchus* and its closest relatives comes from New Guinea, but several species of a terminal clade crossed Lydderker’s, Weber’s and Wallace’s lines and colonized Sulawesi, the Philippines, Greater Sundas and Malaya.

*The cladophorine clade*

Altogether 11 taxa were included in the genomic analysis and we found *Cladophorus* as a sister to other here included genera, i.e., *Pseudodontocerus* (=*Carathrix*) and *Ditua*. Most genera have seven pronotal areoles, male antennae are flabellate to pectinate and the elytra have nine well developed costae. Mitochondrial dataset recovered this clade as the most speciose group (626 mOTUs). *Cladophorus* subclade whose terminal subclade contains characteristic large-bodied species of *Cladophorus*, but as their relatives was recovered a quite high number of small-bodied Metriorrhynchina which do not resemble them. Further splits contain the *Pseudodontocerus* clade and a large clade represented by a high number of serial splits of subclades, some of them containing characteristic representatives of earlier described genera *Ditua* and *Cautiromimus*. The species of *Cladophorus* clade are most diverse in New Guinea, but some *Ditua* species were recorded from the Philippines and Sulawesi, a limited number of species is native to Australia.

The list of Metriorrhynchini genera with complete synonymy

Subtribe Metanoeina Kubecek et al., 2015
*Matsudanoeus* Sklenarova et al., 2014
*Metanoeus* Waterhouse, 1878
*Ochinoeus* Kubecek et al., 2015
*Xylometanoeus* Sklenarova et al. 2014

Subtribe Cautirina Sklenarova et al. 2015
*Caenioxylobanus* Waterhouse, 1878
*Cautires* Waterhouse, 1878
*Prometanoeus* Kleine, 1925
=*Tapromenoeus* Bocak et Bocakova, 1989
*Paracautires* Kazantsev, 2012
*Spartoires* Kazantsev, 2012
*Tricautires* Kazantsev, 2006
*Xylobanus* Waterhouse, 1878

Subtribe Metriorrhynchina Kleine, 1926a

Procautirine clade
*Procautires* Kleine, 1925b
*Xylothrix* Kazantsev, 2015a

Leptotrichaline clade
*Falsolucidota* Pic, 1921
=*Hemiconderis* Kleine, 1926a
*Broxylus* Waterhouse, 1879
=*Samanga* Pic, 1921)
*Mangkutanus* Kubecek et al., 2011
*Sulabanus* Dvorak & Bocak, 2007
*Synchonnus* Waterhouse, 1879
=*Achras* Waterhouse, 1879
=*Enylus* Waterhouse, 1879
=*Strophicus* Waterhouse, 1879
*Wakarumbia* Bocak, 1999a

Cladophorine clade
*Cautiromimus* Pic, 1926
*Cladophorinus* Kleine, 1926
*Cladophorus* Guérin-Méneville, 1830
= *Odontocerus* Guérin-Méneville, 1838
= *Spacekia* Strand, 1936
subgenus *Cladophorus* s. str.
subgenus *Falsocautires* Pic, 1926
*Ditua* Waterhouse, 1879
*Marena* Kazantsev, 2007
*Pseudodontocerus* Pic, 1921
=*Carathrix* Kleine, 1926a

Trichaline clade
*Diatrichalus* Kleine, 1926a
=*Mimoxylobanus* Pic, 1921
*Eniclases* Waterhouse, 1879
= *Trichalolus* Pic, 1923
*Flabellotrichalus* Pic, 1921
= *Stereotrichalus* Kleine, 1926
= *Villosotrichalus* Pic, 1921
subgen. *Flabellotrichalus* s. str.
subgen. *Maibrius* Bocek et Bocak, 2017
*Leptotrichalus* Kleine, 1925
*Lobatang* Bocak, 1998a
subgenus *Lobatang* s. str.
subgenus *Spinotrichalus* Kazantsev, 2010a
*Microtrichalus* Pic, 1921
= *Falsoenylus* Pic, 1926b
*Schizotrichalus* Kleine, 1926
*Trichalus* Waterhouse, 1877
= *Xantheros* Fairmaire 1877

Porrostomine clade
*Metriorrhynchus* Gemminger et Harold, 1869
=*Metriorhynchus* Guérin-Méneville, 1838: 72
=*Dilolycus* Kleine, 1926
=*Flabelloporrostoma* Pic, 1923a
*Porrostoma* Laporte, 1838
*Oriomum* Bocak, 1999b
*Metriorrhynchoides* Kleine, 1926
*Stadenus* Waterhouse, 1879
Metriorrhynchina incertae sedis
*Malacolycus* Kleine, 1943
*Xylobanomimus* Kleine, 1926
*Xylobanomorphus* Kleine, 1935
*Kassemia* Bocak, 1998

**Impact of molecular phylogenetics on the assessment of morphological evolution**
With robust data-rich phylogeny, we can test earlier hypotheses on morphological evolution. Relationships among subtribes have been based on the similarity of larvae and ambiguously supported by Sanger data (Sklenarova et al., 2013, 2014). Our results imply that bipartite larval terga have to be inferred as a synapomorphy of Metriorrhynchini with reversal to entire tergites in Cautirina (Sklenarova et al., 2014). The mapping of the character needs two instead of a single step on the tree and as it violates the principle of parsimony, it could not be inferred with morphological analysis. When most larvae are unknown and their morphology variable, the phylogenetic hypotheses based on incomplete morphological data and uncertain polarity urgently need validation from independent sources of information.

Due to quite high morphological disparity and incorrect interpretation of some morphological characters, there were proposed subfamilies, tribes, and/or subtribes for various subsets of genera now placed in the Metriorrhynchina (Metriorrhynchinae/ini, Cladophorinae/ini, Trichalinae/ini, Dilolycinae/ini, Hemiconderina, and possibly Melanerotini, now in Lycidae *incertae sedis*; see Bocak 2002, Kusy et al. 2019, Kazantsev, 2010). Phylogenomics rejects the possibility to assign a high rank to any of them as numerous coordinated taxa would have to be erected to make all groups reciprocally monophyletic and keep the earlier names valid.

The phylogenomic analyses reinforce the previously identified monophyly of the trichaline clade (=earlier Trichalini part, Tab. 2; Bocek et al. 2017), but simultaneously suggest their delayed origin as a terminal clade and the presence of some 'non-trichaline' lineages as serial sister-groups to trichalines *sensu stricto*. The cladophorine clade contains numerous taxa which had been placed until recently in *Cautires* (Cautirina), a distant lineage in respect to cladophorines. Herein, we further delimit the porrostomine clade which contains *Metriorrhynchus* and *Porrostoma*, but along with them several deeply rooted lineages which have been incorrectly placed to various genera. For the first time, we define the leptotrichaline clade which contains the genera earlier placed to trichalines (*Leptotrichalus*), Hemiconderina (*Falsolucidota*), Lycinae: Calopterini (*Broxylus*), Cautirina (some species of *Xylobanus*). A newly defined group is the procautirine clade. *Procautires* had been for long considered as related to *Cautires*, but it is one of earliest Metriorrhynchina.

Until recently, the classification had been based almost exclusively of the arrangement of pronotal ridges, elytral costae and the shape of antennae. These traits sometimes define a clade, but in many cases their presence and/or absence is misleading as they are highly plastic even within restricted cluster of species (Sklenarova et al. 2015, Bocek & Bocak 2017). Before independent validation, the importance of external morphological characters was apparently overestimated. As a result, the earlier studies coped with uncertain relationships of newly described species at the levels from a tribe down to a genus (Kleine, 1933, Calder 1995, Kazantsev 2015) and hundreds of new combinations had to be proposed to rectify the classification (Kazantsev, 2012; Bocak et al. 2020).

**References (selected studies dealing with Metriorrhynchini)**

Blackburn, T. Further notes on Australian Coleoptera, with descriptions of new species. Trans. R. Soc. S. Aust. 1888, 10, 177–287.

Blackburn, T. Notes on Australian Coleoptera, with descriptions of new species. Proc. Linn. Soc. N. S. W. 1892, 6, 479–550, doi:10.5962/bhl.part.29903.

Blackburn, T. Further notes on Australian Coleoptera, with descriptions of new genera and species. Part XXVI. Trans. R. Soc. S. Aust. 1900, 24, 35–68.

Bocak L. & Bocakova M. 1990a: Revision of the supergeneric classification of the family Lycidae (Coleoptera). *Polskie Pismo Entomol.* **59**: 623-676.

Bocak L. & Bocakova M. 1990b: Revision of the genus Hemiconderis (Coleoptera, Lycidae). *Acta Entomol. Bohemoslov.* **87**: 209-220.

Bocak L. & Bocakova M. 1991: Revision of the genus Eniclases C. O. Waterhouse, 1879 (Coleoptera, Lycidae, Metriorrhynchinae). *Mitt. Münch. Entomol. Ges.* **81**: 203-226.

Bocak L. & Matsuda K. 1998: A review of the Metriorrhynchus thoracicus group (Coleoptera: Lycidae). *G. Ital. Entomol.* **8** (1997): 409-415.

Bocak L. & Matsuda K. 2003. Review of the immature stages of the family Lycidae (Insecta: Coleoptera). *J. Nat. Hist.*
Bocak, L. Nomenclatural notes on taxa of the family Lycidae described by Guérin-Méneville (Insecta: Coleoptera). Ann. Zool. 1998, 48, 245–251.
Bocak, L. A generic revision and phylogenetic analysis of the subtribe Trichalinina (Coleoptera: Lycidae: Metriorrhynchini). Acta Soc. Zool. Boh. 1998, 62, 167–200.
Bocak, L.; Jašš, R. Revision of the genus *Broxylus* Waterhouse (Coleoptera, Lycidae). Dtsch. Entomol. Z. 2004, 51, 65–75, doi:10.1002/mmnd.4810510106.
Bocak, L. A revision of the genus *Microtrichalus* Wat. from the Philippines (Coleoptera: Lycidae). Eur. J.,Entomol. 1998, 95, 417–428.
Bocak L. 1998e: A new species of the genus *Eniclases* Wat. (Coleoptera, Lycidae). *Acta Univ. Pal. Olom., Biologica* **35** (1997): 13-16.
Bocak L. 1999a: A new genus *Oriomum* from New Guinea (Coleoptera, Lycidae, Metriorrhynchini). *Entomol*. *Basiliensia* **21** (1998): 111-114.
Bocak L. 1999b: New taxa of the subtribe Hemiconderinina (Coleoptera, Lycidae) from Indonesia and New Guinea. *Entomol. Bl.* **95**: 166-170.
Bocak, L. A revision of the genus *Diatrichalus* Kleine from New Guinea (Coleoptera: Lycidae). Stuttg. Beitr. Naturk. A 2001, 622, 1–32.
Bocak, L. Revision of the genus *Wakarumbia* (Coleoptera: Lycidae). Eur. J. Entomol. 2000, 97, 271–278, doi:10.14411/eje.2000.043.
Bocak L. 2000a: A revision of the genus Diatrichalus Kleine from the Philippines. *Raffl. Bull. Zool.* **48**(1): 11-16.
Bocak, L. To the knowledge of the genus *Leptotrichalus* Kleine from Sumatra, Borneo and Continental Asia (Coleoptera: Lycidae). Ann. Soc. Entomol. Fr. 2000, 36, 71–184.
Bocak, L. A revision of the genus *Diatrichalus* Kleine from the Philippines (Coleoptera: Lycidae). Raffl. Bull. Zool. 2000, 48, 11–16.
Bocak, L. New species of the genus *Wakarumbia* from Sulawesi (Coleoptera: Lycidae). Eur. J. Entomol. 2001, 49, 1–9.
Bocak, L. A review of the genus *Microtrichalus* Pic from Sumatra, with notes on Oriental and Australian species (Coleoptera: Lycidae). Acta Soc. Zool. Bohem. 1999, 64, 3–16.
Bocak, L. A Revision of *Metriorrhynchus* (Coleoptera: Lycidae) from the Greater Sunda Islands and Continental. Asia. Eur. J. Entomol. 2007, 55, 253–260.
Bocak, L.; Bocakova, M. Revision of the genus *Hemiconderis* (Coleoptera, Lycidae). Acta Entomol. Bohemoslov. 1990, 89, 209–220.
Bocak L. 2001a: A revision of the genus Diatrichalus Kleine from New Guinea (Coleoptera: Lycidae). *Stuttgarter Beitr. Naturk*., Ser. A. **622**: 1-32.

Bocak, L. Revision and phylogenetic analysis of Metriorrhynchinae. Eur. J. Entomol. 2002, 241, 315–351.
Bocak, L.; Bocakova, M. Revision of the supergeneric classification of the family Lycidae (Coleoptera). Pol. Pismo Entomol. 1990, 59, 623–676.
Bocak, L.; Bocakova, M. Revision of the genus Eniclases Waterhouse, 1879 (Coleoptera, Lycidae, Metriorrhynchinae). Mitt. Münch. Entomol. Gess. 1991, 81, 203–226.
Bocak, L.; Bocakova, M. Phylogeny and classification of the family Lycidae (Insecta: Coleoptera). Ann. Zool. 2008, 58, 695–720, doi:10.3161/000345408x396639.
Bocak L., Grebennikov V. V., Sklenarova K. (2014) Cautires apterus, a new species and the first record of wingless male Lycidae (Coleoptera) discovered in the North Pare Mountains, Tanzania. Annales Zoologici 64(1): 1-7.
Bocak, L.; Matsuda, K. Review of the immature stages of the family Lycidae (Insecta: Coleoptera). J. Nat. Hist. 2003, 37, 1463–1507, doi:10.1080/00222930210125362.
Bocak, L.; Matsuda, K.; Yagi, T. A revision of Metriorrhynchus from the Philippines with molecular evidence of an Australian origin of the Oriental Metriorrhynchus fauna (Coleoptera: Lycidae). Eur. J. Entomol. 2006, 103, 115–126, doi:10.14411/eje.2006.017.

Bocak, L.; Yagi, T. Evolution of mimicry patterns in Metriorrhynchus (Coleoptera: Lycidae): The history of dispersal and speciation in South East Asia. Evolution 2010, 64, 39–52.
Bocaková M. & Bocak L. 2007. Lycidae. Pp. 211–224. In Löbl I. & Smetana A. (eds.). Catalogue of Palaearctic Coleoptera.  4. Stenstrup: Apollo Books. 935 pp.Bocek, M.; Bocak, L. Where are species limits in polymorphic mimetic beetles from New Guineanmountains: A case of Eniclases net-winged beetles (Lycidae: Metriorrhynchini). ZooKeys 2016, 593, 15–35.
Bocek, M. New species of Diatrichalus (Coleoptera: Lycidae) from New Guinea and the Moluccas. Zootaxa 2017, 4247, 577–584, doi:10.11646/zootaxa.4247.5.4.
Bocek, M.; Adamkova, K. New species of Moluccan trichaline net-winged beetles, with remarks on the phylogenetic position and distribution of Schizotrichalus (Coleoptera: Lycidae: Metriorrhynchinae). Zootaxa 2019, 4623, 341–350, doi:10.11646/zootaxa.4623.2.8.
Bocek, M.; Bocak, L. The molecular phylogeny and classification of trichaline net-winged beetles (Lycidae: Metriorrhynchini). PeerJ 2017, 5, e3963.
Bocek, M.; Bocak, L. The origins and dispersal history of the trichaline net-winged beetles in South East Asia, Wallacea, New Guinea and Australia. Zool. J. Linn. Soc. 2019, 185, 1079–1094.
Bocek, M.; Motyka, M.; Kusy, D.; Bocak, L.; Kusy Genomic and mitochondrial data identify different species boundaries in aposematically polymorphic eniclases net-winged beetles (Coleoptera: Lycidae). Insects 2019, 10, 295, doi:10.3390/insects10090295.
Boisduval J. B. A. 1835: *Faune entomologique de l'Océanie, comprenant les Coléoptères, les Hemiptères, les Nevroptères, les Hymenoptères et les Diptères*. Bd. 1. Roret, Paris, 267 pp.
Bourgeois J. 1882. Deux notes relatives á différentes espèces de Lycides. Annales de la Société entomologique de France 2(6), 100–102.
Bourgeois J. 1889. Voyage de M. Ch. Alluaud dans le territoire d'Assinie (Afrique occidentale) en juillet et août 1886. 1re Mémoire. Annales de la Société entomologique de France 58, 237–246.
Bourgeois J. 1897. Description de Malacodermes nouveaux recueil- lis en Océanie par M. le Dr. Ph. François. Bulletin de la Société entomologique de France 66, 72–74.
Bourgeois J. 1900. Etude sur les Lycides du Musée Royal d'Histoire Naturelle de Belgique (espèces du Congo et pays voisins). Annales de la Société entomologique de Belgique 44, 139–146.
Bourgeois J. 1902. Diagnoses de Lycides nouveaux ou peu connus. 8e partie. Annales de la Société entomologique de France 71, 735–750.
Bourgeois J. 1905. Malacodermes de la Guinée Espagnole. Memo- rias de la Sociedad espanola de Historia natural. 1. P.189–196. Bourgeois J. 1908. Diagnoses de Lycides nouveaux ou peu connus. 9e partie. Annales de la Société entomologique de France 77, 501–504.
Bourgeois J. 1910. 10. Malacodermata. Wissenschaftliche Ergebnisse der Schwedischen Zoologischen Expedition nach dem Kilimandjaro, dem Meru und den umgebenden Massaisteppen, Deutsch-Ostafrikas, 1905–1906, unter Leitung von Prof. Dr. Yngve Sjøsted 7. Coleoptera. Stockholm: P. Palmquists. P.105–138.Calder, A.A. Coleoptera: Elateroidea. In Zoological Catalogue of Australia. Wells, A., Ed.; CSIRO Publishing: Melbourne, Australia, 1998; Volume 29.6, p. 248.
Calder A. A. 1998: Coleoptera: Elateroidea. In Wells A. (ed.): *Zoological Catalogue of Australia. Volume 26.9.* CSIRO Publishing, Melbourne, xiii + 248 pp.
Castelnau Laporte de, F. L. 1838: Études entomologiques ou descriptiones des insectes nouveaux et observations sur leur synonymie. *Rev. Entomol.* **4:** 5-60.
Dalman, J.W. Appendix ad C.J. Schonherr Synonymiam Insectorum Sistens Descriptiones Novarum Specierum; Schonherr, C.J., Ed.; Officina Lewerentziana: Scarfs, Sweden. Volume 1; Part 3, 1818; 266 pp.
Dudkova, P.; Bocak, L. A review of the Cautires obsoletus species group from Indo–Burma (Coleoptera: Lycidae). Zootaxa 2010, 2527, 28–48, doi:10.11646/zootaxa.2527.1.2.
Dvorak, M.; Bocak, L. Sulabanus gen. nov., a new genus of Lycidae (Coleoptera) from Sulawesi. Zootaxa 2007, 1611, 1–24, doi:10.11646/zootaxa.1611.1.1.

Dvorak, M.; Bocak, L. Ten new species of Wakarumbia Bocak, 1999 from Sulawesi (Coleoptera: Lycidae), with a key to males of the genus. Zootaxa 2009, 2282, 51–61, doi:10.11646/zootaxa.2282.1.2.
Erichson W. F. 1842: Beitrag zur Fauna von Vandiemensland, mit besonderer Ruecksicht auf die geographische Verbreitung der Insecten. *Weigeman Archiv* **8**: 83-287.
Fåhraeus C.H. 1851. In: Boheman C.H. Insecta Caffrariae annis 1838–1845 J.A. Wahlberg collecta, amici auxilio suffutus. Pars I. Fasc. II. Holmiae [=Stockholm]: Norsted Pp. 299–626.
Fabricius J. C. 1775: *Systema Entomologiae, sistens insectorum classes, ordines, genera, species, adiectis synonymis, locis, descriptionibus, observationibus*. Korte, Flensburgi et Lipsiae, xxvii+832 pp.
Fabricius, J.C. Systema Eleutheratorum Secundum Ordines, Genera, Species: Adiectis Synonymis, Locis, Observationibus, Descriptionibus; Bibliopolii Academici: Kiliae, Germania, 1801.
Fairmaire L. 1869. Notes sur les Coléoptères recueillis par Charles Coquerel á Madagascar et sur les côtes d’Afrique. 2e partie. Annales de la Société entomologique de France 9(4), 179–260.
Fairmaire L. M. 1877: Diagnoses des Coléoptères australiens et melanesiens. *Petit Nouv. Entomol.* **2**: 166-167.
Fairmaire L. M. 1879: Descriptions de Coléoptères nouveaux ou peu connus du Musée Godeffroy. *Journ. Mus. Godeffroy* **14**: 80-114.
Fairmaire L. 1880. Description de quelques Coléoptères de Nossi-Bé. Annales de la Société entomologique de France 10(5), 321–340.
Fairmaire L. 1884. Note sur les Coléoptères recueillis par M. Ach. Raffray à Madagascar et descriptions des espèces nouvelles (1re P.). Annales de la Société entomologique de France 4(6), 229–234.
Fairmaire L. 1887a. Coléoptères des voyages de M. G. Revoil chez les Somalis et dans l’Intérieur du Zanquebar. Annales de la Société entomologique de France 6(6), 69–186, 277– 368.
Fairmaire L. 1887b. Diagnoses de Coléoptères nouveaux de Madagascar. Le Naturaliste 9, 56–57.
Fairmaire L. 1896. Matériaux pour la faune coléoptérique de la région malgache. Annales de la Société entomologique de Belgique 40, 336–398.
Fairmaire L. 1901. Matériaux pour la faune coléoptérique de la région malgache (11e note). Revue d’Entomologie 20, 101–248.Gemminger, M.; Harold, E. Catalogus Coleopterorum Hucusque Descriptorum Synonymicus et Systematicus, Autoribus Dr. Gemminger et B. de Harold; E. H. Gummi: Monachii, Germania, 1868.
Gemminger M. & Harold E. von 1869: *Catalogus Coleopterorum hucusque descriptorum synonymicus et systematicus.* Band **6**. Rhipiceridae-Cionidae. E. Deyrolle fils, Paris, pp. 1609-1800.
Gerstaecker A. 1871. Beitrag zur Insectenfauna von Zanzibar. III. Coleoptera. Archiv für Naturgeschichte 37, 42– 86.
Guérin-Méneville, F.E. Crustacés, arachnides et insectes. In Voyage autour du Monde sur la Corvette de Sa Majesté, La Coquille, pendant les annés 1822, 1823, 1824 et 1825. Zoologie, Plates; Duperrey, L.I., Ed.; Bertrand: Paris, France, 1830.
Guérin-Méneville F. L. 1833: Mémoire sur deux nouveaux genres de l’ordre des Coléopteres, et déscriptions des éspeces qui les composent. *Ann. Soc. Entomol. Fr.* **1833**: 155-160.
Guérin-Méneville, F.E. Crustacés, arachnides et insectes. In Voyage autour du monde sur la Corvette de Sa Majesté, La Coquille, pendant les annés 1822, 1823, 1824 et 1825. Zoologie. Vol. 2, Part 2; Duperrey, L.I., Ed.; Arthus Bertrand: Paris, France, 1838.
Jiruskova, A.; Bocak, L. Species delimitation in Cautires (Coleoptera: Lycidae) from Peninsular Malaysia using DNA data and morphology. Ann. Zool. 2015, 65, 239–248, doi:10.3161/00034541anz2015.65.2.007.
Jiruskova,A.;Motyka,M.;Bocak,L.HighdiversityandendemisminthegenusCautiresWaterhouse,1879 (Coleoptera: Lycidae) from the Malay mountain forests, with the descriptions of fourteen new species. Eur. J. Taxon. 2016, 219, 1–29, doi:10.5852/ejt.2016.219.
Jiruskova,A.;Motyka,M.;Bocek,M.;Bocak,L.The Malacca Strait separates distinct faunas of poorly-flying Cautires net-winged beetles. PeerJ 2019, 7, e6511, doi:10.7717/peerj.6511.

Kalousova, R.; Bocak, L. Species delimitation of colour polymorphic Cladophorus (Coleoptera: Lycidae) from New Guinea. Zootaxa 2017, 4320, 505–522, doi:10.11646/zootaxa.4320.3.6.
Kleine, R. Die Lyciden der Philippinen-Inseln. Philipp. J. Sci. 1926, 31, 33–114.
Kazantsev S.V. 2000. New lycids from China (Coleoptera, Lycidae). Miscellania Zoologica. 23, 79–92.
Kazantsev S.V. 2002. New and little known species of Lycidae (Coleoptera) from China. Russian Entomological Journal 11, 253–263.
Kazantsev S.V. 2006a. A new African genus of Metriorrhynchini (Coleoptera: Lycidae). Zoosystematica Rossica 14, 219–221.
Kazantsev S.V. 2006b. New taxa of net-winged beetles (Coleoptera, Lycidae) from Sri Lanka. Eurasian Entomological Journal 5, 127–131 (in Russian).
Kazantsev S.V. 2007. Marena gen.n., new Metriorrhynchini genus from New Guinea (Coleoptera: Lycidae). Russian Entomological Journal 16, 297–300.
Kazantsev S.V. 2010. New taxa of Papuan net-winged beetles (Lycidae, Coleoptera). Latvijas Entomologs 48, 92–100.Kazantsev, S.V. New Xylobanus and related taxa of net-winged beetles from New Guinea (Coleoptera: Lycidae). Russ. Entomol. J. 2015, 24, 107–118, doi:10.15298/rusentj.24.2.02.
Kazantsev S. V. 2010. New taxa of Papuan net-winged beetles (Lycidae, Coeoptera). Latvijas Entomolgs, 48: 92–100.

Kazantsev, S.V. New species of Metriorrhynchus Gemminger et Harold, 1869 and Porrostoma Laporte, 1838 from New Guinea (Coleoptera: Lycidae). Russ. Entomol. J. 2015, 24, 211–233.
Kazantsev, S.V. New and little known species of Falsolucidota Pic, 1921 from New Guinea (Coleoptera: Lycidae). Russ. Entomol. J. 2016, 25, 355–360, doi:10.15298/rusentj.25.4.03.
Kazantsev, S.V. On the status and systematic position of the genera Falsolucidota Pic, 1921 and Macrolycinella Pic, 1922, with notes on Lopheros LeConte, 1881 (Coleoptera: Lycidae). Russ. Entomol. J. 2009, 18, 277–283.
Kazantsev, S.V. New Taxa of Papuan net-winged beetles (Lycidae, Coleoptera). Latv. Entomol. 2010, 48, 92–100.
Kazantsev, S.V. 2015b New species of *Metriorrhynchus* Gemminger et Harold, 1869 and Porrostoma Laporte, 1838 from New Guinea (Coleoptera: Lycidae). Russ. Entomol. J. 2015, 24, 211–233.
Kazantsev, S.V. 2016a. A review of *Ditua* Waterhouse, 1879 with description of seven new species from New Guinea (Coleoptera: Lycidae). Russ. Entomol. J. 25, 239–248. doi:10.15298/rusentj.25.3.03.
Kazantsev, S.V. 2015a New *Xylobanus* and related taxa of net-winged beetles from New Guinea (Coleoptera: Lycidae). Russ. Entomol. J. 2015, 24, 107–118.
Kirsch T. 1875: Neue Käfer aus Malacca. *Mitt. Dresd. Mus.* **1**: 25-58.
Kleine R. 1925a: Die neue Gattung Leptotrichalus*. Philippine J. Sci.* **28**: 295-311.
Kleine R. 1925b: Fauna Buruana. Coleoptera, Fam. Lycidae (2. Beitrag zur Kenntnis der Lycidae). *Treubia* **7**(1): 31-37.
Kleine R. 1925c: Drei neue Lycidengenera des orientalischen Gebietes. 8. Beitrag zur Kenntnis der Lycidae. Stett. Entomol. Zeitung 85: 133-136.
Kleine R. 1926a: Coleoptera, Lycidae. *Nova Guinea, Zoologie* **15**(2): 91-195.
Kleine R. 1926c: Die Lyciden Formosas. *Stett. Ent. Zeit*. **87**: 97-110.
Kleine R. 1926. Bestimmungstabelle der Gattung *Cautires* C.O.Waterhouse. Archiv für Naturgeschichte Ser. A, 92, 118–144.
Kleine R. 1927. Bestimmungstabelle der Gattung *Xylobanus* C.O.Waterhouse. Tijdschrift voor Entomologie 70, 43–72.
Kleine R. 1928: Neue Indische Lycidae nebst faunistische Bemerkungen. *Indian Forest Rec.* **13**: 221-268.
Kleine R. 1930a. Die *Cautires*-Arten des Congo-Museums nebst Bestimmungsschlussel der afrikanischen Arten. Revue de Zoologie et de Botanique Africaines 19. P.4–19.

Kleine R. 1930a: Bestimmungtabelle der Trichalusverwandschaft. *Treubia* **11**: 325-341.
Kleine R. 1930b: Neue Lyciden der malaiischen Subregion. *Journ. Fed. Malay. Stat. Mus.*, **16**: 169-174.
Kleine R. 1930b. Studien zur Gattung *Bulenides* C. O. Wtrh. Treubia 11, 349–355.
Kleine R. 1930c. Neue *Xylobanus*-Arten aus Westafrika nebst Fundnotizen. Revue de Zoologie et de Botanique Africaines 19, 172–176.
Kleine R. 1933a. Pars 123: Lycidae. Coleopterorum Catalogus auspiciis et auxilio W. Junk editus a Schenkling. Berlin: W. Junk. 145 p.
Kleine R. 1933b. Neue Lyciden und Bemerkungen zum Ca Col. Junk-Schenkling Lycidae. Stettiner Entomologische Zeitung 94, 1–20.
Kleine R. 1935. Neue Lyciden aus Afrika. Revue de Zoologie et de Botanique Africaines 26, 425–429.
Kleine R. 1935: Bericht ueber die von Miss Cheesman in British New Guinea gesammelten Brenthiden und Lyciden. *Nova Guinea, Zoologie* **17**: 302-322.
Kleine R. 1936: Check list of Pacific Lycidae. *Bernice Pauhu Bishop Mus. Occ. Pap.* **12**: 1-7.
Kleine R. 1937. Les Lycides du Congo Belge. Annales du Musée du Congo Belge Ser.3. 2, 61–120.
Kleine R. 1939: Neue Lyciden von den orientalischen Inseln und Molukken. *Treubia* **17**: 1-19.
Kleine R. 1940. Neue Brenthiden und Lyciden aus dem Congo- Museum. Revue de Zoologie et de Botanique Africaines 33, 245–255.
Kleine R. 1942. Brenthidae und Lycidae (Addendum). Expl. Parc National Albert, I. Mission G.F. de Witte 1933–1935 8, 1–9.
Kleine R. 1943: Neue Lyciden aus dem Ungarischen National-Museum (Coleopt.) *Acta Mus. nat. Hung., Zoologica* 36: 145-156.
Kleine, R. Bestimmungstabelle der Gattung Trichalus. Treubia 1929, 10, 471–493.
Kleine, R. Fauna Buruana. Coleoptera, Fam. Lycidae. (2. Beitrag zur Kenntnis der Lycidae.). Treubia 1925, 7, 31–37.
Kleine, R. Coleoptera Lycidae. Nova Guinea 1926, 13, 91–195.
Kleine, R. Bericht ueber die von Miss Cheesman in Britisch Neu-Guinea gesammelten Brenthiden und Lyciden. Nova Guinea 1935, 17, 303–322.
Kubecek, V.; Bray, T.C.; Bocak, L. Molecular phylogeny of Metanoeina net-winged beetles identifies Ochinoeus, a new genus from China and Laos (Coleoptera: Lycidae). Zootaxa 2015, 3955, 113–122, doi:10.11646/zootaxa.3955.1.6.
Kubecek, V., Dvorak, M.; Bocak, L. The phylogenetic structure of Metriorrhynchini fauna of Sulawesi, (Coleoptera: Lycidae) with description of a new genus. Zool. Stud. 2011, 50, 645–656.
Kalousova R., Bocak L. 2017. Species delimitation of colour polymorphic Cladophorus (Coleoptera: Lycidae) from New Guinea. Zootaxa, 4320: 505–522.
Kusy, D. A new species of Synchonnus (Coleoptera: Lycidae) from New Guinea, with an identification key to the Papuan species. Acta Entomol. Musei Natl. Pragae 2017, 57, 153–160, doi:10.1515/aemnp-2017-0064.
Kusy, D.; Sklenarova, K.; Bocak, L. The effectiveness of DNA-based delimitation in Synchonnus net- winged beetles (Coleoptera: Lycidae) assessed, and description of 11 new species. Austral Entomol. 2017, 57, 25–39, doi:10.1111/aen.12266.
Kusy, D.; Motyka, M.; Bocek, M.; Masek, M.; Bocak, L. Phylogenomic analysis resolves the relationships among net-winged beetles (Coleoptera: Lycidae) and reveals the parallel evolution of morphological traits. Syst. Entomol. 2019, 44, 911–925, doi:10.1111/syen.12363.
Lea, A.M. Descriptions of new species of Australian Coleoptera. Proc. Linn. Soc. N. S. W. 1895, 9, 589–634.
Lea, A.M. Descriptions of new species of Australian Coleoptera. Part V. Proc. Linn. Soc. N. S. W. 1899, 23, 521–645.
Lea, A.M. The Coleoptera of King Island, Bass Strait. Proc. R. Soc. Vic. 1908, 20, 143–207.

Lea, A.M. Revision of the Australian and Tasmanian Malacodermidae. Trans. Entomol. Soc. Lond. 1909, 45–251.
Lea, A.M. On Coleoptera, mostly from Queensland. Mem. Qld Mus. 1909, 7, 182–240.
Lea, A.M. On Australian Coleoptera of the family Malacodermidae. Trans. R. Soc. S. Aust. 1921, 45, 50–135.
Lea, A.M. Descriptions of new species of Australian Coleoptera. Part XVII. Proc. Linn. Soc. N. S. W. 1921, 46, 351–369, doi:10.5962/bhl.part.14024.
Lea, A.M. Results of Dr. E. Mjöberg’s Swedish scientific expeditions to Australia 1910–1913. Cryptophagidae, Cucujidae, Malacodermidae, Melandryidae, Mordellidae, Rhipidophoridae and Oedemeridae. Ark. Zool. 1922, 14, 1–21.
Lea, A.M. On Coleoptera, mostly from Queensland. Mem. Qld Mus. 1929, 9, 335–363.
Levkanicova, Z.; Bocak, L. Identification of net-winged beetle larvae (Coleoptera: Lycidae) using three mtDNA fragments: A comparison of their utility. Syst. Entomol. 2009, 34, 210–221, doi:10.1111/j.1365-3113.2008.00457.x.
Macleay, W.J. The insects of the Fly River, New Guinea, “Coleoptera”. Proc. Linn. Soc. N. S. W. 1886, 1, 136–157.
Macleay, W.J. The Insects of the Cairns District, Northern Queensland. Proc. Linn. Soc. N. S. W. 1887, 2, 211–238.
Murray A. 1868. List of Coleoptera received from Old Calabar, on the West Coast of Africa. The Annals and Magazine of Natural History, including Zoology, Botany and Geology (series 4) 2. 91–111.
Nakane T. 1968: On Some New Species of Lycidae from Formosa (Insecta, Coleoptera). *Bull. Nat. Science Mus.* **11**(3): 217-224.
Nakane T. 1969a: *Fauna Japonica, Lycidae (Insecta, Coleoptera).* Academic Press of Japan, Tokyo, 224 pp.
Nakane T. 1969b: New Species of Lycidae from Formosa, with Notes on Some Known Species (Insecta: Coleoptera). *Bull. Nat. Science Mus.* **12**(1): 9-16.
Nakane T. 1971: Formosan Lycidae in the Collection of the California Academy of Sciences (Insecta, Coleoptera). *Bull. Nat. Science Mus.* **14**(2): 137-155.
Nakane T. 1980: New or Little-Known Coleoptera from Japan and its Adjacent Regions, xxxii. *Rep. Fac. Science Kagoshima Univ. (Earth Sciences and Biology)* **13**:127et130.
Pic M. 1915: Genres nouveaux, éspèces et variétes nouvelles. *Mélanges exotico-entomologiques* **16**: 2-13.
Pic M. 1921a: Contribution à l'étude des Lycides. *L'Echange* **405**: 5-8 hors-texte.
Pic M. 1921b: Contribution à l'étude des Lycides. *L'Echange* **406**: 9-12 hors-texte.
Pic M. 1922: Contribution à l'étude des Lycides. *L'Echange* **407**: 13-16 hors-texte.
Pic M. 1923: Contribution à l'étude des Lycides. *L'Echange* **412**: 35-36 hors-texte.
Pic M. 1926a: Contribution à l'étude des Lycides. *L'Echange* **425**: 29-30 hors-texte.
Pic M. 1926b: Malacodermes exotiques. *L'Echange* **423**: 22 hors-texte.
Pic M. 1926c: Nouveaux Coléoptères du Globe. *Bull. Soc. Zool. Fr.* **51**: 454.
Pic M. 1930: Malacodermes exotiques. *L'Echange* **442**: 92 hors-texte.
Pic M. 1921–1922. Contribution à l’étude des Lycides. L’Echange, hors texte 37–38, 1–28.
Pic M. 1922. Nouveautés diverses. Mélanges exotico-entomologiques 35, 1–32.
Pic. M. 1924. Lycides africains nouveaux. Annali del Museo Civico di Storia Naturale di Genova 51, 161–164.
Pic M. 1924–1939. Malacodermes exotiques. L’Echange, hors texte 40–55, 1–472.
Pic M. 1925a. Diagnoses préliminaires de Lycides africains [Col.]. Bulletin du Muséum d’Histoire Naturelle. Pp. 184–187.
Pic. M. 1925b. Zoological results of the Swedish Expedition to Central Africa 1921. Insecta. 8. Anobiidae et Malacodermes. Arkiv för Zoologi. Ser.A. 17(5), 1–9.
Pic M. 1928a. Diagnose d’un nouveau Lycide africain [Col.]. Bulletin de la Société entomologique de France p. 112.
Pic M. 1928b. Note sur les Cladophorus [Col., Lycidae]. Bulletin de la Société entomologique de France pp. 199–200.

Pic. M. 1928c. Etudes sur les Lycides. Annali del Museo Civico di Storia Naturale di Genova 52, 116–129.
Pic M. 1928d. Pour aider á l'étude du genre Xylobanus Wa (Col.). Bulletin de la Société zoologique de France 53, 46–50.
Pic. M. 1931. Neue exotische Coleopteren (Malacodermata). Entomologischer Anzeiger. 11, 377–379.
Pic M. 1947. Coléoptères du Globe. L'Echange 63, 1–16.
Pic M. 1950. Coléoptères du Globe. L'Echange 66, 1–16.
Pic M. 1953. Coléoptères du Globe. L'Echange 69, 2–4. Pic, M. Contribution a l'étude des Lycides. L'Echange, Rev. Linn. 1921, 404–406, 1–12.
Pic, M. Malacodermes exotiques. L'Echange, Rev. Linn. 1923, 418–419, 1–84.
Pic, M. Contribution a l'étude des Lycides. L'Echange, Rev. Linn. 1922, 407–410, 13–28.
Potozkaja, V. A. 1981: Morphology and ecology of larvae of Coleopterous genera Aplatopterus Rtt. and Xylobanus C. O. Waterhouse (Coleoptera, Lycidae). *Entomol. Obozr*. **60**: 337-347 (in Russian).
Sklenarova, K.; Chesters, U.; Bocak, L. Phylogeography of poorly dispersing net-winged beetles: A role of drifting India in the origin of afrotropical and oriental fauna. PLoS ONE 2013, 8, e67957, doi:10.1371/journal.pone.0067957.
Sklenarova, K.; Kubecek, V.; Bocak, L. Subtribal classification of Metriorrhynchini (Insecta: Coleoptera: Lycidae): An integrative approach using molecular phylogeny and morphology of adults and larvae. Arthr. Syst. Phyl. 2014, 72, 37–54.
Thomson J. 1858. Voyage au Gabon. Histoire naturelle des Insectes et des Arachnides recueillis pendant un voyage fait au Gabon en 1856 et en 1857 par M. Henry C. Deyrolle sous les auspices de MM. Le Conte de Mnizech et James Thomson précédée de l'histoire de voyage. Deuxième Partie. Insectes. I. Ordre Coléoptères. Famille XVII. Lampyridae. Archives entomologiques ou recueil contenant des illustrations d'Insects nouveaux ou rares. 2, 76–79.
Walker F. 1858: Characters on some apparently undescribed Ceylon Insects. *Ann. Mag. Nat. Hist*. (3)**2**: 280-286.
Waterhouse, C.O. A Monograph of the Australian species of the Coleopterous family Lycidae. Trans. Entomol. Soc. Lond. 1877, 73–86.
Waterhouse, C.O. On the different forms occurring in the Coleopterous family Lycidæ, with descriptions of new genera and species. Trans. Entomol. Soc. Lond. 1878, 26, 95–118, doi:10.1111/j.1365- 2311.1878.tb01944.x.
Waterhouse, C.O. Illustration of the Typical Specimens of Coleoptera in the Collection of the British Museum. Part I.—Lycidae; British Museum: London, UK, 1879; p. 83.
Weiszenstein, M.; Bocak, L. A review of the genus Metanoeus from the Philippines (Coleoptera: Lycidae). Raffl. Bull. Zool. 2011, 59, 163–170.

**References (Supplementary methods section)**

Ababneh, F., Jermiin, L. S., Ma, Ch., Robinson, J. 2006 Matched-pairs tests of homogeneity with applications to homologous nucleotide sequences. Bioinformatics, 22(10), 1225–1231. doi:10.1093/bioinformatics/btl064
Bankevich A et al. 2012 SPAdes: A New Genome Assembly Algorithm and Its Applications to Single-Cell Sequencing. J. Comput. Biol. 19, 455–477. doi:10.1089/cmb.2012.0021
Bininda-Emonds, O.R. 2005 transAlign: using amino acids to facilitate the multiple alignment of protein-coding DNA sequences. BMC Bioinformatics 6, 156. doi:10.1186/1471-2105-6-156
Breinholt JW, Kawahara AY. 2013 Phylotranscriptomics: saturated third codon positions radically influence the estimation of trees based on next-gen data. Genome Biol. Evol. 5, 2082– 2092. doi:10.1093/gbe/evt157
Brower AVZ . 1994. Rapid morphological radiation and convergence among races of the butterfly Heliconius erato inferred from patterns of mitochondrial DNA evolution. PNAS 91: 6491–6495.

Chen S, Zhou Y, Chen, Y, Gu J. 2018 Fastp: an ultra-fast all-in-one FASTQ preprocessor. Bioinformatics 34, i884–i890. doi:10.1093/bioinformatics/bty560
Chernomor O, von Haeseler A, Minh, BQ. 2016 Terrace aware data structure for phylogenomic inference from supermatrices. Syst. Biol. 65, 997–1008. doi:10.1093/sysbio/syw037
Drummond, A.J., Suchard, M.A., Xie, D. & Rambaut, A. 2012 Bayesian phylogenetics with BEAUti and the BEAST 1.7. Molecular Biology and Evolution, 29, 1969–1973.
Fernandez R, Edgecombe GD, Giribet G. 2016 Exploring phylogenetic relationships within myriapoda and the effects of matrix composition and occupancy on phylogenomic reconstruction. Syst. Biol. 65, 871–889. doi:10.1093/sysbio/syw041
Hoang DT, Chernomor O, von Haeseler A, Minh BQ, Vinh LS. 2018 UFBoot2: improving the ultrafast bootstrap approximation. Mol. Biol. Evol. 35, 518–522. doi:10.1093/molbev/msx28
Kalyaanamoorthy S, Minh BQ, Wong TKF, von Haeseler A, Jermiin LS. 2017 ModelFinder: Fast model selection for accurate phylogenetic estimates. Nat. Methods 14, 587–589. doi:10.1038/NMETH.4285
Katoh K, Standley DM. 2013 MAFFT Multiple Sequence Alignment Software Version 7: Improvements in Performance and Usability. Mol. Biol. Evol. 30, 772–780. doi:10.1093/molbev/mst010
Keeling, C. I. et al. 2013. Draft genome of the mountain pine beetle, Dendroctonus ponderosae Hopkins, a major forest pest. Genome Biology, 14(3). doi:10.1186/gb-2013-14-3-r27
Kück P et al. 2010 Parametric and non-parametric masking of randomness in sequence alignments can be improved and leads to better resolved trees. Front. Zool. 7, 10. doi:10.1186/1742-9994-7-10
Kück P, Longo GC. 2014 FASconCAT-G: extensive functions for multiple sequence alignment preparations concerning phylogenetic studies. Front. Zool. B, 81. doi:10.1186/s12983-014-0081-x
Kusy D, Motyka M, Bocek M, Masek M, Bocak L. 2019 Phylogenomic analysis resolves the relationships among net-winged beetles (Coleoptera: Lycidae) and reveals the parallel evolution of morphological traits. Syst. Ent. 44, 911–925. doi:10.1111/syen.12363
Kusy D, Motyka M, Bocek M, Vogler AP, Bocak L. 2018 Genome sequences identify three families of Coleoptera as morphologically derived click beetles (Elateridae). Sci. Rep. 8, 17084. doi:10.1038/s41598-018-35328-0
Lanfear R, Frandsen PB, Wright AM, Senfeld T, Calcott B. 2017 PartitionFinder 2: New Methods for Selecting Partitioned Models of Evolution for Molecular and Morphological Phylogenetic Analyses. Mol. Biol. Evol. 34, 772–773. doi:10.1093/molbev/msw260
McKenna, D. D. et al. 2017 Genome of the Asian longhorned beetle (Anoplophora glabripennis), a globally significant invasive species, reveals key functional and evolutionary innovations at the beetle–plant interface. Genome Biology, 17(1), 227. doi:10.1186/s13059- 016-1088-8
McKenna, D. D., et al. 2019 The evolution and genomic basis of beetle diversity. PNAS, 116, 24729–24737. doi:10.1073/pnas.1909655116
Minh BQ et al. 2020 IQ-TREE 2: New Models and Efficient Methods for Phylogenetic Inference in the Genomic Era. Mol. Biol. Evol. 37, 1530–1534. doi:10.1093/molbev/msaa015
Misof B et al. 2013 Selecting informative subsets of sparse supermatrices increases the chance to find correct trees. BMC Bioinformatics 14, 348. doi:10.1186/1471-2105-14-348
Misof B et al. 2014 Phylogenomics resolves the timing and pattern of insect evolution. Science 346, 763–767. doi:10.1126/science.1257570
Misof B, Misof KA. 2009 A Monte Carlo approach successfully identifies randomness in multiple sequence alignments: a more objective means of data exclusion. Syst. Biol. 58, 21–34. doi:10.1093/sysbio/syp006
Naser-Khdour S, Minh BQ, Zhang W, Stone SA, Lanfear R. 2019 The Prevalence and Impact of Model Violations in Phylogenetic Analysis. Genome Biol. Evol. 11, 3341–3352. doi:10.1093/gbe/evz193
Peters RS et al. 2017 Evolutionary history of the Hymenoptera. Curr. Biol. 27, 1013–1018. doi:10.1016/j.cub.2017.01.027

Petersen M et al. 2017 Orthograph: a versatile tool for mapping coding nucleotide sequences to clusters of orthologous genes. BMC Bioinformatics 18, 111. doi:10.1186/s12859-017-1529-8
Poelchau, M. et al. 2014 The i5k Workspace@NAL—enabling genomic data access, visualization and curation of arthropod genomes. Nucleic Acids Research, 43(D1). doi:10.1093/nar/gku983
Richards, S. et al. 2008 The genome of the model beetle and pest Tribolium castaneum. Nature, 452(7190), 949-955. doi:10.1038/nature06784
Shelton, J. M. et al. 2015 Tools and pipelines for BioNano data: molecule assembly pipeline and FASTA super scaffolding tool. BMC Genomics, 16(1). doi:10.1186/s12864-015-1911-8
Si Q. L., Gascuel, O. 2008 An Improved General Amino Acid Replacement Matrix, Molecular Biology and Evolution, Volume 25, Issue 7, July, Pages 1307–1320, doi:10.1093/molbev/msn067
Stanke M, Waack S. 2003 Gene prediction with a hidden Markov model and a new intron submodel. Bioinformatics 19, 215–225. doi:10.1093/bioinformatics/btg1080
Strimmer, K. & von Haeseler, A. 1997. Likelihood-mapping: a simple method to visualize phylogenetic content of a sequence alignment. PNAS, 94, 6815–6819.
Suyama M, Torrents D, Bork P. 2006 PAL2NAL: robust conversion of protein sequence alignments into the corresponding codon alignments. Nucl. Acids Res. 34, W609–W612. doi:10.1093/nar/gkl315
Vasilikopoulos A et al. 2019 Phylogenomics of the superfamily Dytiscoidea (Coleoptera: Adephaga) with an evaluation of phylogenetic conflict and systematic error. Mol. Phyl. Evol. 135, 270–285. doi:10.1016/j.ympev.2019.02.022
Waterhouse RM et al. 2017 BUSCO Applications from Quality Assessments to Gene Prediction and Phylogenomics. Mol. Biol. Evol. 35, 543–548. doi:10.1093/molbev/msx319
Xie, Y. et al. 2014 Trans: de novo transcriptome assembly with short RNA-Seq reads. Bioinformatics, 30, 1660–1666. doi:10.1093/bioinformatics/btu077
Zdobnov EM et al. 2016 OrthoDBv9.1: cataloging evolutionary and functional annotations for animal, fungal, plant, archaeal, bacterial and viral orthologs. Nucl. Acids Res. 45, D744–D749. doi:10.1093/nar/gkw1119
Zhang, J., Lindsey, A.R.I., Peters, R.S. et al. (2020) Conflicting signal in transcriptomic markers leads to a poorly resolved backbone phylogeny of chalcidoid wasps. Sys. Ento. 45, 783–802 doi:/10.1111/syen.12427.

Table S1. Detailed information on regional sampling.

| Region | Metriorrhynchina # of specimens | Cautirina # of specimens | Metanoeina # of specimens | Metriorrhynchini # of specimens |
|---|---|---|---|---|
| Australian region | 4489 | - | - | 4489 |
| Australian continent | 3964 | - | - | 3964 |
| Australia | 475 | - | - | 475 |
| New South Wales | 64 | - | - | 64 |
| Northern Territory | 18 | - | - | 18 |
| Queensland | 382 | - | - | 382 |
| South Australia | 1 | - | - | 1 |
| Western Australia | 10 | - | - | 10 |
| New Zealand | 1 | - | - | 1 |
| New Guinea | 3461 | - | - | 3461 |
| Solomon Islands | 19 | - | - | 19 |
| Aru Islands | 8 | - | - | 8 |
| Wallacea | 525 | 15 | - | 540 |
| Maluku-Buru | 15 | - | - | 15 |
| Halmahera | 169 | - | - | 169 |
| Sulawesi | 341 | 15 | - | 356 |
| Oriental region | | | | |
| Philippines | 48 | 37 | 14 | 99 |
| Mindanao | 11 | 3 | 9 | 23 |
| Negros | 2 | - | - | 2 |
| Palawan | 29 | 34 | 5 | 68 |
| Sibuyan | 4 | - | - | 4 |
| Luzon | 2 | - | - | 2 |
| Continental Asia | 230 | 1264 | 99 | 1593 |
| Malaya | 34 | 270 | 8 | 348 |
| Java | 21 | 45 | 6 | 72 |
| Bali | 2 | 1 | - | 3 |
| Sumatra | 56 | 201 | 26 | 283 |
| Borneo | 82 | 253 | 22 | 357 |
| Cambodia | 7 | 13 | - | 20 |
| Indo-Burma | 23 | 145 | 10 | 178 |
| China incl. Taiwan | 5 | 100 | 23 | 128 |
| India | - | 59 | 0 | 59 |
| Japan | - | 177 | 4 | 181 |
| Afrotropical region | - | 233 | - | 233 |
| Africa | - | 175 | - | 175 |
| Madagascar | - | 58 | - | 58 |
| Total | 4767 | 1549 | 113 | 6429 |

### Table S2. List of material for phylogenomic analyses.

**Ingroup**

| **Voucher** | **Species** | | **Clade** | **Geographic origin** | **Data source** | **Data type** | **Tissue type** |
|---|---|---|---|---|---|---|---|
| R18010 | *Metriorrhynchus* s. l. | Metriorrhynchina | Porrostomine | New Guinea | This study | ILLUMINA RNA-seq PE-reads | Whole animal |
| R18013 | *Metriorrhynchus* sp. | Metriorrhynchina | Porrostomine | New Guinea | This study | ILLUMINA RNA-seq PE-reads | Whole animal |
| R20001 | *Metriorrhynchus philippinensis* | Metriorrhynchina | Porrostomine | Philippines | This study | ILLUMINA RNA-seq PE-reads | Whole animal |
| R18018 | *Metriorrhynchus* s. l. | Metriorrhynchina | Porrostomine | New Guinea | This study | ILLUMINA RNA-seq PE-reads | Whole animal |
| R18020 | *Metriorrhynchus* s. l. | Metriorrhynchina | Porrostomine | New Guinea | This study | ILLUMINA RNA-seq PE-reads | Whole animal |
| 1_kite | *Porrostoma rhipidium* | Metriorrhynchina | Porrostomine | Australia | McKenna *et al*., 2019 | ILLUMINA RNA-seq PE-reads | Whole animal |
| R18001 | *Cladophorus* sp. | Metriorrhynchina | Cladophorine | New Guinea | This study | ILLUMINA RNA-seq PE-reads | Whole animal |
| R18009 | *Cladophorus* sp. | Metriorrhynchina | Cladophorine | New Guinea | This study | ILLUMINA RNA-seq PE-reads | Whole animal |
| R18007 | *Pseudodontocerus* sp. | Metriorrhynchina | Cladophorine | New Guinea | This study | ILLUMINA RNA-seq PE-reads | Whole animal |
| R18012 | *Ditua* s. l. | Metriorrhynchina | Cladophorine | New Guinea | This study | ILLUMINA RNA-seq PE-reads | Whole animal |
| R18025 | *Ditua* s. l. | Metriorrhynchina | Cladophorine | New Guinea | This study | ILLUMINA RNA-seq PE-reads | Whole animal |
| R18028 | *Ditua* s. l. | Metriorrhynchina | Cladophorine | New Guinea | This study | ILLUMINA RNA-seq PE-reads | Whole animal |
| R18003 | *Ditua* s. l. | Metriorrhynchina | Cladophorine | New Guinea | This study | ILLUMINA RNA-seq PE-reads | Whole animal |
| R18016 | *Ditua* s. l. | Metriorrhynchina | Cladophorine | New Guinea | This study | ILLUMINA RNA-seq PE-reads | Whole animal |
| R18005 | *Ditua* s. l. | Metriorrhynchina | Cladophorine | New Guinea | This study | ILLUMINA RNA-seq PE-reads | Whole animal |
| R18015 | *Ditua* s. l. | Metriorrhynchina | Cladophorine | New Guinea | This study | ILLUMINA RNA-seq PE-reads | Whole animal |
| R18006 | *Ditua* s. l. | Metriorrhynchina | Cladophorine | New Guinea | This study | ILLUMINA RNA-seq PE-reads | Whole animal |
| R18002 | *Microtrichalus* sp. | Metriorrhynchina | Trichaline | New Guinea | This study | ILLUMINA RNA-seq PE-reads | Whole animal |
| R18024 | *Eniclases* sp. | Metriorrhynchina | Trichaline | New Guinea | This study | ILLUMINA RNA-seq PE-reads | Whole animal |
| R18022 | *Diatrichalus* sp. | Metriorrhynchina | Trichaline | New Guinea | This study | ILLUMINA RNA-seq PE-reads | Whole animal |
| R18026 | Trichaline sp. | Metriorrhynchina | Trichaline | New Guinea | This study | ILLUMINA RNA-seq PE-reads | Whole animal |
| JB0085 | Trichaline sp. | Metriorrhynchina | Trichaline | New Guinea | This study | ILLUMINA WGS-seq PE-reads | Thoracic muscles |
| R18004 | *Procautires* sp. | Metriorrhynchina | Procautirine | New Guinea | This study | ILLUMINA RNA-seq PE-reads | Whole animal |
| R18017 | *Procautires* sp. | Metriorrhynchina | Procautirine | New Guinea | This study | ILLUMINA RNA-seq PE-reads | Whole animal |
| R18021 | *Procautires* sp. | Metriorrhynchina | Procautirine | New Guinea | This study | ILLUMINA RNA-seq PE-reads | Whole animal |
| R18023 | *Procautires* sp. | Metriorrhynchina | Procautirine | New Guinea | This study | ILLUMINA RNA-seq PE-reads | Whole animal |
| R18037 | *Broxylus* sp. | Metriorrhynchina | Leptotrichaline | Sulawesi | This study | ILLUMINA RNA-seq PE-reads | Whole animal |
| R18030 | *Leptotrichalus* sp. | Metriorrhynchina | Leptotrichaline | Sulawesi | This study | ILLUMINA RNA-seq PE-reads | Whole animal |
| R18034 | *Wakarumbia* sp. | Metriorrhynchina | Leptotrichaline | Sulawesi | This study | ILLUMINA RNA-seq PE-reads | Whole animal |
| R18036 | *Sulabanus* sp. | Metriorrhynchina | Leptotrichaline | Sulawesi | This study | ILLUMINA RNA-seq PE-reads | Whole animal |
| R18041 | *Sulabanus* sp. | Metriorrhynchina | Leptotrichaline | Sulawesi | This study | ILLUMINA RNA-seq PE-reads | Whole animal |
| R18040 | *Mangkutanus* sp. | Metriorrhynchina | Leptotrichaline | Sulawesi | Kusy *et al*., 2019 | ILLUMINA RNA-seq PE-reads | Whole animal |
| R18039 | *Xylobanus* sp. | Cautirina | | Sulawesi | This study | ILLUMINA RNA-seq PE-reads | Whole animal |
| AJ0013 | *Cautires communis* | Cautirina | | Malaysia | Kusy *et al*., 2019 | ILLUMINA WGS-seq PE-reads | Thoracic muscles |
| G19002 | *Metanoeus* sp. | Metanoeina | | Malaysia | This study | ILLUMINA WGS-seq PE-reads | Thoracic muscles |

**Outgroup**

| | | | | | | | |
|---|---|---|---|---|---|---|---|
| | *Dilophotes* sp. | | | Malaysia | Kusy *et al*., 2019 | ILLUMINA RNA-seq PE-reads | Whole animal |
| | *Dihammatus* sp. | | | Malaysia | Kusy *et al*., 2019 | ILLUMINA RNA-seq PE-reads | Whole animal |
| | *Lycoprogentes* sp. | | | Malaysia | Kusy *et al*., 2019 | ILLUMINA RNA-seq PE-reads | Whole animal |
| | *Libnetis* sp. | | | Malaysia | Kusy *et al*., 2019 | ILLUMINA RNA-seq PE-reads | Whole animal |
| | *Platerodrilus* sp. | | | Malaysia | Kusy *et al*., 2019 | ILLUMINA RNA-seq PE-reads | Whole animal |
| | *Lyropaeus optabilis* | | | Malaysia | Kusy *et al*., 2019 | ILLUMINA RNA-seq PE-reads | Whole animal |
| | *Antennolycus constrictus* | | | Malaysia | Kusy *et al*., 2019 | ILLUMINA RNA-seq PE-reads | Whole animal |

Table S3. Completeness of the mtDNA dataset used for IQtree analysis.
Abbreviations: #INF - number of gene fragments, #MIS - number of missing gene fragments.

| Sample | #INF | #MIS |
|---|---|---|
| A00017 | 3 | 0 |
| A00018 | 3 | 0 |
| A00019 | 3 | 0 |
| A00020 | 1 | 2 |
| A00021 | 1 | 2 |
| A00022 | 2 | 1 |
| A00023 | 2 | 1 |
| A00024 | 3 | 0 |
| A00025 | 3 | 0 |
| A00026 | 2 | 1 |
| A00027 | 2 | 1 |
| A00028 | 1 | 2 |
| A00029 | 3 | 0 |
| A00030 | 3 | 0 |
| A00031 | 3 | 0 |
| A00031 | 3 | 0 |
| A00032 | 3 | 0 |
| A00032 | 3 | 0 |
| A00033 | 2 | 1 |
| A00034 | 3 | 0 |
| A00035 | 3 | 0 |
| A00036 | 3 | 0 |
| A00037 | 3 | 0 |
| A00038 | 3 | 0 |
| A00039 | 3 | 0 |
| A00040 | 3 | 0 |
| A00041 | 3 | 0 |
| A00041 | 3 | 0 |
| A00042 | 2 | 1 |
| A00043 | 3 | 0 |
| A00044 | 3 | 0 |
| A00045 | 3 | 0 |
| A00046 | 3 | 0 |
| A00048 | 2 | 1 |
| A00049 | 3 | 0 |
| A00050 | 2 | 1 |
| A00052 | 3 | 0 |
| A00053 | 3 | 0 |
| A00054 | 2 | 1 |
| A00055 | 3 | 0 |
| A00056 | 3 | 0 |
| A00057 | 2 | 1 |
| A00058 | 3 | 0 |
| A00059 | 3 | 0 |
| A00061 | 3 | 0 |
| A00062 | 3 | 0 |
| A00063 | 2 | 1 |
| A00064 | 2 | 1 |
| A00065 | 2 | 1 |
| A00066 | 3 | 0 |
| A00067 | 3 | 0 |
| A00068 | 3 | 0 |
| A00069 | 3 | 0 |
| A00070 | 3 | 0 |
| A00071 | 3 | 0 |
| A00072 | 2 | 1 |
| A00074 | 1 | 2 |
| A00075 | 3 | 0 |
| A00076 | 3 | 0 |
| A00077 | 3 | 0 |
| A00077 | 3 | 0 |
| A00078 | 3 | 0 |
| A00079 | 3 | 0 |
| A00080 | 2 | 1 |
| A00081 | 3 | 0 |
| A00082 | 2 | 1 |
| A00083 | 3 | 0 |
| A00084 | 3 | 0 |
| A00085 | 3 | 0 |
| A00086 | 3 | 0 |
| A00087 | 3 | 0 |
| A00088 | 3 | 0 |
| A00089 | 3 | 0 |
| A00090 | 3 | 0 |
| A00090 | 3 | 0 |
| A00091 | 3 | 0 |
| A00092 | 3 | 0 |
| A00093 | 3 | 0 |
| A00094 | 3 | 0 |
| A00095 | 2 | 1 |
| A00096 | 2 | 1 |
| A00097 | 3 | 0 |
| A00098 | 3 | 0 |
| A00099 | 3 | 0 |
| A00100 | 3 | 0 |
| A00101 | 1 | 2 |
| A00102 | 2 | 1 |
| A00103 | 1 | 2 |
| A00104 | 2 | 1 |
| A00105 | 2 | 1 |
| A00106 | 3 | 0 |
| A00107 | 1 | 2 |
| A00109 | 3 | 0 |
| A00110 | 2 | 1 |
| A00111 | 3 | 0 |
| A00112 | 3 | 0 |
| A00113 | 3 | 0 |
| A00114 | 2 | 1 |
| A00115 | 2 | 1 |
| A00116 | 2 | 1 |
| A00117 | 2 | 1 |
| A00118 | 3 | 0 |
| A00120 | 2 | 1 |
| A00121 | 3 | 0 |
| A00122 | 3 | 0 |
| A00123 | 3 | 0 |
| A00124 | 3 | 0 |
| A00125 | 3 | 0 |
| A00126 | 3 | 0 |
| A00127 | 3 | 0 |
| A00128 | 3 | 0 |
| A00129 | 2 | 1 |
| A00130 | 2 | 1 |
| A00131 | 1 | 2 |
| A00132 | 3 | 0 |
| A00133 | 2 | 1 |
| A00134 | 3 | 0 |
| A00136 | 3 | 0 |
| A00137 | 2 | 1 |
| A00138 | 3 | 0 |
| A00139 | 3 | 0 |
| A00140 | 3 | 0 |
| A00141 | 1 | 2 |
| A00142 | 3 | 0 |
| A00143 | 3 | 0 |
| A00144 | 3 | 0 |
| A00145 | 3 | 0 |
| A00146 | 3 | 0 |
| A00148 | 3 | 0 |
| A00148 | 3 | 0 |
| A00149 | 3 | 0 |
| A00152 | 2 | 1 |
| A00153 | 2 | 1 |
| A00154 | 3 | 0 |
| A00155 | 3 | 0 |
| A00156 | 3 | 0 |
| A00157 | 2 | 1 |
| A00158 | 3 | 0 |
| A00159 | 2 | 1 |
| A00161 | 2 | 1 |
| A00162 | 3 | 0 |
| A00163 | 3 | 0 |
| A00164 | 3 | 0 |
| A00165 | 3 | 0 |
| A00166 | 3 | 0 |
| A00167 | 3 | 0 |
| A00168 | 3 | 0 |
| A00169 | 3 | 0 |
| A00170 | 3 | 0 |
| A00171 | 3 | 0 |
| A00172 | 2 | 1 |
| A00173 | 3 | 0 |
| A00174 | 3 | 0 |
| A00175 | 3 | 0 |
| A00176 | 3 | 0 |
| A00178 | 3 | 0 |
| A00179 | 3 | 0 |
| A00180 | 3 | 0 |
| A00181 | 3 | 0 |
| A00182 | 2 | 1 |
| A00184 | 2 | 1 |
| A00185 | 2 | 1 |
| A00186 | 2 | 1 |
| A00187 | 3 | 0 |
| A00188 | 2 | 1 |
| A00189 | 3 | 0 |
| A00190 | 2 | 1 |
| A00192 | 2 | 1 |
| A00194 | 2 | 1 |
| A00195 | 3 | 0 |
| A00198 | 2 | 1 |
| A00199 | 3 | 0 |
| A00200 | 3 | 0 |
| A00201 | 3 | 0 |
| A00202 | 3 | 0 |
| A00203 | 3 | 0 |
| A00204 | 3 | 0 |
| A00205 | 3 | 0 |
| A00206 | 2 | 1 |
| A00207 | 1 | 2 |
| A00208 | 3 | 0 |
| A00209 | 3 | 0 |
| A00210 | 3 | 0 |
| A00211 | 3 | 0 |
| A00212 | 3 | 0 |
| A00213 | 3 | 0 |
| A00214 | 3 | 0 |
| A00215 | 3 | 0 |
| A00217 | 3 | 0 |
| A00219 | 3 | 0 |
| A00220 | 3 | 0 |
| A00221 | 3 | 0 |
| A00222 | 3 | 0 |
| A00223 | 3 | 0 |
| A00225 | 3 | 0 |
| A00227 | 3 | 0 |
| A00228 | 3 | 0 |
| A00229 | 3 | 0 |
| A00232 | 3 | 0 |
| A00235 | 3 | 0 |
| A00236 | 3 | 0 |
| A00237 | 3 | 0 |
| A00238 | 3 | 0 |
| A00239 | 3 | 0 |
| A00240 | 3 | 0 |
| A00241 | 3 | 0 |
| A00242 | 3 | 0 |
| A00243 | 3 | 0 |
| A00247 | 3 | 0 |
| A00248 | 3 | 0 |
| A00249 | 3 | 0 |
| A00251 | 3 | 0 |
| A00252 | 3 | 0 |
| A00253 | 3 | 0 |
| A00255 | 3 | 0 |
| A00256 | 3 | 0 |
| A00257 | 3 | 0 |
| A00258 | 3 | 0 |
| A00259 | 3 | 0 |
| A00260 | 3 | 0 |
| A00261 | 3 | 0 |
| A00262 | 3 | 0 |
| A00263 | 3 | 0 |
| A00265 | 2 | 1 |
| A00266 | 3 | 0 |
| A00267 | 3 | 0 |
| A00268 | 3 | 0 |
| A00269 | 3 | 0 |
| A00270 | 3 | 0 |
| A00271 | 3 | 0 |
| A00273 | 3 | 0 |
| A00274 | 2 | 1 |
| A00275 | 3 | 0 |
| A00276 | 3 | 0 |
| A00277 | 3 | 0 |
| A00278 | 3 | 0 |
| A00279 | 2 | 1 |
| A00280 | 3 | 0 |
| A00281 | 3 | 0 |
| A00282 | 3 | 0 |
| A00283 | 3 | 0 |
| A00284 | 3 | 0 |
| A00285 | 3 | 0 |
| A00286 | 3 | 0 |
| A00287 | 3 | 0 |
| A00289 | 3 | 0 |
| A00290 | 3 | 0 |
| A00291 | 3 | 0 |
| A00292 | 2 | 1 |
| A00296 | 3 | 0 |
| A00297 | 3 | 0 |
| A00298 | 3 | 0 |
| A00299 | 3 | 0 |
| A00300 | 3 | 0 |
| A00301 | 3 | 0 |
| A00304 | 3 | 0 |
| A00305 | 3 | 0 |
| A00306 | 3 | 0 |
| A00307 | 3 | 0 |
| A00308 | 1 | 2 |
| A00311 | 3 | 0 |
| A00312 | 3 | 0 |
| A00313 | 3 | 0 |
| A00314 | 3 | 0 |
| A00315 | 3 | 0 |
| A00316 | 3 | 0 |
| A00317 | 3 | 0 |
| A00318 | 3 | 0 |
| A00319 | 3 | 0 |
| A00321 | 3 | 0 |
| A00322 | 3 | 0 |
| A00326 | 2 | 1 |
| A00327 | 3 | 0 |
| A00329 | 3 | 0 |
| A00330 | 3 | 0 |
| A00331 | 3 | 0 |
| A00332 | 3 | 0 |
| A00333 | 3 | 0 |
| A00334 | 3 | 0 |
| A00335 | 3 | 0 |
| A00336 | 3 | 0 |
| A00337 | 2 | 1 |
| A00339 | 3 | 0 |
| A00340 | 2 | 1 |
| A00341 | 3 | 0 |
| A00342 | 3 | 0 |
| A00343 | 3 | 0 |
| A00344 | 3 | 0 |
| A00347 | 3 | 0 |
| A00348 | 3 | 0 |
| A00349 | 3 | 0 |
| A00350 | 3 | 0 |
| A00351 | 3 | 0 |
| A00352 | 2 | 1 |
| A00353 | 3 | 0 |
| A00355 | 3 | 0 |
| A00356 | 2 | 1 |
| A00357 | 3 | 0 |
| A00358 | 3 | 0 |
| A00359 | 3 | 0 |
| A00360 | 3 | 0 |
| A00361 | 3 | 0 |
| A00362 | 3 | 0 |
| A00363 | 3 | 0 |
| A00364 | 3 | 0 |
| A00365 | 2 | 1 |
| A00366 | 3 | 0 |
| A00367 | 3 | 0 |
| A00368 | 3 | 0 |
| A00369 | 3 | 0 |
| A00370 | 3 | 0 |
| A00371 | 3 | 0 |
| A00372 | 3 | 0 |
| A00373 | 3 | 0 |
| A00374 | 3 | 0 |
| A00375 | 3 | 0 |
| A00376 | 3 | 0 |
| A00377 | 3 | 0 |
| A00378 | 3 | 0 |
| A00379 | 3 | 0 |
| A00380 | 3 | 0 |
| A00381 | 3 | 0 |
| A00382 | 3 | 0 |
| A00383 | 3 | 0 |
| A00384 | 3 | 0 |
| A00385 | 3 | 0 |
| A00386 | 3 | 0 |
| A00387 | 3 | 0 |
| A00389 | 3 | 0 |
| A00390 | 3 | 0 |
| A00391 | 3 | 0 |
| A00393 | 3 | 0 |
| A00394 | 3 | 0 |
| A00395 | 2 | 1 |
| A00396 | 3 | 0 |
| A00397 | 3 | 0 |
| A00399 | 3 | 0 |
| A00400 | 3 | 0 |
| A00401 | 3 | 0 |
| A00402 | 3 | 0 |
| A00403 | 3 | 0 |
| A00404 | 2 | 1 |
| A00405 | 2 | 1 |
| A00406 | 2 | 1 |
| A00407 | 3 | 0 |
| A00408 | 2 | 1 |
| A00409 | 3 | 0 |
| A00410 | 3 | 0 |
| A00411 | 2 | 1 |
| A00412 | 2 | 1 |
| A00413 | 3 | 0 |
| A00414 | 3 | 0 |
| A00415 | 3 | 0 |
| A00416 | 3 | 0 |
| A00417 | 3 | 0 |
| A00418 | 3 | 0 |
| A00419 | 3 | 0 |
| A00420 | 3 | 0 |
| A00421 | 3 | 0 |
| A00422 | 3 | 0 |
| A00423 | 3 | 0 |
| A00424 | 3 | 0 |
| A00425 | 3 | 0 |
| A00426 | 3 | 0 |
| A00427 | 3 | 0 |
| A00428 | 3 | 0 |
| A00429 | 3 | 0 |
| A00430 | 3 | 0 |
| A00431 | 3 | 0 |
| A00432 | 3 | 0 |
| A00433 | 3 | 0 |
| A00434 | 3 | 0 |
| A00435 | 3 | 0 |
| A00437 | 2 | 1 |
| A00438 | 3 | 0 |
| A00439 | 2 | 1 |
| A00440 | 3 | 0 |
| A00441 | 3 | 0 |
| A00442 | 3 | 0 |
| A00443 | 3 | 0 |
| A00444 | 3 | 0 |
| A00445 | 3 | 0 |
| A00446 | 3 | 0 |
| A00447 | 3 | 0 |
| A00448 | 3 | 0 |
| A00449 | 3 | 0 |
| A00450 | 3 | 0 |
| A00451 | 3 | 0 |
| A00452 | 2 | 1 |
| A00453 | 2 | 1 |
| A00454 | 3 | 0 |
| A00455 | 3 | 0 |
| A00456 | 3 | 0 |
| A00457 | 3 | 0 |
| A00458 | 3 | 0 |
| A00459 | 3 | 0 |
| A00652 | 2 | 1 |
| A00653 | 2 | 1 |
| A00655 | 2 | 1 |
| A00656 | 3 | 0 |
| A00673 | 3 | 0 |
| A00674 | 1 | 2 |
| A00675 | 1 | 2 |
| A00676 | 1 | 2 |
| A00677 | 1 | 2 |
| A00678 | 1 | 2 |
| A00679 | 3 | 0 |
| A00680 | 1 | 2 |
| A00681 | 1 | 2 |
| A00682 | 3 | 0 |
| A00683 | 3 | 0 |
| A00684 | 1 | 2 |
| A00685 | 1 | 2 |
| A00686 | 1 | 2 |
| A00687 | 1 | 2 |
| A00688 | 1 | 2 |
| A00689 | 2 | 1 |
| A00690 | 3 | 0 |
| A00691 | 3 | 0 |
| A00692 | 3 | 0 |
| A00693 | 1 | 2 |
| A00694 | 1 | 2 |
| A00695 | 3 | 0 |
| A00696 | 3 | 0 |
| A00697 | 1 | 2 |
| A00698 | 3 | 0 |
| A00699 | 3 | 0 |
| A00700 | 1 | 2 |
| A00701 | 2 | 1 |
| A00702 | 3 | 0 |
| A00703 | 1 | 2 |
| A00704 | 1 | 2 |
| A00705 | 1 | 2 |
| A00706 | 1 | 2 |
| A00707 | 1 | 2 |
| A00708 | 1 | 2 |
| A00709 | 1 | 2 |
| A00710 | 3 | 0 |
| A00711 | 1 | 2 |
| A00712 | 2 | 1 |
| A00713 | 2 | 1 |
| A00714 | 1 | 2 |
| A00715 | 1 | 2 |
| A00716 | 1 | 2 |
| A00717 | 3 | 0 |
| A00718 | 1 | 2 |
| A00719 | 1 | 2 |
| A00720 | 3 | 0 |
| A00721 | 2 | 1 |
| A00722 | 2 | 1 |
| A00723 | 3 | 0 |
| A00724 | 3 | 0 |
| A00725 | 1 | 2 |
| A00726 | 3 | 0 |
| A00727 | 3 | 0 |
| A00728 | 3 | 0 |
| A00729 | 2 | 1 |
| A00731 | 2 | 1 |
| A00732 | 2 | 1 |
| A00733 | 1 | 2 |
| A00734 | 1 | 2 |
| A00735 | 1 | 2 |
| A00736 | 3 | 0 |
| A00737 | 3 | 0 |
| A00739 | 3 | 0 |
| A00740 | 1 | 2 |
| A00741 | 1 | 2 |
| A00742 | 1 | 2 |
| A00743 | 1 | 2 |
| A00744 | 2 | 1 |
| A00745 | 3 | 0 |
| A00746 | 1 | 2 |
| A00747 | 1 | 2 |
| A00748 | 3 | 0 |
| A00749 | 1 | 2 |
| A00750 | 1 | 2 |
| A00751 | 3 | 0 |
| A00752 | 1 | 2 |
| A00753 | 1 | 2 |
| A00754 | 1 | 2 |
| A00755 | 1 | 2 |
| A00757 | 1 | 2 |
| A00758 | 1 | 2 |
| A00759 | 2 | 1 |
| A00760 | 3 | 0 |
| A00761 | 1 | 2 |
| A00762 | 3 | 0 |
| A00763 | 3 | 0 |
| A00764 | 1 | 2 |
| A00765 | 1 | 2 |
| A00766 | 1 | 2 |
| A00767 | 2 | 1 |
| A00768 | 2 | 1 |
| A00769 | 1 | 2 |
| A00770 | 3 | 0 |
| A00771 | 1 | 2 |
| A00772 | 3 | 0 |
| A00773 | 1 | 2 |
| A00774 | 1 | 2 |
| A00775 | 1 | 2 |
| A00776 | 3 | 0 |
| A00777 | 2 | 1 |
| A00778 | 3 | 0 |
| A00779 | 3 | 0 |
| A00780 | 1 | 2 |
| A00781 | 1 | 2 |
| A00782 | 1 | 2 |
| A00783 | 3 | 0 |
| A00784 | 1 | 2 |
| A00785 | 3 | 0 |
| A00786 | 1 | 2 |
| A00787 | 1 | 2 |
| A00788 | 1 | 2 |
| A00790 | 3 | 0 |
| A00791 | 1 | 2 |
| A00793 | 1 | 2 |
| A00794 | 1 | 2 |
| A00795 | 1 | 2 |
| A00797 | 1 | 2 |
| A00798 | 1 | 2 |
| A00799 | 3 | 0 |
| A00800 | 1 | 2 |
| A00801 | 1 | 2 |
| A00802 | 1 | 2 |
| A00803 | 1 | 2 |
| A00804 | 1 | 2 |
| A00805 | 1 | 2 |
| A00806 | 1 | 2 |
| A00809 | 1 | 2 |
| A00810 | 1 | 2 |
| A00811 | 1 | 2 |
| A00812 | 1 | 2 |
| A00813 | 1 | 2 |
| A00814 | 1 | 2 |
| A00815 | 3 | 0 |
| A00816 | 1 | 2 |
| A00817 | 1 | 2 |
| A00819 | 2 | 1 |
| A00820 | 1 | 2 |
| A00821 | 3 | 0 |
| A00822 | 1 | 2 |
| A00823 | 1 | 2 |
| A00824 | 3 | 0 |
| A00825 | 1 | 2 |
| A00826 | 1 | 2 |
| A00827 | 1 | 2 |
| A00828 | 3 | 0 |
| A00829 | 2 | 1 |
| A00830 | 1 | 2 |
| A00832 | 1 | 2 |
| A00833 | 1 | 2 |
| A00834 | 1 | 2 |
| A00835 | 1 | 2 |
| A00836 | 1 | 2 |
| A00837 | 1 | 2 |
| A00838 | 1 | 2 |
| A00839 | 3 | 0 |
| A00840 | 1 | 2 |
| A00841 | 1 | 2 |
| A00842 | 1 | 2 |
| A00843 | 3 | 0 |
| A00844 | 3 | 0 |
| A00846 | 2 | 1 |
| A00848 | 1 | 2 |
| A00849 | 3 | 0 |
| A00850 | 3 | 0 |
| A00851 | 3 | 0 |
| A00852 | 1 | 2 |
| A00855 | 2 | 1 |
| A00856 | 2 | 1 |
| A00857 | 1 | 2 |
| A00859 | 2 | 1 |
| A00860 | 2 | 1 |
| A00861 | 2 | 1 |
| A00862 | 3 | 0 |
| A00863 | 3 | 0 |
| A00865 | 1 | 2 |
| A00867 | 1 | 2 |
| A00868 | 1 | 2 |
| A00869 | 2 | 1 |
| A00870 | 1 | 2 |
| A00872 | 1 | 2 |
| A00873 | 1 | 2 |
| A00874 | 1 | 2 |
| A00876 | 1 | 2 |
| A00877 | 1 | 2 |
| A00878 | 3 | 0 |
| A00879 | 3 | 0 |
| A00880 | 1 | 2 |
| A00881 | 1 | 2 |
| A00882 | 2 | 1 |
| A00883 | 2 | 1 |
| A00884 | 1 | 2 |
| A00885 | 3 | 0 |
| A00887 | 1 | 2 |
| A00888 | 2 | 1 |
| A00889 | 1 | 2 |
| A00890 | 1 | 2 |
| A00891 | 3 | 0 |
| A00892 | 1 | 2 |
| A00893 | 1 | 2 |
| A00894 | 1 | 2 |
| A00896 | 3 | 0 |
| A00897 | 2 | 1 |
| A00898 | 3 | 0 |
| A00899 | 1 | 2 |
| A00900 | 1 | 2 |
| A00901 | 2 | 1 |
| A00902 | 1 | 2 |
| A00903 | 1 | 2 |
| A00904 | 1 | 2 |
| A00905 | 1 | 2 |
| A00906 | 3 | 0 |
| A00907 | 1 | 2 |
| A00909 | 3 | 0 |
| A00910 | 3 | 0 |
| A00911 | 1 | 2 |
| A00912 | 3 | 0 |
| A00913 | 1 | 2 |
| A00914 | 1 | 2 |
| A00915 | 1 | 2 |
| A00916 | 1 | 2 |
| A00917 | 1 | 2 |
| A00918 | 1 | 2 |
| A00919 | 1 | 2 |
| A00920 | 1 | 2 |
| A00921 | 1 | 2 |
| A00922 | 3 | 0 |
| A00923 | 1 | 2 |
| A00924 | 3 | 0 |
| A00925 | 1 | 2 |
| A00926 | 1 | 2 |
| A00928 | 1 | 2 |
| A00929 | 1 | 2 |
| A00930 | 2 | 1 |
| A00931 | 1 | 2 |
| A00932 | 1 | 2 |
| A00933 | 3 | 0 |
| A00934 | 1 | 2 |
| A00935 | 1 | 2 |
| A00936 | 1 | 2 |
| A00937 | 1 | 2 |
| A00938 | 1 | 2 |
| A00940 | 1 | 2 |
| A00941 | 2 | 1 |
| A00942 | 3 | 0 |
| A00943 | 1 | 2 |
| A00944 | 1 | 2 |
| A00945 | 1 | 2 |
| A00946 | 1 | 2 |
| A00947 | 2 | 1 |
| A00948 | 2 | 1 |
| A00949 | 3 | 0 |
| A00950 | 2 | 1 |
| A00951 | 1 | 2 |
| A00952 | 3 | 0 |
| A00953 | 1 | 2 |
| A00954 | 2 | 1 |
| A00955 | 2 | 1 |
| A00956 | 2 | 1 |
| A00957 | 2 | 1 |
| A00958 | 2 | 1 |
| A00959 | 2 | 1 |
| A00960 | 2 | 1 |
| A00961 | 1 | 2 |
| A00962 | 2 | 1 |
| A00963 | 2 | 1 |
| A00964 | 3 | 0 |
| A00965 | 1 | 2 |
| A00966 | 1 | 2 |
| A00967 | 1 | 2 |
| A00968 | 1 | 2 |
| A00969 | 1 | 2 |
| A00970 | 3 | 0 |
| A00971 | 3 | 0 |
| A00972 | 1 | 2 |
| A00973 | 1 | 2 |
| A00974 | 3 | 0 |
| A00975 | 3 | 0 |
| A00976 | 3 | 0 |
| A00977 | 1 | 2 |
| A00978 | 1 | 2 |
| A00979 | 3 | 0 |
| A00980 | 1 | 2 |
| A00981 | 1 | 2 |
| A00982 | 3 | 0 |
| A00983 | 2 | 1 |
| A00984 | 3 | 0 |
| A00985 | 2 | 1 |
| A00986 | 2 | 1 |
| A00987 | 2 | 1 |
| A00988 | 2 | 1 |
| A00989 | 2 | 1 |
| A00990 | 3 | 0 |
| A00991 | 3 | 0 |
| A00992 | 3 | 0 |
| A00993 | 3 | 0 |
| A00994 | 1 | 2 |
| A00995 | 1 | 2 |
| A00996 | 3 | 0 |
| A00998 | 3 | 0 |
| A00999 | 2 | 1 |
| A01000 | 3 | 0 |
| A01001 | 3 | 0 |
| A01002 | 3 | 0 |
| A01003 | 3 | 0 |
| A01004 | 3 | 0 |
| A01005 | 3 | 0 |
| A01006 | 2 | 1 |
| A01007 | 3 | 0 |
| A01008 | 3 | 0 |
| A01009 | 1 | 2 |
| A01010 | 3 | 0 |
| A01011 | 3 | 0 |
| A01012 | 3 | 0 |
| A01013 | 1 | 2 |
| A01014 | 1 | 2 |
| A01015 | 2 | 1 |
| A01016 | 2 | 1 |
| A01017 | 3 | 0 |
| A01018 | 1 | 2 |
| A01019 | 3 | 0 |
| A01020 | 3 | 0 |
| A01021 | 1 | 2 |
| A01022 | 1 | 2 |
| A01023 | 1 | 2 |
| A01024 | 1 | 2 |
| A01025 | 3 | 0 |
| A01026 | 3 | 0 |
| A01027 | 1 | 2 |
| A01028 | 3 | 0 |
| A01029 | 1 | 2 |
| A01030 | 3 | 0 |
| A01031 | 1 | 2 |
| A01032 | 1 | 2 |
| A01034 | 1 | 2 |
| A01035 | 3 | 0 |
| A01038 | 1 | 2 |
| A01039 | 1 | 2 |
| A01040 | 3 | 0 |
| A01041 | 2 | 1 |
| A01042 | 3 | 0 |
| A01043 | 1 | 2 |
| A01044 | 2 | 1 |
| A01045 | 1 | 2 |
| A01046 | 1 | 2 |
| A01047 | 1 | 2 |
| A01049 | 1 | 2 |
| A01051 | 1 | 2 |
| A01052 | 3 | 0 |
| A01053 | 2 | 1 |
| A01054 | 2 | 1 |
| A01055 | 2 | 1 |
| A01056 | 2 | 1 |
| A01057 | 1 | 2 |
| A01058 | 1 | 2 |
| A01063 | 3 | 0 |
| A01064 | 1 | 2 |
| A01065 | 2 | 1 |
| A01066 | 3 | 0 |
| A01067 | 1 | 2 |
| A01068 | 1 | 2 |
| A01069 | 1 | 2 |
| A01070 | 3 | 0 |
| A01071 | 1 | 2 |
| A01072 | 1 | 2 |
| A01073 | 3 | 0 |
| A01074 | 1 | 2 |
| A01075 | 1 | 2 |
| A01076 | 1 | 2 |
| A01077 | 3 | 0 |
| A01078 | 1 | 2 |
| A01079 | 3 | 0 |
| A01080 | 1 | 2 |
| A01081 | 2 | 1 |
| A01082 | 2 | 1 |
| A01083 | 3 | 0 |
| A01084 | 3 | 0 |
| A01085 | 1 | 2 |
| A01086 | 1 | 2 |
| A01087 | 3 | 0 |
| A01088 | 3 | 0 |
| A01089 | 1 | 2 |
| A01090 | 1 | 2 |
| A01091 | 1 | 2 |
| A01092 | 1 | 2 |
| A01093 | 1 | 2 |
| A01094 | 3 | 0 |
| A01095 | 1 | 2 |
| A01096 | 3 | 0 |
| A01097 | 3 | 0 |
| A01098 | 1 | 2 |
| A01101 | 1 | 2 |
| A01102 | 1 | 2 |
| A01104 | 1 | 2 |
| A01105 | 1 | 2 |
| A01106 | 1 | 2 |
| A01107 | 3 | 0 |
| A01108 | 2 | 1 |
| A01109 | 2 | 1 |
| A01110 | 1 | 2 |
| A01111 | 1 | 2 |
| A01113 | 1 | 2 |
| A01114 | 1 | 2 |
| A01115 | 1 | 2 |
| A01116 | 3 | 0 |
| A01117 | 1 | 2 |
| A01118 | 3 | 0 |
| A01119 | 3 | 0 |
| A01120 | 1 | 2 |
| A01121 | 2 | 1 |
| A01122 | 1 | 2 |
| A01123 | 3 | 0 |
| A01124 | 3 | 0 |
| A01125 | 3 | 0 |
| A01126 | 1 | 2 |
| A01127 | 2 | 1 |
| A01128 | 2 | 1 |
| A01129 | 3 | 0 |
| A01130 | 2 | 1 |
| A01131 | 2 | 1 |
| A01132 | 1 | 2 |
| A01133 | 1 | 2 |
| A01134 | 1 | 2 |
| A01135 | 1 | 2 |
| A01136 | 2 | 1 |
| A01137 | 1 | 2 |
| A01138 | 1 | 2 |
| A01139 | 1 | 2 |
| A01140 | 1 | 2 |
| A01141 | 1 | 2 |
| A01142 | 1 | 2 |
| A01143 | 3 | 0 |
| A01144 | 3 | 0 |
| A01145 | 1 | 2 |
| A01146 | 1 | 2 |
| A01147 | 3 | 0 |
| A01148 | 3 | 0 |
| A01149 | 3 | 0 |
| A01150 | 3 | 0 |
| A01151 | 1 | 2 |
| A01152 | 1 | 2 |
| A01153 | 3 | 0 |
| A01154 | 1 | 2 |
| A01155 | 1 | 2 |
| A01156 | 1 | 2 |
| A01157 | 1 | 2 |
| A01158 | 1 | 2 |
| A01159 | 1 | 2 |
| A01160 | 1 | 2 |
| A01161 | 1 | 2 |
| A01162 | 1 | 2 |
| A01163 | 1 | 2 |
| A01164 | 1 | 2 |
| A01165 | 3 | 0 |
| A01166 | 1 | 2 |
| A01167 | 3 | 0 |
| A01168 | 3 | 0 |
| A01169 | 1 | 2 |
| A01170 | 1 | 2 |
| A01171 | 1 | 2 |
| A01172 | 1 | 2 |
| A01174 | 3 | 0 |
| A01175 | 1 | 2 |
| A01176 | 1 | 2 |
| A01177 | 1 | 2 |
| A01178 | 1 | 2 |
| A01179 | 3 | 0 |
| A01180 | 1 | 2 |
| A01181 | 1 | 2 |
| A01182 | 1 | 2 |
| A01184 | 3 | 0 |
| A01185 | 1 | 2 |
| A01186 | 1 | 2 |
| A01187 | 3 | 0 |
| A01188 | 3 | 0 |
| A01189 | 3 | 0 |
| A01190 | 3 | 0 |
| A01191 | 3 | 0 |
| A01192 | 1 | 2 |
| A01193 | 3 | 0 |
| A01196 | 1 | 2 |
| A01197 | 1 | 2 |
| A01199 | 1 | 2 |
| A01200 | 3 | 0 |
| A01201 | 1 | 2 |
| A01202 | 1 | 2 |
| A01205 | 1 | 2 |
| A01207 | 2 | 1 |
| A01208 | 1 | 2 |
| A01209 | 1 | 2 |
| A01210 | 1 | 2 |
| A01211 | 1 | 2 |
| A01212 | 1 | 2 |
| A01214 | 3 | 0 |
| A01215 | 1 | 2 |
| A01216 | 2 | 1 |
| A01217 | 1 | 2 |
| A01218 | 2 | 1 |
| A01219 | 1 | 2 |
| A01220 | 1 | 2 |
| A01221 | 2 | 1 |
| A01222 | 1 | 2 |
| A01223 | 1 | 2 |
| A01224 | 1 | 2 |
| A01225 | 1 | 2 |
| A01226 | 1 | 2 |
| A01227 | 2 | 1 |
| A01228 | 3 | 0 |
| A01229 | 2 | 1 |
| A01230 | 1 | 2 |
| A01231 | 3 | 0 |
| A01232 | 1 | 2 |
| A01233 | 1 | 2 |
| A01234 | 1 | 2 |
| A01235 | 1 | 2 |
| A01236 | 1 | 2 |
| A01237 | 1 | 2 |
| A01238 | 3 | 0 |
| A01239 | 3 | 0 |
| A01240 | 3 | 0 |
| A01241 | 2 | 1 |
| A01242 | 1 | 2 |
| A01243 | 1 | 2 |
| A01244 | 3 | 0 |
| A01245 | 1 | 2 |
| A01246 | 1 | 2 |
| A01247 | 2 | 1 |
| A01248 | 1 | 2 |
| A01249 | 1 | 2 |
| A01250 | 1 | 2 |
| A01251 | 1 | 2 |
| A01252 | 2 | 1 |
| A01253 | 1 | 2 |
| A01254 | 2 | 1 |
| A01255 | 1 | 2 |
| A01256 | 1 | 2 |
| A01258 | 1 | 2 |
| A01259 | 2 | 1 |
| A01260 | 1 | 2 |
| A01261 | 1 | 2 |
| A01262 | 1 | 2 |
| A01263 | 1 | 2 |
| A01264 | 1 | 2 |
| A01265 | 1 | 2 |
| A01266 | 2 | 1 |
| A01267 | 1 | 2 |
| A01268 | 1 | 2 |
| A01269 | 1 | 2 |
| A01270 | 3 | 0 |
| A01271 | 1 | 2 |
| A01273 | 3 | 0 |
| A01274 | 1 | 2 |
| A01275 | 2 | 1 |
| A01276 | 2 | 1 |
| A01278 | 1 | 2 |
| A01280 | 1 | 2 |
| A01282 | 3 | 0 |
| A01283 | 3 | 0 |
| A01284 | 1 | 2 |
| A01286 | 1 | 2 |
| A01287 | 3 | 0 |
| A01288 | 1 | 2 |
| A01289 | 1 | 2 |
| A01290 | 2 | 1 |
| A01293 | 1 | 2 |
| A01294 | 1 | 2 |
| A01295 | 2 | 1 |
| A01296 | 1 | 2 |
| A01297 | 3 | 0 |
| A01298 | 1 | 2 |
| A01299 | 1 | 2 |
| A01302 | 1 | 2 |
| A01303 | 2 | 1 |
| A01304 | 3 | 0 |
| A01305 | 1 | 2 |
| A01306 | 3 | 0 |
| A01307 | 3 | 0 |
| A01308 | 2 | 1 |
| A01309 | 2 | 1 |
| A01310 | 1 | 2 |
| A01311 | 1 | 2 |
| A01312 | 2 | 1 |
| A01313 | 2 | 1 |
| A01316 | 1 | 2 |
| A01317 | 3 | 0 |
| A01318 | 1 | 2 |
| A01319 | 1 | 2 |
| A01320 | 1 | 2 |
| A01321 | 1 | 2 |
| A01322 | 1 | 2 |
| A01323 | 1 | 2 |
| A01324 | 1 | 2 |
| A01325 | 3 | 0 |
| A01326 | 3 | 0 |
| A01327 | 1 | 2 |
| A01328 | 2 | 1 |
| A01329 | 1 | 2 |
| A01330 | 1 | 2 |
| A01331 | 1 | 2 |
| A01332 | 3 | 0 |
| A01333 | 1 | 2 |
| A01334 | 1 | 2 |
| A01335 | 1 | 2 |
| A01336 | 1 | 2 |
| A01337 | 3 | 0 |
| A01340 | 3 | 0 |
| A01341 | 1 | 2 |
| A01342 | 3 | 0 |
| A01343 | 1 | 2 |
| A01344 | 3 | 0 |
| A01345 | 2 | 1 |
| A01346 | 3 | 0 |
| A01347 | 1 | 2 |
| A01348 | 1 | 2 |
| A01349 | 3 | 0 |
| A01350 | 1 | 2 |
| A01351 | 1 | 2 |
| A01352 | 1 | 2 |
| A01353 | 1 | 2 |
| A01354 | 1 | 2 |
| A01355 | 1 | 2 |
| A01356 | 3 | 0 |
| A01357 | 1 | 2 |
| A01358 | 1 | 2 |
| A01361 | 2 | 1 |
| A01362 | 1 | 2 |
| A01363 | 1 | 2 |
| A01364 | 3 | 0 |
| A01365 | 1 | 2 |
| A01366 | 1 | 2 |
| A01367 | 1 | 2 |
| A01368 | 2 | 1 |
| A01369 | 1 | 2 |
| A01373 | 1 | 2 |
| A01375 | 1 | 2 |
| A01376 | 3 | 0 |
| A01377 | 1 | 2 |
| A01378 | 3 | 0 |
| A01379 | 1 | 2 |
| A01380 | 3 | 0 |
| A01381 | 3 | 0 |
| A01382 | 1 | 2 |
| A01383 | 3 | 0 |
| A01385 | 3 | 0 |
| A01386 | 3 | 0 |
| A01387 | 2 | 1 |
| A01388 | 1 | 2 |
| A01389 | 1 | 2 |
| A01390 | 1 | 2 |
| A01392 | 3 | 0 |
| A01393 | 3 | 0 |
| A01394 | 1 | 2 |
| A01395 | 1 | 2 |
| A01396 | 3 | 0 |
| A01397 | 1 | 2 |
| A01398 | 3 | 0 |
| A01400 | 1 | 2 |
| A01401 | 1 | 2 |

A01402 1 2
A01403 1 2
A01404 1 2
A01408 1 2
A01409 3 0
A01410 2 1
A01411 1 2
A01412 1 2
A01413 1 2
A01414 3 0
A01416 3 0
A01417 1 2
A01418 1 2
A01419 1 2
A01420 3 0
A01421 1 2
A01422 1 2
A01423 1 2
A01424 1 2
A01425 3 0
A01426 1 2
A01427 3 0
A01428 1 2
A01429 3 0
A01430 1 2
A01431 3 0
A01432 1 2
A01433 3 0
A01434 1 2
A01435 1 2
A01436 1 2
A01437 3 0
A01438 1 2
A01440 1 2
A01441 2 1
A01442 2 1
A01443 3 0
A01444 1 2
A01445 1 2
A01446 1 2
A01447 1 2
A01448 1 2
A01449 1 2
A01450 1 2
A01451 1 2
A01452 1 2
A01453 2 1
A01454 2 1
A01455 1 2
A01456 1 2
A01457 1 2
A01458 1 2
A01459 1 2
A01460 1 2
A01461 2 1
A01462 3 0
A01463 1 2
A01464 1 2
A01465 1 2
A01466 2 1
A01467 3 0
A01468 3 0
A01469 1 2
A01470 3 0
A01471 3 0
A01472 1 2
A01473 2 1
A01474 2 1
A01475 1 2
A01476 3 0
A01478 2 1
A01480 1 2
A01481 3 0
A01482 1 2
A01483 1 2
A01484 3 0
A01485 2 1
A01486 3 0
A01487 1 2
A01488 2 1
A01489 1 2
A01491 1 2
A01492 2 1
A01493 2 1
A01494 2 1
A01496 1 2
A01497 3 0
A01498 1 2
A01499 1 2
A01500 1 2
A01501 1 2
A01502 1 2
A01503 1 2
A01504 3 0
A01505 3 0
A01506 1 2
A01507 3 0
A01508 2 1
A01509 2 1
A01510 3 0
A01511 2 1
A01512 3 0
A01513 3 0
A01514 3 0
A01515 2 1
A01516 1 2
A01517 3 0
A01518 3 0
A01519 3 0
A01520 3 0
A01521 3 0
A01522 1 2

A01523 1 2
A01524 1 2
A01525 3 0
A01526 2 1
A01527 2 1
A01528 3 0
A01529 3 0
A01530 1 2
A01531 1 2
A01532 1 2
A01535 3 0
A01536 1 2
A01537 1 2
A01538 1 2
A01539 1 2
A01545 1 2
A01546 1 2
A01547 1 2
A01548 3 0
A01549 1 2
A01550 2 1
A01551 3 0
A01552 1 2
A01553 1 2
A01554 2 1
A01555 1 2
A01556 1 2
A01558 3 0
A01559 1 2
A01562 1 2
A01563 1 2
A01564 1 2
A01565 1 2
A01566 1 2
A01567 1 2
A01568 3 0
A01569 1 2
A01570 3 0
A01571 1 2
A01572 1 2
A01574 1 2
A01575 3 0
A01576 3 0
A01580 1 2
A01581 3 0
A01586 3 0
A01587 1 2
A01588 1 2
A01589 1 2
A01591 1 2
A01593 1 2
A01594 3 0
A01595 1 2
A01596 3 0
A01597 1 2
A01598 3 0
A01599 3 0
A01602 1 2
A01603 1 2
A01604 3 0
A01605 1 2
A01606 1 2
A01607 1 2
A01608 1 2
A01609 1 2
A01610 3 0
A01611 3 0
A01612 3 0
A01613 3 0
A01614 1 2
A01616 1 2
A01617 1 2
A01618 1 2
A01619 1 2
A01620 3 0
A01621 1 2
A01623 1 2
A01624 1 2
A01625 1 2
A01626 1 2
A01629 1 2
A01632 3 0
A01633 1 2
A01634 1 2
A01635 3 0
A01636 1 2
A01637 1 2
A01638 1 2
A01639 3 0
A01640 3 0
A01641 1 2
A01642 1 2
A01644 1 2
A01646 2 1
A01647 3 0
A01648 1 2
A01649 1 2
A01650 1 2
A01651 1 2
A01652 1 2
A01653 3 0
A01654 1 2
A01655 1 2
A01656 1 2
A01657 1 2
A01658 1 2
A01659 1 2
A01660 2 1
A01661 3 0
A01662 3 0
A01663 3 0
A01664 1 2

A01665 1 2
A01666 1 2
A01667 1 2
A01668 1 2
A01669 1 2
A01670 1 2
A01671 1 2
A01672 2 1
A01673 1 2
A01674 1 2
A01675 1 2
A01677 1 2
A01678 1 2
A01679 2 1
A01680 2 1
A01681 1 2
A01682 1 2
A01683 2 1
A01684 1 2
A01685 1 2
A01687 2 1
A01688 1 2
A01689 1 2
A01690 1 2
A01691 1 2
A01692 1 2
A01693 1 2
A01694 1 2
A01695 1 2
A01696 1 2
A01697 2 1
A01698 2 1
A01699 1 2
A01700 2 1
A01701 1 2
A01702 1 2
A01703 1 2
A01704 1 2
A01705 1 2
A01706 1 2
A01707 1 2
A01709 1 2
A01710 1 2
A01711 1 2
A01713 1 2
A01715 1 2
A01716 1 2
A01722 1 2
A01723 1 2
A01724 1 2
A01725 1 2
A01726 1 2
A01727 1 2
A01729 1 2
A01730 1 2
A01732 1 2
A01733 1 2
A01734 1 2
A01735 1 2
A01736 1 2
A01737 2 1
A01740 1 2
A01741 1 2
A01742 1 2
A01743 1 2
A01744 1 2
A01745 1 2
A01746 1 2
A01747 1 2
A01748 1 2
A01749 1 2
A01750 1 2
A01751 1 2
A01752 1 2
A01753 1 2
A01754 1 2
A01763 1 2
A01765 1 2
A01774 2 1
A01776 1 2
A01777 1 2
A01778 1 2
A01779 1 2
A01780 1 2
A01781 1 2
A01784 1 2
A01785 1 2
A01786 1 2
A01787 1 2
A01788 1 2
A01789 1 2
A01791 1 2
A01792 1 2
A01794 1 2
A01797 1 2
A01799 3 0
A01800 1 2
A01801 1 2
A01802 1 2
A01803 1 2
A01805 1 2
A01806 3 0
A01807 1 2
A01809 1 2
A01810 1 2
A01811 1 2
A01812 1 2
A01814 1 2
A01815 1 2
A01816 1 2
A01817 1 2
A01819 3 0

A01822 1 2
A01824 1 2
A01826 1 2
A01827 2 1
A01828 1 2
A01829 1 2
A01830 1 2
A01832 1 2
A01849 1 2
A01850 1 2
A01851 1 2
A01852 1 2
A01854 3 0
A01855 3 0
A01856 1 2
A01857 1 2
A01858 3 0
A01859 3 0
A01860 3 0
A01861 2 1
A01862 1 2
A01863 1 2
A01864 3 0
A01865 3 0
A01866 1 2
A01867 1 2
A01868 3 0
A01869 3 0
A01870 1 2
A01871 1 2
A01873 1 2
A01874 3 0
A01875 3 0
A01876 2 1
A01877 3 0
A01884 1 2
A01885 1 2
A01886 1 2
A01887 2 1
A01888 3 0
A01889 1 2
A01890 1 2
A01892 1 2
A01893 3 0
A01894 3 0
A01895 3 0
A01896 1 2
A01897 1 2
A01898 1 2
A01899 1 2
A01900 1 2
A01901 3 0
A01902 3 0
A01904 3 0
A01905 3 0
A01906 1 2
A01907 1 2
A01908 1 2
A01909 1 2
A01910 2 1
A01911 1 2
A01912 1 2
A01913 1 2
A01914 1 2
A01915 1 2
A01916 2 1
A01917 1 2
A01918 3 0
A01919 3 0
A01920 3 0
A01921 1 2
A01922 3 0
A01923 1 2
A01924 3 0
A01925 3 0
A01926 2 1
A01927 1 2
A01928 1 2
A01929 3 0
A01930 3 0
A01932 2 1
A01933 3 0
A01934 3 0
A01935 3 0
A01936 3 0
A01937 3 0
A01938 3 0
A01939 3 0
A01940 3 0
A01941 3 0
A01943 1 2
A01944 3 0
A01945 1 2
A01946 3 0
A01947 3 0
A01948 2 1
A01949 3 0
A01950 3 0
A01951 3 0
A01952 3 0
A01953 3 0
A01954 3 0
A01955 3 0
A01956 3 0
A01957 1 2
A01958 3 0
A01959 1 2
A01960 3 0
A01962 1 2
A01963 1 2
A01964 1 2
A01966 3 0

A01967 1 2
A01968 3 0
A01969 1 2
A01970 1 2
A01971 1 2
A01972 1 2
A01973 1 2
A01974 1 2
A01975 3 0
A01976 3 0
A01977 2 1
A01978 3 0
A01979 3 0
A01980 3 0
A01981 3 0
A01982 1 2
A01984 2 1
A01985 3 0
A01986 3 0
A01987 1 2
A01988 1 2
A01989 1 2
A01991 2 1
A01992 3 0
A01993 2 1
A01994 3 0
A01995 3 0
A01997 3 0
A01998 2 1
A02000 1 2
A02001 3 0
A02002 2 1
A02003 3 0
A02004 2 1
A02005 2 1
A02006 3 0
A02007 2 1
A02008 3 0
A02009 2 1
A02010 3 0
A02011 3 0
A02012 3 0
A02013 2 1
A02014 3 0
A02015 3 0
A02016 3 0
A02017 3 0
A02018 3 0
A02019 2 1
A02020 2 1
A02021 1 2
A02022 3 0
A02023 2 1
A02024 3 0
A02025 2 1
A02026 2 1
A02027 2 1
A02028 3 0
A02029 2 1
A02030 1 2
A02031 1 2
A02032 3 0
A02034 3 0
A02035 3 0
A02036 3 0
A02037 2 1
A02038 3 0
A02039 2 1
A02040 3 0
A02041 2 1
A02042 1 2
A02043 3 0
A02044 2 1
A02045 3 0
A02046 3 0
A02047 1 2
A02048 3 0
A02049 3 0
A02050 3 0
A02051 1 2
A02052 3 0
A02053 3 0
A02054 3 0
A02055 1 2
A02056 3 0
A02057 3 0
A02058 2 1
A02059 3 0
A02060 3 0
A02061 3 0
A02062 1 2
A02063 3 0
A02064 3 0
A02065 3 0
A02066 3 0
A02067 1 2
A02068 3 0
A02069 1 2
A02070 3 0
A02071 2 1
A02072 1 2
A02073 3 0
A02074 1 2
A02075 1 2
A02076 3 0
A02077 3 0
A02078 1 2
A02079 3 0
A02080 3 0
A02081 1 2
A02082 3 0
A02083 1 2

A02084 1 2
A02085 1 2
A02086 1 2
A02088 1 2
A02090 1 2
A02091 1 2
A02092 1 2
A02093 1 2
A02094 2 1
A02095 1 2
A02096 3 0
A02097 3 0
A02098 3 0
A02099 3 0
A02100 2 1
A02101 3 0
A02102 1 2
A02103 3 0
A02104 3 0
A02105 1 2
A02107 1 2
A02108 1 2
A02109 2 1
A02110 3 0
A02111 2 1
A02112 1 2
A02113 3 0
A02114 2 1
A02115 2 1
A02116 1 2
A02117 2 1
A02118 3 0
A02119 3 0
A02120 3 0
A02121 3 0
A02122 3 0
A02123 3 0
A02124 1 2
A02125 2 1
A02126 2 1
A02127 1 2
A02128 2 1
A02129 3 0
A02130 3 0
A02131 3 0
A02132 1 2
A02133 1 2
A02134 1 2
A02135 2 1
A02136 2 1
A02137 1 2
A02139 2 1
A02142 1 2
A02143 2 1
A02144 2 1
A02145 3 0
A02146 1 2
A02147 2 1
A02148 2 1
A02149 3 0
A02150 2 1
A02151 1 2
A02152 2 1
A02153 2 1
A02154 2 1
A02162 3 0
A02163 3 0
A02164 3 0
A02165 3 0
A02166 3 0
A02167 3 0
A02168 3 0
A02169 1 2
A02170 2 1
A02171 1 2
A02172 1 2
A02173 2 1
A02174 3 0
A02175 1 2
A02176 1 2
A02177 1 2
A02178 3 0
A02179 1 2
A02180 2 1
A02181 2 1
A02182 1 2
A02183 2 1
A02184 3 0
A02185 3 0
A02186 3 0
A02187 3 0
A02191 3 0
A02192 3 0
A02193 2 1
A02196 3 0
A02197 2 1
A02198 1 2
A02199 1 2
A02202 3 0
A02203 1 2
A02204 3 0
A02205 1 2
A02206 1 2
A02207 2 1
A02208 1 2
A02209 3 0
A02210 2 1
A02211 3 0
A02212 3 0
A02213 3 0
A02214 2 1
A02216 2 1

A02217 1 2
A02218 1 2
A02219 3 0
A02221 3 0
A02222 1 2
A02223 1 2
A02224 2 1
A02225 1 2
A02226 2 1
A02227 2 1
A02228 1 2
A02229 1 2
A02230 1 2
A02231 2 1
A02232 2 1
A02233 1 2
A02235 1 2
A02236 2 1
A02237 2 1
A02238 1 2
A02239 2 1
A02240 1 2
A02242 2 1
A02243 2 1
A02244 1 2
A02245 1 2
A02246 2 1
A02247 2 1
A02248 1 2
A02249 2 1
A02250 1 2
A02251 1 2
A02252 1 2
A02253 1 2
A02254 1 2
A02255 1 2
A02256 1 2
A02257 1 2
A02258 1 2
A02259 1 2
A02260 1 2
A02261 2 1
A02262 2 1
A02263 1 2
A02264 1 2
A02265 2 1
A02266 1 2
A02267 1 2
A02268 1 2
A02269 3 0
A02270 3 0
A02271 3 0
A02273 1 2
A02274 1 2
A02275 3 0
A02276 1 2
A02277 1 2
A02278 2 1
A02279 1 2
A02280 2 1
A02281 1 2
A02282 2 1
A02283 1 2
A02284 1 2
A02285 2 1
A02286 2 1
A02287 1 2
A02290 2 1
A02334 2 1
A02335 2 1
A02336 3 0
A02337 3 0
A02338 2 1
A02339 1 2
A02341 3 0
A02343 1 2
A02344 2 1
A02345 3 0
A02346 1 2
A02347 3 0
A02348 2 1
A02349 1 2
A02350 1 2
A02351 2 1
A02352 2 1
A02353 3 0
A02354 2 1
A02355 2 1
A02356 2 1
A02359 1 2
A02360 1 2
A02361 2 1
A02362 1 2
A02363 2 1
A02364 1 2
A02365 1 2
A02367 1 2
A02368 2 1
A02369 3 0
A02370 1 2
A02371 3 0
A02372 3 0
A02373 3 0
A02375 3 0
A02376 1 2
A02377 2 1
A02378 2 1
A02379 2 1
A02381 1 2
A02383 2 1
A02384 3 0
A02385 3 0

A02387 1 2
A02388 1 2
A02389 3 0
A02391 3 0
A02392 3 0
A02393 1 2
A02394 2 1
A02395 1 2
A02396 2 1
A02397 1 2
A02398 3 0
A02399 1 2
A02400 1 2
A02401 3 0
A02402 1 2
A02403 1 2
A02404 3 0
A02405 1 2
A02406 1 2
A02408 1 2
A02409 1 2
A02411 1 2
A02412 3 0
A02413 2 1
A02414 3 0
A02416 2 1
A02417 2 1
A02418 2 1
A02420 3 0
A02421 1 2
A02422 3 0
A02423 2 1
A02424 1 2
A02425 1 2
A02426 1 2
A02427 1 2
A02428 3 0
A02429 1 2
A02430 1 2
A02431 1 2
A02432 1 2
A02433 1 2
A02434 1 2
A02435 1 2
A02436 1 2
A02437 1 2
A02438 3 0
A02439 1 2
A02440 1 2
A02441 1 2
A02442 1 2
A02443 2 1
A02444 1 2
A02445 1 2
A02446 3 0
A02447 1 2
A02448 1 2
A02449 1 2
A02450 3 0
A02451 1 2
A02452 1 2
A02453 2 1
A02454 1 2
A02455 1 2
A02456 1 2
A02457 1 2
A02458 2 1
A02459 2 1
A02460 1 2
A02461 1 2
A02462 1 2
A02463 2 1
A02464 2 1
A02465 3 0
A02466 1 2
A02467 3 0
A02468 1 2
A02469 1 2
A02470 2 1
A02471 1 2
A02472 1 2
A02473 2 1
A02474 3 0
A02475 3 0
A02476 3 0
A02477 3 0
A02478 1 2
A02483 1 2
A02484 1 2
A02485 1 2
A02486 2 1
A02487 2 1
A02488 1 2
A02489 1 2
A02490 2 1
A02491 1 2
A02492 1 2
A02493 1 2
A02494 1 2
A02495 2 1
A02496 1 2
A02497 1 2
A02498 1 2
A02499 2 1
A02501 1 2
A02502 1 2
A02504 1 2
A02505 1 2
A02506 3 0
A02507 1 2
A02509 1 2
A02510 2 1

A02511 2 1
A02512 3 0
A02513 1 2
A02514 1 2
A02515 1 2
A02516 3 0
A02517 1 2
A02518 1 2
A02519 3 0
A02520 2 1
A02522 1 2
A02523 1 2
A02524 2 1
A02525 2 1
A02526 1 2
A02527 1 2
A02528 1 2
A02529 1 2
A02530 3 0
A02531 3 0
A02532 1 2
A02533 1 2
A02534 1 2
A02535 1 2
A02536 1 2
A02538 3 0
A02539 3 0
A02540 1 2
A02541 3 0
A02542 1 2
A02543 1 2
A02544 1 2
A02545 3 0
A02546 3 0
A02548 3 0
A02549 1 2
A02550 1 2
A02551 3 0
A02552 3 0
A02553 3 0
A02554 3 0
A02555 1 2
A02556 1 2
A02557 2 1
A02558 3 0
A02559 1 2
A02560 1 2
A02562 1 2
A02563 1 2
A02564 1 2
A02565 1 2
A02566 1 2
A02567 1 2
A02568 3 0
A02569 3 0
A02570 1 2
A02571 1 2
A02572 3 0
A02573 1 2
A02574 1 2
A02575 2 1
A02576 1 2
A02577 3 0
A02578 2 1
A02579 1 2
A02580 1 2
A02581 3 0
A02583 1 2
A02584 3 0
A02585 1 2
A02586 3 0
A02587 1 2
A02588 1 2
A02589 3 0
A02591 1 2
A02592 3 0
A02594 3 0
A02595 2 1
A02596 1 2
A02597 3 0
A02598 3 0
A02599 2 1
A02600 3 0
A02601 2 1
A02602 3 0
A02603 3 0
A02604 3 0
A02605 3 0
A02606 3 0
A02607 3 0
A02608 3 0
A02609 3 0
A02610 3 0
A02611 1 2
A02612 1 2
A02613 3 0
A02614 3 0
A02615 3 0
A02616 3 0
A02617 1 2
A02618 2 1
A02619 2 1
A02620 1 2
A02621 1 2
A02622 2 1
A02623 2 1
A02624 2 1
A02625 2 1
A02626 2 1
A02628 2 1
A02629 1 2
A02630 1 2

A02634 2 1
A02635 2 1
A02637 2 1
A02638 3 0
A02639 3 0
A02640 1 2
A02641 2 1
A02642 2 1
A02643 2 1
A02644 2 1
A02645 2 1
A02646 2 1
A02647 1 2
A02648 2 1
A02649 2 1
A02650 2 1
A02651 2 1
A02654 1 2
A02655 1 2
A02656 3 0
A02657 3 0
A02658 3 0
A02659 3 0
A02660 3 0
A02661 2 1
A02662 1 2
A02663 1 2
A02664 3 0
A02665 2 1
A02666 2 1
A02667 3 0
A02668 2 1
A02669 3 0
A02670 2 1
A02671 2 1
A02672 1 2
A02674 2 1
A02675 2 1
A02676 1 2
A02678 1 2
A02679 3 0
A02680 3 0
A02681 3 0
A02682 2 1
A02683 3 0
A02684 3 0
A02685 1 2
A02686 2 1
A02687 1 2
A02688 1 2
A02689 1 2
A02690 3 0
A02691 3 0
A02692 1 2
A02693 1 2
A02694 2 1
A02695 1 2
A02696 1 2
A02697 2 1
A02698 2 1
A02701 2 1
A02702 3 0
A02704 1 2
A02705 2 1
A02706 1 2
A02707 1 2
A02708 1 2
A02709 1 2
A02710 2 1
A02712 1 2
A02713 2 1
A02714 1 2
A02718 1 2
A02719 1 2
A02720 2 1
A02721 3 0
A02722 1 2
A02723 1 2
A02724 1 2
A02725 1 2
A02726 3 0
A02727 2 1
A02729 3 0
A02730 2 1
A02731 3 0
A02732 3 0
A02733 3 0
A02734 3 0
A02735 3 0
A02736 1 2
A02737 1 2
A02738 1 2
A02739 3 0
A02740 3 0
A02741 3 0
A02742 3 0
A02743 1 2
A02744 2 1
A02745 3 0
A02746 3 0
A02747 1 2
A02748 3 0
A02749 2 1
A02750 3 0
A02751 2 1
A02752 3 0
A02753 2 1
A02754 1 2
A02755 3 0
A02756 3 0
A02757 3 0
A02758 1 2

A02759 2 1
A02760 2 1
A02761 2 1
A02762 2 1
A02763 1 2
A02764 1 2
A02765 1 2
A02766 3 0
A02767 3 0
A02768 1 2
A02769 3 0
A02770 3 0
A02771 3 0
A02772 1 2
A02773 1 2
A02774 1 2
A02775 1 2
A02778 1 2
A02780 1 2
A02781 1 2
A02782 1 2
A02783 1 2
A02784 1 2
A02786 1 2
A02787 3 0
A02788 1 2
A02789 1 2
A02791 1 2
A02792 3 0
A02793 1 2
A02794 2 1
A02795 3 0
A02796 3 0
A02797 1 2
A02798 1 2
A02799 1 2
A02801 1 2
A02802 2 1
A02803 3 0
A02804 2 1
A02805 1 2
A02806 2 1
A02807 3 0
A02808 3 0
A02809 3 0
A02810 1 2
A02811 1 2
A02812 1 2
A02813 3 0
A02814 3 0
A02815 2 1
A02816 2 1
A02817 1 2
A02818 3 0
A02819 3 0
A02820 3 0
A02821 3 0
A02822 2 1
A02823 2 1
A02824 3 0
A02825 1 2
A02826 3 0
A02827 1 2
A02828 1 2
A02829 1 2
A02830 1 2
A02832 3 0
A02833 1 2
A02834 1 2
A02836 3 0
A02837 1 2
A02838 1 2
A02839 1 2
A02840 1 2
A02841 2 1
A02842 1 2
A02843 1 2
A02844 1 2
A02845 1 2
A02846 2 1
A02847 1 2
A02848 1 2
A02851 1 2
A02853 3 0
A02855 1 2
A02856 1 2
A02857 1 2
A02859 3 0
A02860 2 1
A02861 2 1
A02862 3 0
A02864 1 2
A02866 3 0
A02867 1 2
A02868 1 2
A02869 1 2
A02870 3 0
A02871 3 0
A02872 3 0
A02873 1 2
A02874 3 0
A02875 1 2
A02876 1 2
A02877 3 0
A02878 1 2
A02879 1 2
A02880 1 2
A02881 1 2
A02882 3 0
A02883 3 0
A02884 3 0
A02885 1 2
A02886 3 0
A02887 3 0
A02888 1 2
A02889 3 0
A02890 1 2
A02891 1 2
A02892 3 0
A02893 1 2
A02894 1 2
A02895 3 0
A02896 1 2
A02897 1 2
A02898 3 0
A02899 3 0
A02900 2 1
A02901 3 0
A02902 1 2
A02903 1 2
A02904 1 2
A02905 1 2
A02906 1 2
A02907 1 2
A02909 1 2
A02910 2 1
A02911 2 1
A02913 2 1
A02921 2 1
A02922 2 1
A02923 2 1
A02924 2 1
A02925 2 1
A02926 2 1
A02927 2 1
A02928 2 1
A02930 1 2
A02931 2 1
A02933 1 2
A02935 2 1
A02938 2 1
A02939 2 1
A02940 2 1
A02941 2 1
A02944 2 1
A02945 2 1
A02948 1 2
A02950 1 2
A02951 2 1
A02957 2 1
A02958 1 2
A02962 2 1
A02964 1 2
A02966 2 1
A02967 2 1
A02969 2 1
A02972 1 2
A02973 2 1
A02975 2 1
A02976 2 1
A02982 2 1
A02983 2 1
A02986 2 1
A02987 1 2
A02988 2 1
A02990 2 1
A02991 2 1
A02994 2 1
A02995 2 1
A02998 2 1
A02999 2 1
A03000 1 2
A03001 3 0
A03002 3 0
A03003 1 2
A03004 3 0
A03005 3 0
A03006 1 2
A03007 1 2
A03008 2 1
A03009 1 2
A03010 1 2
A03011 1 2
A03012 1 2
A03013 1 2
A03014 3 0
A03015 3 0
A03016 3 0
A03017 1 2
A03018 1 2
A03019 1 2
A03020 1 2
A03021 1 2
A03022 1 2
A03023 2 1
A03024 1 2
A03025 1 2
A03026 2 1
A03027 3 0
A03028 2 1
A03029 3 0
A03030 3 0
A03031 3 0
A03032 3 0
A03033 2 1
A03034 1 2
A03035 1 2
A03036 3 0
A03037 1 2
A03038 3 0
A03039 3 0
A03041 1 2
A03042 1 2
A03043 1 2
A03044 1 2
A03045 3 0
A03046 3 0
A03047 3 0
A03048 2 1
A03049 2 1
A03050 3 0
A03051 2 1
A03052 1 2
A03053 1 2
A03054 2 1
A03055 2 1
A03056 2 1
A03057 2 1
A03058 1 2
A03059 1 2
A03060 1 2
A03061 1 2
A03062 2 1
A03063 2 1
A03064 3 0
A03065 3 0
A03066 3 0
A03067 3 0
A03068 3 0
A03069 1 2
A03070 2 1
A03071 3 0
A03072 3 0
A03073 3 0
A03074 1 2
A03075 1 2
A03076 1 2
A03077 3 0
A03078 2 1
A03079 1 2
A03080 1 2
A03081 1 2
A03082 1 2
A03083 2 1
A03084 3 0
A03085 1 2
A03086 2 1
A03087 1 2
A03088 1 2
A03089 1 2
A03090 2 1
A03091 1 2
A03092 1 2
A03093 1 2
A03094 1 2
A03095 1 2
A03096 1 2
A03097 2 1
A03098 1 2
A03099 1 2
A03100 1 2
A03101 1 2
A03102 1 2
A03103 1 2
A03104 1 2
A03105 2 1
A03106 1 2
A03107 1 2
A03108 1 2
A03109 1 2
A03110 1 2
A03111 1 2
A03112 1 2
A03114 1 2
A03115 1 2
A03116 3 0
A03117 3 0
A03118 2 1
A03119 2 1
A03120 2 1
A03121 1 2
A03122 1 2
A03123 1 2
A03124 1 2
A03126 1 2
A03127 1 2
A03128 2 1
A03129 2 1
A03130 2 1
A03131 1 2
A03132 3 0
A03133 3 0
A03134 2 1
A03135 2 1
A03136 1 2
A03137 1 2
A03138 3 0
A03139 1 2
A03141 1 2
A03142 3 0
A03144 1 2
A03145 1 2
A03146 3 0
A03147 1 2
A03148 1 2
A03149 3 0
A03150 1 2
A03151 1 2
A03152 1 2
A03153 1 2
A03154 1 2
A03155 1 2
A03156 1 2
A03157 1 2
A03158 1 2
A03161 1 2
A03162 1 2
A03163 1 2
A03164 2 1
A03165 3 0
A03166 1 2
A03167 1 2
A03169 3 0
A03170 2 1
A03171 2 1
A03172 1 2
A03173 1 2
A03174 1 2
A03175 1 2
A03176 1 2
A03177 1 2
A03178 2 1
A03179 1 2
A03180 2 1
A03181 3 0
A03182 1 2
A03183 3 0
A03185 1 2
A03186 2 1
A03187 3 0
A03189 1 2
A03190 3 0
A03191 3 0
A03192 3 0
A03193 1 2
A03194 2 1
A03195 1 2
A03196 1 2
A03197 1 2
A03198 1 2
A03199 1 2
A03200 1 2
A03201 1 2
A03202 3 0
A03203 3 0
A03204 3 0
A03205 3 0
A03206 3 0
A03207 3 0
A03208 3 0
A03209 2 1
A03210 2 1
A03211 2 1
A03213 3 0
A03214 3 0
A03215 2 1
A03216 3 0
A03218 1 2
A03219 1 2
A03220 1 2
A03221 1 2
A03222 1 2
A03223 1 2
A03224 2 1
A03225 3 0
A03228 1 2
A03229 3 0
A03230 3 0
A03231 3 0
A03232 1 2
A03233 1 2
A03234 3 0
A03235 1 2
A03236 1 2
A03237 2 1
A03238 2 1
A03239 3 0
A03240 3 0
A03241 1 2
A03242 3 0
A03243 3 0
A03245 2 1
A03248 1 2
A03249 1 2
A03251 1 2
A03252 1 2
A03253 1 2
A03254 2 1
A03255 1 2
A03256 1 2
A03257 2 1
A03258 3 0
A03259 2 1
A03260 2 1
A03261 2 1
A03262 3 0
A03263 3 0
A03264 3 0
A03267 1 2
A03268 2 1
A03269 1 2
A03270 3 0
A03271 1 2
A03272 3 0
A03273 1 2
A03274 1 2
A03275 3 0
A03276 1 2
A03277 1 2
A03278 3 0
A03279 3 0
A03280 3 0
A03281 3 0
A03282 1 2
A03283 1 2
A03284 1 2
A03285 1 2
A03286 1 2
A03287 3 0
A03288 2 1
A03289 1 2
A03290 1 2
A03291 3 0
A03292 3 0
A03294 1 2
A03295 1 2
A03296 3 0
A03297 3 0
A03298 3 0
A03299 3 0
A03300 3 0
A03301 3 0
A03302 2 1
A03303 3 0
A03304 3 0
A03305 3 0
A03306 3 0
A03307 3 0
A03308 3 0
A03309 3 0
A03310 1 2
A03311 1 2
A03312 1 2
A03313 1 2
A03314 1 2
A03315 3 0
A03316 3 0
A03317 3 0
A03318 2 1
A03319 1 2
A03320 1 2
A03321 2 1
A03323 1 2
A03324 1 2
A03325 3 0
A03326 1 2
A03327 3 0
A03328 3 0
A03329 1 2
A03330 3 0
A03331 1 2
A03332 3 0
A03333 1 2
A03334 3 0
A03335 1 2
A03336 3 0
A03337 3 0
A03338 3 0
A03339 1 2
A03341 1 2
A03342 3 0
A03343 3 0
A03345 1 2
A03346 3 0
A03347 3 0
A03348 3 0
A03349 3 0
A03350 3 0
A03351 3 0
A03352 3 0
A03353 2 1
A03354 1 2
A03355 1 2
A03356 1 2
A03357 1 2
A03358 1 2
A03359 1 2
A03360 3 0
A03361 3 0
A03362 3 0
A03363 3 0
A03364 1 2
A03365 1 2
A03366 1 2
A03367 1 2
A03368 1 2
A03369 1 2
A03370 1 2
A03373 1 2
A03375 2 1
A03376 3 0
A03377 1 2
A03378 1 2
A03379 3 0
A03380 3 0
A03381 3 0
A03382 3 0
A03383 1 2
A03384 3 0
A03385 3 0
A03386 3 0
A03387 1 2
A03388 3 0
A03389 1 2
A03390 3 0
A03391 3 0
A03392 1 2
A03394 1 2
A03395 1 2
A03396 3 0
A03397 1 2
A03398 3 0
A03399 3 0
A03400 3 0
A03401 3 0
A03403 1 2
A03404 1 2
A03405 1 2
A03406 3 0
A03407 3 0
A03409 3 0
A03410 1 2
A03411 3 0
A03412 1 2
A03413 1 2
A03416 1 2
A03417 3 0
A03418 3 0
A03419 3 0
A03420 3 0
A03421 1 2
A03422 1 2
A03423 3 0
A03424 3 0
A03425 1 2
A03426 1 2
A03427 3 0
A03428 3 0
A03429 3 0
A03430 1 2
A03431 3 0
A03432 1 2
A03433 1 2
A03434 1 2
A03435 1 2
A03436 1 2
A03437 1 2
A03438 1 2
A03439 1 2
A03440 1 2
A03441 3 0
A03442 3 0
A03443 1 2
A03444 1 2
A03445 1 2
A03446 1 2
A03447 1 2
A03448 3 0
A03449 3 0
A03450 1 2
A03451 3 0
A03452 1 2
A03453 1 2
A03454 1 2
A03455 1 2
A03456 1 2
A03457 1 2
A03458 2 1
A03459 1 2
A03460 1 2
A03461 1 2
A03462 1 2
A03463 3 0
A03464 3 0
A03465 1 2
A03466 3 0
A03467 1 2
A03468 1 2
A03469 1 2
A03470 1 2
A03471 1 2
A03473 3 0
A03474 3 0
A03475 3 0
A03476 2 1
A03477 1 2
A03481 1 2
A03482 3 0
A03483 1 2
A03484 1 2
A03485 3 0
A03486 1 2
A03487 3 0
A03488 3 0
A03489 3 0
A03490 3 0
A03491 1 2
A03492 3 0
A03493 3 0
A03494 3 0
A03495 1 2
A03496 1 2
A03497 1 2
A03500 3 0
A03502 3 0
A03503 3 0
A03504 3 0
A03505 2 1
A03506 2 1
A03507 2 1
A03508 3 0
A03509 1 2
A03510 3 0
A03511 2 1
A03512 1 2
A03513 1 2
A03514 3 0
A03515 1 2
A03516 1 2
A03517 3 0
A03518 1 2
A03519 3 0
A03520 3 0
A03521 1 2
A03522 3 0
A03523 3 0
A03524 1 2
A03525 1 2
A03526 3 0
A03527 1 2
A03528 1 2
A03529 1 2
A03530 1 2
A03531 1 2
A03533 1 2
A03534 3 0
A03535 1 2
A03536 3 0
A03537 3 0
A03538 1 2
A03539 1 2
A03540 3 0
A03541 3 0
A03542 3 0
A03543 3 0
A03544 3 0
A03545 3 0
A03546 2 1
A03547 1 2
A03548 3 0
A03549 1 2
A03550 1 2
A03551 2 1
A03552 1 2
A03553 1 2
A03554 1 2
A03555 1 2
A03556 1 2
A03557 1 2
A03558 3 0
A03559 3 0
A03560 1 2
A03561 1 2
A03563 1 2
A03564 3 0
A03565 1 2
A03566 1 2
A03567 1 2
A03568 1 2
A03569 1 2
A03570 1 2
A03571 1 2
A03572 1 2
A03573 1 2
A03574 3 0
A03575 1 2
A03576 1 2
A03577 3 0
A03578 3 0
A03579 1 2
A03580 1 2
A03581 1 2
A03582 1 2
A03583 1 2
A03584 1 2
A03585 3 0
A03586 3 0
A03587 1 2
A03588 1 2
A03589 1 2
A03590 1 2
A03591 3 0
A03592 3 0
A03593 3 0
A03594 1 2
A03595 1 2
A03596 1 2
A03597 1 2
A03598 3 0
A03599 1 2
A03600 1 2
A03601 1 2
A03602 3 0
A03603 1 2
A03604 1 2
A03605 1 2
A03606 1 2
A03607 1 2
A03608 1 2
A03609 3 0
A03610 3 0
A03611 2 1
A03612 2 1
A03613 1 2
A03614 1 2
A03615 3 0
A03616 1 2
A03618 3 0
A03619 3 0
A03620 3 0
A03621 1 2
A03623 3 0
A03624 3 0
A03625 1 2
A03626 2 1
A03627 3 0
A03628 1 2
A03629 1 2
A03630 1 2
A03631 3 0
A03632 3 0
A03633 1 2
A03634 3 0
A03635 3 0
A03636 1 2
A03637 1 2
A03638 1 2
A03639 1 2
A03640 3 0
A03641 2 1
A03642 3 0
A03643 3 0
A03644 3 0
A03645 1 2
A03646 3 0
A03647 3 0
A03648 3 0
A03649 1 2
A03650 1 2
A03651 3 0
A03652 3 0
A03656 3 0
A03657 3 0
A03660 3 0
A03661 3 0
A03662 3 0
A03674 1 2
A03675 1 2
A03676 3 0
A03679 1 2
A03680 1 2
A03681 3 0
A03684 1 2
A03685 2 1
A03686 1 2
A03687 1 2
A03690 3 0
A03691 3 0
A03694 3 0
A03698 1 2
A03700 3 0
A03701 3 0
A03704 3 0
A03705 3 0
A03706 3 0
A03707 3 0
A03709 2 1
A03711 2 1
A03712 1 2
A03713 1 2
A03714 1 2
A03715 1 2
A03716 3 0
A03717 1 2
A03718 1 2
A03719 1 2
A03720 1 2
A03721 1 2
A03722 1 2
A03723 1 2
A03724 1 2
A03725 1 2
A03726 1 2
A03727 1 2
A03728 1 2
A03729 3 0
A03730 1 2
A03731 3 0
A03732 2 1
A03733 1 2
A03734 3 0
A03735 3 0
A03736 3 0
A03737 1 2
A03738 1 2
A03739 1 2
A03740 1 2
A03741 1 2
A03742 1 2
A03744 1 2
A03745 1 2
A03746 1 2
A03747 1 2
A03748 1 2
A03749 1 2
A03750 1 2
A03751 1 2
A03753 1 2
A03778 1 2
A03779 1 2
A03780 1 2
A03781 1 2
A03782 1 2
A03784 1 2
A03785 1 2
A03786 1 2
A03787 1 2
A03788 1 2
A03790 1 2
A03792 1 2
A03793 1 2
A03794 1 2
A03795 1 2
A03796 1 2
A03797 1 2
A03798 1 2
A03799 1 2
A03800 1 2
A03801 1 2
A03803 1 2
A03805 1 2
A03806 1 2
A03807 1 2
A03808 1 2
A03809 1 2
A03810 1 2
A03811 1 2
A03813 1 2
A03814 1 2
A03815 1 2
A03816 1 2
A03817 1 2
A03818 1 2
A03819 1 2
A03820 1 2
A03821 1 2
A03823 1 2
A03825 1 2
A03826 1 2
A03827 1 2
A03828 1 2
A03829 1 2
A03830 1 2
A03831 1 2
A03833 1 2
A03834 1 2
A03835 1 2
A03836 1 2
A03837 1 2
A03838 1 2
A03839 1 2
A03840 1 2
A03841 1 2
A03842 1 2
A03843 1 2
A03845 1 2
A03847 1 2
A03848 1 2
A03849 1 2
A03850 1 2
A03852 1 2
A03853 1 2
A03854 1 2
A03855 1 2
A03856 1 2
A03857 1 2
A03858 1 2
A03859 1 2
A03860 1 2
A03863 1 2
A03865 1 2
A03867 1 2
A03868 1 2
A03869 1 2
A03870 1 2
A03872 1 2
A03873 1 2
A03876 1 2
A03877 1 2
A03878 1 2
A03879 1 2
A03881 1 2
A03882 1 2
A03883 1 2
A03884 1 2
A03885 1 2
A03886 1 2
A03888 1 2
A03889 1 2
A03890 1 2
A03891 1 2
A03892 1 2
A03895 1 2
A03896 1 2
A03897 1 2
A03898 1 2
A03899 1 2
A03900 1 2
A03901 1 2
A03902 1 2
A03904 1 2
A03905 1 2
A03906 1 2
A03907 1 2
A03908 1 2
A03909 1 2
A03910 1 2
A03911 1 2
A03915 1 2
A03916 1 2
A03918 1 2
A03919 1 2
A03920 1 2
A03922 1 2
A03923 1 2
A03924 1 2
A03925 1 2
A03926 1 2
A03927 1 2
A03928 1 2
A03929 1 2
A03931 1 2
A03932 1 2
A03933 1 2
A03934 1 2
A03935 1 2
A03936 1 2
A03937 1 2
A03938 1 2
A03939 1 2
A03940 1 2
A03941 1 2
A03942 1 2
A03943 1 2
A03944 1 2
A03945 1 2
A03946 1 2
A03948 1 2
A03949 1 2
A03950 1 2
A03951 1 2
A03952 1 2
A03953 1 2
A03954 1 2
A03955 1 2
A03956 1 2
A03957 1 2
A03958 1 2
A03959 1 2
A03960 1 2
A03961 1 2
A03962 1 2
A03963 1 2
A03965 1 2
A03967 1 2
A03968 1 2
A03969 1 2
A03971 1 2
A03972 1 2
A03973 1 2
A03975 1 2
A03976 1 2
A03977 1 2
A03978 1 2
A03979 1 2
A03980 1 2
A03981 1 2
A03983 1 2
A03984 1 2
A03985 1 2
A03986 1 2
A03987 1 2
A03988 1 2
A03989 1 2
A03991 1 2
A03992 1 2
A03993 1 2
A03994 1 2
A03995 1 2
A03996 1 2
A03997 1 2
A03998 1 2
A03999 1 2
A04000 1 2
A04002 1 2
A04004 1 2
A04005 1 2
A04006 1 2
A04007 1 2
A04008 1 2
A04009 1 2
A04010 1 2
A04011 1 2
A04012 1 2
A04013 1 2
A04014 1 2
A04015 1 2
A04016 1 2
A04017 1 2
A04018 1 2
A04019 1 2
A04020 1 2
A04021 1 2
A04022 1 2
A04023 1 2
A04024 1 2
A04025 1 2
A04026 1 2
A04027 1 2
A04028 1 2
A04029 1 2
A04030 1 2
A04031 1 2
A04032 1 2
A04033 1 2
A04034 1 2
A04035 1 2
A04037 1 2
A04039 1 2
A04040 1 2
A04041 1 2
A04042 1 2
A04043 1 2
A04044 1 2
A04045 1 2
A04046 1 2
A04047 1 2
A04048 1 2
A04049 1 2
A04050 1 2
A04051 1 2
A04052 1 2
A04053 1 2
A04054 1 2
A04055 1 2
A04056 1 2
A04057 1 2
A04058 1 2
A04059 1 2
A04060 1 2
A04062 1 2
A04063 1 2
A04064 1 2
A04065 1 2
A04066 1 2
A04067 1 2
A04068 1 2
A04069 1 2
A04070 1 2
A04071 1 2
A04072 1 2
A04073 1 2
A04074 1 2
A04075 1 2
A04076 1 2
A04077 1 2
A04078 1 2
A04079 1 2
A04080 1 2
A04081 1 2

| | | | | | | | | | |
|---|---|---|---|---|---|---|---|---|---|
| A04082 1 2 | AJ0061 3 0 | BM0189 3 0 | DK0131 1 2 | JB0064 1 2 | JB0206 1 2 | JB0336 1 2 | JB0472 1 2 | JB0598 1 2 | JB0722 1 2 |
| A04083 1 2 | AJ0062 3 0 | BM0190 3 0 | DK0132 1 2 | JB0068 1 2 | JB0207 1 2 | JB0337 1 2 | JB0473 1 2 | JB0599 1 2 | JB0727 1 2 |
| A04084 1 2 | AJ0063 3 0 | BM0191 3 0 | DK0133 3 0 | JB0069 1 2 | JB0208 1 2 | JB0338 1 2 | JB0474 1 2 | JB0600 1 2 | JB0728 1 2 |
| A04085 1 2 | AJ0064 3 0 | BM0192 3 0 | DK0134 1 2 | JB0070 1 2 | JB0209 1 2 | JB0339 1 2 | JB0475 1 2 | JB0602 1 2 | JB0733 1 2 |
| A04086 1 2 | AJ0065 3 0 | DK0002 3 0 | DK0135 1 2 | JB0071 1 2 | JB0211 1 2 | JB0340 1 2 | JB0476 1 2 | JB0604 1 2 | JB0734 1 2 |
| A04087 1 2 | AJ0066 2 1 | DK0004 1 2 | DK0136 1 2 | JB0072 1 2 | JB0212 1 2 | JB0341 1 2 | JB0478 1 2 | JB0606 1 2 | JB0735 1 2 |
| A04089 1 2 | AJ0067 2 1 | DK0005 1 2 | DK0137 1 2 | JB0073 1 2 | JB0213 1 2 | JB0342 1 2 | JB0479 1 2 | JB0607 1 2 | JB0736 1 2 |
| A04090 1 2 | AJ0068 2 1 | DK0006 3 0 | DK0138 1 2 | JB0075 1 2 | JB0214 1 2 | JB0343 1 2 | JB0481 1 2 | JB0608 1 2 | JB0737 1 2 |
| A04091 1 2 | AJ0069 3 0 | DK0007 3 0 | DK0139 3 0 | JB0077 1 2 | JB0215 1 2 | JB0345 1 2 | JB0482 1 2 | JB0609 1 2 | JB0738 1 2 |
| A04092 1 2 | AJ0070 3 0 | DK0008 1 2 | DK0140 1 2 | JB0078 1 2 | JB0216 1 2 | JB0346 1 2 | JB0483 1 2 | JB0610 1 2 | JB0739 1 2 |
| A04093 1 2 | AJ0071 3 0 | DK0009 1 2 | DK0141 1 2 | JB0079 1 2 | JB0220 1 2 | JB0348 1 2 | JB0484 1 2 | JB0611 1 2 | JB0740 1 2 |
| A04094 1 2 | AJ0072 3 0 | DK0010 3 0 | DK0142 1 2 | JB0080 1 2 | JB0221 1 2 | JB0349 1 2 | JB0487 1 2 | JB0612 1 2 | JB0741 1 2 |
| A04095 1 2 | AJ0073 2 1 | DK0011 1 2 | DK0144 1 2 | JB0081 1 2 | JB0222 1 2 | JB0350 1 2 | JB0488 1 2 | JB0613 1 2 | JB0742 1 2 |
| A04096 1 2 | AJ0074 2 1 | DK0012 1 2 | DK0145 3 0 | JB0082 1 2 | JB0223 1 2 | JB0351 1 2 | JB0490 1 2 | JB0614 1 2 | JB0743 1 2 |
| A04097 1 2 | AJ0075 2 1 | DK0014 1 2 | DK0146 1 2 | JB0083 1 2 | JB0224 1 2 | JB0352 1 2 | JB0491 1 2 | JB0615 1 2 | JB0745 1 2 |
| A04098 1 2 | AJ0076 2 1 | DK0016 1 2 | DK0147 1 2 | JB0084 1 2 | JB0225 1 2 | JB0353 1 2 | JB0492 1 2 | JB0616 1 2 | JB0746 1 2 |
| A04100 1 2 | AJ0077 3 0 | DK0017 1 2 | DK0148 1 2 | JB0085 1 2 | JB0226 1 2 | JB0354 1 2 | JB0493 1 2 | JB0617 1 2 | JB0747 1 2 |
| A04101 1 2 | AJ0078 2 1 | DK0018 1 2 | DK0149 3 0 | JB0086 1 2 | JB0227 1 2 | JB0355 1 2 | JB0494 1 2 | JB0618 1 2 | JB0748 1 2 |
| A04102 1 2 | AJ0079 3 0 | DK0019 3 0 | DK0150 1 2 | JB0088 1 2 | JB0228 1 2 | JB0356 1 2 | JB0496 1 2 | JB0619 1 2 | JB0749 1 2 |
| A04103 1 2 | AJ0080 3 0 | DK0022 3 0 | DK0152 1 2 | JB0089 1 2 | JB0232 1 2 | JB0357 1 2 | JB0498 1 2 | JB0621 1 2 | JB0750 1 2 |
| A04104 1 2 | AJ0081 3 0 | DK0023 3 0 | DK0153 1 2 | JB0090 1 2 | JB0233 1 2 | JB0359 1 2 | JB0500 1 2 | JB0622 1 2 | JB0752 1 2 |
| A04106 1 2 | AJ0082 3 0 | DK0025 1 2 | DK0154 1 2 | JB0091 1 2 | JB0234 1 2 | JB0360 1 2 | JB0502 1 2 | JB0623 1 2 | JB0753 1 2 |
| A04107 1 2 | AJ0083 3 0 | DK0026 1 2 | DK0155 3 0 | JB0092 1 2 | JB0235 1 2 | JB0361 1 2 | JB0503 1 2 | JB0624 1 2 | JB0754 1 2 |
| A04109 1 2 | AJ0084 3 0 | DK0027 2 1 | DK0156 1 2 | JB0095 1 2 | JB0236 1 2 | JB0362 1 2 | JB0504 1 2 | JB0625 1 2 | JB0755 1 2 |
| A04110 1 2 | AJ0085 3 0 | DK0028 1 2 | DK0157 1 2 | JB0096 1 2 | JB0237 1 2 | JB0363 1 2 | JB0505 1 2 | JB0626 1 2 | JB0756 1 2 |
| A04111 1 2 | AJ0086 3 0 | DK0029 1 2 | DK0158 1 2 | JB0097 1 2 | JB0238 1 2 | JB0364 1 2 | JB0506 1 2 | JB0627 1 2 | JB0758 1 2 |
| A04112 1 2 | AJ0087 3 0 | DK0030 1 2 | DK0159 1 2 | JB0098 1 2 | JB0240 1 2 | JB0365 1 2 | JB0507 1 2 | JB0628 1 2 | JB0759 1 2 |
| A04113 1 2 | AJ0088 3 0 | DK0031 1 2 | DK0160 1 2 | JB0100 1 2 | JB0241 1 2 | JB0366 1 2 | JB0508 1 2 | JB0629 1 2 | JB0760 1 2 |
| A04115 1 2 | AJ0090 3 0 | DK0032 1 2 | DK0161 3 0 | JB0101 1 2 | JB0242 1 2 | JB0368 1 2 | JB0509 1 2 | JB0630 1 2 | JB0761 1 2 |
| A04117 1 2 | AJ0091 3 0 | DK0033 3 0 | DK0162 3 0 | JB0102 1 2 | JB0243 1 2 | JB0369 1 2 | JB0510 1 2 | JB0631 1 2 | JB0766 1 2 |
| A04118 1 2 | AJ0092 3 0 | DK0034 1 2 | DK0163 1 2 | JB0103 1 2 | JB0244 1 2 | JB0370 1 2 | JB0511 1 2 | JB0632 1 2 | JB0767 1 2 |
| A04119 1 2 | AJ0093 3 0 | DK0035 1 2 | DK0164 1 2 | JB0104 1 2 | JB0246 1 2 | JB0371 1 2 | JB0512 1 2 | JB0634 1 2 | JB0768 1 2 |
| A04120 1 2 | AJ0094 3 0 | DK0038 1 2 | DK0165 1 2 | JB0105 1 2 | JB0247 1 2 | JB0372 1 2 | JB0513 1 2 | JB0635 1 2 | JB0769 1 2 |
| A04121 1 2 | AJ0095 3 0 | DK0039 3 0 | DK0166 1 2 | JB0107 1 2 | JB0248 1 2 | JB0373 1 2 | JB0514 1 2 | JB0636 1 2 | JB0770 1 2 |
| A04123 1 2 | AJ0096 2 1 | DK0040 1 2 | DK0167 1 2 | JB0108 1 2 | JB0249 1 2 | JB0374 1 2 | JB0515 1 2 | JB0637 1 2 | JB0771 1 2 |
| A04125 1 2 | BM0071 3 0 | DK0041 3 0 | DK0168 1 2 | JB0110 1 2 | JB0250 1 2 | JB0377 1 2 | JB0516 1 2 | JB0638 1 2 | JB0772 1 2 |
| A04126 1 2 | BM0072 3 0 | DK0042 1 2 | DK0169 3 0 | JB0111 1 2 | JB0251 1 2 | JB0378 1 2 | JB0518 1 2 | JB0639 1 2 | JB0773 1 2 |
| A04128 1 2 | BM0073 3 0 | DK0043 3 0 | DK0170 1 2 | JB0112 1 2 | JB0252 1 2 | JB0379 1 2 | JB0519 1 2 | JB0640 1 2 | JB0774 1 2 |
| A04131 1 2 | BM0074 3 0 | DK0044 3 0 | DK0171 1 2 | JB0113 1 2 | JB0253 1 2 | JB0380 1 2 | JB0520 1 2 | JB0641 1 2 | JB0775 1 2 |
| A04132 1 2 | BM0076 3 0 | DK0045 3 0 | DK0172 3 0 | JB0114 1 2 | JB0254 1 2 | JB0381 1 2 | JB0521 1 2 | JB0643 1 2 | JB0776 1 2 |
| A04133 1 2 | BM0077 3 0 | DK0046 2 1 | DK0173 1 2 | JB0115 1 2 | JB0255 1 2 | JB0383 1 2 | JB0522 1 2 | JB0644 1 2 | JB0777 1 2 |
| A04134 1 2 | BM0078 3 0 | DK0047 1 2 | DK0174 1 2 | JB0116 1 2 | JB0256 1 2 | JB0384 1 2 | JB0523 1 2 | JB0645 1 2 | JB0778 1 2 |
| A04135 1 2 | BM0079 3 0 | DK0048 1 2 | DK0176 3 0 | JB0117 1 2 | JB0257 1 2 | JB0385 1 2 | JB0524 1 2 | JB0646 1 2 | JB0779 1 2 |
| A04136 1 2 | BM0081 3 0 | DK0049 1 2 | DK0177 1 2 | JB0118 1 2 | JB0258 1 2 | JB0387 1 2 | JB0525 1 2 | JB0647 1 2 | JB0780 1 2 |
| A04137 1 2 | BM0082 3 0 | DK0050 1 2 | DK0178 1 2 | JB0119 1 2 | JB0259 1 2 | JB0389 1 2 | JB0526 1 2 | JB0650 1 2 | JB0781 1 2 |
| A04138 1 2 | BM0083 3 0 | DK0051 1 2 | DK0179 1 2 | JB0121 1 2 | JB0260 1 2 | JB0390 1 2 | JB0527 1 2 | JB0651 1 2 | JB0782 1 2 |
| A04139 1 2 | BM0085 2 1 | DK0052 1 2 | DK0180 1 2 | JB0122 1 2 | JB0261 1 2 | JB0392 1 2 | JB0528 1 2 | JB0652 1 2 | JB0783 1 2 |
| A04144 1 2 | BM0100 3 0 | DK0053 1 2 | DK0181 1 2 | JB0123 1 2 | JB0262 1 2 | JB0393 1 2 | JB0529 1 2 | JB0653 1 2 | JB0784 1 2 |
| A04149 1 2 | BM0101 3 0 | DK0055 1 2 | DK0182 1 2 | JB0124 1 2 | JB0263 1 2 | JB0394 1 2 | JB0530 1 2 | JB0654 1 2 | JB0785 1 2 |
| A04150 1 2 | BM0102 3 0 | DK0056 1 2 | DK0183 1 2 | JB0125 1 2 | JB0264 1 2 | JB0396 1 2 | JB0531 1 2 | JB0655 1 2 | JB0786 1 2 |
| A04151 1 2 | BM0103 3 0 | DK0058 1 2 | DK0184 1 2 | JB0126 1 2 | JB0265 1 2 | JB0397 1 2 | JB0532 1 2 | JB0656 1 2 | JB0787 1 2 |
| A04152 1 2 | BM0104 3 0 | DK0060 1 2 | DK0185 1 2 | JB0127 1 2 | JB0267 1 2 | JB0398 1 2 | JB0533 1 2 | JB0657 1 2 | JB0788 1 2 |
| AJ0001 3 0 | BM0105 3 0 | DK0061 3 0 | DK0186 3 0 | JB0128 1 2 | JB0268 1 2 | JB0399 1 2 | JB0534 1 2 | JB0658 1 2 | JB0789 1 2 |
| AJ0002 3 0 | BM0108 3 0 | DK0062 1 2 | DK0187 3 0 | JB0129 1 2 | JB0269 1 2 | JB0400 1 2 | JB0535 1 2 | JB0660 1 2 | JB0790 1 2 |
| AJ0003 3 0 | BM0109 3 0 | DK0065 3 0 | DK0188 1 2 | JB0130 1 2 | JB0270 1 2 | JB0401 1 2 | JB0536 1 2 | JB0661 1 2 | JB0791 1 2 |
| AJ0004 3 0 | BM0110 3 0 | DK0066 1 2 | DK0189 1 2 | JB0132 1 2 | JB0271 1 2 | JB0402 1 2 | JB0537 1 2 | JB0662 1 2 | JB0792 1 2 |
| AJ0005 3 0 | BM0111 3 0 | DK0067 3 0 | DK0190 1 2 | JB0135 1 2 | JB0272 1 2 | JB0404 1 2 | JB0538 1 2 | JB0663 1 2 | JB0793 1 2 |
| AJ0006 3 0 | BM0113 3 0 | DK0068 1 2 | DK0191 1 2 | JB0136 1 2 | JB0273 1 2 | JB0405 1 2 | JB0539 1 2 | JB0665 1 2 | JB0794 1 2 |
| AJ0007 3 0 | BM0114 3 0 | DK0069 1 2 | DK0192 3 0 | JB0137 1 2 | JB0274 1 2 | JB0406 1 2 | JB0540 1 2 | JB0666 1 2 | JB0795 1 2 |
| AJ0008 3 0 | BM0115 3 0 | DK0070 3 0 | G19002 1 2 | JB0138 1 2 | JB0275 1 2 | JB0407 1 2 | JB0541 1 2 | JB0667 1 2 | JB0796 1 2 |
| AJ0009 3 0 | BM0116 3 0 | DK0071 1 2 | JB0004 1 2 | JB0139 1 2 | JB0276 1 2 | JB0408 1 2 | JB0542 1 2 | JB0668 1 2 | JB0797 1 2 |
| AJ0010 3 0 | BM0117 3 0 | DK0072 1 2 | JB0006 1 2 | JB0140 1 2 | JB0277 1 2 | JB0409 1 2 | JB0543 1 2 | JB0669 1 2 | JB0798 1 2 |
| AJ0011 3 0 | BM0118 3 0 | DK0075 1 2 | JB0007 1 2 | JB0141 1 2 | JB0278 1 2 | JB0410 1 2 | JB0544 1 2 | JB0670 1 2 | JB0799 1 2 |
| AJ0012 3 0 | BM0119 3 0 | DK0076 3 0 | JB0008 1 2 | JB0142 1 2 | JB0279 1 2 | JB0412 1 2 | JB0545 1 2 | JB0671 1 2 | JB0800 1 2 |
| AJ0013 3 0 | BM0120 3 0 | DK0077 1 2 | JB0009 1 2 | JB0143 1 2 | JB0280 1 2 | JB0413 1 2 | JB0547 1 2 | JB0673 1 2 | JB0801 1 2 |
| AJ0014 3 0 | BM0121 3 0 | DK0078 1 2 | JB0010 1 2 | JB0144 1 2 | JB0281 1 2 | JB0414 1 2 | JB0548 1 2 | JB0674 1 2 | JB0802 1 2 |
| AJ0015 3 0 | BM0122 3 0 | DK0079 1 2 | JB0011 1 2 | JB0145 1 2 | JB0282 1 2 | JB0416 1 2 | JB0549 1 2 | JB0675 1 2 | JB0803 1 2 |
| AJ0016 3 0 | BM0123 3 0 | DK0081 1 2 | JB0012 1 2 | JB0146 1 2 | JB0283 1 2 | JB0418 1 2 | JB0550 1 2 | JB0676 1 2 | JB0804 1 2 |
| AJ0017 3 0 | BM0124 3 0 | DK0082 1 2 | JB0013 1 2 | JB0147 1 2 | JB0284 1 2 | JB0419 1 2 | JB0551 1 2 | JB0677 1 2 | JB0805 1 2 |
| AJ0018 3 0 | BM0126 3 0 | DK0084 3 0 | JB0014 1 2 | JB0148 1 2 | JB0285 1 2 | JB0420 1 2 | JB0552 1 2 | JB0678 1 2 | JB0806 1 2 |
| AJ0019 3 0 | BM0127 3 0 | DK0085 1 2 | JB0015 1 2 | JB0149 1 2 | JB0286 1 2 | JB0421 1 2 | JB0553 1 2 | JB0679 1 2 | JB0807 1 2 |
| AJ0020 3 0 | BM0133 3 0 | DK0086 3 0 | JB0016 1 2 | JB0150 1 2 | JB0288 1 2 | JB0423 1 2 | JB0554 1 2 | JB0680 1 2 | JB0808 1 2 |
| AJ0021 3 0 | BM0134 3 0 | DK0087 1 2 | JB0017 1 2 | JB0151 1 2 | JB0289 1 2 | JB0424 1 2 | JB0555 1 2 | JB0681 1 2 | JB0809 1 2 |
| AJ0022 3 0 | BM0135 3 0 | DK0088 1 2 | JB0018 1 2 | JB0152 1 2 | JB0290 1 2 | JB0425 1 2 | JB0556 1 2 | JB0682 1 2 | JB0810 1 2 |
| AJ0023 3 0 | BM0136 3 0 | DK0090 1 2 | JB0020 1 2 | JB0154 1 2 | JB0291 1 2 | JB0426 1 2 | JB0557 1 2 | JB0683 1 2 | JB0811 1 2 |
| AJ0024 3 0 | BM0138 3 0 | DK0091 1 2 | JB0021 1 2 | JB0155 1 2 | JB0292 1 2 | JB0427 1 2 | JB0558 1 2 | JB0684 1 2 | JB0812 1 2 |
| AJ0025 3 0 | BM0139 3 0 | DK0092 1 2 | JB0023 1 2 | JB0156 1 2 | JB0293 1 2 | JB0428 1 2 | JB0559 1 2 | JB0685 1 2 | JB0813 1 2 |
| AJ0026 3 0 | BM0140 3 0 | DK0093 3 0 | JB0024 1 2 | JB0157 1 2 | JB0294 1 2 | JB0430 1 2 | JB0562 1 2 | JB0686 1 2 | JB0814 1 2 |
| AJ0027 3 0 | BM0141 3 0 | DK0097 1 2 | JB0025 1 2 | JB0158 1 2 | JB0295 1 2 | JB0431 1 2 | JB0563 1 2 | JB0687 1 2 | JB0817 1 2 |
| AJ0028 3 0 | BM0145 3 0 | DK0098 3 0 | JB0026 1 2 | JB0160 1 2 | JB0296 1 2 | JB0432 1 2 | JB0564 1 2 | JB0688 1 2 | JB0818 1 2 |
| AJ0029 3 0 | BM0146 3 0 | DK0099 1 2 | JB0027 1 2 | JB0161 1 2 | JB0297 1 2 | JB0433 1 2 | JB0565 1 2 | JB0689 1 2 | JB0819 1 2 |
| AJ0030 3 0 | BM0147 3 0 | DK0100 1 2 | JB0028 1 2 | JB0164 1 2 | JB0298 1 2 | JB0434 1 2 | JB0566 1 2 | JB0690 1 2 | JB0820 1 2 |
| AJ0031 3 0 | BM0148 3 0 | DK0101 3 0 | JB0029 1 2 | JB0165 1 2 | JB0300 1 2 | JB0435 1 2 | JB0567 1 2 | JB0691 1 2 | JB0821 1 2 |
| AJ0032 3 0 | BM0149 3 0 | DK0102 1 2 | JB0030 1 2 | JB0166 1 2 | JB0301 1 2 | JB0436 1 2 | JB0568 1 2 | JB0692 1 2 | JB0822 1 2 |
| AJ0033 3 0 | BM0150 3 0 | DK0103 1 2 | JB0031 1 2 | JB0167 1 2 | JB0302 1 2 | JB0437 1 2 | JB0569 1 2 | JB0693 1 2 | JB0823 1 2 |
| AJ0034 3 0 | BM0151 3 0 | DK0104 1 2 | JB0032 1 2 | JB0168 1 2 | JB0304 1 2 | JB0438 1 2 | JB0570 1 2 | JB0694 1 2 | JB0824 1 2 |
| AJ0035 3 0 | BM0152 3 0 | DK0105 1 2 | JB0033 1 2 | JB0169 1 2 | JB0306 1 2 | JB0439 1 2 | JB0571 1 2 | JB0695 1 2 | JB0825 1 2 |
| AJ0036 3 0 | BM0153 3 0 | DK0106 1 2 | JB0034 1 2 | JB0170 1 2 | JB0307 1 2 | JB0440 1 2 | JB0572 1 2 | JB0696 1 2 | JB0826 1 2 |
| AJ0037 3 0 | BM0154 3 0 | DK0107 3 0 | JB0035 1 2 | JB0172 1 2 | JB0309 1 2 | JB0441 1 2 | JB0573 1 2 | JB0697 1 2 | JB0827 1 2 |
| AJ0038 3 0 | BM0156 3 0 | DK0108 1 2 | JB0036 1 2 | JB0173 1 2 | JB0310 1 2 | JB0442 1 2 | JB0575 1 2 | JB0699 1 2 | JB0828 1 2 |
| AJ0039 3 0 | BM0157 3 0 | DK0109 1 2 | JB0037 1 2 | JB0174 1 2 | JB0311 1 2 | JB0444 1 2 | JB0576 1 2 | JB0700 1 2 | JB0829 1 2 |
| AJ0040 3 0 | BM0158 3 0 | DK0110 3 0 | JB0038 1 2 | JB0175 1 2 | JB0313 1 2 | JB0445 1 2 | JB0577 1 2 | JB0701 1 2 | JB0830 1 2 |
| AJ0041 3 0 | BM0159 3 0 | DK0111 1 2 | JB0039 1 2 | JB0177 1 2 | JB0314 1 2 | JB0446 1 2 | JB0578 1 2 | JB0702 1 2 | JB0831 1 2 |
| AJ0042 3 0 | BM0160 3 0 | DK0112 3 0 | JB0040 1 2 | JB0179 1 2 | JB0315 1 2 | JB0447 1 2 | JB0579 1 2 | JB0703 1 2 | JB0832 1 2 |
| AJ0043 3 0 | BM0162 3 0 | DK0113 1 2 | JB0041 1 2 | JB0180 1 2 | JB0316 1 2 | JB0448 1 2 | JB0580 1 2 | JB0704 1 2 | JB0833 1 2 |
| AJ0044 3 0 | BM0164 3 0 | DK0114 1 2 | JB0042 1 2 | JB0181 1 2 | JB0317 1 2 | JB0449 1 2 | JB0581 1 2 | JB0705 1 2 | JB0834 1 2 |
| AJ0045 3 0 | BM0165 3 0 | DK0115 1 2 | JB0043 1 2 | JB0182 1 2 | JB0318 1 2 | JB0450 1 2 | JB0582 1 2 | JB0706 1 2 | JB0835 1 2 |
| AJ0046 3 0 | BM0166 3 0 | DK0116 3 0 | JB0044 1 2 | JB0183 1 2 | JB0319 1 2 | JB0452 1 2 | JB0583 1 2 | JB0707 1 2 | JB0837 1 2 |
| AJ0047 3 0 | BM0167 3 0 | DK0117 3 0 | JB0045 1 2 | JB0186 1 2 | JB0320 1 2 | JB0454 1 2 | JB0584 1 2 | JB0708 1 2 | JB0838 1 2 |
| AJ0048 3 0 | BM0168 3 0 | DK0118 1 2 | JB0046 1 2 | JB0187 1 2 | JB0321 1 2 | JB0456 1 2 | JB0585 1 2 | JB0709 1 2 | JB0840 1 2 |
| AJ0049 3 0 | BM0169 3 0 | DK0119 3 0 | JB0048 1 2 | JB0188 1 2 | JB0322 1 2 | JB0458 1 2 | JB0586 1 2 | JB0710 1 2 | JB0841 1 2 |
| AJ0050 3 0 | BM0174 3 0 | DK0120 3 0 | JB0049 1 2 | JB0189 1 2 | JB0323 1 2 | JB0459 1 2 | JB0587 1 2 | JB0711 1 2 | JB0842 1 2 |
| AJ0051 3 0 | BM0175 3 0 | DK0121 3 0 | JB0050 1 2 | JB0190 1 2 | JB0325 1 2 | JB0462 1 2 | JB0588 1 2 | JB0712 1 2 | JB0843 1 2 |
| AJ0052 3 0 | BM0177 3 0 | DK0122 1 2 | JB0052 1 2 | JB0191 1 2 | JB0326 1 2 | JB0463 1 2 | JB0589 1 2 | JB0713 1 2 | JB0844 1 2 |
| AJ0053 3 0 | BM0178 3 0 | DK0123 1 2 | JB0053 1 2 | JB0192 1 2 | JB0327 1 2 | JB0464 1 2 | JB0590 1 2 | JB0714 1 2 | JB0846 1 2 |
| AJ0054 3 0 | BM0179 3 0 | DK0124 1 2 | JB0054 1 2 | JB0193 1 2 | JB0329 1 2 | JB0465 1 2 | JB0591 1 2 | JB0715 1 2 | JB0847 1 2 |
| AJ0055 3 0 | BM0180 3 0 | DK0125 3 0 | JB0056 1 2 | JB0197 1 2 | JB0330 1 2 | JB0466 1 2 | JB0592 1 2 | JB0716 1 2 | JB0848 1 2 |
| AJ0056 3 0 | BM0183 3 0 | DK0126 1 2 | JB0057 1 2 | JB0199 1 2 | JB0331 1 2 | JB0467 1 2 | JB0593 1 2 | JB0717 1 2 | JB0849 1 2 |
| AJ0057 3 0 | BM0185 3 0 | DK0127 1 2 | JB0058 1 2 | JB0200 1 2 | JB0332 1 2 | JB0468 1 2 | JB0594 1 2 | JB0718 1 2 | JB0850 1 2 |
| AJ0058 3 0 | BM0186 3 0 | DK0128 3 0 | JB0059 1 2 | JB0202 1 2 | JB0333 1 2 | JB0469 1 2 | JB0595 1 2 | JB0719 1 2 | JB0851 1 2 |
| AJ0059 3 0 | BM0187 3 0 | DK0129 1 2 | JB0060 1 2 | JB0204 1 2 | JB0334 1 2 | JB0470 1 2 | JB0596 1 2 | JB0720 1 2 | JB0852 1 2 |
| AJ0060 3 0 | BM0188 3 0 | DK0130 1 2 | JB0062 1 2 | JB0205 1 2 | JB0335 1 2 | JB0471 1 2 | JB0597 1 2 | JB0721 1 2 | JB0853 1 2 |

| | | |
|---|---|---|
| JB0854 | 1 | 2 |
| JB0855 | 1 | 2 |
| JB0856 | 1 | 2 |
| JB0857 | 1 | 2 |
| JB0858 | 1 | 2 |
| JB0859 | 1 | 2 |
| JB0861 | 1 | 2 |
| JB0863 | 1 | 2 |
| JB0864 | 1 | 2 |
| JB0865 | 1 | 2 |
| JB0867 | 1 | 2 |
| JB0868 | 1 | 2 |
| JB0870 | 1 | 2 |
| JB0871 | 1 | 2 |
| JB0872 | 1 | 2 |
| JB0873 | 1 | 2 |
| JB0874 | 1 | 2 |
| JB0875 | 1 | 2 |
| JB0876 | 1 | 2 |
| JB0877 | 1 | 2 |
| JB0878 | 1 | 2 |
| JB0879 | 1 | 2 |
| JB0881 | 1 | 2 |
| JB0882 | 1 | 2 |
| JB0883 | 1 | 2 |
| JB0884 | 1 | 2 |
| JB0885 | 1 | 2 |
| JB0886 | 1 | 2 |
| JB0889 | 1 | 2 |
| JB0890 | 1 | 2 |
| JB0891 | 1 | 2 |
| JB0893 | 1 | 2 |
| JB0894 | 1 | 2 |
| JB0895 | 1 | 2 |
| JB0896 | 1 | 2 |
| JB0897 | 1 | 2 |
| JB0898 | 1 | 2 |
| LB0001 | 2 | 1 |
| LB0002 | 2 | 1 |
| LB0003 | 2 | 1 |
| LB0004 | 2 | 1 |
| LB0005 | 2 | 1 |
| LB0006 | 2 | 1 |
| LB0007 | 2 | 1 |
| LB0008 | 2 | 1 |
| LB0009 | 3 | 0 |
| LB0010 | 2 | 1 |
| LB0011 | 3 | 0 |
| LB0012 | 2 | 1 |
| LB0013 | 2 | 1 |
| LB0014 | 2 | 1 |
| LB0015 | 2 | 1 |
| LB0016 | 2 | 1 |
| LB0017 | 3 | 0 |
| LB0018 | 2 | 1 |
| LB0019 | 2 | 1 |
| LB0020 | 2 | 1 |
| LB0021 | 2 | 1 |
| LB0022 | 2 | 1 |
| LB0023 | 2 | 1 |
| LB0024 | 1 | 2 |
| LB0025 | 1 | 2 |
| LB0026 | 3 | 0 |
| LB0027 | 2 | 1 |
| LB0028 | 2 | 1 |
| LB0029 | 2 | 1 |
| LB0030 | 3 | 0 |
| LB0032 | 1 | 2 |
| LB0035 | 2 | 1 |
| LB0036 | 2 | 1 |
| LB0037 | 3 | 0 |
| LB0038 | 1 | 2 |
| LB0039 | 1 | 2 |
| LB0040 | 3 | 0 |
| LB0041 | 1 | 2 |
| LB0042 | 2 | 1 |
| LB0043 | 3 | 0 |
| LB0044 | 3 | 0 |
| LB0045 | 2 | 1 |
| LB0046 | 2 | 1 |
| LB0047 | 3 | 0 |
| LB0049 | 1 | 2 |
| LB0050 | 3 | 0 |
| LB0051 | 1 | 2 |
| LB0052 | 3 | 0 |
| LB0053 | 1 | 2 |
| LB0054 | 2 | 1 |
| LB0055 | 1 | 2 |
| LB0056 | 3 | 0 |
| LB0057 | 1 | 2 |
| LB0058 | 2 | 1 |
| LB0059 | 2 | 1 |
| LB0060 | 3 | 0 |
| LB0061 | 2 | 1 |
| LB0062 | 1 | 2 |
| LB0064 | 3 | 0 |
| LB0065 | 1 | 2 |
| LB0066 | 3 | 0 |
| LB0067 | 1 | 2 |
| LB0068 | 3 | 0 |
| LB0069 | 3 | 0 |
| LB0070 | 3 | 0 |
| LB0071 | 3 | 0 |
| LB0072 | 1 | 2 |
| LB0074 | 3 | 0 |
| LB0076 | 2 | 1 |
| LB0077 | 1 | 2 |
| LB0078 | 1 | 2 |
| LB0079 | 3 | 0 |
| LB0079 | 3 | 0 |
| LB0080 | 3 | 0 |
| LB0081 | 3 | 0 |
| LB0082 | 2 | 1 |
| LB0083 | 1 | 2 |
| LB0084 | 3 | 0 |
| LB0085 | 1 | 2 |
| LB0086 | 1 | 2 |
| LB0087 | 2 | 1 |
| LB0088 | 3 | 0 |
| LB0089 | 1 | 2 |
| LB0090 | 3 | 0 |
| LB0092 | 1 | 2 |
| LB0096 | 1 | 2 |
| LB0099 | 1 | 2 |
| LB0100 | 1 | 2 |
| LB0101 | 1 | 2 |
| LB0104 | 3 | 0 |
| LB0105 | 3 | 0 |
| LB0106 | 1 | 2 |
| LB0107 | 1 | 2 |
| LB0108 | 1 | 2 |
| LB0109 | 3 | 0 |
| LB0111 | 1 | 2 |
| LB0112 | 1 | 2 |
| LB0113 | 1 | 2 |
| LB0114 | 1 | 2 |
| LB0115 | 1 | 2 |
| LB0116 | 1 | 2 |
| LB0117 | 1 | 2 |
| LB0118 | 1 | 2 |
| LB0119 | 1 | 2 |
| LB0120 | 3 | 0 |
| LB0121 | 3 | 0 |
| LB0122 | 3 | 0 |
| LB0123 | 3 | 0 |
| LB0125 | 3 | 0 |
| LB0126 | 1 | 2 |
| LB0127 | 1 | 2 |
| LB0128 | 1 | 2 |
| LB0129 | 1 | 2 |
| LB0130 | 1 | 2 |
| LB0131 | 3 | 0 |
| LB0132 | 3 | 0 |
| LB0133 | 1 | 2 |
| LB0134 | 1 | 2 |
| LB0135 | 1 | 2 |
| LB0136 | 1 | 2 |
| LB0137 | 1 | 2 |
| LB0138 | 1 | 2 |
| LB0142 | 2 | 1 |
| LB0143 | 1 | 2 |
| LB0144 | 1 | 2 |
| LB0145 | 1 | 2 |
| LB0146 | 1 | 2 |
| LB0147 | 3 | 0 |
| LB0148 | 1 | 2 |
| LB0149 | 2 | 1 |
| LB0150 | 2 | 1 |
| LB0151 | 2 | 1 |
| LB0152 | 3 | 0 |
| LB0153 | 3 | 0 |
| LB0154 | 3 | 0 |
| LB0158 | 1 | 2 |
| LB0159 | 1 | 2 |
| LB0160 | 1 | 2 |
| LB0161 | 2 | 1 |
| LB0162 | 1 | 2 |
| LB0163 | 1 | 2 |
| LB0164 | 3 | 0 |
| LB0165 | 2 | 1 |
| LB0166 | 2 | 1 |
| LB0167 | 2 | 1 |
| LB0168 | 2 | 1 |
| LB0169 | 2 | 1 |
| LB0170 | 2 | 1 |
| LB0171 | 1 | 2 |
| LB0172 | 2 | 1 |
| LB0173 | 2 | 1 |
| LB0174 | 2 | 1 |
| LB0176 | 2 | 1 |
| LB0177 | 3 | 0 |
| LB0178 | 3 | 0 |
| LB0179 | 1 | 2 |
| LB0180 | 1 | 2 |
| LB0181 | 1 | 2 |
| LB0182 | 1 | 2 |
| LB0183 | 1 | 2 |
| LB0184 | 3 | 0 |
| LB0185 | 1 | 2 |
| LB0186 | 1 | 2 |
| LB0187 | 1 | 2 |
| LB0188 | 3 | 0 |
| LB0189 | 3 | 0 |
| LB0190 | 1 | 2 |
| LB0195 | 3 | 0 |
| LB0196 | 1 | 2 |
| LB0197 | 1 | 2 |
| LB0198 | 2 | 1 |
| LB0199 | 3 | 0 |
| LB0202 | 2 | 1 |
| LB0203 | 2 | 1 |
| LB0205 | 3 | 0 |
| LB0206 | 3 | 0 |
| LB0207 | 2 | 1 |
| LB0208 | 3 | 0 |
| LB0209 | 2 | 1 |
| LB0210 | 1 | 2 |
| LB0211 | 2 | 1 |
| LB0212 | 2 | 1 |
| LB0213 | 1 | 2 |
| LB0214 | 1 | 2 |
| LB0215 | 1 | 2 |
| LB0216 | 1 | 2 |
| LB0217 | 3 | 0 |
| LB0218 | 1 | 2 |
| LB0219 | 3 | 0 |
| LB0220 | 3 | 0 |
| LB0220 | 3 | 0 |
| LB0221 | 3 | 0 |
| LB0222 | 1 | 2 |
| LB0223 | 1 | 2 |
| LB0224 | 3 | 0 |
| LB0225 | 1 | 2 |
| LB0226 | 1 | 2 |
| LB0227 | 1 | 2 |
| LB0228 | 1 | 2 |
| LB0229 | 1 | 2 |
| LB0231 | 1 | 2 |
| LB0232 | 2 | 1 |
| LB0234 | 2 | 1 |
| LB0235 | 1 | 2 |
| LB0237 | 1 | 2 |
| LB0238 | 1 | 2 |
| LB0239 | 1 | 2 |
| LB0240 | 1 | 2 |
| LB0241 | 2 | 1 |
| LB0242 | 1 | 2 |
| LB0246 | 3 | 0 |
| LB0247 | 1 | 2 |
| LB0248 | 3 | 0 |
| LB0249 | 2 | 1 |
| LB0250 | 2 | 1 |
| LB0251 | 1 | 2 |
| LB0254 | 1 | 2 |
| LB0255 | 2 | 1 |
| LB0257 | 1 | 2 |
| LB0258 | 1 | 2 |
| LB0259 | 1 | 2 |
| LB0260 | 2 | 1 |
| LB0261 | 1 | 2 |
| LB0262 | 3 | 0 |
| LB0264 | 1 | 2 |
| LB0268 | 1 | 2 |
| LB0270 | 1 | 2 |
| LB0271 | 2 | 1 |
| LB0272 | 2 | 1 |
| LB0273 | 1 | 2 |
| LB0274 | 3 | 0 |
| LB0275 | 2 | 1 |
| LB0276 | 1 | 2 |
| LB0278 | 1 | 2 |
| LB0279 | 1 | 2 |
| LB0280 | 1 | 2 |
| LB0281 | 2 | 1 |
| LB0283 | 1 | 2 |
| LB0284 | 2 | 1 |
| LB0285 | 1 | 2 |
| LB0286 | 1 | 2 |
| LB0287 | 1 | 2 |
| LB0288 | 2 | 1 |
| LB0289 | 1 | 2 |
| LB0290 | 3 | 0 |
| LB0292 | 1 | 2 |
| LB0293 | 2 | 1 |
| LB0294 | 3 | 0 |
| LB0295 | 3 | 0 |
| LB0296 | 1 | 2 |
| LB0297 | 3 | 0 |
| LB0297 | 3 | 0 |
| LB0299 | 1 | 2 |
| LB0300 | 2 | 1 |
| LB0301 | 1 | 2 |
| LB0302 | 1 | 2 |
| LB0304 | 1 | 2 |
| LB0305 | 1 | 2 |
| LB0307 | 1 | 2 |
| LB0308 | 2 | 1 |
| LB0309 | 2 | 1 |
| LB0310 | 1 | 2 |
| LB0311 | 2 | 1 |
| LB0312 | 2 | 1 |
| LB0313 | 1 | 2 |
| LB0314 | 3 | 0 |
| LB0315 | 3 | 0 |
| LB0316 | 1 | 2 |
| LB0317 | 2 | 1 |
| LB0318 | 1 | 2 |
| LB0319 | 1 | 2 |
| LB0320 | 1 | 2 |
| LB0322 | 1 | 2 |
| LB0333 | 1 | 2 |
| LB0334 | 2 | 1 |
| LB0335 | 3 | 0 |
| LB0336 | 1 | 2 |
| LB0337 | 1 | 2 |
| LB0338 | 1 | 2 |
| LB0339 | 3 | 0 |
| LB0340 | 1 | 2 |
| LB0341 | 1 | 2 |
| LB0342 | 3 | 0 |
| LB0343 | 1 | 2 |
| LB0344 | 1 | 2 |
| LB0345 | 1 | 2 |
| LB0346 | 3 | 0 |
| LB0349 | 2 | 1 |
| LB0350 | 2 | 1 |
| LB0351 | 2 | 1 |
| LB0352 | 1 | 2 |
| LB0353 | 1 | 2 |
| LB0354 | 1 | 2 |
| LB0355 | 3 | 0 |
| LB0356 | 2 | 1 |
| LB0357 | 2 | 1 |
| LB0359 | 2 | 1 |
| LB0360 | 2 | 1 |
| LB0361 | 2 | 1 |
| LB0362 | 1 | 2 |
| LB0363 | 1 | 2 |
| LB0364 | 1 | 2 |
| LB0365 | 1 | 2 |
| LB0366 | 3 | 0 |
| LB0367 | 2 | 1 |
| LB0369 | 2 | 1 |
| LB0372 | 3 | 0 |
| LB0373 | 3 | 0 |
| LB0374 | 3 | 0 |
| LB0375 | 3 | 0 |
| LB0376 | 3 | 0 |
| LB0378 | 3 | 0 |
| LB0379 | 3 | 0 |
| LB0380 | 1 | 2 |
| LB0381 | 1 | 2 |
| LB0382 | 1 | 2 |
| LB0383 | 2 | 1 |
| LB0385 | 2 | 1 |
| LB0386 | 1 | 2 |
| LB0387 | 1 | 2 |
| LB0388 | 2 | 1 |
| LB0390 | 2 | 1 |
| LB0391 | 1 | 2 |
| LB0392 | 1 | 2 |
| LB0393 | 2 | 1 |
| LB0394 | 1 | 2 |
| LB0395 | 3 | 0 |
| LB0396 | 3 | 0 |
| LB0397 | 1 | 2 |
| LB0398 | 1 | 2 |
| LB0399 | 1 | 2 |
| LB0401 | 1 | 2 |
| LB0402 | 3 | 0 |
| LB0403 | 3 | 0 |
| LB0404 | 1 | 2 |
| LB0405 | 1 | 2 |
| LB0406 | 1 | 2 |
| LB0408 | 3 | 0 |
| LB0409 | 1 | 2 |
| LB0410 | 1 | 2 |
| LB0411 | 3 | 0 |
| LB0412 | 3 | 0 |
| LB0413 | 1 | 2 |
| LB0414 | 1 | 2 |
| LB0416 | 1 | 2 |
| LB0417 | 1 | 2 |
| LB0418 | 1 | 2 |
| LB0419 | 3 | 0 |
| LB0420 | 1 | 2 |
| LB0421 | 1 | 2 |
| LB0422 | 1 | 2 |
| LB0423 | 1 | 2 |
| LB0424 | 1 | 2 |
| LB0425 | 3 | 0 |
| LB0426 | 1 | 2 |
| LB0427 | 1 | 2 |
| LB0428 | 1 | 2 |
| LB0429 | 1 | 2 |
| LB0430 | 1 | 2 |
| LB0431 | 1 | 2 |
| LB0432 | 1 | 2 |
| LB0433 | 1 | 2 |
| LB0434 | 3 | 0 |
| LB0435 | 1 | 2 |
| LB0436 | 2 | 1 |
| LB0437 | 1 | 2 |
| LB0438 | 2 | 1 |
| LB0439 | 2 | 1 |
| LB0440 | 1 | 2 |
| LB0441 | 1 | 2 |
| LB0442 | 1 | 2 |
| LB0443 | 1 | 2 |
| LB0444 | 1 | 2 |
| LB0446 | 2 | 1 |
| LB0447 | 2 | 1 |
| LB0448 | 2 | 1 |
| LB0449 | 2 | 1 |
| LB0450 | 2 | 1 |
| LB0451 | 2 | 1 |
| LB0452 | 2 | 1 |
| LB0453 | 2 | 1 |
| LB0454 | 2 | 1 |
| LB0455 | 2 | 1 |
| LB0456 | 2 | 1 |
| LB0457 | 2 | 1 |
| LB0458 | 2 | 1 |
| LB0459 | 2 | 1 |
| LB0460 | 2 | 1 |
| LB0461 | 2 | 1 |
| LB0462 | 2 | 1 |
| LB0463 | 2 | 1 |
| LB0464 | 2 | 1 |
| LB0465 | 2 | 1 |
| LB0466 | 2 | 1 |
| LB0467 | 2 | 1 |
| LB0468 | 2 | 1 |
| LB0469 | 2 | 1 |
| LB0470 | 1 | 2 |
| LB0471 | 1 | 2 |
| LB0473 | 1 | 2 |
| LB0475 | 2 | 1 |
| LB0476 | 2 | 1 |
| LB0477 | 1 | 2 |
| LB0478 | 2 | 1 |
| LB0479 | 2 | 1 |
| LB0480 | 2 | 1 |
| LB0481 | 1 | 2 |
| LB0482 | 1 | 2 |
| LB0483 | 2 | 1 |
| LB0484 | 2 | 1 |
| LB0485 | 2 | 1 |
| LB0486 | 2 | 1 |
| LB0487 | 1 | 2 |
| LB0488 | 2 | 1 |
| LB0489 | 2 | 1 |
| LB0490 | 1 | 2 |
| LB0491 | 1 | 2 |
| LB0492 | 1 | 2 |
| LB0493 | 1 | 2 |
| LB0494 | 1 | 2 |
| LB0495 | 2 | 1 |
| LB0496 | 2 | 1 |
| LB0498 | 2 | 1 |
| LB0499 | 2 | 1 |
| LB0500 | 2 | 1 |
| LB0501 | 2 | 1 |
| LB0502 | 2 | 1 |
| LB0503 | 2 | 1 |
| LB0504 | 2 | 1 |
| LB0505 | 2 | 1 |
| LB0506 | 2 | 1 |
| LB0507 | 2 | 1 |
| LB0508 | 2 | 1 |
| LB0509 | 2 | 1 |
| LB0510 | 1 | 2 |
| LB0511 | 1 | 2 |
| LB0512 | 1 | 2 |
| LB0513 | 1 | 2 |
| LB0514 | 2 | 1 |
| LB0517 | 2 | 1 |
| LB0518 | 2 | 1 |
| LB0519 | 2 | 1 |
| LB0523 | 2 | 1 |
| LB0524 | 2 | 1 |
| LB0525 | 2 | 1 |
| LB0526 | 2 | 1 |
| LB0527 | 2 | 1 |
| LB0528 | 2 | 1 |
| LB0530 | 1 | 2 |
| LB0531 | 2 | 1 |
| LB0532 | 2 | 1 |
| LB0533 | 1 | 2 |
| LB0535 | 2 | 1 |
| LB0536 | 2 | 1 |
| LB0537 | 2 | 1 |
| LB0538 | 2 | 1 |
| LB0539 | 1 | 2 |
| LB0544 | 2 | 1 |
| LB0545 | 1 | 2 |
| LB0547 | 2 | 1 |
| LB0548 | 1 | 2 |
| LB0549 | 1 | 2 |
| LB0550 | 2 | 1 |
| LB0551 | 1 | 2 |
| LB0553 | 1 | 2 |
| LB0557 | 1 | 2 |
| LB0558 | 2 | 1 |
| LB0561 | 1 | 2 |
| LB0562 | 2 | 1 |
| LB0563 | 1 | 2 |
| LB0564 | 1 | 2 |
| LB0565 | 1 | 2 |
| LB0566 | 1 | 2 |
| LB0567 | 1 | 2 |
| LB0568 | 1 | 2 |
| LB0569 | 1 | 2 |
| LB0596 | 1 | 2 |
| LB0597 | 2 | 1 |
| LB0598 | 1 | 2 |
| LB0599 | 2 | 1 |
| LB0600 | 2 | 1 |
| LB0601 | 2 | 1 |
| LB0602 | 2 | 1 |
| LB0603 | 2 | 1 |
| LB0604 | 2 | 1 |
| LB0605 | 2 | 1 |
| LB0606 | 2 | 1 |
| LB0607 | 2 | 1 |
| LB0L05 | 3 | 0 |
| LB0L06 | 3 | 0 |
| LB0L14 | 3 | 0 |
| LB0L23 | 3 | 0 |
| LB0L24 | 2 | 1 |
| LB2282 | 1 | 2 |
| LB2283 | 2 | 1 |
| MD0001 | 1 | 2 |
| MD0002 | 1 | 2 |
| MD0003 | 1 | 2 |
| MD0033 | 3 | 0 |
| MD0004 | 1 | 2 |
| MD0005 | 1 | 2 |
| MD0006 | 1 | 2 |
| MD0069 | 3 | 0 |
| MD0007 | 1 | 2 |
| MD0071 | 3 | 0 |
| MD0008 | 1 | 2 |
| MD0010 | 1 | 2 |
| MD0012 | 1 | 2 |
| MD0013 | 1 | 2 |
| MD0013 | 3 | 0 |
| MD0014 | 1 | 2 |
| MD0016 | 1 | 2 |
| MD0021 | 1 | 2 |
| MD0024 | 1 | 2 |
| MD0025 | 1 | 2 |
| MD0026 | 1 | 2 |
| MD0028 | 1 | 2 |
| MD0029 | 1 | 2 |
| MD0029 | 1 | 2 |
| MD0030 | 1 | 2 |
| MD0030 | 1 | 2 |
| MD0031 | 1 | 2 |
| MD0032 | 1 | 2 |
| MD0033 | 1 | 2 |
| MD0033 | 1 | 2 |
| MD0034 | 1 | 2 |
| MD0034 | 1 | 2 |
| MD0035 | 1 | 2 |
| MD0036 | 1 | 2 |
| MD0036 | 1 | 2 |
| MD0037 | 1 | 2 |
| MD0039 | 1 | 2 |
| MD0042 | 1 | 2 |
| MD0044 | 1 | 2 |
| MD0044 | 1 | 2 |
| MD0045 | 1 | 2 |
| MD0046 | 1 | 2 |
| MD0047 | 1 | 2 |
| MD0048 | 1 | 2 |
| MD0049 | 1 | 2 |
| MD0056 | 1 | 2 |
| MD0058 | 1 | 2 |
| MD0059 | 1 | 2 |
| MD0060 | 1 | 2 |
| MD0061 | 1 | 2 |
| MD0063 | 1 | 2 |
| MD0064 | 1 | 2 |
| MD0064 | 1 | 2 |
| MD0065 | 1 | 2 |
| MD0065 | 1 | 2 |
| MD0066 | 1 | 2 |
| MD0067 | 1 | 2 |
| MD0067 | 1 | 2 |
| MD0068 | 1 | 2 |
| MD0069 | 1 | 2 |
| MD0069 | 1 | 2 |
| MD0071 | 1 | 2 |
| MD0071 | 1 | 2 |
| MD0072 | 1 | 2 |
| MD0073 | 1 | 2 |
| MD0074 | 1 | 2 |
| MD0075 | 1 | 2 |
| MD0076 | 1 | 2 |
| MD0077 | 1 | 2 |
| MD0078 | 1 | 2 |
| MD0081 | 1 | 2 |
| MD0082 | 1 | 2 |
| MD0083 | 1 | 2 |
| MD0085 | 1 | 2 |
| MD0087 | 1 | 2 |
| MD0089 | 1 | 2 |
| MD0090 | 1 | 2 |
| MD0091 | 1 | 2 |
| MD0092 | 1 | 2 |
| MD0093 | 1 | 2 |
| MD0094 | 1 | 2 |
| MD0095 | 1 | 2 |
| MD0096 | 1 | 2 |
| MD0097 | 1 | 2 |
| MD0097 | 1 | 2 |
| MD0098 | 1 | 2 |
| MD0098 | 1 | 2 |
| MD0099 | 1 | 2 |
| MD0099 | 1 | 2 |
| MD0100 | 1 | 2 |
| MD0101 | 1 | 2 |
| MD0101 | 1 | 2 |
| MD0102 | 1 | 2 |
| MD0103 | 1 | 2 |
| MD0104 | 1 | 2 |
| MD0105 | 1 | 2 |
| MD0106 | 1 | 2 |
| MD0106 | 1 | 2 |
| MD0107 | 1 | 2 |
| MD0107 | 1 | 2 |
| MD0108 | 1 | 2 |
| MD0109 | 1 | 2 |
| MD0109 | 1 | 2 |
| MD0110 | 1 | 2 |
| MD0111 | 1 | 2 |
| MD0111 | 1 | 2 |
| MD0112 | 1 | 2 |
| MD0113 | 1 | 2 |
| MD0114 | 1 | 2 |
| MD0115 | 1 | 2 |
| MD0116 | 1 | 2 |
| MD0117 | 1 | 2 |
| MD0118 | 1 | 2 |
| MD0118 | 1 | 2 |
| MD0119 | 1 | 2 |
| MD0119 | 1 | 2 |
| MD0120 | 1 | 2 |
| MD0121 | 1 | 2 |
| MD0122 | 1 | 2 |
| MD0123 | 1 | 2 |
| MD0124 | 1 | 2 |
| MD0125 | 1 | 2 |
| MD0126 | 1 | 2 |
| MD0126 | 1 | 2 |
| MD0127 | 1 | 2 |
| MD0127 | 1 | 2 |
| MD0128 | 1 | 2 |
| MD0129 | 1 | 2 |
| MD0129 | 1 | 2 |
| MD0130 | 1 | 2 |
| MD0130 | 1 | 2 |
| MD0131 | 1 | 2 |
| MD0132 | 1 | 2 |
| MD0132 | 1 | 2 |
| MD0133 | 1 | 2 |
| MD0133 | 1 | 2 |
| MD0134 | 1 | 2 |
| MD0134 | 1 | 2 |
| MD0135 | 1 | 2 |
| MD0135 | 1 | 2 |
| MD0136 | 1 | 2 |
| MD0136 | 1 | 2 |
| MD0137 | 1 | 2 |
| MD0137 | 1 | 2 |
| MD0138 | 1 | 2 |
| MD0139 | 1 | 2 |
| MD0140 | 1 | 2 |
| MD0140 | 1 | 2 |
| MD0142 | 1 | 2 |
| MD0143 | 1 | 2 |
| MD0143 | 1 | 2 |
| MD0144 | 1 | 2 |
| MD0145 | 1 | 2 |
| MD0145 | 1 | 2 |
| MD0146 | 1 | 2 |
| MD0147 | 1 | 2 |
| MD0150 | 1 | 2 |
| MD0152 | 1 | 2 |
| MD0153 | 1 | 2 |
| MD0154 | 1 | 2 |
| MD0155 | 1 | 2 |
| MD0155 | 1 | 2 |
| MD0156 | 1 | 2 |
| MD0156 | 1 | 2 |
| MD0157 | 1 | 2 |
| MD0157 | 1 | 2 |
| MD0158 | 1 | 2 |
| MD0159 | 1 | 2 |
| MD0161 | 1 | 2 |
| MD0162 | 1 | 2 |
| MD0163 | 1 | 2 |
| MD0167 | 1 | 2 |
| MD0169 | 1 | 2 |
| MD0170 | 1 | 2 |
| MD0171 | 1 | 2 |
| MD0172 | 1 | 2 |
| MD0173 | 1 | 2 |
| MD0174 | 1 | 2 |
| MD0175 | 1 | 2 |
| MD0176 | 1 | 2 |
| MD0177 | 1 | 2 |
| MD0178 | 1 | 2 |
| MD0179 | 1 | 2 |
| MD0180 | 1 | 2 |
| MD0181 | 1 | 2 |
| MD0182 | 1 | 2 |
| MD0183 | 1 | 2 |
| MD0184 | 1 | 2 |
| MD0185 | 1 | 2 |
| MD0186 | 1 | 2 |
| MD0187 | 1 | 2 |
| MD0188 | 1 | 2 |
| MD0189 | 1 | 2 |
| MD0190 | 1 | 2 |
| MD0191 | 1 | 2 |
| MD0192 | 1 | 2 |
| MD0193 | 1 | 2 |
| MD0194 | 1 | 2 |
| MD0198 | 1 | 2 |
| MD0199 | 1 | 2 |
| MD0200 | 1 | 2 |
| MD0201 | 1 | 2 |
| MK0001 | 1 | 2 |
| MK0002 | 3 | 0 |
| MK0003 | 2 | 1 |
| MK0004 | 3 | 0 |
| MK0005 | 2 | 1 |
| MK0006 | 1 | 2 |
| MK0008 | 1 | 2 |
| MK0009 | 2 | 1 |
| MK0010 | 1 | 2 |
| MK0011 | 1 | 2 |
| MK0012 | 1 | 2 |
| MK0013 | 3 | 0 |
| MK0014 | 1 | 2 |
| MK0015 | 1 | 2 |
| MK0016 | 1 | 2 |
| MK0041 | 3 | 0 |
| MK0061 | 1 | 2 |
| MK0104 | 3 | 0 |
| MK0129 | 3 | 0 |
| MK0130 | 1 | 2 |
| MK0131 | 1 | 2 |
| MK0132 | 1 | 2 |
| MK0133 | 1 | 2 |
| MK0215 | 3 | 0 |
| MK0231 | 2 | 1 |
| MK0236 | 1 | 2 |
| MK0243 | 1 | 2 |
| MK0244 | 3 | 0 |
| MK0245 | 1 | 2 |
| MK0327 | 3 | 0 |
| MK0334 | 3 | 0 |
| MK0336 | 2 | 1 |
| MK0344 | 1 | 2 |
| MK0358 | 2 | 1 |
| MK0359 | 1 | 2 |
| MK0361 | 3 | 0 |
| MK0362 | 1 | 2 |
| MK0363 | 3 | 0 |
| MK0390 | 2 | 1 |
| MK0392 | 3 | 0 |
| MK0393 | 2 | 1 |
| MK0394 | 2 | 1 |
| MK0406 | 3 | 0 |
| MK0421 | 2 | 1 |
| MK0423 | 3 | 0 |
| MK0424 | 1 | 2 |
| MK0425 | 1 | 2 |
| MK0426 | 2 | 1 |
| MK0427 | 2 | 1 |
| MK0428 | 1 | 2 |
| MK0430 | 1 | 2 |
| MK0431 | 1 | 2 |
| MK0432 | 1 | 2 |
| MK0433 | 1 | 2 |
| MK0434 | 1 | 2 |
| MK0435 | 3 | 0 |
| MK0438 | 1 | 2 |
| MK0439 | 3 | 0 |
| MK0440 | 2 | 1 |
| MK0441 | 2 | 1 |
| MK0442 | 3 | 0 |
| MK0443 | 1 | 2 |
| MK0444 | 1 | 2 |
| MK0445 | 3 | 0 |
| MK0447 | 1 | 2 |
| MK0448 | 2 | 1 |
| MK0449 | 2 | 1 |
| MK0450 | 1 | 2 |
| MK0451 | 2 | 1 |
| MK0452 | 2 | 1 |
| MK0453 | 2 | 1 |
| MK0454 | 3 | 0 |
| MK0456 | 1 | 2 |
| MK0457 | 1 | 2 |
| MK0458 | 1 | 2 |
| MK0459 | 2 | 1 |
| MK0460 | 1 | 2 |
| MK0461 | 2 | 1 |
| MK0462 | 1 | 2 |
| MK0463 | 1 | 2 |
| MK0464 | 3 | 0 |
| MK0465 | 2 | 1 |
| MK0466 | 2 | 1 |
| MK0467 | 1 | 2 |
| MK0469 | 1 | 2 |
| MK0470 | 3 | 0 |
| MK0471 | 3 | 0 |
| MK0472 | 1 | 2 |
| MK0473 | 3 | 0 |
| MK0474 | 3 | 0 |
| MK0475 | 3 | 0 |
| MK0476 | 3 | 0 |
| MK0477 | 2 | 1 |
| MK0478 | 1 | 2 |
| MK0479 | 2 | 1 |
| MK0480 | 2 | 1 |
| MK0481 | 1 | 2 |
| MK0482 | 1 | 2 |
| MK0483 | 1 | 2 |
| MK0484 | 1 | 2 |
| MK0485 | 1 | 2 |
| MK0486 | 1 | 2 |
| MK0487 | 1 | 2 |
| MK0488 | 3 | 0 |
| MK0489 | 3 | 0 |
| MK0490 | 3 | 0 |
| MK0492 | 3 | 0 |
| MK0493 | 2 | 1 |
| MK0494 | 1 | 2 |
| MK0496 | 1 | 2 |
| MK0497 | 1 | 2 |
| MK0498 | 1 | 2 |
| MK0500 | 1 | 2 |
| MK0501 | 3 | 0 |
| MK0502 | 2 | 1 |
| MK0522 | 3 | 0 |
| MK0532 | 3 | 0 |
| MK0533 | 2 | 1 |
| MK0534 | 2 | 1 |
| MK0535 | 2 | 1 |
| MK0536 | 2 | 1 |
| MK0537 | 2 | 1 |
| MK0539 | 2 | 1 |
| MK0540 | 1 | 2 |
| MK0541 | 1 | 2 |
| MK0542 | 1 | 2 |
| MK0543 | 2 | 1 |
| MK0544 | 2 | 1 |
| MK0545 | 1 | 2 |
| MK0546 | 1 | 2 |
| MK0547 | 1 | 2 |
| MK0548 | 1 | 2 |
| MK0549 | 1 | 2 |
| MK0550 | 1 | 2 |
| MK0551 | 2 | 1 |
| MK0552 | 2 | 1 |
| MK0553 | 1 | 2 |
| MK0561 | 2 | 1 |
| MK0563 | 1 | 2 |
| MK0564 | 1 | 2 |
| MK0566 | 1 | 2 |
| MK0567 | 1 | 2 |
| MK0568 | 2 | 1 |
| MK0569 | 1 | 2 |
| MK0577 | 1 | 2 |
| MK0578 | 1 | 2 |
| MK0641 | 3 | 0 |
| MK0661 | 1 | 2 |
| MK0662 | 3 | 0 |
| MK0663 | 1 | 2 |
| MK0671 | 1 | 2 |
| MK0672 | 1 | 2 |
| MK0673 | 1 | 2 |
| MK0674 | 3 | 0 |
| MK0675 | 3 | 0 |
| MK0676 | 1 | 2 |
| MK0678 | 1 | 2 |
| MK0679 | 1 | 2 |
| MK0680 | 3 | 0 |
| MK0685 | 3 | 0 |
| MK0686 | 3 | 0 |
| MK0693 | 1 | 2 |
| MK0694 | 1 | 2 |
| MK0700 | 2 | 1 |
| MK0704 | 1 | 2 |
| MK0705 | 1 | 2 |
| MK0706 | 1 | 2 |
| MK0707 | 1 | 2 |
| MK0708 | 1 | 2 |
| MK0713 | 1 | 2 |
| MK0714 | 3 | 0 |
| MK0715 | 3 | 0 |
| MK0716 | 3 | 0 |
| MK0717 | 1 | 2 |
| MK0719 | 3 | 0 |
| MK0720 | 1 | 2 |
| MK0721 | 1 | 2 |
| MK0734 | 3 | 0 |
| MK0747 | 3 | 0 |
| MK0748 | 1 | 2 |
| MK0749 | 3 | 0 |
| MK0750 | 3 | 0 |
| MK0751 | 3 | 0 |
| MK0752 | 2 | 1 |
| MK0753 | 3 | 0 |
| MK0754 | 3 | 0 |
| MK0755 | 1 | 2 |
| MK0756 | 1 | 2 |
| MK0757 | 3 | 0 |
| MK0758 | 3 | 0 |
| MK0759 | 1 | 2 |
| MK0760 | 2 | 1 |
| MK0761 | 1 | 2 |
| MK0762 | 3 | 0 |
| MK0763 | 1 | 2 |
| MK0765 | 1 | 2 |
| MK0766 | 3 | 0 |
| MK0767 | 3 | 0 |
| MK0768 | 3 | 0 |
| MK0769 | 3 | 0 |
| MK0770 | 3 | 0 |
| MK0771 | 1 | 2 |
| MK0772 | 1 | 2 |
| MK0773 | 1 | 2 |
| MK0774 | 3 | 0 |
| MK0775 | 3 | 0 |
| MK0776 | 3 | 0 |
| MK0777 | 1 | 2 |
| MK0778 | 3 | 0 |
| MK0779 | 3 | 0 |
| MK0780 | 3 | 0 |
| MK0781 | 3 | 0 |
| MK0782 | 1 | 2 |
| MK0783 | 1 | 2 |
| MK0784 | 1 | 2 |
| MK0785 | 1 | 2 |
| MK0786 | 1 | 2 |
| MK0787 | 1 | 2 |
| MK0788 | 1 | 2 |
| MK0789 | 3 | 0 |
| MK0791 | 2 | 1 |
| MK0792 | 3 | 0 |
| MK0793 | 2 | 1 |
| MK0796 | 3 | 0 |
| MK0797 | 3 | 0 |
| MK0798 | 1 | 2 |
| MK0799 | 1 | 2 |
| MK0800 | 3 | 0 |
| MK0801 | 3 | 0 |
| MK0802 | 1 | 2 |
| MK0803 | 1 | 2 |
| MK0804 | 1 | 2 |
| MK0805 | 3 | 0 |
| MK0806 | 3 | 0 |
| MK0807 | 3 | 0 |
| MK0808 | 3 | 0 |
| MK0809 | 3 | 0 |
| MK0810 | 3 | 0 |
| MK0811 | 1 | 2 |
| MK0812 | 1 | 2 |
| MK0813 | 1 | 2 |
| MK0815 | 3 | 0 |
| MK0816 | 2 | 1 |
| MK0817 | 3 | 0 |
| MK0818 | 1 | 2 |
| MK0819 | 1 | 2 |
| MK0820 | 1 | 2 |
| MK0821 | 1 | 2 |
| MK0822 | 1 | 2 |
| MK0823 | 3 | 0 |
| MK0824 | 1 | 2 |
| MK0825 | 2 | 1 |
| MK0826 | 2 | 1 |
| MK0827 | 2 | 1 |
| MK0828 | 1 | 2 |
| MK0829 | 1 | 2 |
| MK0830 | 1 | 2 |
| MK0831 | 1 | 2 |
| MK0832 | 1 | 2 |
| MK0833 | 1 | 2 |
| MK0834 | 1 | 2 |
| MK0835 | 3 | 0 |
| MK0836 | 1 | 2 |
| MK0837 | 1 | 2 |
| MK0838 | 1 | 2 |
| MK0839 | 3 | 0 |
| MK0840 | 3 | 0 |
| MK0841 | 2 | 1 |
| MK0842 | 3 | 0 |
| MK0843 | 1 | 2 |
| MK0844 | 1 | 2 |
| MK0845 | 3 | 0 |
| MK0846 | 1 | 2 |
| MK0847 | 1 | 2 |
| MK0849 | 2 | 1 |
| MK0850 | 2 | 1 |
| MK0851 | 3 | 0 |
| MK0852 | 2 | 1 |
| MK0853 | 1 | 2 |
| MK0854 | 1 | 2 |
| MK0855 | 3 | 0 |
| MK0856 | 1 | 2 |
| MK0857 | 1 | 2 |
| MK0858 | 1 | 2 |
| MK0859 | 3 | 0 |
| MK0860 | 1 | 2 |
| MK0861 | 3 | 0 |
| MK0862 | 3 | 0 |
| MK0863 | 2 | 1 |
| MK0864 | 2 | 1 |
| MK0865 | 3 | 0 |
| MK0866 | 3 | 0 |
| MK0867 | 1 | 2 |
| MK0868 | 1 | 2 |
| MK0869 | 3 | 0 |
| MK0870 | 3 | 0 |
| MK0871 | 3 | 0 |
| MK0872 | 2 | 1 |
| MK0873 | 2 | 1 |
| MK0874 | 3 | 0 |
| MK0875 | 3 | 0 |
| MK0876 | 3 | 0 |
| MK0877 | 1 | 2 |
| MK0878 | 2 | 1 |
| MK0879 | 1 | 2 |
| MK0880 | 1 | 2 |
| MK0881 | 1 | 2 |
| MK0882 | 3 | 0 |
| MK0883 | 1 | 2 |
| MK0884 | 1 | 2 |
| MK0885 | 3 | 0 |
| MK0886 | 3 | 0 |
| MK0887 | 3 | 0 |
| MK0888 | 1 | 2 |
| MK0889 | 1 | 2 |
| MK0890 | 1 | 2 |
| MK0891 | 3 | 0 |
| MK0892 | 1 | 2 |
| MK0893 | 3 | 0 |
| MK0894 | 3 | 0 |
| MK0895 | 1 | 2 |
| MK0896 | 1 | 2 |
| MK0897 | 3 | 0 |
| MK0898 | 1 | 2 |
| MK0900 | 1 | 2 |
| MK0901 | 1 | 2 |
| MK0902 | 3 | 0 |
| MK0903 | 2 | 1 |
| MK0904 | 2 | 1 |
| MK0905 | 1 | 2 |
| MK0906 | 1 | 2 |
| MK0908 | 1 | 2 |
| MK0911 | 1 | 2 |
| MK0912 | 1 | 2 |
| MK0914 | 1 | 2 |
| MK0915 | 1 | 2 |
| MK0916 | 1 | 2 |
| MK0919 | 3 | 0 |
| MK0920 | 1 | 2 |
| MK0921 | 1 | 2 |
| MK0922 | 3 | 0 |
| MK0923 | 3 | 0 |
| MK0924 | 3 | 0 |
| MK0929 | 3 | 0 |
| MK0930 | 3 | 0 |
| MK0931 | 3 | 0 |
| MK0932 | 3 | 0 |
| MK0933 | 1 | 2 |
| MK0934 | 2 | 1 |
| MK0935 | 3 | 0 |
| MK0936 | 3 | 0 |
| MK0937 | 1 | 2 |
| MK0938 | 3 | 0 |
| MK0939 | 3 | 0 |
| MK0940 | 2 | 1 |
| MK0941 | 3 | 0 |
| MK0942 | 1 | 2 |
| MK0943 | 1 | 2 |
| MK0944 | 3 | 0 |
| MK0945 | 3 | 0 |
| MK0946 | 3 | 0 |
| MK0947 | 1 | 2 |
| MK0948 | 1 | 2 |
| MK0949 | 1 | 2 |
| MK0952 | 3 | 0 |
| MK0953 | 1 | 2 |
| MK0954 | 1 | 2 |
| MK0955 | 1 | 2 |
| MK0956 | 1 | 2 |
| MK0957 | 3 | 0 |
| MK0958 | 3 | 0 |
| MK0959 | 2 | 1 |
| MK0960 | 3 | 0 |
| MK0961 | 3 | 0 |
| MK0962 | 2 | 1 |
| MK0963 | 1 | 2 |
| MK0965 | 3 | 0 |
| MK0966 | 2 | 1 |
| MK0967 | 1 | 2 |
| MK0968 | 2 | 1 |
| MK0969 | 1 | 2 |
| MK1002 | 3 | 0 |
| MK1003 | 3 | 0 |
| MK1004 | 3 | 0 |
| MK1005 | 2 | 1 |
| MK1006 | 3 | 0 |

| | | |
|---|---|---|
| MK1007 | 3 | 0 |
| MK1008 | 3 | 0 |
| MK1010 | 3 | 0 |
| MK1011 | 3 | 0 |
| MK1012 | 1 | 2 |
| MK1013 | 3 | 0 |
| MK1014 | 2 | 1 |
| MK1015 | 1 | 2 |
| MK1016 | 1 | 2 |
| MK1017 | 1 | 2 |
| MK1018 | 3 | 0 |
| MK1019 | 3 | 0 |
| MK1020 | 3 | 0 |
| MK1021 | 1 | 2 |
| MK1022 | 1 | 2 |
| MK1023 | 3 | 0 |
| MK1024 | 3 | 0 |
| MK1025 | 3 | 0 |
| MK1026 | 2 | 1 |
| MK1027 | 3 | 0 |
| MK1028 | 2 | 1 |
| MK1029 | 3 | 0 |
| MK1030 | 3 | 0 |
| MK1031 | 2 | 1 |
| MK1032 | 2 | 1 |
| MK1033 | 1 | 2 |
| MK1034 | 1 | 2 |
| MK1035 | 1 | 2 |
| MK1036 | 3 | 0 |
| MK1037 | 2 | 1 |
| MK1038 | 3 | 0 |
| MK1039 | 1 | 2 |
| MK1040 | 1 | 2 |
| MK1041 | 3 | 0 |
| MK1042 | 3 | 0 |
| MK1043 | 3 | 0 |
| MK1044 | 2 | 1 |
| MK1045 | 1 | 2 |
| MK1046 | 3 | 0 |
| MK1047 | 1 | 2 |
| MK1048 | 1 | 2 |
| MK1049 | 1 | 2 |
| MK1050 | 1 | 2 |
| MK1051 | 3 | 0 |
| MK1111 | 3 | 0 |
| MK1149 | 3 | 0 |
| MK1151 | 2 | 1 |
| MK1152 | 3 | 0 |
| MK1153 | 2 | 1 |
| MK1154 | 1 | 2 |
| MK1155 | 2 | 1 |
| MK1156 | 3 | 0 |
| MK1157 | 3 | 0 |
| MK1158 | 3 | 0 |
| MK1159 | 2 | 1 |
| MK1160 | 3 | 0 |
| MK1161 | 1 | 2 |
| MK1162 | 2 | 1 |
| MK1163 | 3 | 0 |
| MK1164 | 2 | 1 |
| MK1165 | 3 | 0 |
| MK1166 | 3 | 0 |
| MK1167 | 1 | 2 |
| MK1168 | 3 | 0 |
| MK1169 | 3 | 0 |
| MK1170 | 3 | 0 |
| MK1171 | 3 | 0 |
| MK1172 | 3 | 0 |
| MK1173 | 3 | 0 |
| MK1174 | 3 | 0 |
| MK1175 | 3 | 0 |
| MK1176 | 2 | 1 |
| MK1177 | 2 | 1 |
| MK1178 | 3 | 0 |
| MK1179 | 1 | 2 |
| MK1180 | 1 | 2 |
| MK1181 | 2 | 1 |
| MK1182 | 3 | 0 |
| MK1183 | 3 | 0 |
| MK1184 | 1 | 2 |
| MK1185 | 1 | 2 |
| MK1186 | 3 | 0 |
| MK1187 | 3 | 0 |
| MK1188 | 3 | 0 |
| MK1189 | 3 | 0 |
| MK1190 | 3 | 0 |
| MK1191 | 3 | 0 |
| MK1192 | 3 | 0 |
| MK1193 | 2 | 1 |
| MK1194 | 3 | 0 |
| MK1195 | 1 | 2 |
| MK1196 | 1 | 2 |
| MK1197 | 3 | 0 |
| MK1198 | 3 | 0 |
| MK1199 | 3 | 0 |
| MM0001 | 3 | 0 |
| MM0002 | 2 | 1 |
| MM0003 | 2 | 1 |
| MM0004 | 2 | 1 |
| MM0005 | 2 | 1 |
| MM0006 | 2 | 1 |
| MM0007 | 2 | 1 |
| MM0009 | 2 | 1 |
| MM0010 | 2 | 1 |
| MM0012 | 3 | 0 |
| MM0013 | 2 | 1 |
| MM0014 | 2 | 1 |
| MM0015 | 2 | 1 |
| MM0016 | 2 | 1 |
| MM0017 | 2 | 1 |
| MM0018 | 3 | 0 |
| MM0019 | 2 | 1 |
| MM0020 | 2 | 1 |
| MM0021 | 2 | 1 |
| MM0022 | 2 | 1 |
| MM0023 | 2 | 1 |
| MM0024 | 2 | 1 |
| MM0026 | 3 | 0 |
| MM0027 | 2 | 1 |
| MM0029 | 1 | 2 |
| MM0030 | 2 | 1 |
| MM0031 | 2 | 1 |
| MM0032 | 2 | 1 |
| MM0033 | 2 | 1 |
| MM0034 | 3 | 0 |
| MM0035 | 3 | 0 |
| MM0036 | 3 | 0 |
| MM0037 | 2 | 1 |
| MM0038 | 3 | 0 |
| MM0039 | 2 | 1 |
| MM0040 | 3 | 0 |
| MM0041 | 3 | 0 |
| MM0044 | 2 | 1 |
| MM0045 | 2 | 1 |
| MM0047 | 1 | 2 |
| MM0048 | 2 | 1 |
| MM0051 | 2 | 1 |
| MM0053 | 2 | 1 |
| MM0054 | 2 | 1 |
| MM0055 | 2 | 1 |
| MM0056 | 2 | 1 |
| MM0057 | 1 | 2 |
| MM0058 | 3 | 0 |
| MM0059 | 3 | 0 |
| MM0060 | 2 | 1 |
| MM0061 | 2 | 1 |
| MW0001 | 3 | 0 |
| MW0002 | 3 | 0 |
| MW0003 | 3 | 0 |
| MW0004 | 3 | 0 |
| MW0005 | 2 | 1 |
| MW0006 | 3 | 0 |
| MW0007 | 3 | 0 |
| MW0008 | 3 | 0 |
| MW0009 | 3 | 0 |
| MW0010 | 2 | 1 |
| MW0011 | 2 | 1 |
| MW0012 | 1 | 2 |
| MW0013 | 3 | 0 |
| MW0014 | 2 | 1 |
| MW0015 | 1 | 2 |
| MW0016 | 2 | 1 |
| MW0017 | 1 | 2 |
| MW0018 | 3 | 0 |
| MW0019 | 3 | 0 |
| MW0020 | 3 | 0 |
| MW0021 | 3 | 0 |
| MW0022 | 1 | 2 |
| MW0023 | 3 | 0 |
| MW0024 | 3 | 0 |
| MW0025 | 3 | 0 |
| MW0026 | 3 | 0 |
| MW0027 | 2 | 1 |
| MW0028 | 2 | 1 |
| MW0029 | 1 | 2 |
| MW0030 | 2 | 1 |
| MW0031 | 3 | 0 |
| MW0032 | 3 | 0 |
| MW0033 | 2 | 1 |
| MW0036 | 2 | 1 |
| MW0037 | 3 | 0 |
| MW0038 | 3 | 0 |
| MW0042 | 3 | 0 |
| MW0043 | 3 | 0 |
| TV0001 | 2 | 1 |
| TV0002 | 1 | 2 |
| TV0003 | 2 | 1 |
| TV0004 | 2 | 1 |
| TV0005 | 2 | 1 |
| TV0006 | 3 | 0 |
| TV0007 | 3 | 0 |
| TV0008 | 3 | 0 |
| TV0009 | 2 | 1 |
| TV0010 | 3 | 0 |
| TV0012 | 3 | 0 |
| TV0013 | 3 | 0 |
| TV0014 | 3 | 0 |
| TV0017 | 2 | 1 |
| TV0018 | 3 | 0 |
| TV0019 | 2 | 1 |
| TV0020 | 3 | 0 |
| TV0021 | 3 | 0 |
| TV0022 | 3 | 0 |
| TV0023 | 3 | 0 |
| TV0024 | 3 | 0 |
| TV0025 | 3 | 0 |
| TV0026 | 3 | 0 |
| TV0027 | 3 | 0 |
| TV0028 | 3 | 0 |
| TV0030 | 3 | 0 |
| TV0031 | 3 | 0 |
| TV0032 | 3 | 0 |
| TV0033 | 3 | 0 |
| TV0036 | 3 | 0 |
| TV0037 | 3 | 0 |
| TV0038 | 3 | 0 |
| TV0039 | 2 | 1 |
| TV0040 | 3 | 0 |
| TV0041 | 3 | 0 |
| TV0042 | 3 | 0 |
| TV0043 | 3 | 0 |
| TV0044 | 3 | 0 |
| TV0045 | 3 | 0 |
| TV0046 | 3 | 0 |
| TV0047 | 3 | 0 |
| TV0048 | 3 | 0 |
| TV0049 | 3 | 0 |
| TV0050 | 2 | 1 |
| TV0051 | 2 | 1 |
| TV0052 | 2 | 1 |
| TV0053 | 3 | 0 |
| TV0054 | 3 | 0 |
| TV0055 | 3 | 0 |
| TV0059 | 3 | 0 |
| TV0060 | 3 | 0 |
| TV0061 | 3 | 0 |
| TV0062 | 3 | 0 |
| TV0063 | 3 | 0 |
| TV0064 | 3 | 0 |
| TV0065 | 3 | 0 |
| TV0066 | 3 | 0 |
| TV0067 | 3 | 0 |
| TV0068 | 3 | 0 |
| TV0069 | 3 | 0 |
| TV0070 | 3 | 0 |
| TV0071 | 3 | 0 |
| TV0072 | 3 | 0 |
| TV0073 | 3 | 0 |
| TV0074 | 3 | 0 |
| TV0075 | 3 | 0 |
| TV0076 | 3 | 0 |
| TV0077 | 2 | 1 |
| TV0078 | 3 | 0 |
| TV0079 | 3 | 0 |
| TV0080 | 2 | 1 |
| TV0081 | 3 | 0 |
| TV0082 | 3 | 0 |
| TV0083 | 2 | 1 |
| TV0084 | 3 | 0 |
| TV0085 | 3 | 0 |
| TV0086 | 3 | 0 |
| TV0087 | 3 | 0 |
| TV0088 | 3 | 0 |
| TV0089 | 2 | 1 |
| TV0090 | 3 | 0 |
| TV0091 | 1 | 2 |
| TV0092 | 3 | 0 |
| TV0093 | 3 | 0 |
| TV0094 | 3 | 0 |
| TV0095 | 2 | 1 |
| TV0096 | 3 | 0 |
| VK0001 | 2 | 1 |
| VK0002 | 3 | 0 |
| VK0003 | 2 | 1 |
| VK0004 | 2 | 1 |
| VK0005 | 2 | 1 |
| VK0006 | 3 | 0 |
| VK0007 | 2 | 1 |
| VK0008 | 2 | 1 |
| VK0009 | 3 | 0 |
| VK0010 | 3 | 0 |
| VK0011 | 2 | 1 |
| VK0012 | 3 | 0 |
| VK0013 | 2 | 1 |
| VK0014 | 3 | 0 |
| VK0015 | 2 | 1 |
| VK0016 | 3 | 0 |
| VK0017 | 2 | 1 |
| VK0018 | 2 | 1 |
| VK0019 | 2 | 1 |
| VK0028 | 2 | 1 |
| VK0029 | 2 | 1 |
| VK0038 | 2 | 1 |
| VK0039 | 2 | 1 |
| VK0040 | 3 | 0 |
| VK0041 | 2 | 1 |
| VK0042 | 2 | 1 |
| VK0043 | 2 | 1 |
| VK0044 | 2 | 1 |
| VK0045 | 3 | 0 |
| VK0046 | 2 | 1 |
| VK0047 | 2 | 1 |
| VK0048 | 2 | 1 |
| VK0049 | 2 | 1 |
| VK0050 | 2 | 1 |
| VK0051 | 2 | 1 |
| VK0054 | 2 | 1 |
| VK0056 | 2 | 1 |
| VK0059 | 2 | 1 |
| VK0060 | 2 | 1 |
| VK0061 | 2 | 1 |
| VK0062 | 3 | 0 |
| VK0063 | 3 | 0 |
| VK0064 | 2 | 1 |
| VK0065 | 2 | 1 |
| VK0066 | 2 | 1 |
| VK0067 | 3 | 0 |
| VK0068 | 2 | 1 |
| VK0069 | 2 | 1 |
| VK0070 | 3 | 0 |
| VK0071 | 2 | 1 |
| VK0072 | 2 | 1 |
| VK0074 | 2 | 1 |
| VK0075 | 3 | 0 |
| VK0077 | 3 | 0 |
| VK0078 | 2 | 1 |
| VK0080 | 2 | 1 |
| VK0081 | 3 | 0 |
| VK0082 | 3 | 0 |
| VK0083 | 3 | 0 |
| VK0084 | 3 | 0 |
| VK0085 | 2 | 1 |
| VK0087 | 3 | 0 |
| VK0088 | 2 | 1 |
| VK0089 | 2 | 1 |
| VK0090 | 2 | 1 |
| VK0091 | 2 | 1 |
| VK0093 | 2 | 1 |
| VK0094 | 3 | 0 |
| VK0095 | 2 | 1 |
| VK0098 | 3 | 0 |
| VK0099 | 2 | 1 |
| VK0100 | 3 | 0 |
| VK0116 | 2 | 1 |
| VK0117 | 2 | 1 |
| VK0126 | 2 | 1 |
| VK0127 | 2 | 1 |
| VK0128 | 3 | 0 |
| VK0129 | 2 | 1 |
| VK0130 | 3 | 0 |
| VK0131 | 2 | 1 |
| VK0132 | 3 | 0 |
| VK0133 | 3 | 0 |
| VK0134 | 2 | 1 |
| VK0135 | 3 | 0 |
| VK0136 | 2 | 1 |
| VK0137 | 2 | 1 |
| VK0138 | 3 | 0 |
| VK0141 | 2 | 1 |
| VK0142 | 2 | 1 |
| VK0143 | 2 | 1 |
| VK0144 | 2 | 1 |
| VK0145 | 2 | 1 |
| VK0146 | 3 | 0 |
| VK0147 | 2 | 1 |
| VK0148 | 2 | 1 |
| VK0149 | 1 | 2 |
| VK0150 | 2 | 1 |
| VK0151 | 2 | 1 |
| VK0152 | 3 | 0 |
| VK0153 | 3 | 0 |
| VK0154 | 2 | 1 |
| VK0155 | 3 | 0 |
| VK0156 | 2 | 1 |
| VK0157 | 3 | 0 |
| VK0158 | 3 | 0 |
| VK0159 | 3 | 0 |
| VK0160 | 3 | 0 |
| VK0161 | 2 | 1 |
| VK0162 | 2 | 1 |
| VK0163 | 3 | 0 |
| VK0164 | 2 | 1 |
| VK0165 | 2 | 1 |
| VK0166 | 3 | 0 |
| VK0167 | 2 | 1 |
| VK0168 | 2 | 1 |
| VK0169 | 2 | 1 |
| VK0170 | 2 | 1 |
| VK0171 | 2 | 1 |
| VK0172 | 2 | 1 |
| VK0173 | 3 | 0 |
| VK0174 | 2 | 1 |
| VK0175 | 3 | 0 |
| VK0176 | 2 | 1 |
| VK0177 | 3 | 0 |
| VK0178 | 2 | 1 |
| VK0179 | 3 | 0 |
| VK0180 | 2 | 1 |
| VK0181 | 1 | 2 |
| VK0182 | 2 | 1 |
| VK0183 | 2 | 1 |
| VK0184 | 3 | 0 |
| VK0185 | 3 | 0 |
| VK0186 | 2 | 1 |
| VK0187 | 3 | 0 |
| VK0188 | 2 | 1 |
| VK0189 | 2 | 1 |
| VK0190 | 3 | 0 |
| VK0191 | 2 | 1 |
| VK0192 | 2 | 1 |
| VK0193 | 2 | 1 |
| VK0194 | 2 | 1 |
| VK0195 | 2 | 1 |
| VK0196 | 3 | 0 |
| VK0197 | 3 | 0 |
| VK0198 | 2 | 1 |
| VK0199 | 2 | 1 |
| VK0200 | 3 | 0 |
| VK0201 | 2 | 1 |
| VK0202 | 2 | 1 |
| VK0203 | 3 | 0 |
| VK0204 | 2 | 1 |
| VK0205 | 2 | 1 |
| VK0206 | 3 | 0 |
| VK0207 | 3 | 0 |
| VK0208 | 2 | 1 |
| VK0209 | 2 | 1 |
| VK0210 | 1 | 2 |
| VK0211 | 2 | 1 |
| VK0212 | 3 | 0 |
| VK0213 | 2 | 1 |
| VK0214 | 2 | 1 |
| VK0215 | 3 | 0 |
| VK0216 | 2 | 1 |
| VK0218 | 2 | 1 |
| VK0219 | 3 | 0 |
| VK0220 | 1 | 2 |
| VK0221 | 1 | 2 |
| VK0222 | 3 | 0 |
| VK0223 | 2 | 1 |
| VK0224 | 2 | 1 |
| VK0225 | 1 | 2 |
| VK0226 | 3 | 0 |
| VK0227 | 2 | 1 |
| VK0228 | 2 | 1 |
| VK0229 | 3 | 0 |
| VK0230 | 2 | 1 |
| VK0231 | 3 | 0 |
| VK0232 | 2 | 1 |
| VK0233 | 3 | 0 |
| VK0235 | 2 | 1 |
| VK0236 | 2 | 1 |
| VK0237 | 2 | 1 |
| VK0239 | 2 | 1 |
| VK0240 | 2 | 1 |
| VK0241 | 2 | 1 |
| VK0242 | 2 | 1 |
| VK0243 | 2 | 1 |
| VK0244 | 2 | 1 |
| VK0245 | 2 | 1 |
| VK0246 | 2 | 1 |
| VK0247 | 3 | 0 |
| VK0248 | 2 | 1 |
| VK0249 | 3 | 0 |
| VK0250 | 1 | 2 |
| VK0251 | 2 | 1 |
| VK0252 | 2 | 1 |
| VK0253 | 3 | 0 |
| VK0254 | 3 | 0 |
| VK0255 | 2 | 1 |
| VK0256 | 3 | 0 |
| VK0257 | 2 | 1 |
| VK0258 | 2 | 1 |
| VK0259 | 2 | 1 |
| VK0260 | 3 | 0 |
| VK0261 | 3 | 0 |
| VK0262 | 2 | 1 |
| VK0263 | 2 | 1 |
| VK0264 | 2 | 1 |
| VK0265 | 2 | 1 |
| VK0266 | 2 | 1 |
| VK0267 | 2 | 1 |
| VK0268 | 2 | 1 |
| VK0269 | 2 | 1 |
| VK0270 | 3 | 0 |
| VK0271 | 2 | 1 |
| VK0272 | 2 | 1 |
| VK0273 | 2 | 1 |
| VK0274 | 2 | 1 |
| VK0275 | 2 | 1 |
| VK0276 | 2 | 1 |
| VK0277 | 2 | 1 |
| VK0278 | 2 | 1 |
| VK0280 | 3 | 0 |
| VK0281 | 2 | 1 |
| VK0283 | 2 | 1 |
| VK0284 | 2 | 1 |
| VK0285 | 2 | 1 |
| VK0286 | 1 | 2 |
| VK0289 | 2 | 1 |
| VK0303 | 2 | 1 |
| VK0307 | 2 | 1 |
| VK0308 | 3 | 0 |
| VK0309 | 3 | 0 |
| VK0310 | 2 | 1 |
| VK0311 | 2 | 1 |
| VK0312 | 2 | 1 |
| VK0313 | 2 | 1 |
| VK0315 | 2 | 1 |
| VK0317 | 2 | 1 |
| VK0318 | 2 | 1 |
| VK0319 | 2 | 1 |
| VK0320 | 3 | 0 |
| VK0321 | 2 | 1 |
| VK0322 | 2 | 1 |
| VK0323 | 2 | 1 |
| VK0324 | 2 | 1 |
| VK0325 | 2 | 1 |
| VK0326 | 3 | 0 |
| VK0327 | 2 | 1 |
| VK0328 | 1 | 2 |
| VK0329 | 2 | 1 |
| VK0331 | 2 | 1 |
| VK0332 | 1 | 2 |
| VK0334 | 2 | 1 |
| VK0335 | 2 | 1 |
| VK0336 | 2 | 1 |
| VK0337 | 3 | 0 |
| VK0338 | 3 | 0 |
| VK0340 | 3 | 0 |
| VK0341 | 2 | 1 |
| VK0342 | 2 | 1 |
| VK0343 | 2 | 1 |
| VK0345 | 2 | 1 |
| VK0349 | 2 | 1 |
| VK0352 | 2 | 1 |
| VK0353 | 2 | 1 |
| VK0354 | 2 | 1 |
| VK0355 | 2 | 1 |
| VK0356 | 2 | 1 |
| VK0357 | 1 | 2 |
| VK0358 | 2 | 1 |
| VK0359 | 3 | 0 |
| VK0360 | 2 | 1 |
| VK0361 | 2 | 1 |
| VK0362 | 2 | 1 |
| VK0363 | 2 | 1 |
| VK0364 | 3 | 0 |
| VK0365 | 2 | 1 |
| VK0366 | 2 | 1 |
| VK0367 | 2 | 1 |
| VK0368 | 2 | 1 |
| VK0369 | 2 | 1 |
| VK0370 | 2 | 1 |
| VK0371 | 2 | 1 |
| VK0372 | 2 | 1 |
| VK0373 | 3 | 0 |
| VK0378 | 1 | 2 |
| VK0379 | 1 | 2 |
| VK0382 | 2 | 1 |
| VK0384 | 2 | 1 |
| VK0386 | 2 | 1 |
| VK0387 | 3 | 0 |
| VK0388 | 2 | 1 |
| VK0389 | 1 | 2 |
| VK0390 | 1 | 2 |
| VK0391 | 2 | 1 |
| VK0392 | 3 | 0 |
| VK0393 | 2 | 1 |
| VK0395 | 1 | 2 |
| VK0397 | 2 | 1 |
| VK0398 | 2 | 1 |
| VK0399 | 2 | 1 |
| VK0400 | 2 | 1 |
| VK0401 | 3 | 0 |
| VK0402 | 2 | 1 |
| VK0403 | 3 | 0 |
| VK0404 | 2 | 1 |
| VK0405 | 2 | 1 |
| VK0406 | 3 | 0 |
| VK0407 | 2 | 1 |
| VK0408 | 3 | 0 |
| VK0409 | 2 | 1 |
| VK0410 | 2 | 1 |
| VK0411 | 2 | 1 |
| VK0412 | 3 | 0 |
| VK0413 | 2 | 1 |
| VK0414 | 2 | 1 |
| VK0415 | 3 | 0 |
| VK0416 | 2 | 1 |
| VK0417 | 2 | 1 |
| VK0418 | 2 | 1 |
| VK0419 | 3 | 0 |
| VK0420 | 3 | 0 |
| VK0421 | 2 | 1 |
| VK0422 | 2 | 1 |
| VK0423 | 2 | 1 |
| VK0424 | 3 | 0 |
| VK0425 | 2 | 1 |
| VK0426 | 3 | 0 |
| VK0427 | 3 | 0 |
| VK0428 | 2 | 1 |
| VK0430 | 3 | 0 |
| VK0432 | 1 | 2 |
| VK0433 | 2 | 1 |
| VK0434 | 2 | 1 |
| VK0435 | 2 | 1 |
| VK0437 | 2 | 1 |
| VK0438 | 3 | 0 |
| VK0439 | 3 | 0 |
| VK0440 | 2 | 1 |
| VK0441 | 3 | 0 |
| VK0442 | 2 | 1 |
| VK0443 | 2 | 1 |
| VK0444 | 2 | 1 |
| VK0445 | 3 | 0 |
| VK0446 | 2 | 1 |
| VK0447 | 2 | 1 |
| VK0449 | 2 | 1 |
| VK0450 | 2 | 1 |
| VK0451 | 3 | 0 |
| VK0452 | 2 | 1 |
| VK0453 | 2 | 1 |
| VK0454 | 3 | 0 |
| VK0455 | 2 | 1 |
| VK0456 | 2 | 1 |
| VK0457 | 2 | 1 |
| VK0459 | 2 | 1 |
| VK0460 | 2 | 1 |
| VK0463 | 2 | 1 |
| VK0465 | 3 | 0 |
| VK0467 | 2 | 1 |
| VK0468 | 3 | 0 |
| VK0469 | 2 | 1 |
| VK0470 | 1 | 2 |
| VK0471 | 3 | 0 |
| VK0472 | 3 | 0 |
| VK0476 | 3 | 0 |
| VK0477 | 2 | 1 |
| VK0478 | 2 | 1 |
| VK0480 | 2 | 1 |
| VK0481 | 3 | 0 |
| VK0483 | 2 | 1 |
| VK0484 | 2 | 1 |
| VK0485 | 3 | 0 |
| VK0486 | 3 | 0 |
| VK0487 | 2 | 1 |
| VK0488 | 2 | 1 |
| VK0489 | 2 | 1 |
| VK0490 | 2 | 1 |
| VK0491 | 3 | 0 |
| VK0492 | 2 | 1 |
| VK0493 | 2 | 1 |
| VK0494 | 2 | 1 |
| VK0495 | 2 | 1 |
| VK0496 | 2 | 1 |
| VK0497 | 2 | 1 |
| VK0498 | 2 | 1 |
| VK0499 | 2 | 1 |
| VK0500 | 2 | 1 |
| VK0501 | 3 | 0 |
| VK0502 | 2 | 1 |
| VK0503 | 1 | 2 |
| VK0504 | 3 | 0 |
| VK0505 | 1 | 2 |
| VK0506 | 2 | 1 |
| VK0507 | 1 | 2 |
| VK0508 | 1 | 2 |
| VK0510 | 2 | 1 |
| VK0512 | 2 | 1 |
| VK0514 | 2 | 1 |
| VK0515 | 2 | 1 |
| VK0516 | 2 | 1 |
| VK0517 | 2 | 1 |
| VK0518 | 2 | 1 |
| VK0519 | 3 | 0 |
| VK0520 | 2 | 1 |
| VK0521 | 2 | 1 |
| VK0523 | 2 | 1 |
| VK0524 | 2 | 1 |
| VK0525 | 2 | 1 |
| VK0526 | 2 | 1 |
| VK0527 | 3 | 0 |
| VK0528 | 2 | 1 |
| VK0529 | 3 | 0 |
| VK0530 | 3 | 0 |
| VK0531 | 3 | 0 |
| VK0533 | 3 | 0 |
| VK0534 | 2 | 1 |
| VK0535 | 3 | 0 |
| VK0537 | 2 | 1 |
| VK0539 | 2 | 1 |
| VK0540 | 2 | 1 |
| VK0541 | 2 | 1 |
| VK0543 | 2 | 1 |
| VK0544 | 3 | 0 |
| VK0546 | 3 | 0 |
| VK0547 | 2 | 1 |
| VK0548 | 2 | 1 |
| VK0550 | 2 | 1 |
| VK0551 | 3 | 0 |
| VK0552 | 2 | 1 |
| VK0553 | 2 | 1 |
| VK0554 | 2 | 1 |
| VK0555 | 2 | 1 |
| VK0556 | 1 | 2 |
| VK0557 | 3 | 0 |
| VK0558 | 3 | 0 |
| VK0559 | 2 | 1 |
| VK0560 | 2 | 1 |
| VK0561 | 3 | 0 |
| VK0563 | 2 | 1 |
| VK0564 | 2 | 1 |
| VK0565 | 2 | 1 |
| VK0566 | 2 | 1 |
| VK0567 | 2 | 1 |
| VK0568 | 2 | 1 |
| VK0569 | 2 | 1 |
| VK0570 | 3 | 0 |
| VK0571 | 2 | 1 |
| VK0572 | 2 | 1 |
| VK0573 | 2 | 1 |
| VK0574 | 2 | 1 |
| VK0575 | 2 | 1 |
| VK0576 | 2 | 1 |
| VK0577 | 2 | 1 |
| VK0578 | 3 | 0 |
| VK0579 | 2 | 1 |
| VK0580 | 2 | 1 |
| VK0581 | 1 | 2 |
| VK0582 | 3 | 0 |
| VK0583 | 2 | 1 |
| VK0586 | 2 | 1 |
| VK0595 | 2 | 1 |
| VK0596 | 2 | 1 |
| VK0597 | 2 | 1 |
| VK0598 | 2 | 1 |
| VK0599 | 2 | 1 |
| VK0603 | 3 | 0 |
| VK0604 | 2 | 1 |
| VK0605 | 2 | 1 |
| VK0606 | 2 | 1 |
| VK0607 | 3 | 0 |
| VK0608 | 2 | 1 |
| VK0609 | 3 | 0 |
| VK0610 | 2 | 1 |
| VK0611 | 3 | 0 |
| VK0612 | 3 | 0 |
| VK0613 | 2 | 1 |
| VK0614 | 3 | 0 |
| VK0615 | 3 | 0 |
| VK0616 | 2 | 1 |
| VK0618 | 2 | 1 |
| VK0619 | 2 | 1 |
| VK0620 | 2 | 1 |
| VK0621 | 3 | 0 |
| VK0622 | 2 | 1 |
| VK0623 | 2 | 1 |
| VK0625 | 2 | 1 |
| VK0626 | 2 | 1 |
| VK0627 | 2 | 1 |
| VK0628 | 3 | 0 |
| VK0634 | 2 | 1 |
| VK0635 | 2 | 1 |
| VK0636 | 2 | 1 |
| VK0637 | 2 | 1 |
| VK0638 | 3 | 0 |
| VK0640 | 2 | 1 |
| VK0641 | 2 | 1 |
| VK0642 | 3 | 0 |
| VK0644 | 3 | 0 |
| VK0645 | 2 | 1 |
| VK0646 | 3 | 0 |
| VK0647 | 3 | 0 |
| VK0650 | 2 | 1 |
| VK0651 | 1 | 2 |
| VK0652 | 2 | 1 |
| VK0654 | 2 | 1 |
| VK0655 | 3 | 0 |
| VK0656 | 2 | 1 |
| VK0657 | 3 | 0 |
| VK0660 | 2 | 1 |
| VK0663 | 2 | 1 |
| VK0664 | 2 | 1 |
| VK0665 | 1 | 2 |
| VK0668 | 2 | 1 |
| VK0670 | 3 | 0 |
| VK0672 | 2 | 1 |
| VK0673 | 2 | 1 |
| VK0674 | 3 | 0 |
| VK0677 | 2 | 1 |
| VK0678 | 2 | 1 |
| VK0680 | 1 | 2 |
| VK0681 | 3 | 0 |
| VK0682 | 1 | 2 |
| VK0683 | 3 | 0 |
| VK0684 | 2 | 1 |
| VK0685 | 2 | 1 |
| VK0686 | 2 | 1 |
| VK0687 | 2 | 1 |
| VK0688 | 2 | 1 |
| VK0689 | 3 | 0 |
| VK0690 | 2 | 1 |
| VK0692 | 3 | 0 |
| VK0693 | 2 | 1 |
| VK0694 | 2 | 1 |
| VK0695 | 2 | 1 |
| VK0696 | 2 | 1 |
| VK0697 | 3 | 0 |
| VK0698 | 2 | 1 |
| VK0699 | 2 | 1 |
| VK0700 | 2 | 1 |
| VK0701 | 2 | 1 |
| VK0702 | 2 | 1 |
| VK0703 | 2 | 1 |
| VK0704 | 3 | 0 |
| VK0705 | 1 | 2 |
| VK0706 | 3 | 0 |
| VK0707 | 2 | 1 |
| VK0708 | 2 | 1 |
| VK0709 | 2 | 1 |
| VK0710 | 2 | 1 |
| VK0711 | 2 | 1 |
| VK0712 | 2 | 1 |
| VK0713 | 3 | 0 |
| VK0715 | 2 | 1 |
| VK0716 | 2 | 1 |
| VK0717 | 3 | 0 |
| VK0718 | 3 | 0 |
| VK0719 | 2 | 1 |
| VK0721 | 2 | 1 |
| VK0722 | 2 | 1 |
| VK0723 | 2 | 1 |
| VK0724 | 2 | 1 |
| VK0725 | 1 | 2 |
| VK0726 | 3 | 0 |
| VK0727 | 2 | 1 |
| VK0728 | 1 | 2 |
| VK0729 | 2 | 1 |
| VK0735 | 3 | 0 |
| VK0736 | 3 | 0 |
| VK0737 | 3 | 0 |
| VK0738 | 3 | 0 |
| VK0739 | 3 | 0 |
| VK0740 | 3 | 0 |
| VK0741 | 2 | 1 |
| VK0742 | 2 | 1 |
| VK0743 | 3 | 0 |
| VK0744 | 3 | 0 |
| VK0745 | 3 | 0 |
| VK0746 | 3 | 0 |
| VK0747 | 3 | 0 |
| VK0748 | 3 | 0 |

Table S4. Overview of official gene sets of six reference species used for ortholog assessment. Number of genes correspond with OrthoDB 9.1.

| Species | Accession | # of contigs | Source | Download Date | Reference |
|---|---|---|---|---|---|
| *Onthophagus taurus* | PRJNA167478 | 17483 | i5K | 05.03.2017 | 1 |
| *Tribolium castaneum* | PRJNA12540 | 16631 | iBeetle | 05.03.2017 | 2,3 |
| *Dendroctonus ponderosae* | PRJNA360270 | 13088 | ENS Metazoa | 05.03.2017 | 4 |
| *Anoploph ora glabripennis* | PRJNA167479 | 22035 | i5K | 05.03.2017 | 5 |
| *Leptinotarsa decemlin*eata | PRJNA171749 | 24671 | i5K | 05.03.2017 | 1 |
| *Agrilus planipennis* | PRJNA230921 | 15497 | i5K | 05.03.2017 | 1 |

Supplementary Table S5. Detailed information and statistics of each generated dataset.

| Dataset Name | Number of taxa | Number of partitions | Number of alignment sites | Completeness score (Ca) AliStat | Percentage of pairwise p-values < 0.05 for the Bowker's test | (SV) MARE Matrix saturation | (IC) MARE Information content | Partition scheme optimalization |
|---|---|---|---|---|---|---|---|---|
| amino acids | | | | | | | | |
| A-4109-AA | 42 | 4109 | 1892691 | 0.772127 | 98.61% | 0.888 | 0.506 | Yes |
| D-3370-AA_Bacoca | 42 | 3370 | 1672362 | 0.780363 | 97.91% | 0.880 | 0.511 | |
| F-1490-AA_Bacoca_decisive | 42 | 1490 | 673102 | 0.925082 | 94.08% | 1 | 0.594 | |
| J-2129-AA_MARE | 42 | 2129 | 959741 | 0.876331 | 90.82% | 0.964 | 0.648 | |
| nucleoides | | | | | | | | |
| B-4109-NT | 42 | 4109 | 5678073 | 0.772133 | 100% | NA | NA | |
| C-4109-NT12 | 42 | 4109 | 3785382 | 0.772133 | 99.77% | NA | NA | |
| E-4109-NT2 | 42 | 4109 | 1892691 | 0.772134 | 92.45% | NA | NA | |
| G-NT-1767_MaxSymTest | 42 | 1767 | 2413164 | 0.713912 | 99.88% | NA | NA | |
| H-NT-1645_MaxSymTest$_{marginal}$ | 42 | 1645 | 2233485 | 0.712208 | 99.77% | NA | NA | |
| I-NT-3905_MaxSymTest$_{Internal}$ | 42 | 3905 | 5377449 | 0.771645 | 100% | NA | NA | |

Table S6. Characteristics of concatenated super-matrices (mitochondrial fragments) and used models of the DNA evolution (descriptions of columns abbreviation listed bellow tables).

Complete dataset

| Subset | Seqs | Sites | Infor | Invar | Model |
|---|---|---|---|---|---|
| *nad5* | 3205 | 987 | 809 | 88 | GTR+F+I+G4 |
| *cox* | 5935 | 1068 | 812 | 134 | GTR+F+I+G4 |
| *16s* | 2381 | 875 | 577 | 201 | GTR+F+I+G4 |

98% OTUs delimitation dataset

| Subset | Seqs | Sites | Infor | Invar | Model |
|---|---|---|---|---|---|
| *nad5* | 1252 | 969 | 776 | 115 | GTR+F+I+G4 |
| tRNA *Leu* | 2245 | 62 | 34 | 17 | GTR+F+G4 |
| *cox1* | 2395 | 783 | 536 | 141 | GTR+F+I+G4 |
| *cox2* | 2192 | 216 | 194 | 11 | GTR+F+I+G4 |
| *16s* | 1115 | 834 | 512 | 237 | GTR+F+I+G4 |

Abbreviations:
Seq: Number of sequences
Site: Number of bases
Infor: Number of parsimony-informative sites
Invar: Number of invariant sites
Model: Best-fit model according to BIC using ModelFinder.

Table S8. Overview of the revised Metriorrhynchini generic classification and distribution. Abbreviations: AFR – Sub-Saharan Africa; AUS – Australia, JAP – Japan; LSU – Lesser Sundas; MDG – Madagascar; MOL – Moluccas; NG – New Guinea; ORI – Oriental region; PAL – Eastern Palearctic region; SUL – Sulawesi; PHI – Philippines; SRL – Sri Lanka; WAL – Wallacea; [e] – endemic.

**Subtribe Metanoeina Kubecek *et al.*, 2015**

*Matsudanoeus* Sklenarova *et al.*, 2014 (JAP)
*Metanoeus* Waterhouse, 1878 (ORI)
*Ochinoeus* Kubecek *et al*., 2015 (JAP)
*Xylometanoeus* Sklenarova *et al.*, 2014 (ORI, PAL)

**Subtribe Cautirina Sklenarova *et al.*, 2015**

*Caenioxylobanus* Waterhouse, 1878 (MDG)
*Cautires* Waterhouse, 1878 (SUL, ORI, AFR, MDG, PAL)
*Prometanoeus* Kleine, 1925 (SRL)[e]
*Paracautires* Kazantsev, 2012 (AFR)[e]
*Spartoires* Kazantsev, 2012 (AFR)[e]
*Tricautires* Kazantsev, 2006 (AFR)[e]
*Xylobanus* Waterhouse, 1878 (SUL, ORI, PAL)

**Subtribe Metriorrhynchina Kleine, 1926a**

*Procautirine clade*
*Procautires* Kleine, 1925b (AUS, NG, MOL)
*Xylothrix* Kazantsev, 2015 (NG)[e]
*Leptotrichaline clade*
*Falsolucidota* Pic, 1921 (NG)[e]
*Broxylus* Waterhouse, 1879 (SUL)[e]
*Leptotrichalus* Kleine, 1925 (WAL, ORI)
*Mangkutanus* Kubecek *et al*., 2011 (SUL)[e]
*Sulabanus* Dvorak & Bocak, 2007 (SUL)[e]
*Synchonnus* Waterhouse, 1879 (AUS, NG)
*Wakarumbia* Bocak, 1999 (SUL)[e]
*Cladophorine clade*
*Cautiromimus* Pic, 1926 (NG, MOL)
*Cladophorinus* Kleine, 1926 (NG)[e]
*Cladophorus* Guérin-Méneville, 1830 (NG)[e]
*Ditua* Waterhouse, 1879 (NG, MOL, SUL, PHI)
*Marena* Kazantsev, 2007 (NG)[e]
*Pseudodontocerus* Pic, 1921 (NG)[e]
*Trichaline clade*
*Diatrichalus* Kleine, 1926 (AUS, NG, WAL, ORI)
*Eniclases* Waterhouse, 1879 (NG, MOL)
*Flabellotrichalus* Pic, 1921 (NG, MOL)
*Lobatang* Bocak, 1998 (NG, WAL)
*Microtrichalus* Pic, 1921 (AUS, NG, WAL, ORI)
*Schizotrichalus* Kleine, 1926 (NG, MOL)
*Trichalus* Waterhouse, 1877 (AUS, NG, WAL, ORI)
*Porrostomine clade*
*Metriorrhynchus* Gemminger & Harold, 1869 (AUS, NG, WAL, ORI)
*Porrostoma* Laporte, 1838 (AUS, NG, MOL)
*Oriomum* Bocak, 1999 (NG)[e]
*Metriorrhynchoides* Kleine, 1926 (NG)[e]
*Stadenus* Waterhouse, 1879 (AUS)[e]
Metriorrhynchina *incertae sedis*
*Malacolycus* Kleine, 1943 (NG, LSU)
*Xylobanomimus* Kleine, 1926 (NG)[e]
*Xylobanomorphus* Kleine, 1935 (NG)[e]
*Kassemia Bocak, 1998a (NG)*[e]

Table S7. Comparison of the Robinson-Fould distances among tree searches with constrained topology and tree searches with unconstrained topology. Green values represent top 10% RF- distances, and red values shows bottom 10% RF- distances. The blue part of the graph represents constrained trees values, the grey part of the graph represents unconstrained trees values, red part of the graph represents constrained/unconstrained trees values.

| | | 1 | 2 | 3 | 4 | 5 | 6 | 7 | 8 | 9 | 10 | 11 | 12 | 13 | 14 | 15 | 16 | 17 | 18 | 19 | 20 | 21 | 22 | 23 | 24 | 25 | 26 | 27 | 28 | 29 | 30 | 31 | 32 | 33 | 34 | 35 | 36 | 37 | 38 |
|---|---|---|---|---|---|---|---|---|---|---|---|---|---|---|---|---|---|---|---|---|---|---|---|---|---|---|---|---|---|---|---|---|---|---|---|---|---|---|---|
| uncon | 1 | - | | | | | | | | | | | | | | | | | | | | | | | | | | | | | | | | | | | | | |
| uncon | 2 | 0.139 | - | | | | | | | | | | | | | | | | | | | | | | | | | | | | | | | | | | | | |
| uncon | 3 | 0.141 | 0.150 | - | | | | | | | | | | | | | | | | | | | | | | | | | | | | | | | | | | | |
| uncon | 4 | 0.149 | 0.141 | 0.158 | - | | | | | | | | | | | | | | | | | | | | | | | | | | | | | | | | | | |
| uncon | 5 | 0.150 | 0.142 | 0.153 | 0.152 | - | | | | | | | | | | | | | | | | | | | | | | | | | | | | | | | | | |
| uncon | 6 | 0.150 | 0.153 | 0.158 | 0.170 | 0.166 | - | | | | | | | | | | | | | | | | | | | | | | | | | | | | | | | | |
| uncon | 7 | 0.153 | 0.165 | 0.163 | 0.174 | 0.168 | 0.171 | - | | | | | | | | | | | | | | | | | | | | | | | | | | | | | | | |
| uncon | 8 | 0.151 | 0.153 | 0.154 | 0.152 | 0.144 | 0.158 | 0.152 | - | | | | | | | | | | | | | | | | | | | | | | | | | | | | | | |
| uncon | 9 | 0.145 | 0.143 | 0.143 | 0.163 | 0.145 | 0.156 | 0.156 | 0.156 | - | | | | | | | | | | | | | | | | | | | | | | | | | | | | | |
| uncon | 10 | 0.143 | 0.151 | 0.149 | 0.161 | 0.139 | 0.161 | 0.161 | 0.162 | 0.141 | - | | | | | | | | | | | | | | | | | | | | | | | | | | | | |
| uncon | 11 | 0.150 | 0.151 | 0.152 | 0.148 | 0.149 | 0.166 | 0.171 | 0.153 | 0.154 | 0.150 | - | | | | | | | | | | | | | | | | | | | | | | | | | | | |
| uncon | 12 | 0.150 | 0.159 | 0.160 | 0.157 | 0.145 | 0.161 | 0.173 | 0.149 | 0.152 | 0.154 | 0.159 | - | | | | | | | | | | | | | | | | | | | | | | | | | | |
| uncon | 13 | 0.143 | 0.141 | 0.153 | 0.153 | 0.157 | 0.156 | 0.166 | 0.143 | 0.151 | 0.157 | 0.153 | 0.158 | - | | | | | | | | | | | | | | | | | | | | | | | | | |
| uncon | 14 | 0.155 | 0.158 | 0.160 | 0.159 | 0.148 | 0.179 | 0.176 | 0.149 | 0.154 | 0.160 | 0.158 | 0.160 | 0.163 | - | | | | | | | | | | | | | | | | | | | | | | | | |
| uncon | 15 | 0.160 | 0.158 | 0.161 | 0.172 | 0.164 | 0.168 | 0.167 | 0.173 | 0.149 | 0.164 | 0.178 | 0.179 | 0.171 | 0.178 | - | | | | | | | | | | | | | | | | | | | | | | | |
| uncon | 16 | 0.162 | 0.158 | 0.168 | 0.157 | 0.153 | 0.172 | 0.176 | 0.155 | 0.159 | 0.159 | 0.161 | 0.157 | 0.163 | 0.167 | 0.168 | - | | | | | | | | | | | | | | | | | | | | | | |
| uncon | 17 | 0.153 | 0.156 | 0.165 | 0.156 | 0.151 | 0.167 | 0.166 | 0.152 | 0.153 | 0.160 | 0.158 | 0.154 | 0.158 | 0.155 | 0.175 | 0.171 | - | | | | | | | | | | | | | | | | | | | | | |
| uncon | 18 | 0.153 | 0.147 | 0.148 | 0.152 | 0.160 | 0.163 | 0.173 | 0.154 | 0.154 | 0.163 | 0.167 | 0.162 | 0.161 | 0.170 | 0.158 | 0.152 | 0.166 | - | | | | | | | | | | | | | | | | | | | | |
| uncon | 19 | 0.143 | 0.153 | 0.153 | 0.160 | 0.153 | 0.161 | 0.155 | 0.153 | 0.157 | 0.145 | 0.153 | 0.154 | 0.152 | 0.161 | 0.166 | 0.171 | 0.151 | 0.153 | - | | | | | | | | | | | | | | | | | | | |
| con | 20 | 0.142 | 0.152 | 0.158 | 0.163 | 0.159 | 0.161 | 0.162 | 0.152 | 0.144 | 0.152 | 0.166 | 0.157 | 0.150 | 0.148 | 0.174 | 0.174 | 0.158 | 0.157 | 0.152 | - | | | | | | | | | | | | | | | | | | |
| con | 21 | 0.153 | 0.157 | 0.161 | 0.177 | 0.164 | 0.173 | 0.176 | 0.169 | 0.156 | 0.167 | 0.173 | 0.170 | 0.166 | 0.168 | 0.175 | 0.179 | 0.177 | 0.175 | 0.160 | 0.152 | - | | | | | | | | | | | | | | | | | |
| con | 22 | 0.156 | 0.157 | 0.159 | 0.168 | 0.162 | 0.169 | 0.177 | 0.162 | 0.153 | 0.155 | 0.155 | 0.164 | 0.162 | 0.168 | 0.176 | 0.171 | 0.166 | 0.166 | 0.160 | 0.167 | 0.176 | - | | | | | | | | | | | | | | | | |
| con | 23 | 0.148 | 0.156 | 0.154 | 0.172 | 0.151 | 0.168 | 0.168 | 0.160 | 0.153 | 0.149 | 0.163 | 0.151 | 0.163 | 0.162 | 0.166 | 0.169 | 0.161 | 0.158 | 0.143 | 0.149 | 0.162 | 0.161 | - | | | | | | | | | | | | | | | |
| con | 24 | 0.151 | 0.148 | 0.163 | 0.165 | 0.173 | 0.168 | 0.167 | 0.168 | 0.162 | 0.167 | 0.173 | 0.175 | 0.158 | 0.177 | 0.170 | 0.170 | 0.172 | 0.157 | 0.168 | 0.172 | 0.171 | 0.180 | 0.173 | - | | | | | | | | | | | | | | |
| con | 25 | 0.154 | 0.160 | 0.164 | 0.172 | 0.156 | 0.171 | 0.166 | 0.171 | 0.149 | 0.152 | 0.167 | 0.166 | 0.164 | 0.169 | 0.170 | 0.172 | 0.172 | 0.172 | 0.159 | 0.162 | 0.177 | 0.166 | 0.158 | 0.169 | - | | | | | | | | | | | | | |
| con | 26 | 0.150 | 0.152 | 0.157 | 0.157 | 0.171 | 0.176 | 0.168 | 0.163 | 0.160 | 0.161 | 0.164 | 0.167 | 0.157 | 0.168 | 0.171 | 0.171 | 0.158 | 0.160 | 0.152 | 0.169 | 0.176 | 0.159 | 0.160 | 0.157 | 0.158 | - | | | | | | | | | | | | |
| con | 27 | 0.151 | 0.153 | 0.152 | 0.167 | 0.158 | 0.160 | 0.159 | 0.151 | 0.151 | 0.154 | 0.156 | 0.162 | 0.160 | 0.166 | 0.161 | 0.165 | 0.164 | 0.161 | 0.154 | 0.156 | 0.173 | 0.157 | 0.151 | 0.176 | 0.156 | 0.160 | - | | | | | | | | | | | |
| con | 28 | 0.156 | 0.145 | 0.169 | 0.165 | 0.163 | 0.172 | 0.169 | 0.171 | 0.152 | 0.157 | 0.162 | 0.172 | 0.159 | 0.168 | 0.171 | 0.162 | 0.167 | 0.165 | 0.168 | 0.162 | 0.177 | 0.167 | 0.166 | 0.175 | 0.167 | 0.164 | 0.165 | - | | | | | | | | | | |
| con | 29 | 0.151 | 0.155 | 0.173 | 0.157 | 0.154 | 0.180 | 0.173 | 0.161 | 0.159 | 0.158 | 0.161 | 0.158 | 0.168 | 0.156 | 0.177 | 0.162 | 0.157 | 0.171 | 0.166 | 0.159 | 0.171 | 0.163 | 0.161 | 0.175 | 0.167 | 0.167 | 0.168 | 0.165 | - | | | | | | | | | |
| con | 30 | 0.149 | 0.161 | 0.153 | 0.167 | 0.156 | 0.159 | 0.170 | 0.157 | 0.149 | 0.159 | 0.161 | 0.161 | 0.155 | 0.170 | 0.168 | 0.169 | 0.152 | 0.157 | 0.155 | 0.159 | 0.164 | 0.155 | 0.159 | 0.154 | 0.162 | 0.163 | 0.159 | 0.171 | 0.168 | - | | | | | | | | |
| con | 31 | 0.166 | 0.169 | 0.174 | 0.178 | 0.166 | 0.174 | 0.179 | 0.169 | 0.160 | 0.171 | 0.181 | 0.168 | 0.162 | 0.166 | 0.185 | 0.179 | 0.176 | 0.164 | 0.170 | 0.151 | 0.177 | 0.179 | 0.163 | 0.177 | 0.166 | 0.177 | 0.161 | 0.175 | 0.176 | 0.170 | - | | | | | | | |
| con | 32 | 0.156 | 0.162 | 0.163 | 0.176 | 0.168 | 0.173 | 0.176 | 0.160 | 0.158 | 0.169 | 0.173 | 0.168 | 0.166 | 0.170 | 0.179 | 0.169 | 0.168 | 0.161 | 0.165 | 0.163 | 0.172 | 0.158 | 0.158 | 0.178 | 0.169 | 0.164 | 0.158 | 0.173 | 0.177 | 0.167 | 0.176 | - | | | | | | |
| con | 33 | 0.148 | 0.153 | 0.160 | 0.167 | 0.160 | 0.159 | 0.155 | 0.158 | 0.146 | 0.149 | 0.162 | 0.167 | 0.153 | 0.168 | 0.161 | 0.171 | 0.168 | 0.163 | 0.158 | 0.153 | 0.165 | 0.163 | 0.156 | 0.165 | 0.155 | 0.163 | 0.153 | 0.169 | 0.160 | 0.163 | 0.168 | 0.167 | - | | | | | |
| con | 34 | 0.135 | 0.152 | 0.154 | 0.166 | 0.156 | 0.167 | 0.156 | 0.159 | 0.148 | 0.151 | 0.157 | 0.163 | 0.158 | 0.172 | 0.158 | 0.163 | 0.158 | 0.160 | 0.150 | 0.149 | 0.163 | 0.162 | 0.155 | 0.164 | 0.161 | 0.157 | 0.147 | 0.168 | 0.154 | 0.154 | 0.173 | 0.165 | 0.161 | - | | | | |
| con | 35 | 0.145 | 0.151 | 0.156 | 0.163 | 0.158 | 0.167 | 0.164 | 0.161 | 0.160 | 0.159 | 0.163 | 0.163 | 0.153 | 0.165 | 0.179 | 0.176 | 0.160 | 0.165 | 0.153 | 0.154 | 0.163 | 0.161 | 0.156 | 0.173 | 0.163 | 0.161 | 0.163 | 0.167 | 0.164 | 0.160 | 0.167 | 0.161 | 0.168 | 0.159 | - | | | |
| con | 36 | 0.143 | 0.154 | 0.152 | 0.164 | 0.153 | 0.152 | 0.168 | 0.161 | 0.148 | 0.143 | 0.163 | 0.157 | 0.154 | 0.162 | 0.164 | 0.166 | 0.158 | 0.163 | 0.150 | 0.155 | 0.162 | 0.168 | 0.159 | 0.168 | 0.159 | 0.168 | 0.152 | 0.162 | 0.170 | 0.152 | 0.159 | 0.166 | 0.156 | 0.151 | 0.161 | - | | |
| con | 37 | 0.160 | 0.151 | 0.161 | 0.172 | 0.161 | 0.166 | 0.164 | 0.158 | 0.152 | 0.153 | 0.163 | 0.160 | 0.156 | 0.158 | 0.165 | 0.168 | 0.164 | 0.167 | 0.152 | 0.159 | 0.156 | 0.165 | 0.163 | 0.159 | 0.159 | 0.165 | 0.156 | 0.173 | 0.169 | 0.154 | 0.164 | 0.161 | 0.164 | 0.154 | 0.161 | 0.150 | - | |
| con | 38 | 0.156 | 0.158 | 0.161 | 0.153 | 0.151 | 0.166 | 0.168 | 0.146 | 0.154 | 0.154 | 0.152 | 0.159 | 0.145 | 0.159 | 0.181 | 0.166 | 0.149 | 0.172 | 0.158 | 0.158 | 0.174 | 0.168 | 0.169 | 0.166 | 0.159 | 0.169 | 0.164 | 0.169 | 0.163 | 0.154 | 0.174 | 0.174 | 0.161 | 0.163 | 0.161 | 0.155 | 0.157 | - |

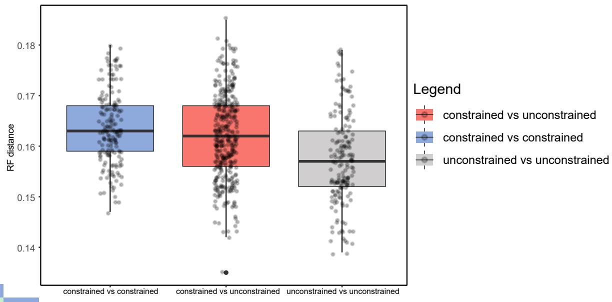

A)

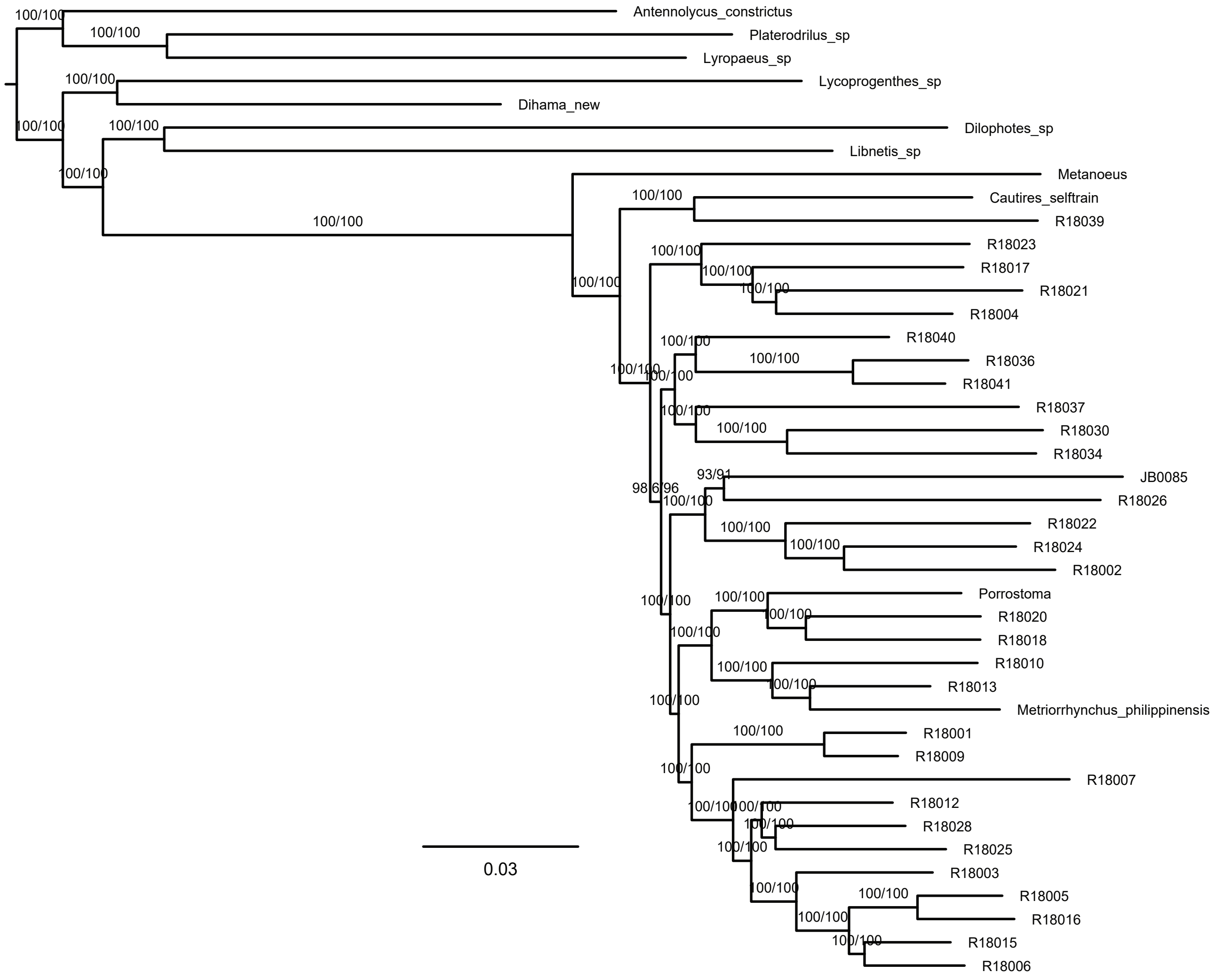

B)

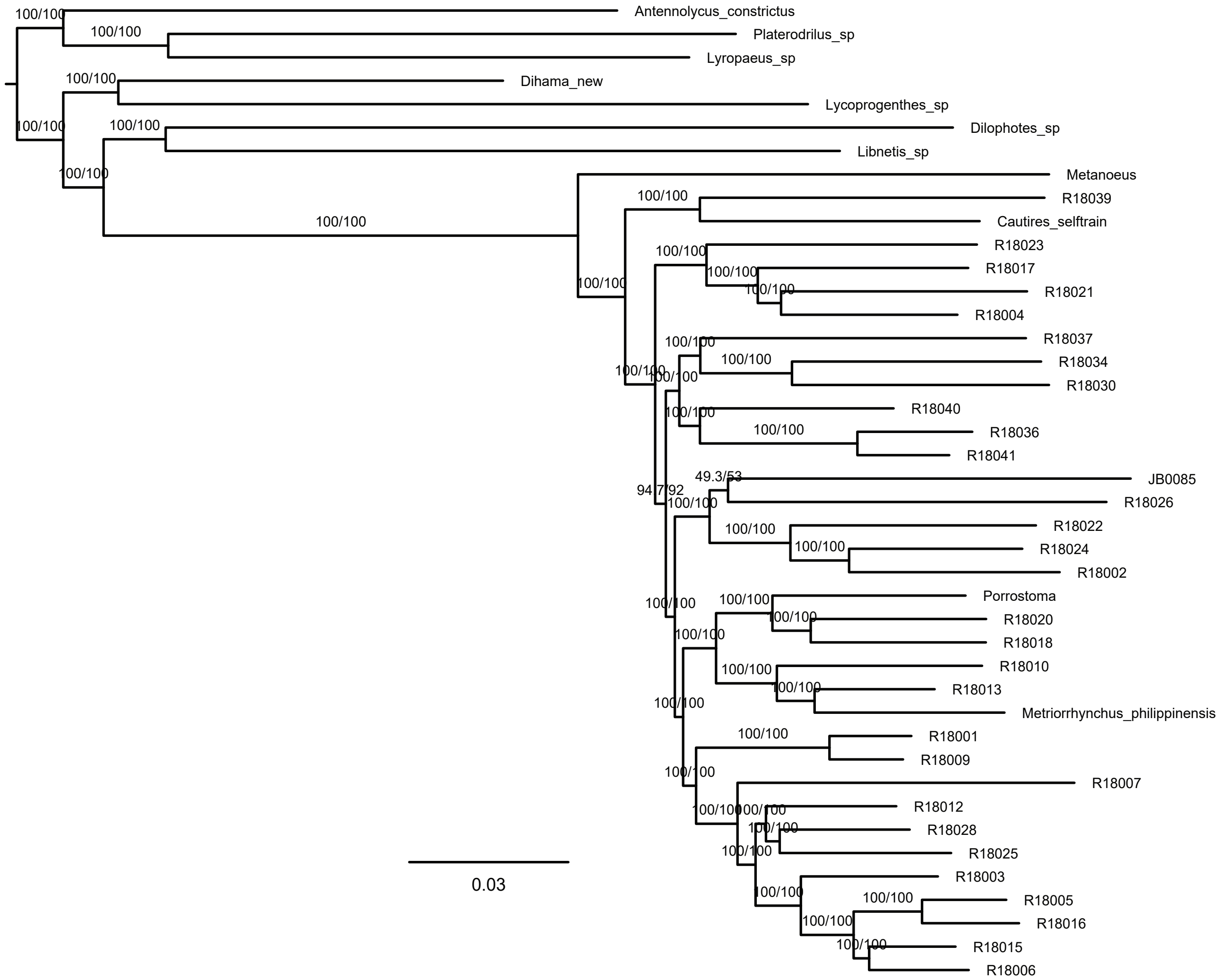

Figure S1. Maximum likelihood trees from IQ-TREE amino acid analysis of dataset A-4109-AA A) with partitioning by gene and B) without partitioning. The depicted branch support values represent SH-aLRT and ultrafast bootstrap.

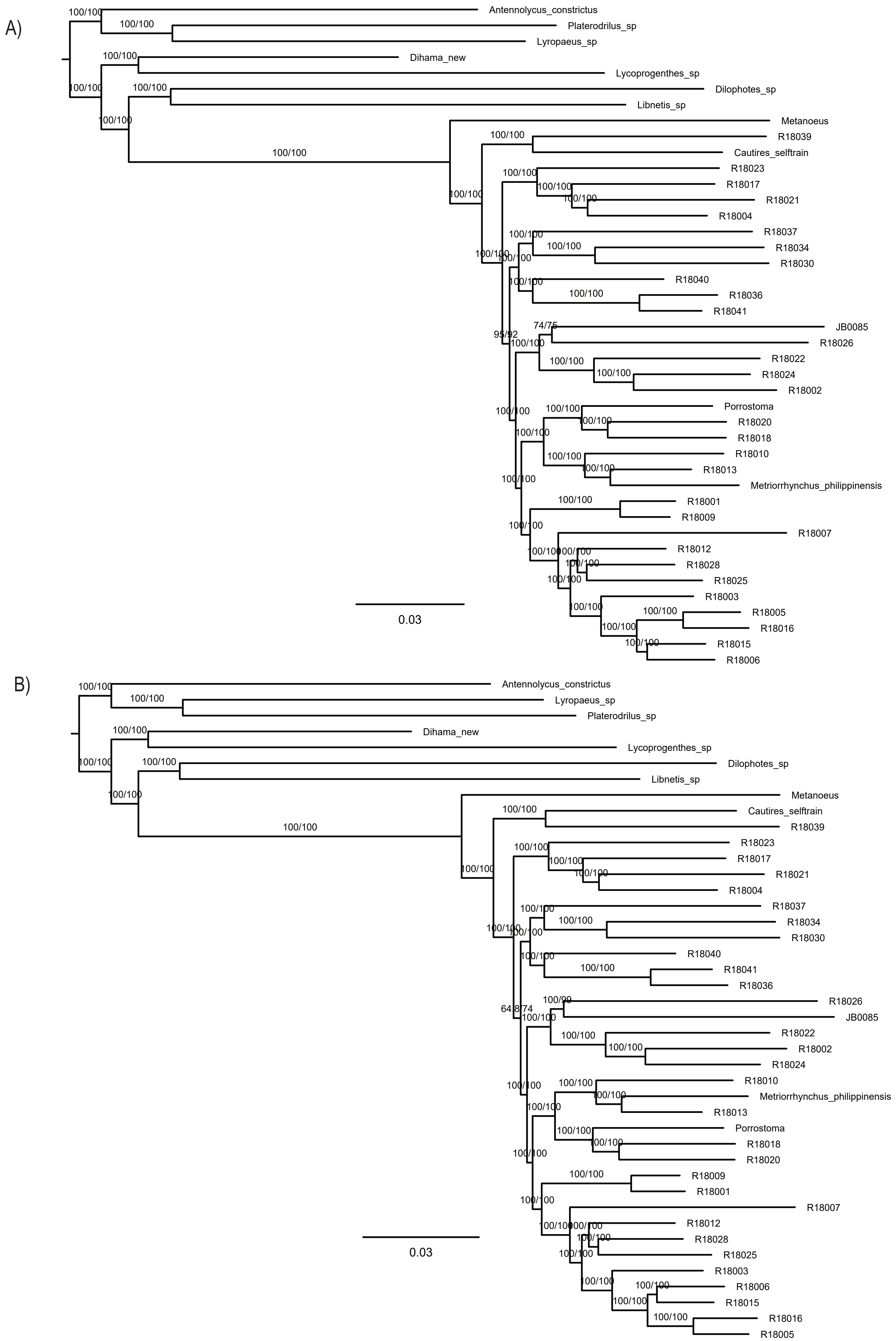

Figure S2. Maximum likelihood trees from IQ-TREE amino acid analysis of datasets A) A-4109-AA with optimization of the partitioning scheme and B) D-3370-AA_Bacoca with partitioning by gene. The depicted branch support values represent SH-aLRT and ultrafast bootstrap.

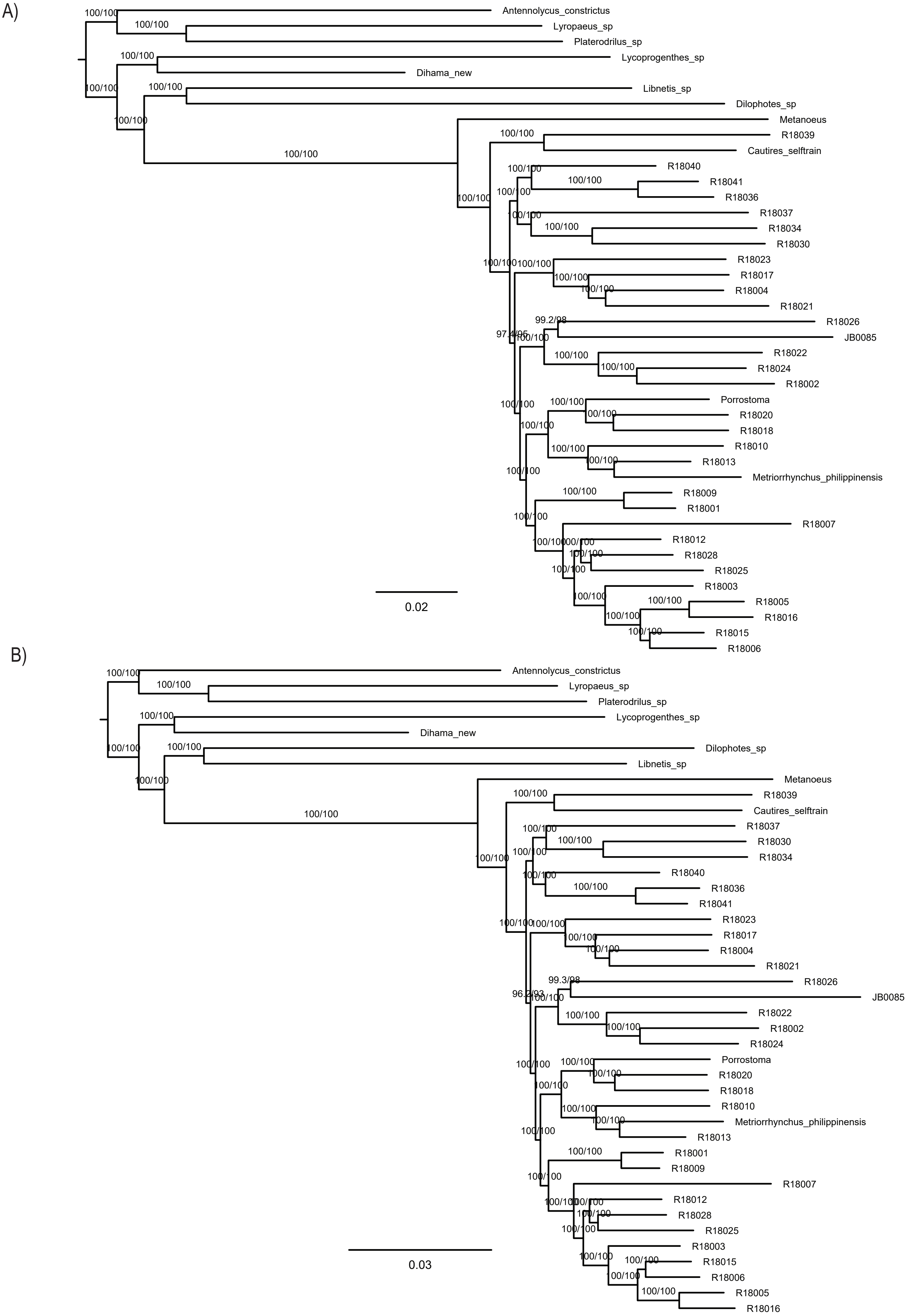

Figure S3. Maximum likelihood trees from IQ-TREE amino acid analysis of datasets A) F-1490-AA_Bacoca_decisive and B) J-2129-AA_MARE both datsets were partitioned by gene. The depicted branch support values represent SH-aLRT and ultrafast bootstrap.

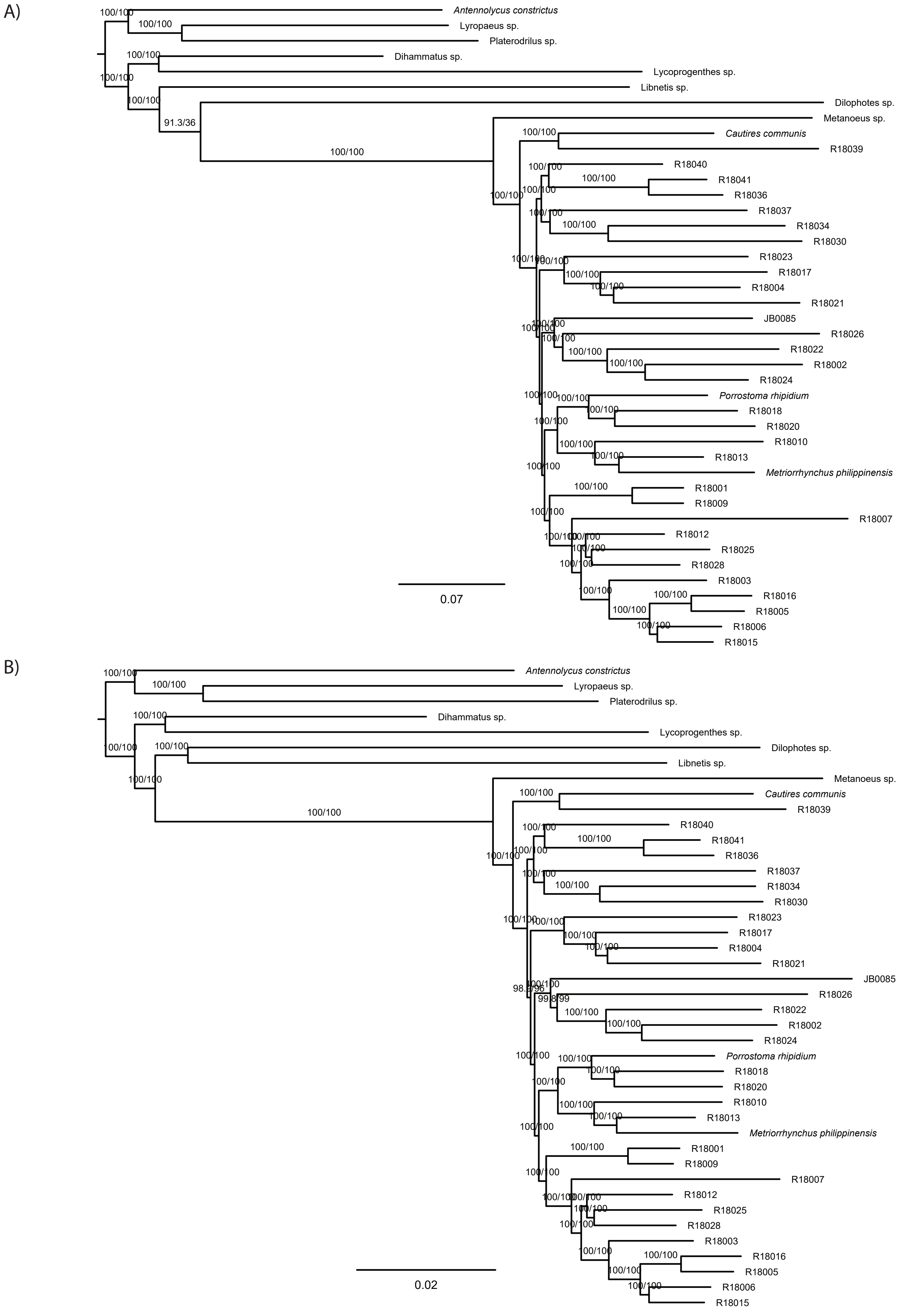

Figure S4. Maximum likelihood trees from IQ-TREE nucleotide analysis of datasets A) B-4109-NT and B) E-4109-NT2 using only second codon positions, both datsets were partitioned by gene. The depicted branch support values represent SH-aLRT and ultrafast bootstrap.

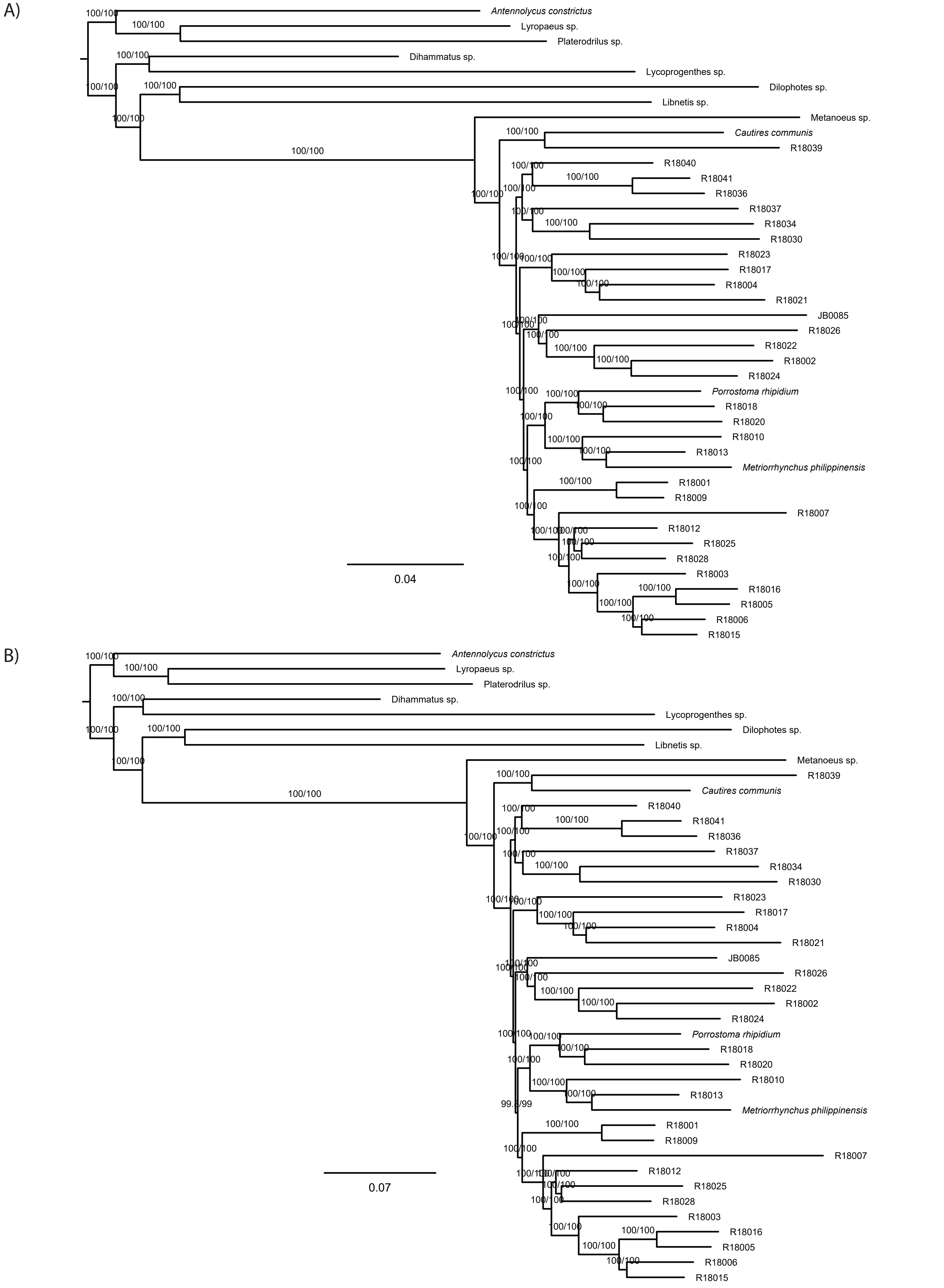

Figure S5. Maximum likelihood trees from IQ-TREE nucleotide analysis of datasets A) C-4109-NT12 using codonpositions 1+2 and B) G-NT-1767_MaxSymTest, both datsets were partitioned by gene. The depicted branch support values represent SH-aLRT and ultrafast bootstrap.

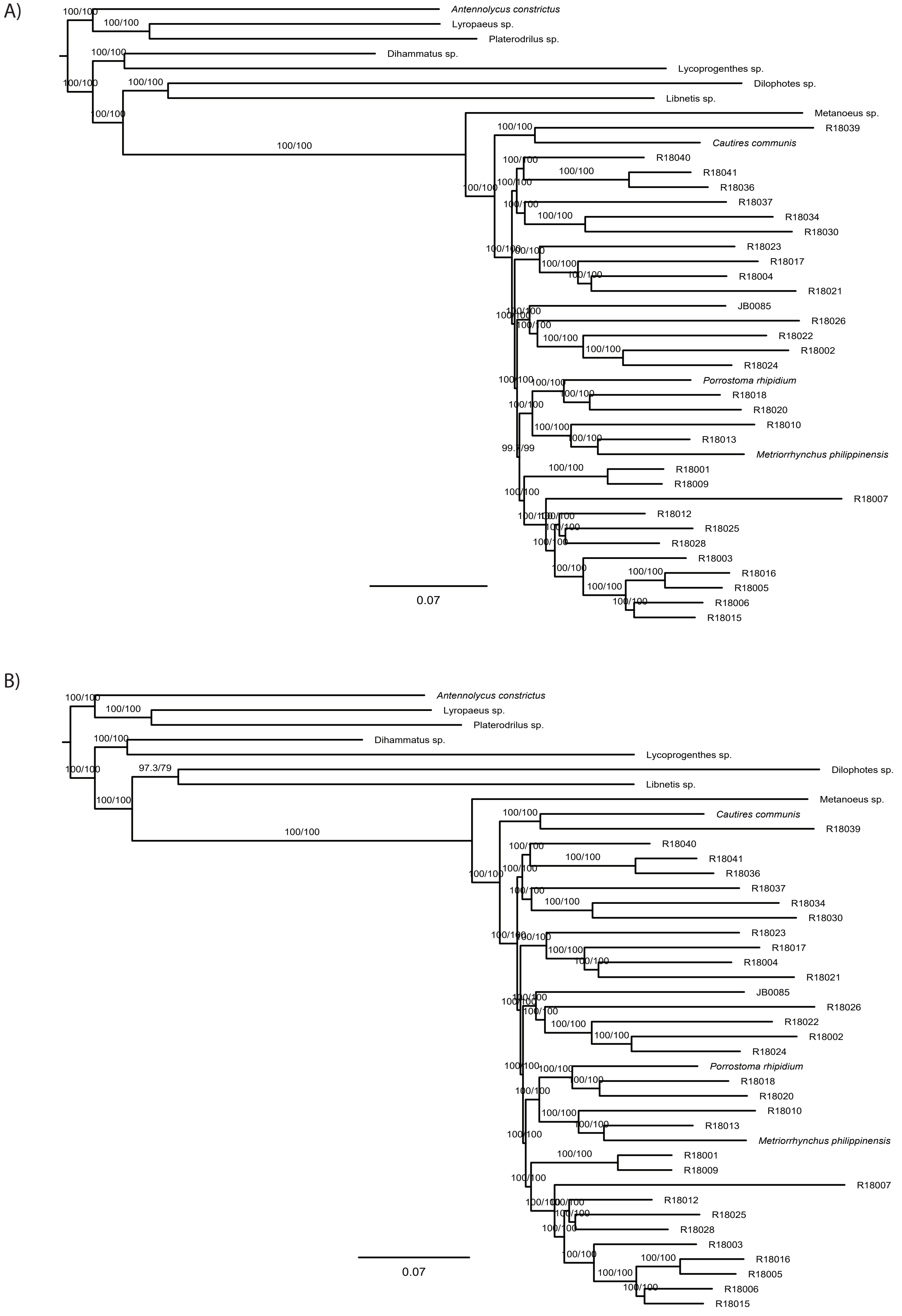

Figure S6. Maximum likelihood trees from IQ-TREE nucleotide analysis of datasets A) H-NT-1645_MaxSymTestmarginal and B) I-NT-3905_MaxSymTestInternal, both datsets were partitioned by gene. The depicted branch support values represent SH-aLRT and ultrafast bootstrap.

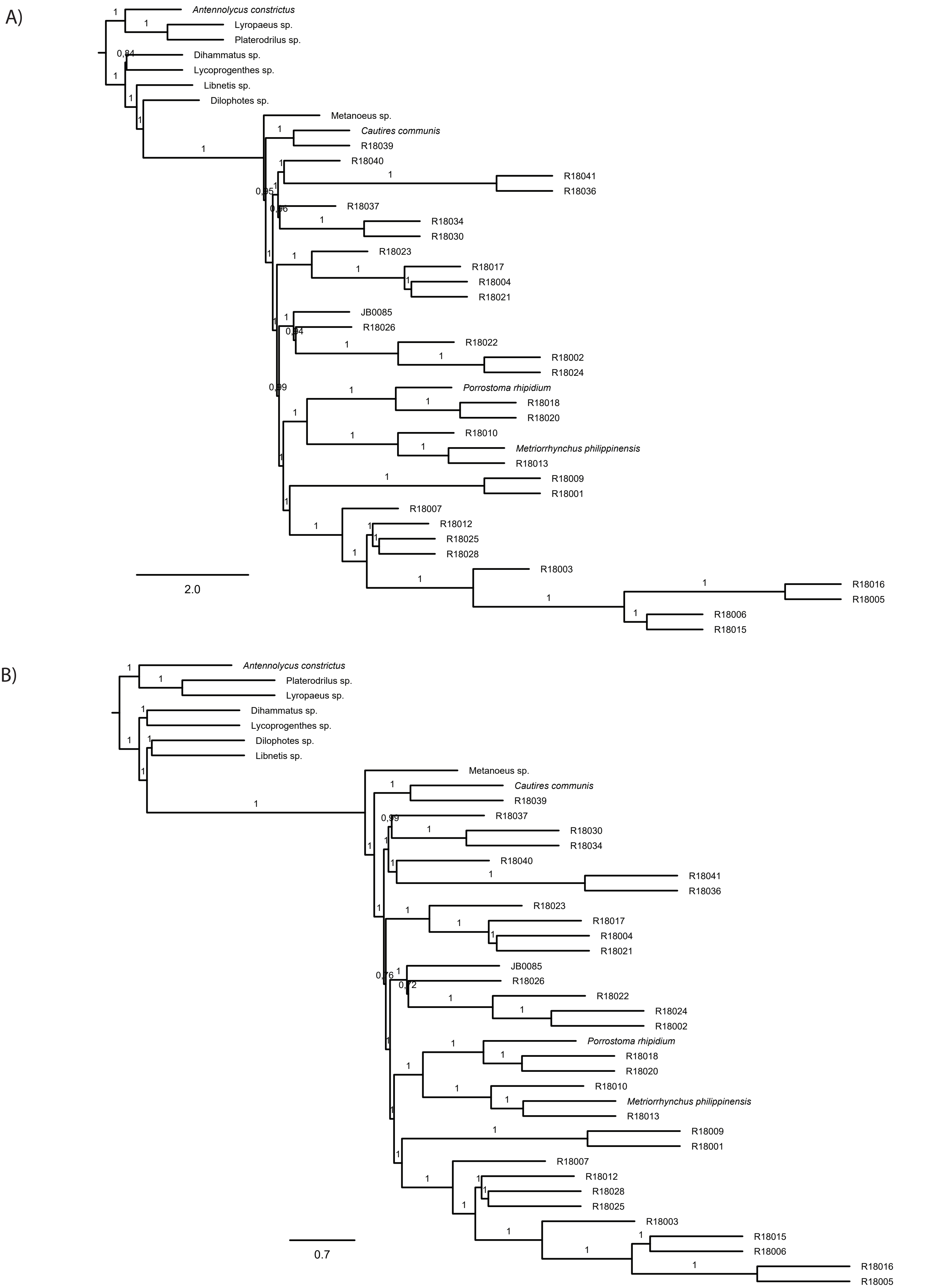

Figure S7. A) and B) ASTRAL species trees with branch lengths in coalescent units as resulted from the analyses of individual IQ-TREE maximum likelihood gene trees of A) nucleotide dataset B-4109-NT and B) amino acid dataset A-4199-AA. Numbers on nodes show local posterior probabilities (pp1). Quartet support for the alternative topologies (q1, q2 and q3), total number of induced quartet trees in the gene trees that support the alternative topologies (f1, f2, f3) and local posterior probabilities (pp1, pp2, pp3) are available at Dryad repository.

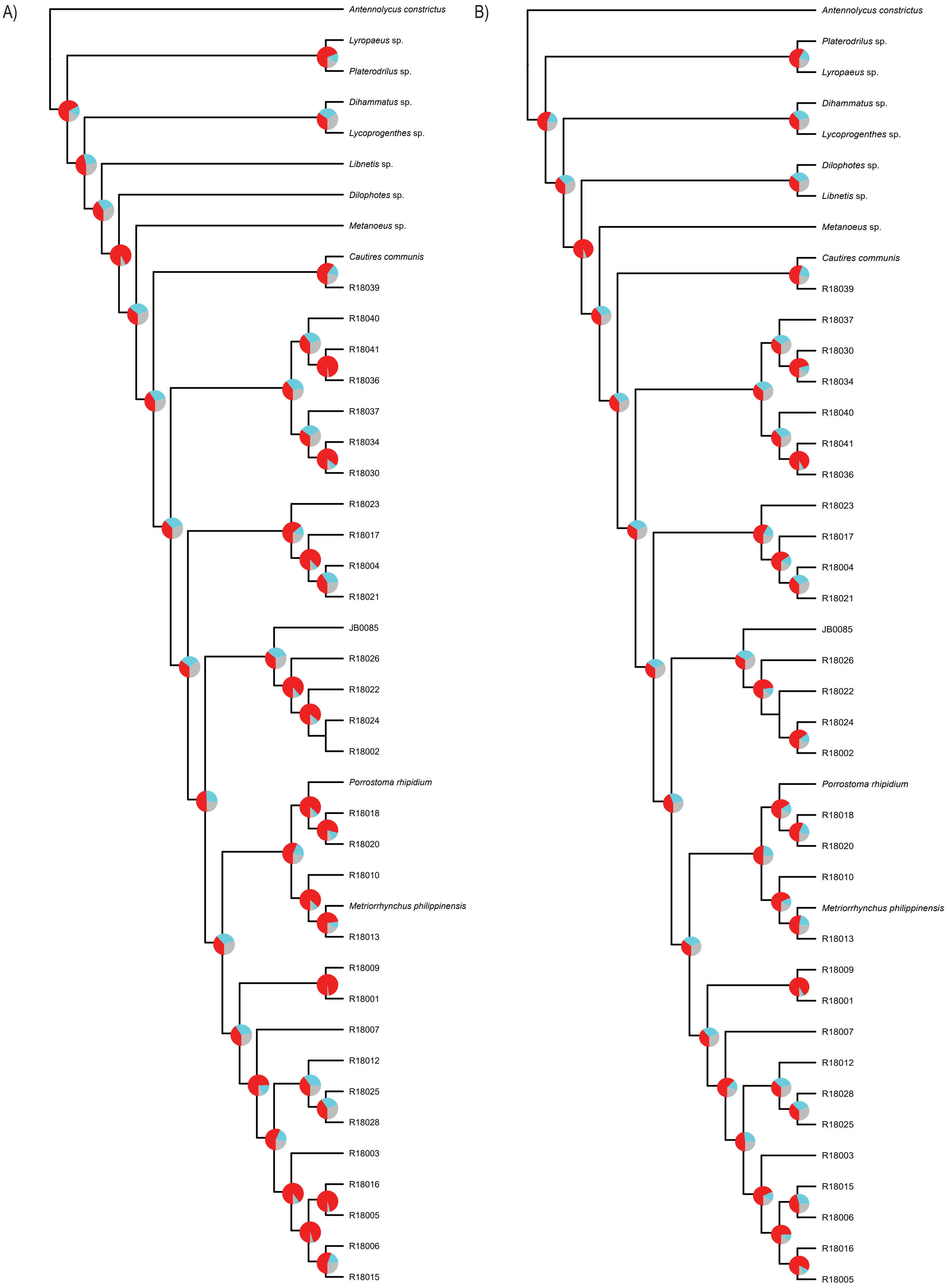

Figure S8. Phylogenetic relationships of Metriorrhynchini, resulted from the summary coalescent phylogenetic analysis with ASTRAL, when analyzing the full set of gene trees (4109 gene trees inferred at the A) nucleotide and B) amino acids level). Pie charts on branches show ASTRAL quartet supports q1,q2,q3 (quartet-based frequencies of alternative quadripartition topologies around a given internode).Tree topologies correspond with Figure S7.

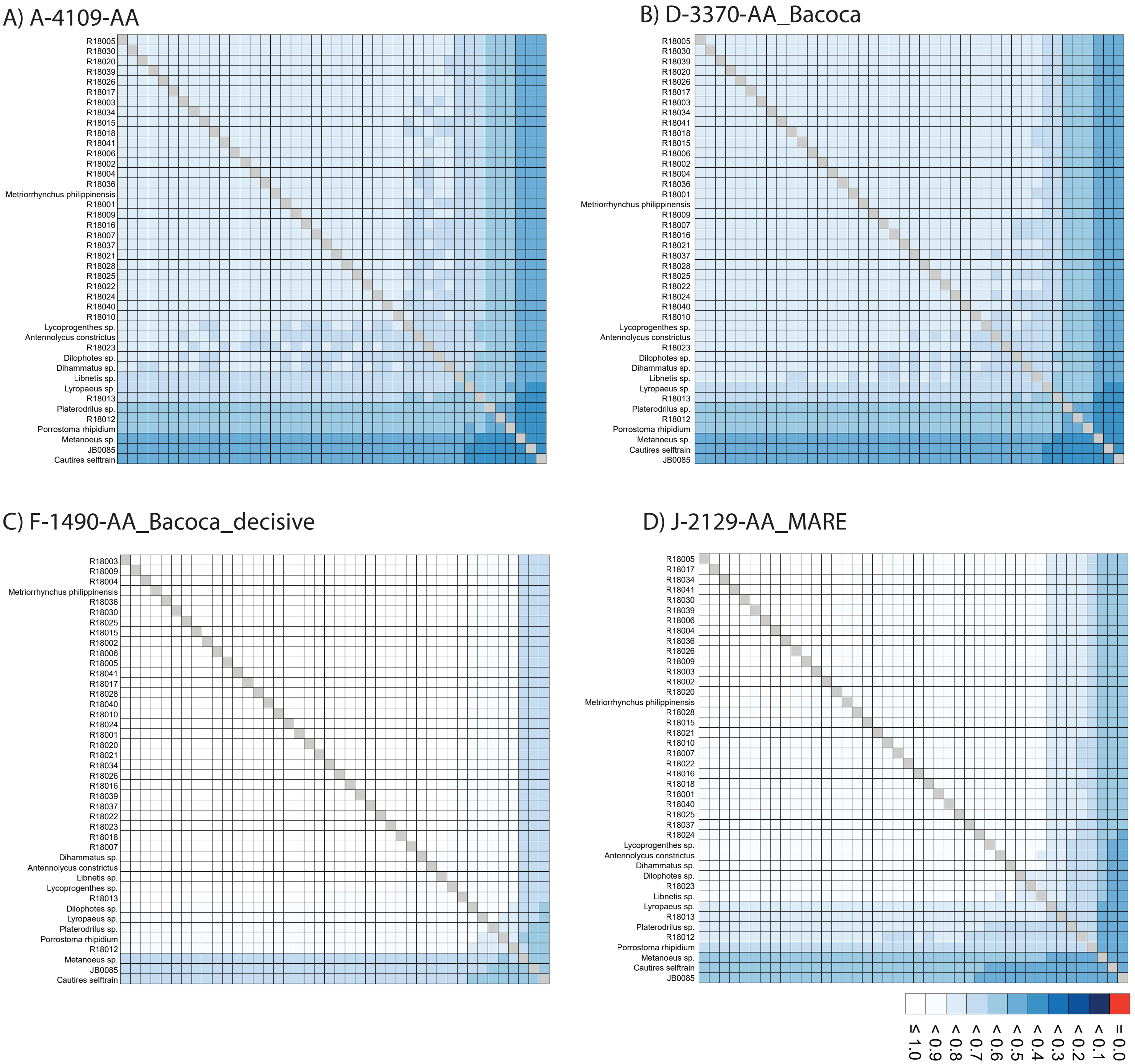

Figure S9. A-D AliStat rectangular heatmaps showing pairwise alignment completeness scores for all species included in the analyzed amino acid supermatrixes. The abbreviations of datasets correspond to Table SX. Values closer to 1 indicate higher completeness scores for the pairwise sequence comparisons.

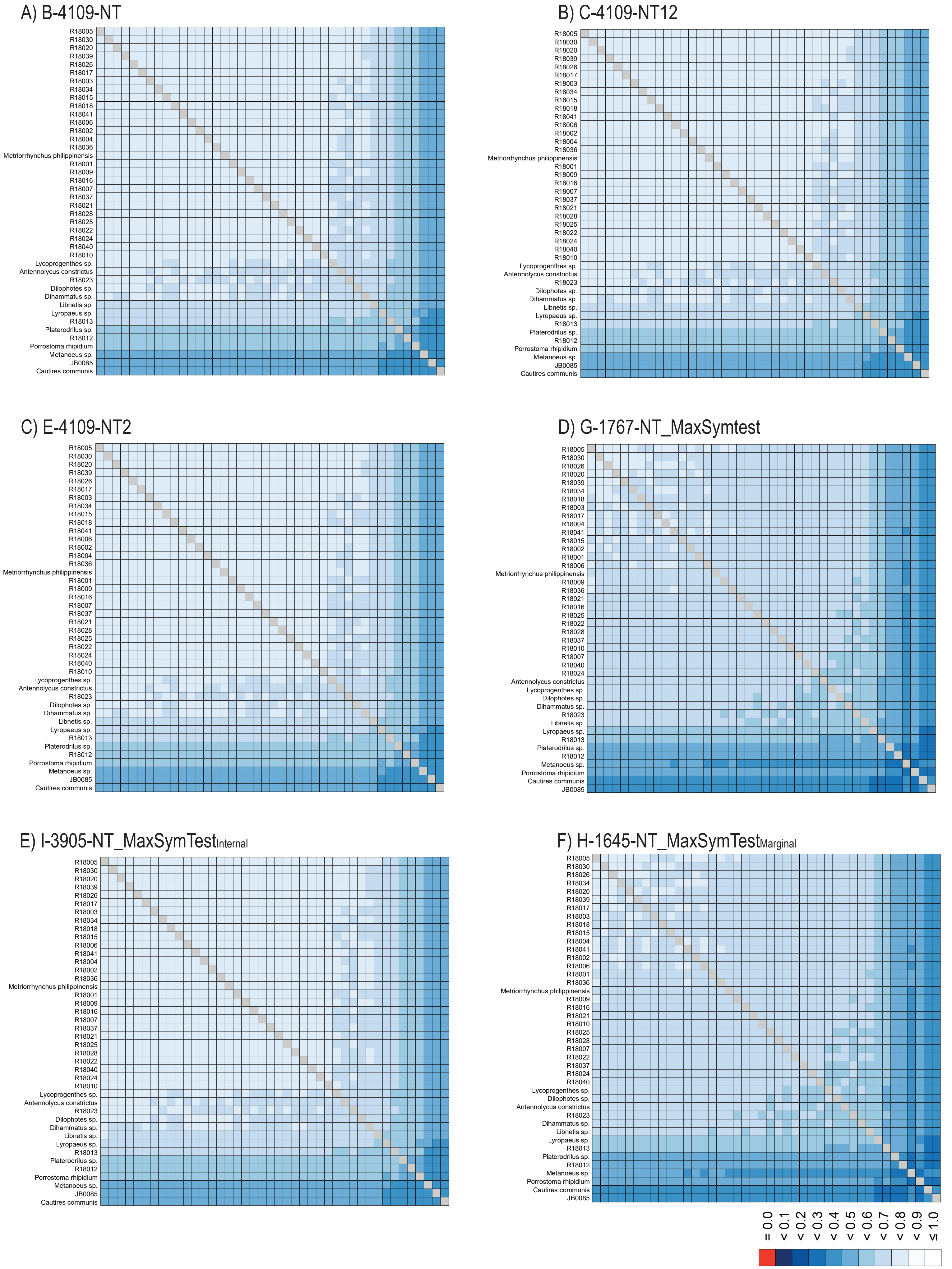

Figure S10. A-F AliStat rectangular heatmaps showing pairwise alignment completeness scores for all species included in the analyzed nucleotide supermatrixes. The abbreviations of datasets correspond to Table SX. Values closer to 1 indicate higher completeness scores for the pairwise sequence comparisons.

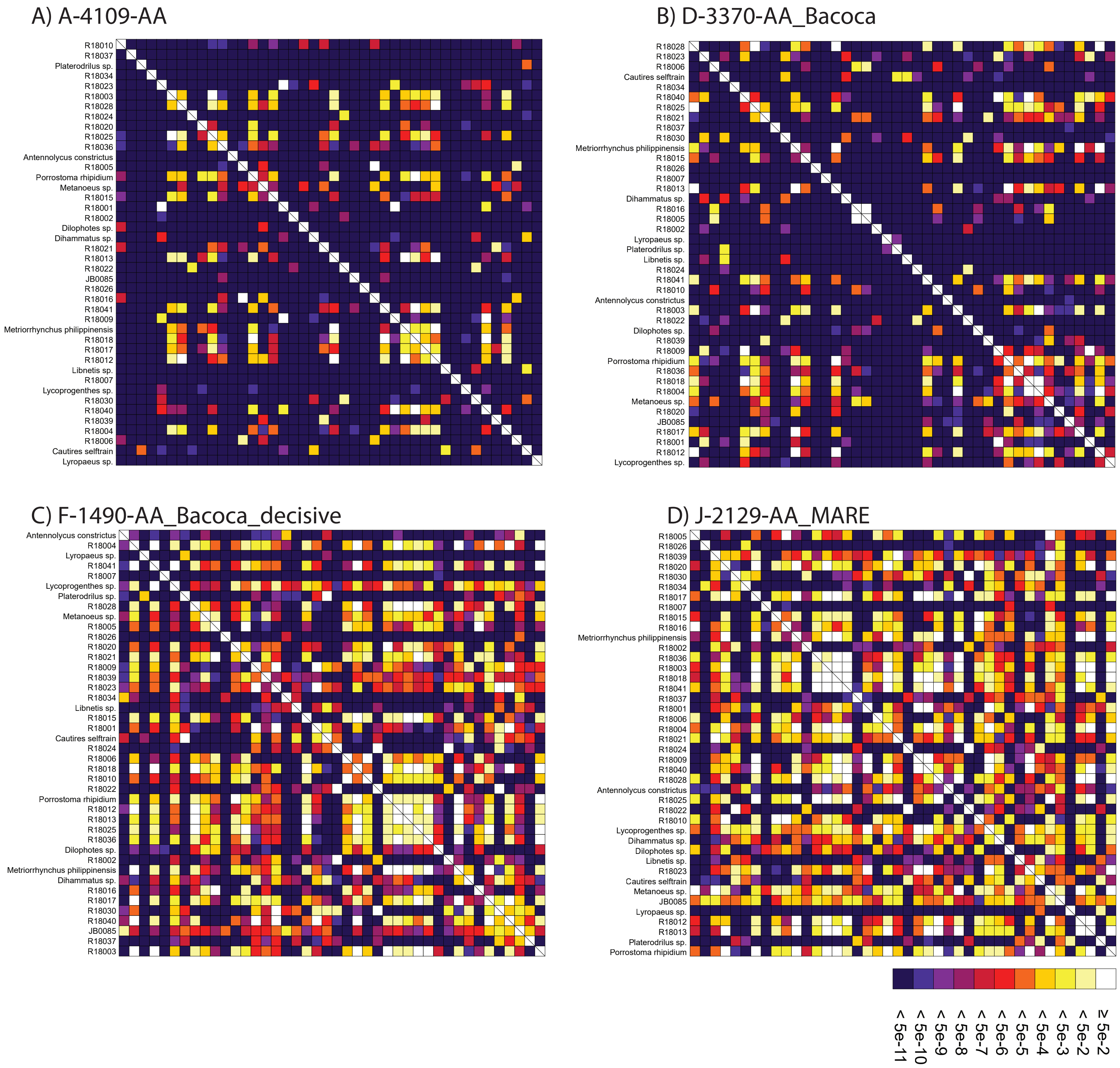

Figure S11. A-D Rectangular heat map calculated with SymTest showing p-values for the pairwise Bowker's tests in the analyzed amino acid supermatrixes. The abbreviations of datasets correspond to Table SX. Darker boxes indicate lower p-values and thus larger deviation from evolution under SRH conditions.

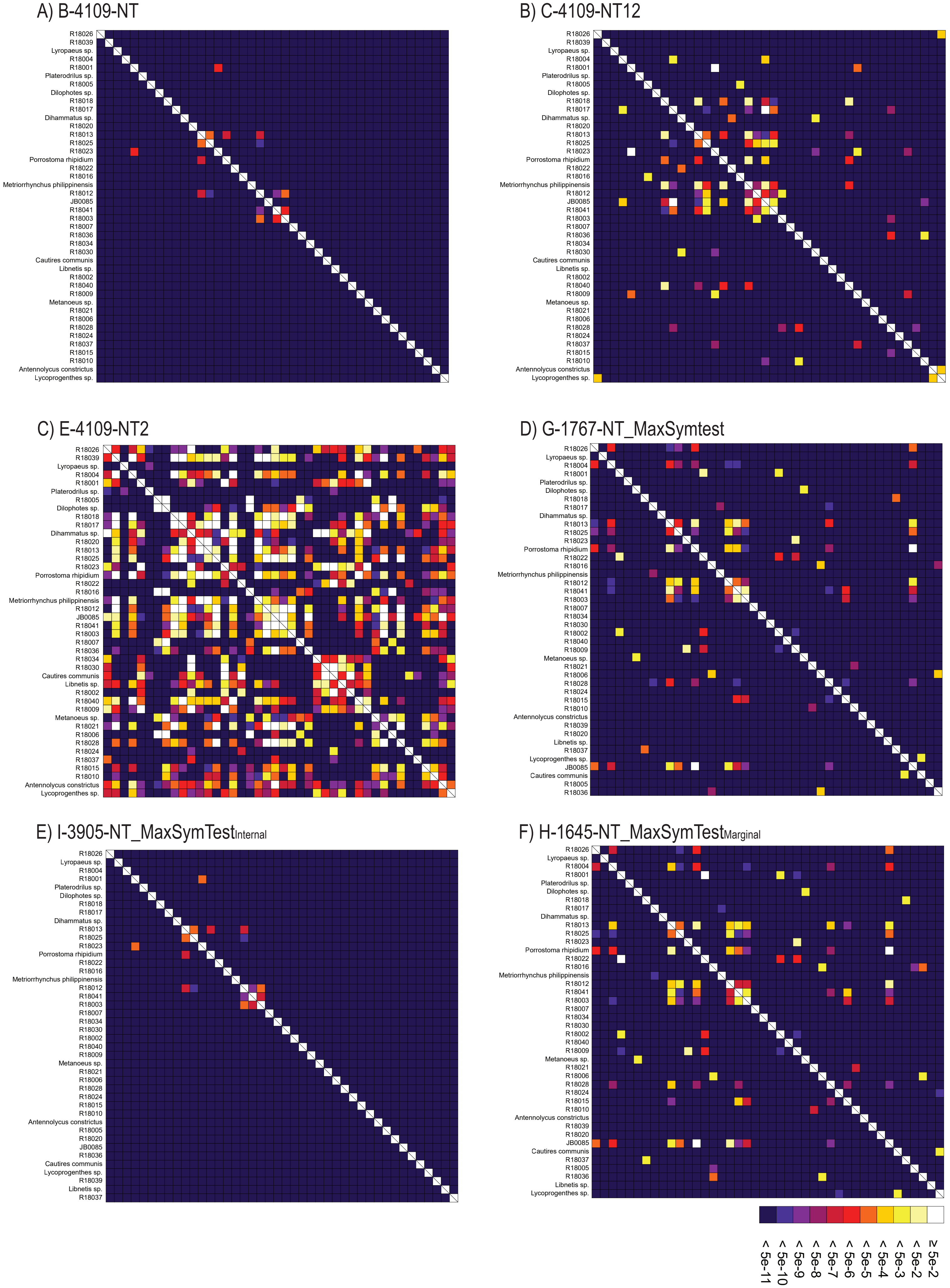

Figure S12. A-F Rectangular heat map calculated with SymTest showing p-values for the pairwise Bowker's tests in the analyzed nucleotide supermatrixes. The abbreviations of datasets correspond to Table SX. Darker boxes indicate lower p-values and thus larger deviation from evolution under SRH conditions.

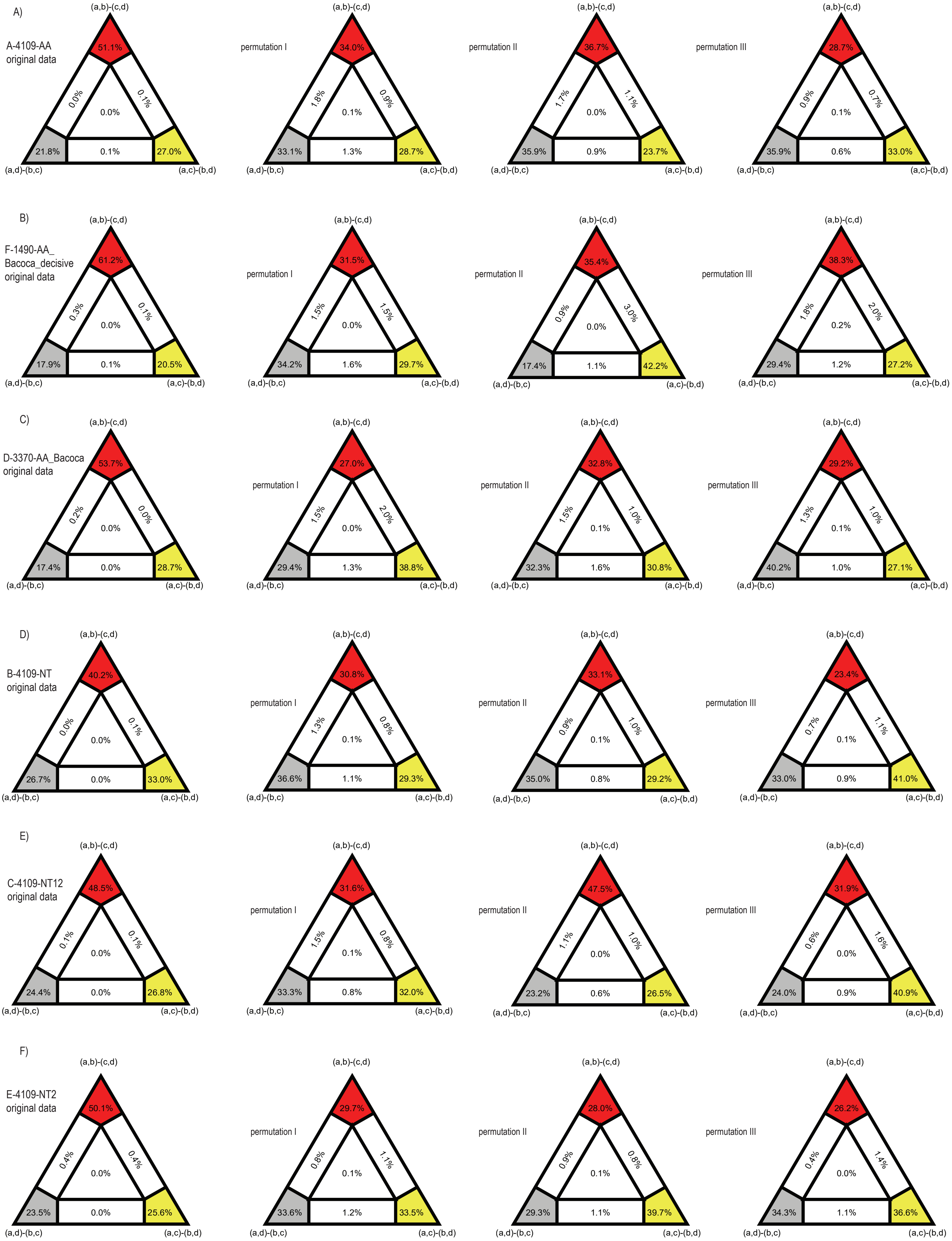

a) Cautirina + Metanoeina
b) Leptotrichaline clade
c) Procautirine clade
d) Trichaline, Porrostomine, Cladophorine clades

Figure S13. Results of FcLM analyses testing alternative phylogenetic hypotheses about placement of Procautirine and Leptotrichaline clades applied for various supermatrixes. The first column shows the results of FcLM when the original data were analyzed. The second column shows the results of FcLM after phylogenetic signal had been eliminated from data. The third column show the results of FcLM after elimination of phylogenetic signal and inhomogeneous amino-acid or nucleotide composition. The fourth column show the results of FcLM after elimination of phylogenetic signal, inhomogeneous amino-acid or nucleotide composition and with randomized data coverage within all meta-partitions (see Supplementary methods for details)

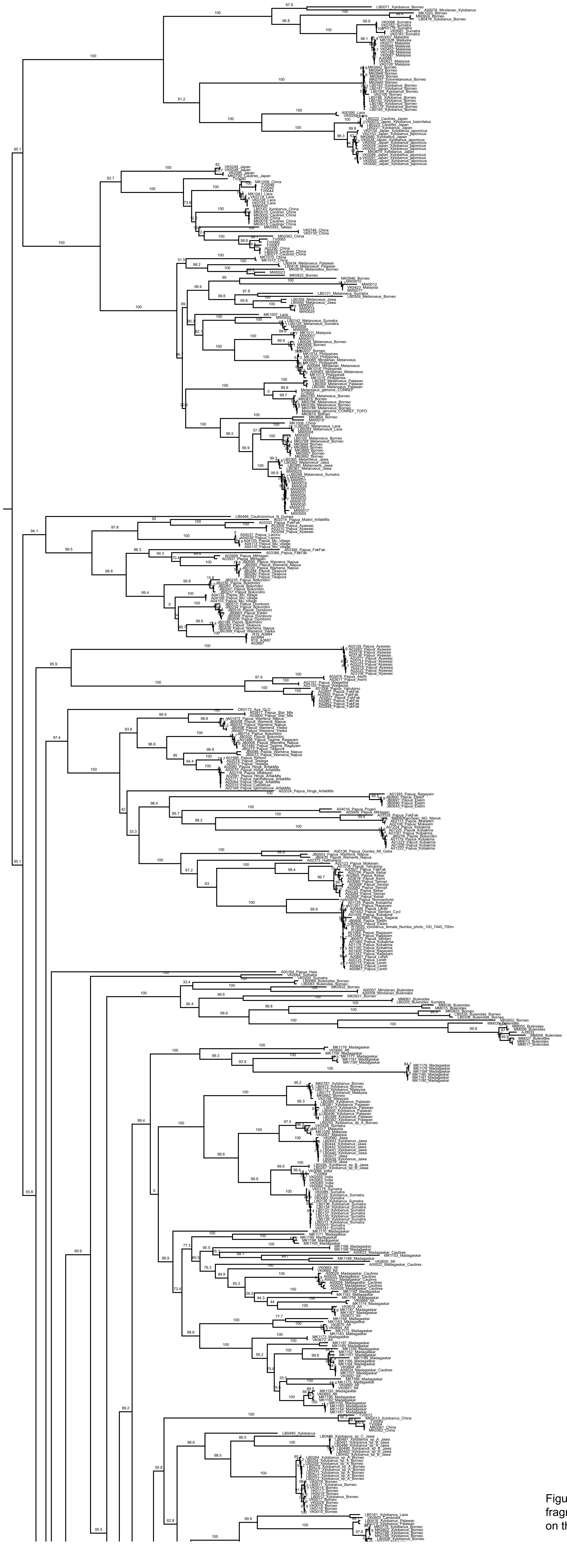

Figure S14. Maximum likelihood tree recovered by the analysis of the fulldataset (mitochondrial fragments). Depicted numbers above branches represent SH-alrt support values. The figure continues on the next page.

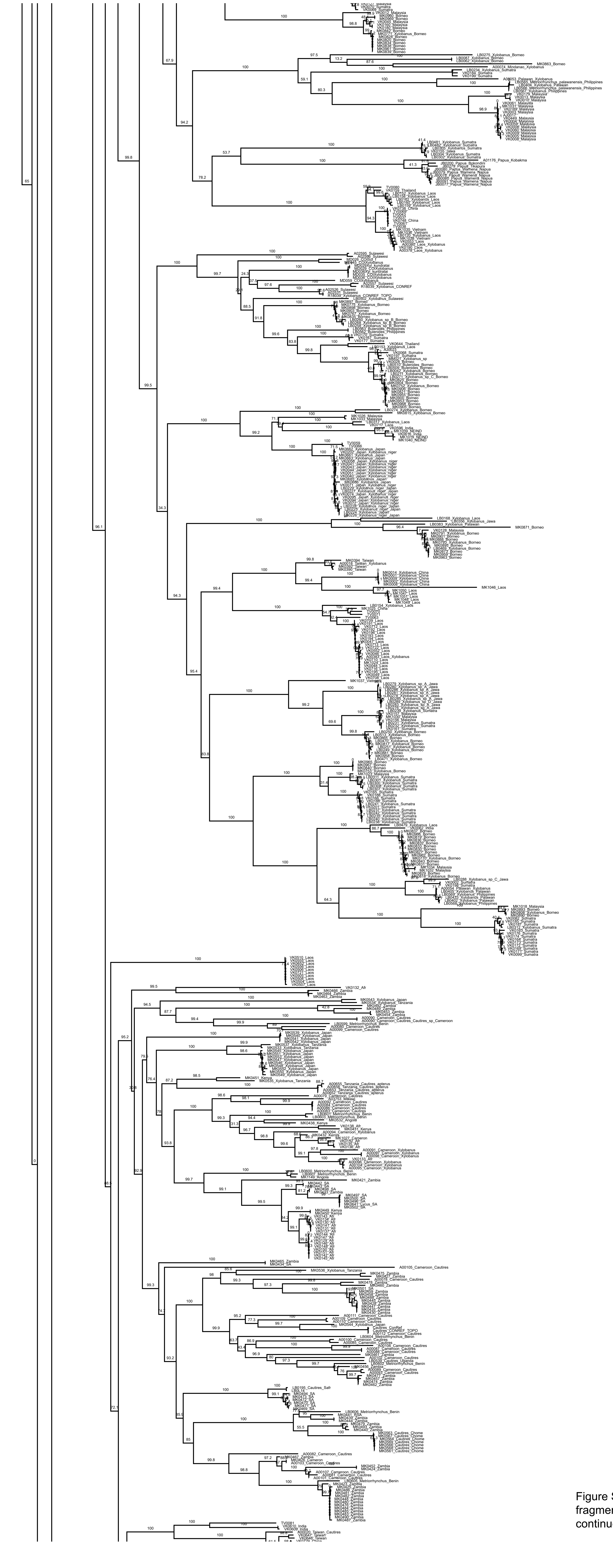

Figure S14. Maximum likelihood tree recovered by the analysis of the fulldataset (mitochondrial fragments). Depicted numbers above branches represent SH-alrt support values. The figure continues on the next page.

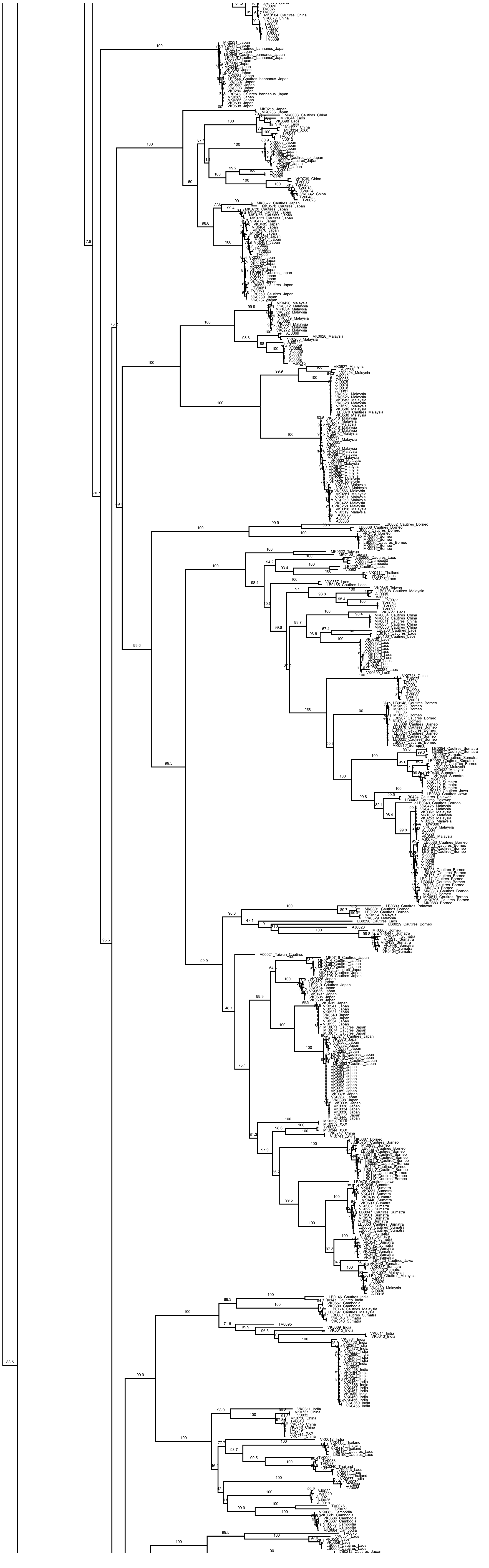

Figure S14. Maximum likelihood tree recovered by the analysis of the fulldataset (mitochondrial fragments). Depicted numbers above branches represent SH-alrt support values. The figure continues on the next page.

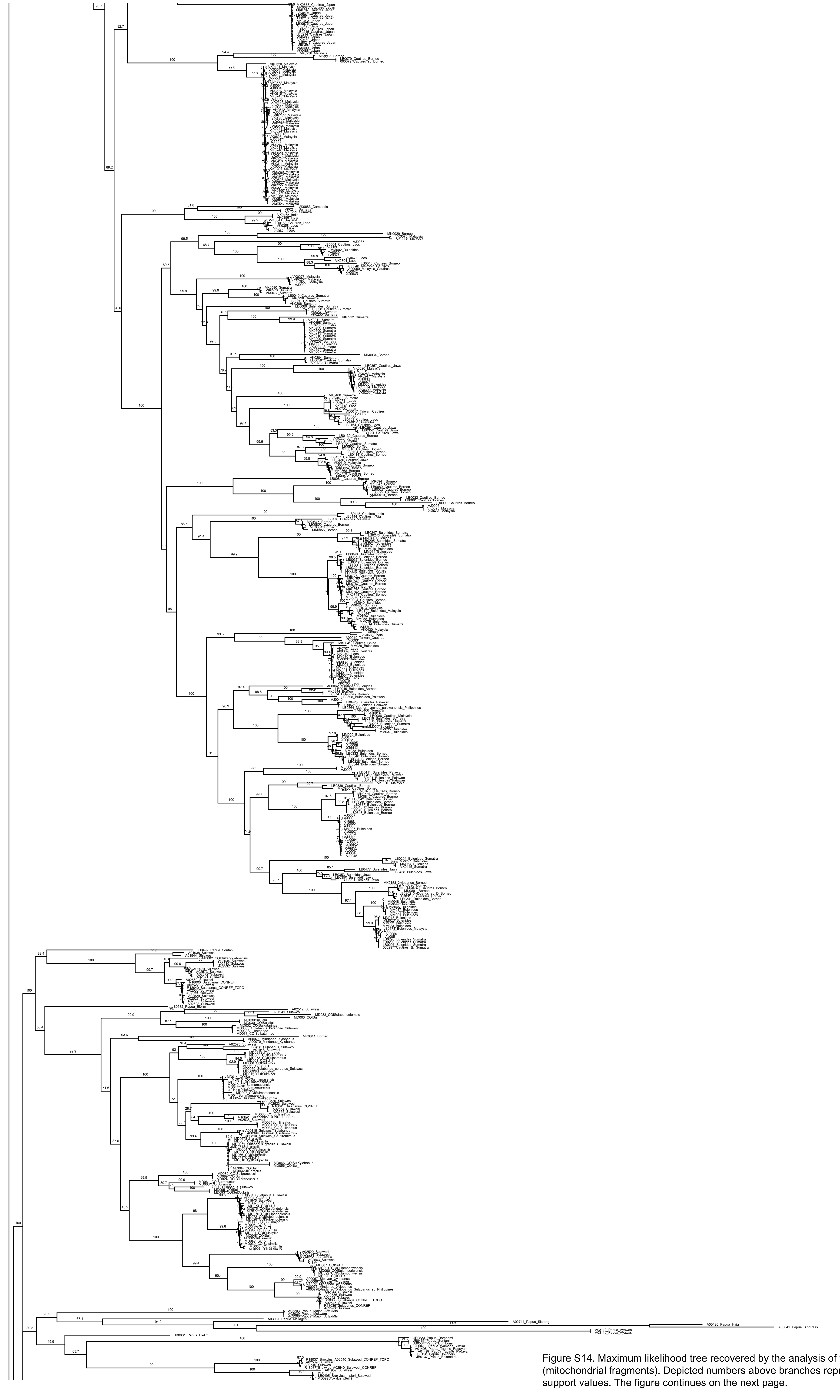

Figure S14. Maximum likelihood tree recovered by the analysis of the fulldataset (mitochondrial fragments). Depicted numbers above branches represent SH-alrt support values. The figure continues on the next page.

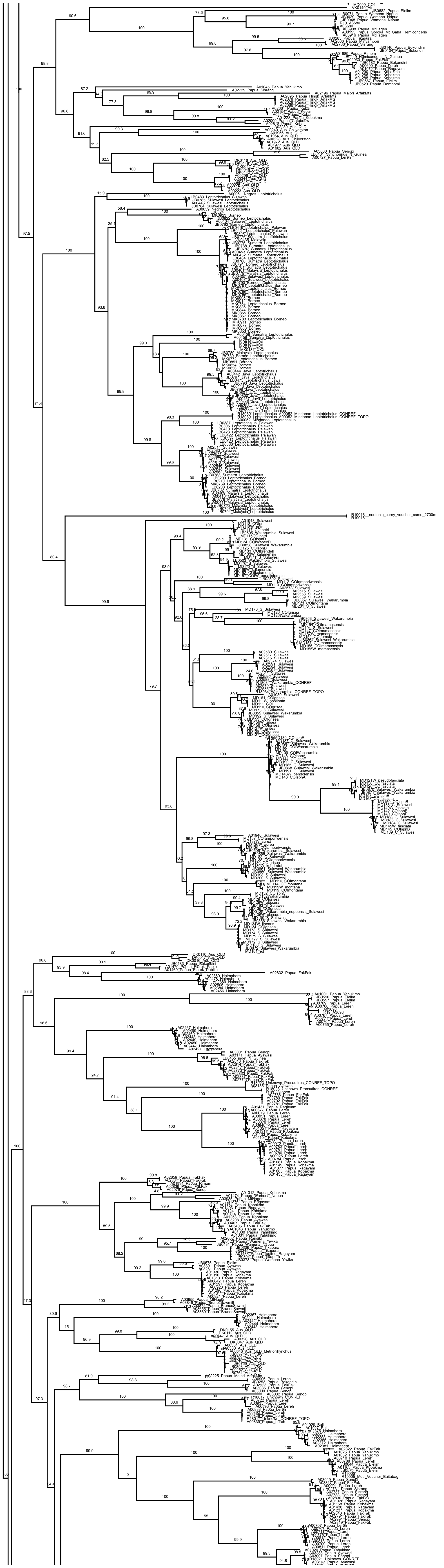

Figure S14. Maximum likelihood tree recovered by the analysis of the fulldataset (mitochondrial fragments). Depicted numbers above branches represent SH-alrt support values. The figure continues on the next page.

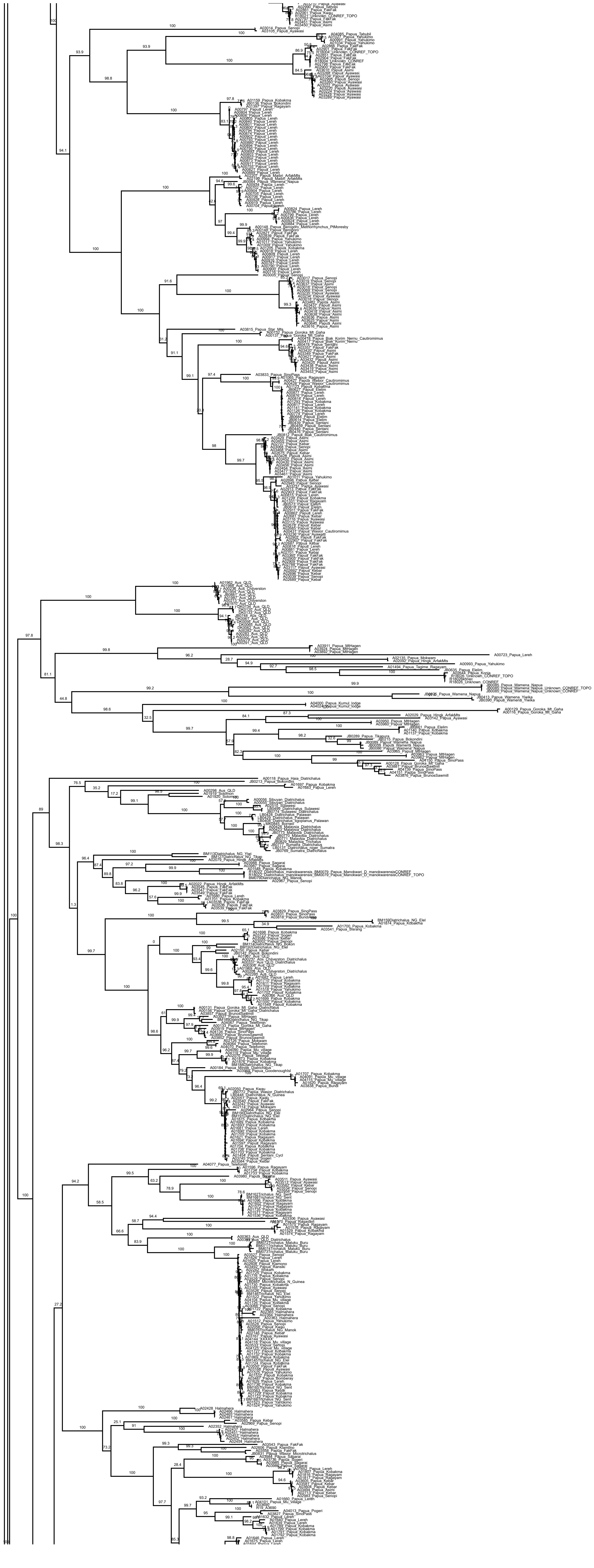

Figure S14. Maximum likelihood tree recovered by the analysis of the fulldataset (mitochondrial fragments). Depicted numbers above branches represent SH-alrt support values. The figure continues on the next page.

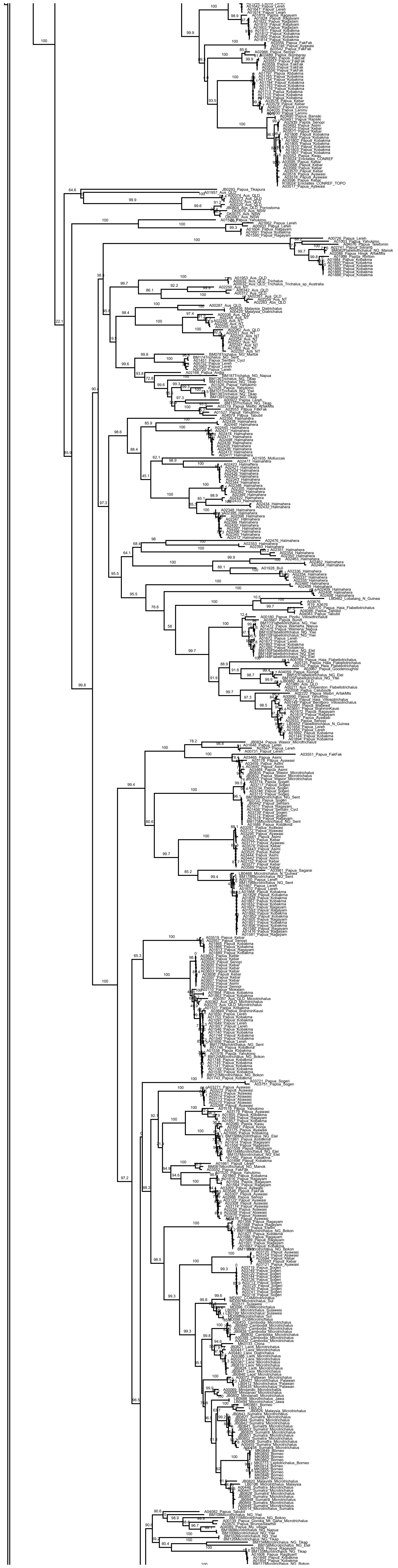

Figure S14. Maximum likelihood tree recovered by the analysis of the fulldataset (mitochondrial fragments). Depicted numbers above branches represent SH-alrt support values. The figure continues on the next page.

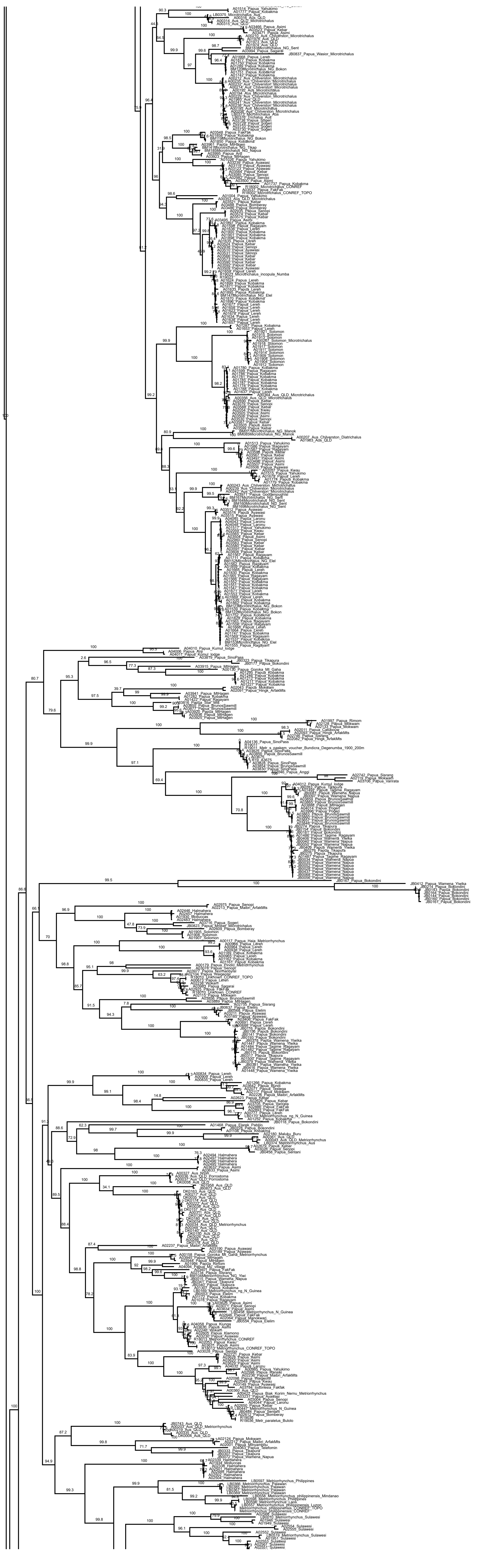

Figure S14. Maximum likelihood tree recovered by the analysis of the fulldataset (mitochondrial fragments). Depicted numbers above branches represent SH-alrt support values. The figure continues on the next page.

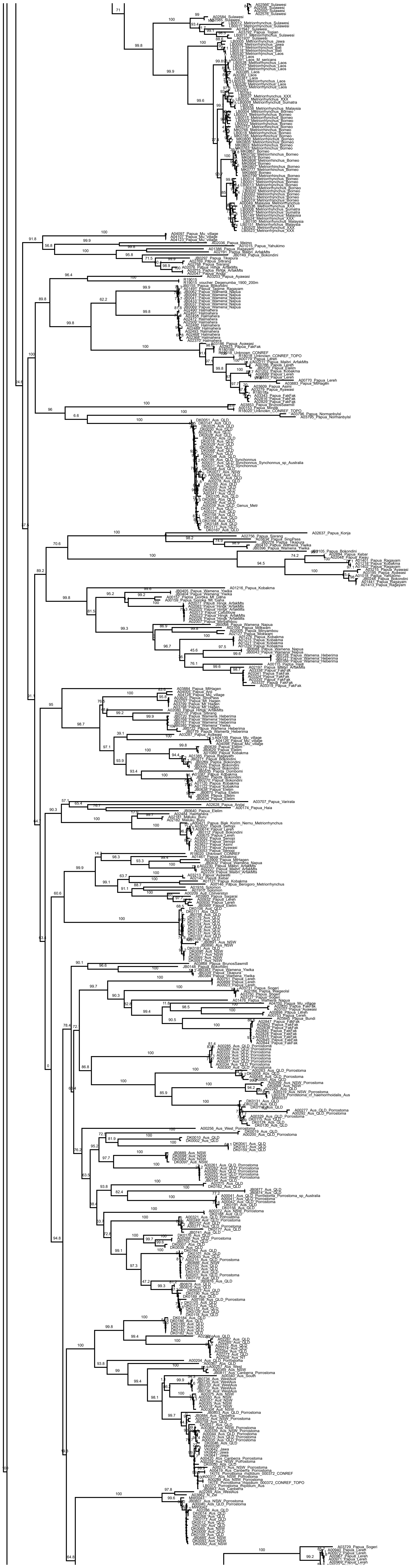

Figure S14. Maximum likelihood tree recovered by the analysis of the fulldataset (mitochondrial fragments). Depicted numbers above branches represent SH-alrt support values. The figure continues on the next page.

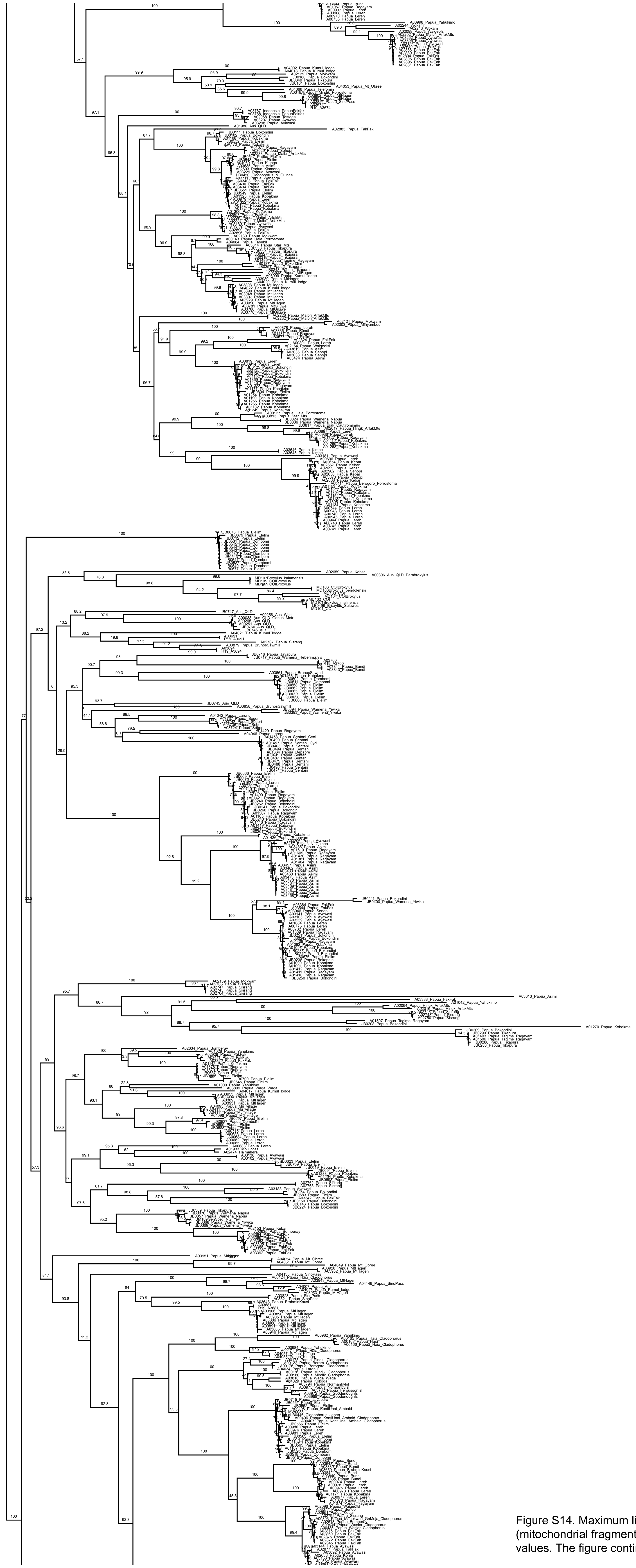

Figure S14. Maximum likelihood tree recovered by the analysis of the fulldataset (mitochondrial fragments). Depicted numbers above branches represent SH-alrt support values. The figure continues on the next page.

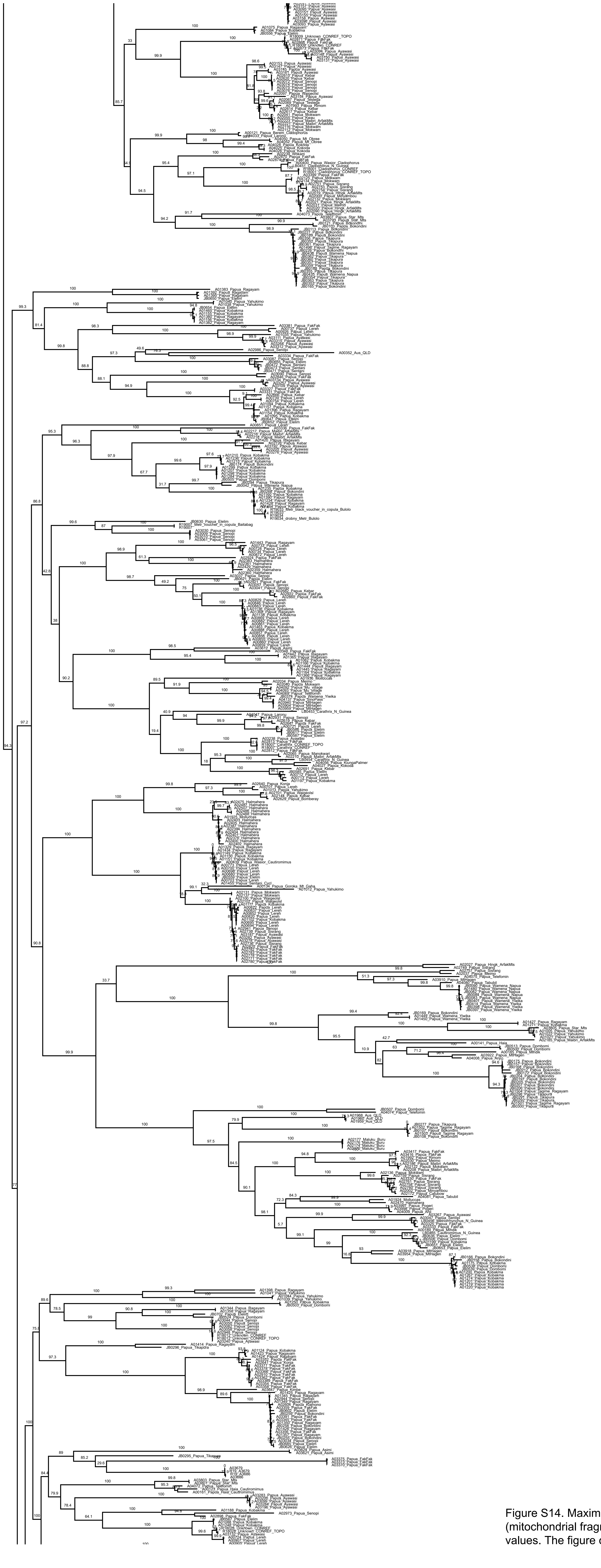

Figure S14. Maximum likelihood tree recovered by the analysis of the fulldataset (mitochondrial fragments). Depicted numbers above branches represent SH-alrt support values. The figure continues on the next page.

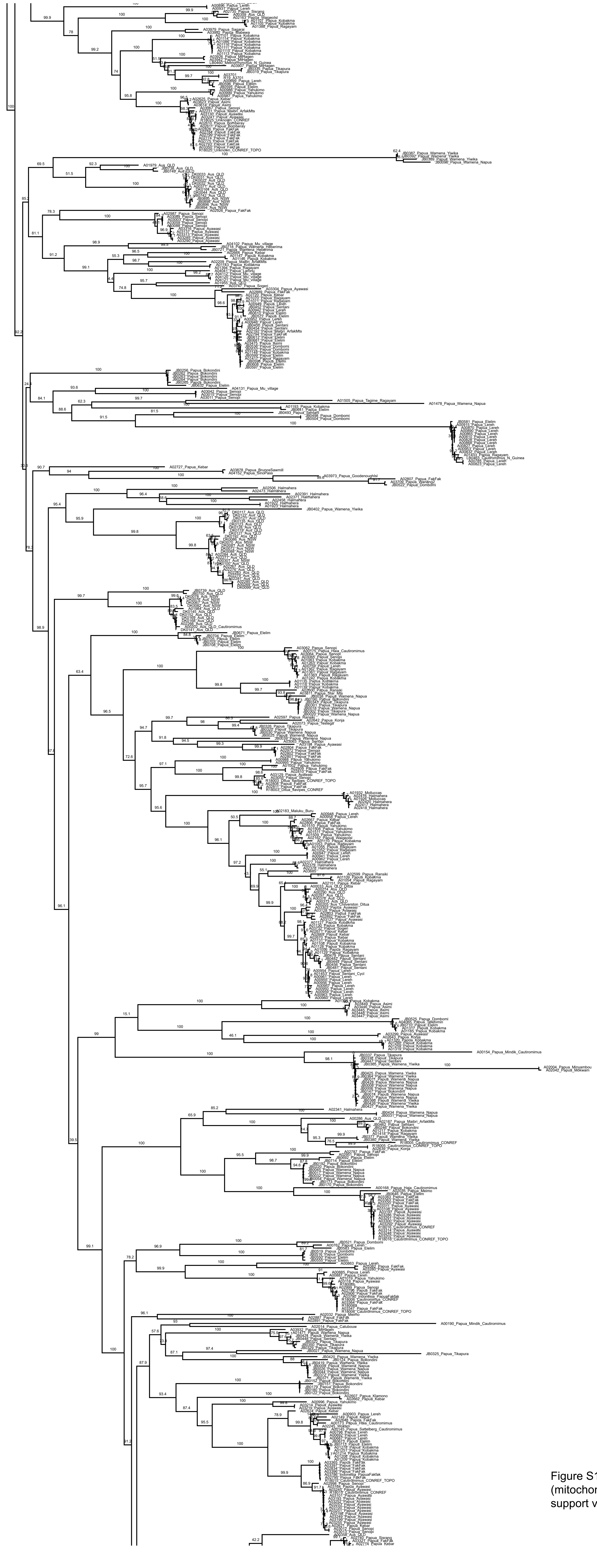

Figure S14. Maximum likelihood tree recovered by the analysis of the fulldataset (mitochondrial fragments). Depicted numbers above branches represent SH-alrt support values. The figure continues on the next page.

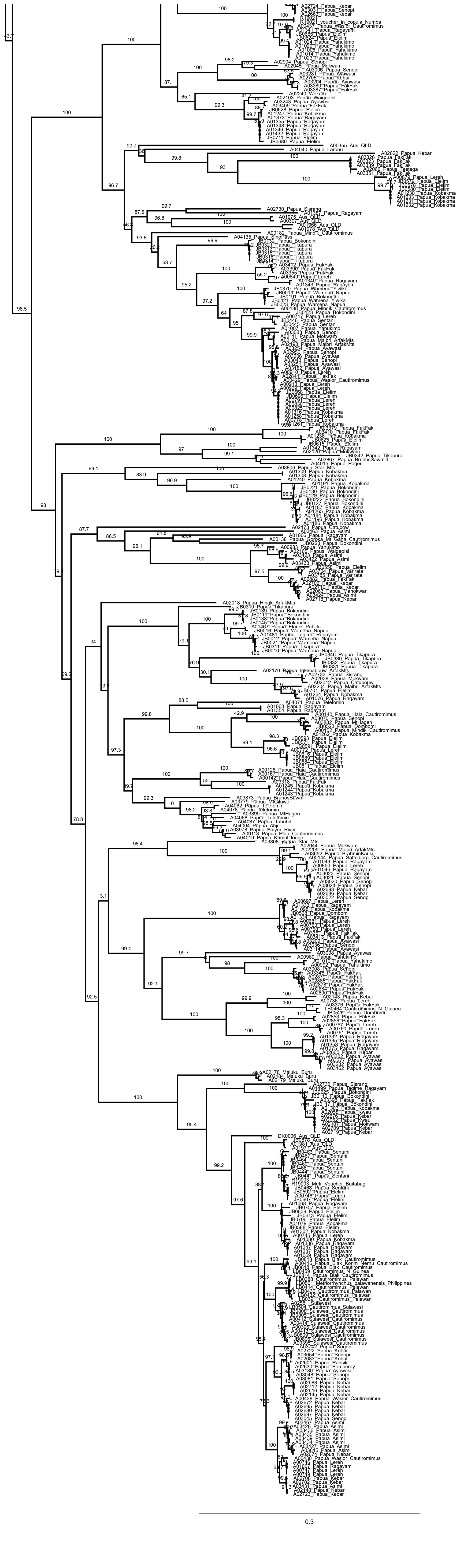

Figure S14. Maximum likelihood tree recovered by the analysis of the fulldataset (mitochondrial fragments). Depicted numbers above branches represent SH-alrt support values.

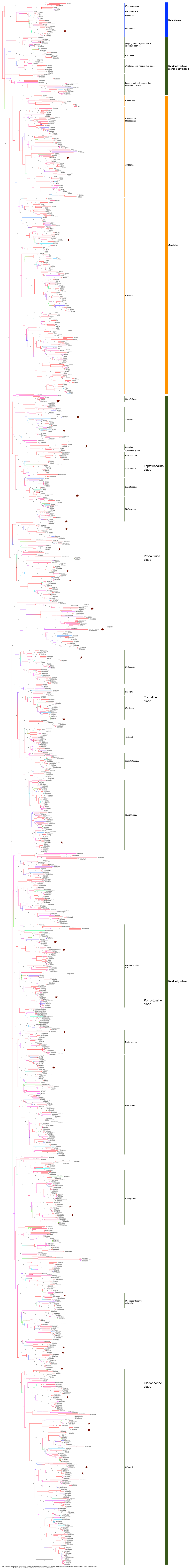

Figure S15. Maximum likelihood tree recovered by the analysis of the reduced dataset (98% similarity OTUs). Depicted numbers above braches represent SH-aLRT support values. The taxa with the constrained position in the tree are marked with red star.

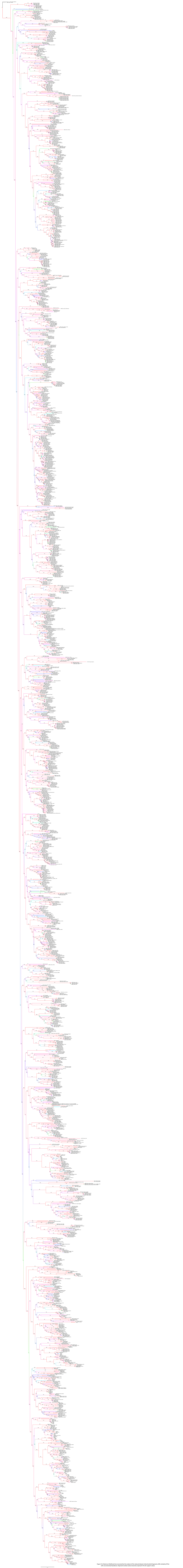

Figure S16. Maximum likelihood tree recovered by the analysis of the reduced dataset (mitochondrial fragments, 98% similarity OTUs) with unconstrained backbone. Depicted numbers above branches represent SH-alrt support values.

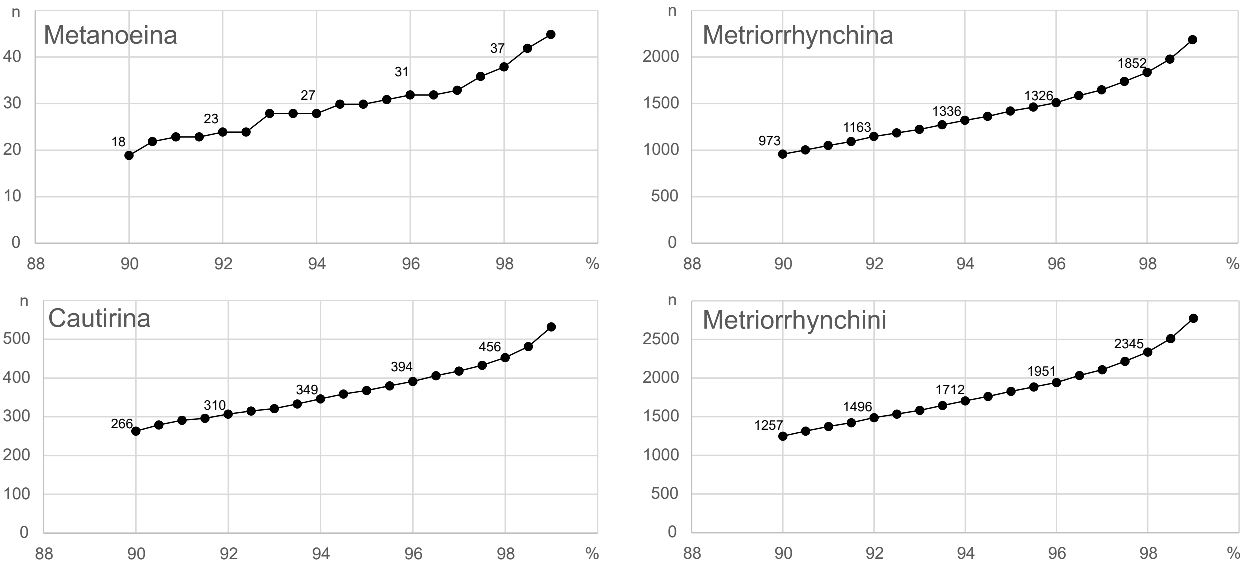

Figure S17. Tribal and subtribal mOTUs delimitation using CD-hit-est. Axis X represent the number of the operational taxonomic units, whereas axis Y shows the delimitation threshold.

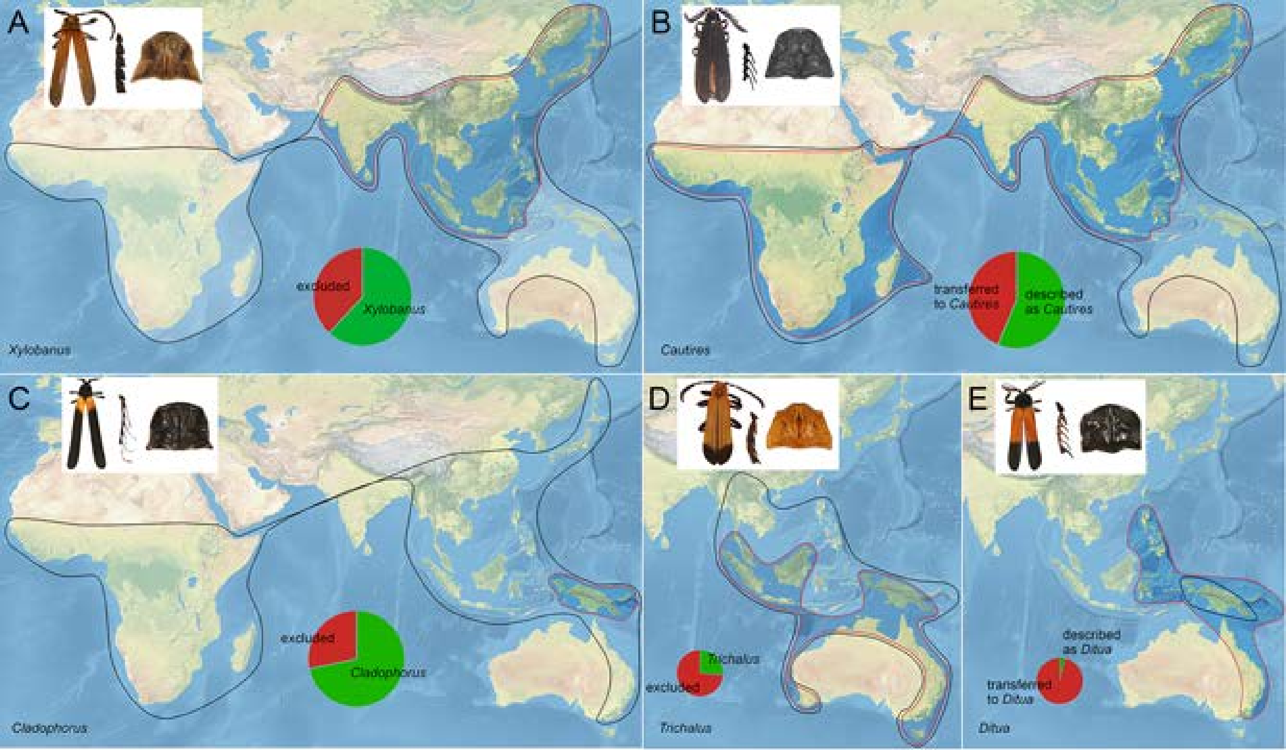

Figure S18. A Distribution of selected Metriorrhynchini genera (originally assumed and revised).
